# Interactors and effects of overexpressing YlxR/RpnM, a conserved RNA binding protein in cyanobacteria

**DOI:** 10.1101/2024.01.06.574455

**Authors:** Luisa Hemm, Anna Miucci, Matthias Riediger, Stefan Tholen, Alexander Kraus, Jens Georg, Oliver Schilling, Wolfgang R. Hess

## Abstract

Throughout the tree of life RNA-binding proteins play important roles, but they are poorly characterized in cyanobacteria. Structural prediction suggests an RNA-binding interface for the protein YlxR/Ssr1238 in the cyanobacterium *Synechocystis* 6803. Two pairs of cysteine residues are arranged as possibly coordinating an Fe-S cluster and appear widely conserved in the homologous proteins of other cyanobacteria. Overexpression of Ssr1238 for 24 h led to higher levels of RNase P RNA, tRNAs, and stress-related mRNAs. Co-immunoprecipitation of proteins followed by MS analysis and sequencing of UV crosslinked, co-immunoprecipitated RNA samples identified potential interaction partners of Ssr1238. The most enriched transcript was RNase P RNA, and RnpA, the protein component of RNase P, was among the most highly enriched proteins. A second highly enriched transcript derived from gene *ssl3177*, which encodes a central enzyme in cell wall remodeling during cell division. The data also showed a strong connection to the RNA maturation and modification system indicated by co-precipitation of RNA modifying enzymes, riboendonuclease E and enolase. Surprisingly, cyanophycin synthetase and urease were highly enriched as well. In conclusion, Ssr1238 specifically binds to two different transcripts and participates in the coordination of RNA maturation, translation, cell division, and aspects of nitrogen metabolism. Our results are consistent with recent findings that the *B. subtilis* YlxR protein functions as an RNase P modulator (RnpM), but suggest additional functionalities and extend its proposed role to the phylum cyanobacteria.

## Introduction

### RNA-binding proteins and cyanobacteria

RNA-binding proteins (RBPs) are crucial regulators of gene expression in all domains of life. Hundreds of previously unknown RBPs have recently been described from yeast to humans [1]. In bacteria, RBPs accomplish functions as divergent as selectively protecting specific RNAs from degradation or recruiting the RNA degradosome to interact with a particular mRNA for degradation. Bacterial RBPs regulate the initiation of translation, terminate transcription (Rho) and are involved in matchmaking between an sRNA and its target mRNA (RNA chaperones such as Hfq or ProQ) or scaffolding (for a recent review, see [2]).

Cyanobacteria are the only bacteria whose physiology is based on oxygenic photosynthesis. Therefore, they are physiologically and genetically distant from gram-positive and gram-negative model bacteria such as *Bacillus subtilis* and *Escherichia coli*. A well-established model for cyanobacteria is the unicellular *Synechocystis* sp. PCC 6803 (*Synechocystis* 6803). *Synechocystis* 6803 was the first photosynthetic organism for which the total genome sequence was obtained [3], and extensive datasets exist on the genome-wide regulation of gene expression [4] and active promoters [5], and on the suite of co-fractionating RNAs and proteins using gradient profiling by sequencing (Grad-seq) [6]. Several sRNAs and riboswitches have been described in *Synechocystis* 6803, which regulate the responses to high light intensity [7], low iron [8], or are involved in the control of mixotrophy [9], or nitrogen metabolism [10,11]. However, no RNA chaperones or RBPs that mediate sRNA/mRNA duplex formation have been described in *Synechocystis* 6803 or any other cyanobacteria. Although a structural homolog of Hfq exists in most cyanobacteria, residues that are known to interact with RNA are not well conserved [12], and RNA-binding activity of Ssr3341, the Hfq homolog in *Synechocystis* 6803 [13], was not detected [14]. In contrast, RNA binding was verified *in vitro* [15] for some proteins with an RNA-recognition motif (RRM), which are encoded by a small gene family in all cyanobacteria [16–21]. Some RRM domain-containing RBPs (in *Synechocystis* 6803 Rbp2 and Rbp3) were found to be involved in the targeting of certain mRNAs to the thylakoid membrane system [22]. Overall, these observations suggest that most aspects concerning the regulation by RBPs and their overall functions still await discovery in cyanobacteria.

### The Ssr1238/Slr0743a protein - an uncharacterized YlxR homolog

The *Synechocystis* 6803 protein Ssr1238, also referred to as Slr0743a in some genome annotations, is an 84 amino acid basic protein of the YlxR/ DUF448 family with a calculated isoelectric point of 11.09. In our previous work, it was predicted as a potential RBP based on Grad-seq results and a high support vector machine score for the prediction of RBPs from their amino acid sequences [6]. Ssr1238 is more distant to the YlxR proteins in *Bacillus subtilis* and *Streptococcus pneumoniae* or *Clostridioides difficile* with 24%, 24%, or 29% identical amino acids (**Figure 1A**).

**Figure 1.**
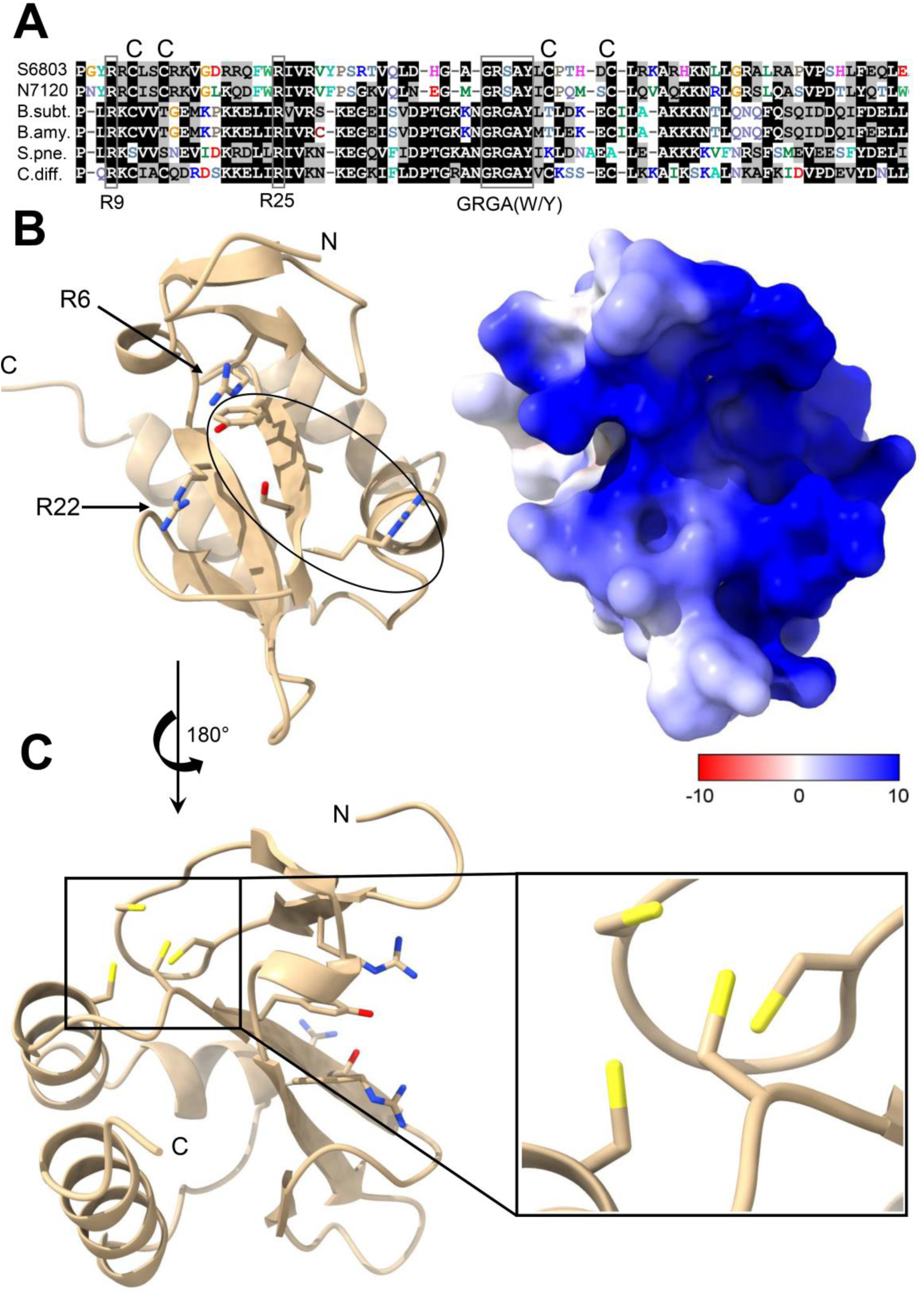
Conservation of potential RNA-binding residues and two pairs of cysteine residues in YlxR/Ssr1238. **A.** Multiple sequence alignment of YlxR homologs from different species. Only the stretch matching to residues 3 to 77 of the 84 amino acid protein in *Synechocystis* 6803 is shown. The homologs are from the two cyanobacteria *Synechocystis* (S6803; Ssr1238) and *Nostoc* sp. PCC 7120 (N7120; Asr2830), compared to sequences from *Bacillus subtilis* subsp. subtilis str. 168 (B.subt.; NP_389543), *Bacillus amyloliquefaciens* (B.amy.; WP_158185884), *Streptococcus pneumoniae* (S.pne.; BEL22271) and *Clostridioides difficile* (C.diff.; CEJ97888). Identical and at least 50% conserved residues are shaded black. Residues R9, R25 and the GRGA(W/Y) motif within the *S. pneumonia* homolog are indicated as these have been discussed to be involved in RNA binding [23]. Capital Cs on top of the alignment indicate two pairs of cysteine residues shared between Ssr1238 and homologs in other cyanobacteria and in *C. difficile*. These cysteine pairs are conserved among cyanobacterial homologs (***SI Appendix* Dataset 1**). **B.** Left panel: The structure of *Synechocystis* Ssr1238/YlxR predicted by AlphaFold [29,30] with the conserved residues R6 and R22 (corresponding to R9 and R25 of the *S. pneumonia* homolog) and the GRSAY motif (circled). N and C termini are indicated. Right panel: Electrostatic surface representation of YlxR/Ssr1238 highlighting the electropositive (blue) RNA-binding surface. **C.** Modeling of the YlxR/Ssr1238 structure suggests an arrangement of the cysteine residues (yellow) compatible with binding of an Fe-S cluster. The four well-conserved cysteine residues are boxed and enlarged to the right.

Initial evidence for YlxR proteins as RBPs was inferred from the structural analysis of the homolog in *S. pneumoniae* [23] and the in-gradient distribution of the *C. difficile* protein [24]. The *B. subtilis* YlxR protein has been reported as a nucleoid-associated protein that regulates nearly 400 genes [25–27], and more recently to bind to the RNA subunit of the ribonuclease P and modulate its activity [28]. Consequently, the *B. subtilis* YlxR protein has been renamed as RNase P modulator (RnpM) [28].

However, residues K11, V13, T15, and D53, which have been shown to be critical for RNA binding in *Bacillus* RnpM, are not conserved in the *Synechocystis* 6803 protein Ssr1238. In addition, two pairs of cysteine residues are present in Ssr1238, but are lacking in RnpM and the homologs in other *Bacillus* and *Streptococcus* strains (**Figure 1A**). Therefore, the function of cyanobacterial YlxR homologs is not clear.

To gain insight into the possible functions of YlxR homologs in cyanobacteria, we have identified RNA molecules and proteins interacting with the *Synechocystis* 6803 Ssr1238/YlxR protein *in vivo*. We report its interaction with both the RNase P RNA and protein subunits and at least one unrelated transcript. Co-immunoprecipitation (co-IP) experiments enriched ribosomal proteins, several enzymes related to RNA modification and turnover, and, surprisingly, nitrogen metabolism. Ssr1238/YlxR overexpression had a positive effect on the accumulation level of all 44 tRNAs and stimulated the expression of a small number of stress-related mRNAs.

## Results

### YlxR/Ssr1238 homologs in cyanobacteria carry two pairs of widely conserved cysteine residues

The *ssr1238* gene is located downstream of the *rimP* and *nusA* genes, encoding the ribosome maturation factor RimP and the transcription termination factor NusA, and upstream of the infB gene encoding the translation initiation factor InfB (arrangement *rimP-nusA-ssr1238-infB*) [6], thus suggesting a relationship to transcription and translation. Moreover, cyanobacterial genomes are widely syntenic around the *ssr1238* locus and strikingly similar arrangements exist in Firmicutes and Actinobacteria indicating a conserved common functional context of these genes.

A major difference compared to the *Bacillus* RnpM protein are two pairs of cysteine residues present in YlxR/Ssr1238 (C8/C11 and C45/C50; **Figure 1A**). To evaluate the conservation of these residues in cyanobacterial homologs, we searched the NCBI database with the YlxR/Ssr1238 sequence by blastP, yielding 1,062 homologs. Homologs were also detected in the genomes of diatom-associated symbionts *Calothrix rhizosoleniae* (GenBank accession WP_088239620.1) and *Richelia intracellularis* (WP_008232964.1), of UCYN-A *Candidatus Atelocyanobacterium thalassa* endosymbionts (WP_040054730.1, WP_012954106.1), and in the endosymbiont chromatophore genomes of photosynthetic *Paulinella longichromatophora* (AUG32252.1), *Paulinella chromatophora* (YP_002048766.1), and *Paulinella micropora* (APP87879.1, YP_009530348.1). These data suggest that homologs of *ylxR/ssr1238* belong to the core genome of cyanobacteria. With the exception of a few outliers which appear N-terminally truncated, the alignment of these proteins indicated the total conservation of both cysteine pairs in homologs throughout the cyanobacterial phylum (***SI Appendix* Dataset 1**).

Another difference to the *Bacillus* protein is that residues K11, V13, T15, and D53, which are critical for RNA binding in RnpM [28], appear to have been replaced in Ssr1238/YlxR (residues R7, L9, C11 and T47, **Figure 1A**). However, residues corresponding to R9, R19 and the GRGA(W/Y motif of the *S. pneumoniae* YlxR are well conserved. These were previously discussed as involved in RNA binding based on structural analysis of the *S. pneumoniae* YlxR protein [23] and appear widely conserved also in the homologs of other cyanobacteria (**Figure 1A, *SI Appendix* Dataset 1**). Prediction of the Ssr1238/YlxR structure by AlphaFold [29,30] revealed that these residues are part of an highly electropositive cleft (**Figure 1B**), a likely RNA-binding interface.

The structural analysis furthermore predicted the conserved cysteine pairs to be arranged spatially close and facing outward on the other side of the protein (**Figure 1C**). The arrangement of these four cysteine residues appears compatible with binding of an Fe-S cluster, indicating a major difference to the homologs in *Bacillus* and *Streptococcus*.

### UV Crosslinking and sequencing provides evidence of YlxR/Ssr1238 interacting with two distinct transcripts

To determine potential RNA interaction partners of YlxR/Ssr1238, a pVZ322 plasmid was constructed in which a sequence encoding a triple FLAG epitope tag was fused to the 3’ end of the *ssr1238* reading frame controlled by the P_rha_ promoter, which can be activated by the addition of rhamnose [31]. Conjugation of this plasmid into *Synechocystis* 6803 yielded strain P*_rha_*_*ssr1238-*3×FLAG. As control, a construct encoding a tagged superfolder green fluorescent protein (sfGFP) was engineered, yielding strain P*_rha_*_*sfGFP-*3×FLAG.

The strains were cultivated in triplicate under identical conditions and cells were subjected to the crosslinking, immunoprecipitation and RNA sequencing procedure (CLIP-Seq). After crosslinking with UV light, the Ssr1238 and sfGFP proteins were pulled down using magnetic beads binding to the FLAG epitope tag that was fused to the C-terminal ends of the proteins. Purification of sfGFP and Ssr1238 was controlled by SDS-PAGE and western blotting, showing signals at the expected molecular masses of ∼30 kDa (sfGFP_3×FLAG) and ∼13 kDa (Ssr1238_3×FLAG; ***SI Appendix* Fig. S1**).

After protein pull-down, crosslinked RNA was extracted from the beads. As control for total transcript levels, total RNA of both strains was extracted in triplicate, yielding 12 samples in total. We obtained ∼160 ± 60 ng of UV-clipped RNA from the Ssr1238_3×FLAG purification and 15 ± 4 ng of RNA for the sfGFP_3×FLAG negative control. For total RNA control samples, 4 to 24 μg of RNA was obtained. The ten-fold higher yield from the Ssr1238_3×FLAG purification compared to the sfGFP_3×FLAG sample indicated that RNA was bound by the former but less so by the latter. Comparison of the extracted RNA samples on a fragment analyzer revealed the enrichment of an RNA species of ∼450 nt in the CLIP RNA of P*_rha_*_*ssr1238*-3×FLAG, but not in the RNA extracted from the UV-treated P*_rha_*_*sfGFP-*3×FLAG strain or in either of the unclipped controls (***SI Appendix* Fig. S2**).

Next, cDNA libraries were prepared and sequenced on an Illumina NextSeq 500 system with 75 bp read length providing ∼10,000,000 reads per sample.

The galaxy system [32] was used to analyze the sequencing data. The workflow is illustrated in ***SI Appendix* Fig. S3.**

To identify crosslinked RNAs that specifically interacted with the Ssr1238_3×FLAG protein (indicated as XL-Ssr1238), the CLIP seq data for the Ssr1238_3×FLAG expression strain were related to the corresponding data obtained for the strain expressing sfGFP_3×FLAG (indicated as XL-sfGFP). To obtain quantitative data, log_2_ fold changes (log_2_ FC) were calculated and plotted over the adjusted *p*-values (**Figure 2A**). In the volcano plot, the transcript fragments with the highest log_2_ FC compared to the sfGFP control are labelled. With the lowest observed *p*-value and a log_2_FC of 5.969, RNase P RNA was the most-enriched transcript. The coverage pattern along the *rnpB* gene, from where RNase P RNA originates, showed contiguous coverage, with three distinct peaks (**Figure 2B**). In contrast, the coverage of *ssr1238* mRNA was almost identical in XL-Ssr1238 and in the cDNA analysis of total RNA with overexpressed *ssr1238* (OE-Ssr1238), and the coverage for both samples was much higher compared than that of the two sfGFP samples (XL-sfGFP and OE-sfGFP), consistent with its overexpression (**Figure 2C**). Therefore, *ssr1238* mRNA was not specifically enriched owing to crosslinking with YfrX/Ssr1238.

**Figure 2.**
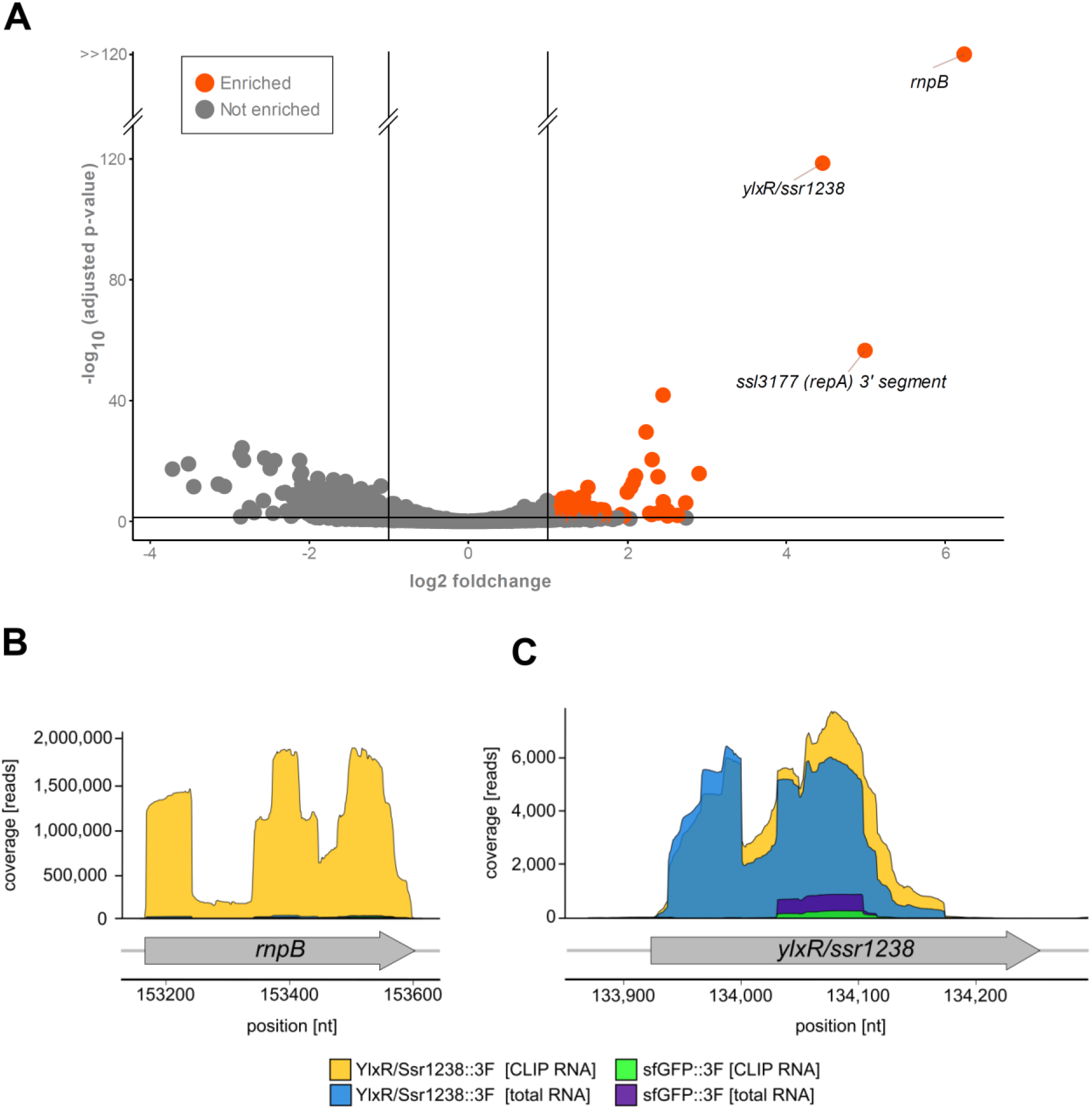
Transcripts enriched with YlxR/Ssr1238 after UV crosslinking in *Synechocystis.* **A.** Volcano plot showing transcripts enriched in the Ssr1238*_*3×FLAG CLIP RNA samples compared to the sfGFP*_*3×FLAG CLIP RNA samples (n=3). The - log_10_ adjusted *p*-values are plotted against the log_2_ fold changes. Red dots indicate enriched transcripts and gray dots indicate non-enriched transcripts. Gene IDs are given for the three most enriched transcripts (*rnpB*, RNase P RNA; *ylxR/ssr1238*, mRNA of the overexpressed gene; *ssl3177 (repA*), 3’ segment of coding sequence and 3’ UTR; see Figure 3 for further details). The full list is provided in ***SI Appendix* Table S3**. **B.** Read coverage for the 4 different samples at the *rnpB* locus. **C.** Read coverage at the *ylxR/ssr1238* gene. Read coverages are given along the y-axis, genomic positions (GenBank accession NC_000911.1) along the x-axis. The different samples compared are color-coded as indicated in the legend (CLIP RNA, crosslinked RNA; total RNA, RNA-seq data from cultures overexpressing *ylxR/ssr1238* or *sfGFP*), n=3.

The list of all transcripts enriched in XL-Ssr1238 RNA compared to XL-sfGFP is given in ***SI Appendix* Table S3** and the list of transcripts enriched in XL-Ssr1238 RNA compared to OE-Ssr1238 in ***SI Appendix* Table S4**. In both comparisons we chose a minimum log_2_ fold change (log_2_FC) ≥1 and an adjusted *p*-value (padj) ≤0.05.

In addition to RNase P RNA, we noticed another particularly highly enriched transcript segment and several other slightly enriched peaks in the cDNA libraries generated from RNA crosslinked to YlxR/Ssr1238. The highly enriched transcript segment derived from the 3’ region of gene *ssl3177* with a log_2_FC +4.99 in the comparison XL- Ssr1238 / XL-sfGFP and a log_2_FC of +4.01 in the comparison XL-Ssr1238 / OE- Ssr1238. The identified transcript segment interacting with YlxR/Ssr1238 was approximately 45 nt in length (**Figure 3A**). The segment contains a potentially strong helical structure of 37 nt (minimum free energy of -13.10 kcal/mol) and encompasses the last two codons and the *ssl3177* stop codon, together with the first 27 nt of the 3’UTR (**Figure 3B**). Therefore, it cannot constitute a classical 3’ UTR-located Rho-independent terminator. Comparison of sequences from homologous loci in other *Synechocystis* strains revealed strong sequence and structure conservation (**Figure 3C**). These strains differ by several hundred genes from *Synechocystis* 6803 [33–35]. For instance, *Synechocystis* 6803 and *Synechocystis* 6714 share 2838 protein-coding genes, while 845 genes are unique for *Synechocystis* 6803 and 895 genes for *Synechocystis* 6714 [34]. Therefore, the conservation of the RNA fragment crosslinked to YlxR/Ssr1238 indicates that it has a common relevant function in the different *Synechocystis* strains.

**Figure 3.**
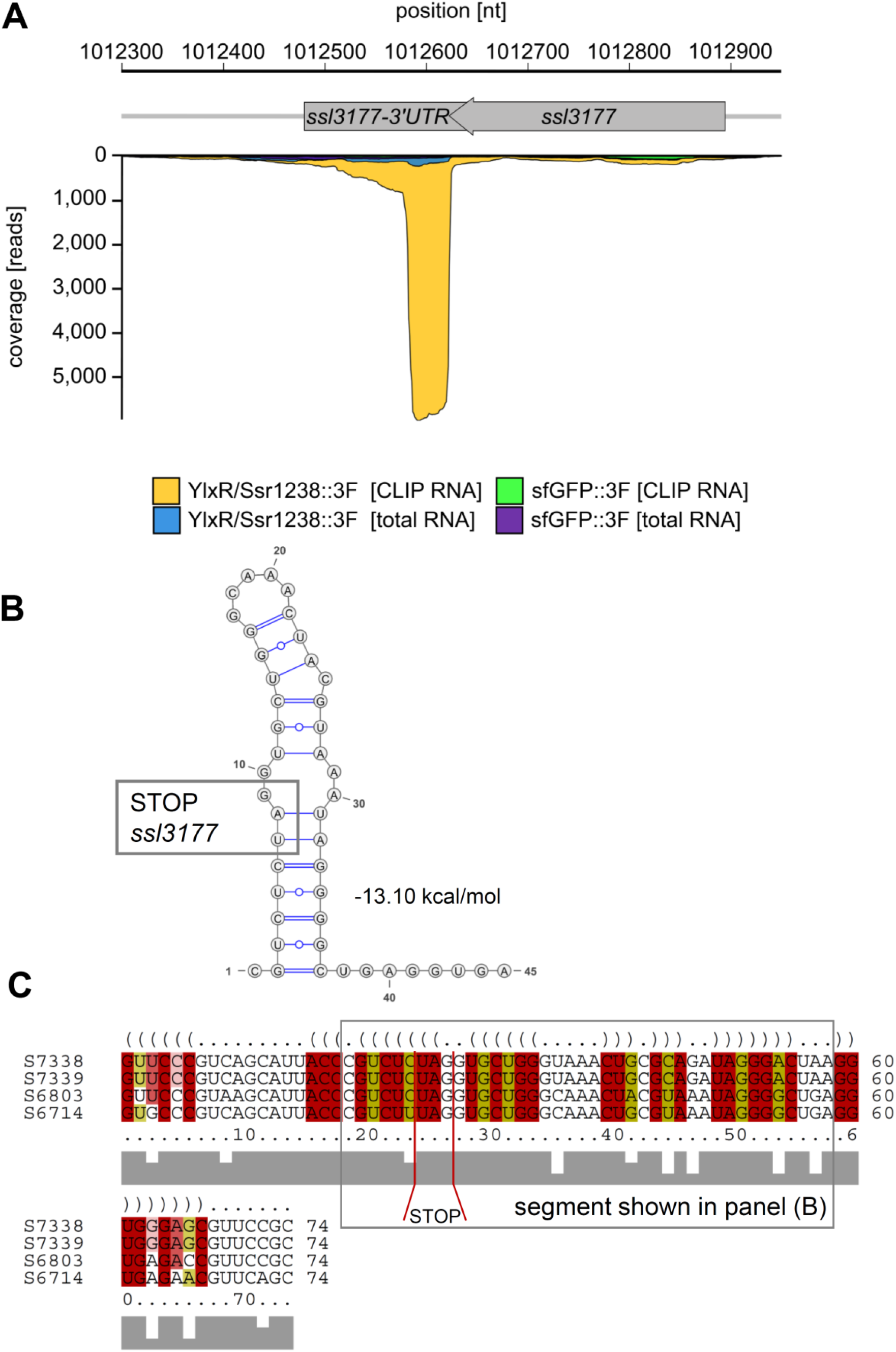
Interaction of YlxR/Ssr1238 with an *ssl3177* mRNA 3’ end-derived transcript segment. **A.** Crosslinked transcript segment (grey, XL YlxR/Ssr1438, all colors as in Figure 2B and C). **B.** A stem-loop secondary structure within the enriched transcript segment as predicted by RNAfold [84] accessed at http://rna.tbi.univie.ac.at//cgi-bin/RNAWebSuite/RNAfold.cgi. The *ssl3177* stop codon and the calculated minimum folding energy are indicated. **C.** LocaRNA [85] alignment of the segment shown in panel (B) including some up- and downstream positions with homologous sequences from *Synechocystis* spp. PCC 6714, PCC 7338 and PCC 7339. The segment from panel (B) is boxed and the stop codon indicated. The webserver at https://rna.informatik.uni-freiburg.de/LocARNA/Input.jsp was used.

However, annotation of *ssl3177* is conflicting. In most *Synechocystis* 6803 genome annotations it is a 90-codon reading frame encoding a protein with a truncated (partial) rare lipoprotein A domain. The gene *ssl3177* has remained uncharacterized experimentally thus far, but it has been found to be highly regulated in the context of carbon metabolism manipulation [36], iron starvation [37], osmotic stress [38], and in response to the depletion of the FtsH1/3 proteolytic complex [39], pointing to a role of Ssl3177 in several different acclimation processes. Indeed, comparison of Ssl3177 to likely homologs (***SI Appendix* Fig. S4**) revealed that the annotation of *ssl3177* should be corrected. The re-annotated coding sequence begins with an alternative TTG start codon (positions 1,012,946 to 1,012,948 on the reverse strand; GenBank accession NC_000911.1) and encodes the 110 amino acid rare lipoprotein A (gene *repA;* note that *Synechocystis* 6803 has a second rare lipoprotein A gene, called *rlpA, slr0423*), a central enzyme involved in cell wall remodeling during cell division [40]. This function is also consistent with the genomic location of *repA*/*ssl3177* downstream of *ftsZ*.

We conclude that YlxR/Ssr1238 binds to at least two distinct transcripts of *Synechocystis* 6803 *in vivo*. One of these transcripts is RNase P RNA and the other is an RNA fragment derived from the 3’ end of the *ssl3177* gene encoding a rare lipoprotein A homolog.

### Overexpression of YlxR/Ssr1238 leads to transcriptome changes

The 24 h induction period provided an opportunity for insight into possible consequences of increased YlxR/Ssr1238 expression. We calculated log_2_FC for the comparison of transcript levels between OE-Ssr1238 and OE-sfGFP (**Figure 4**). This comparison indicated that the transcript levels of 68 genes were upregulated and another 57 genes were downregulated when applying a a log_2_FC ≥1 and a *p*_adj_ ≤0.05 (***SI Appendix* Table S5**).

**Figure 4.**
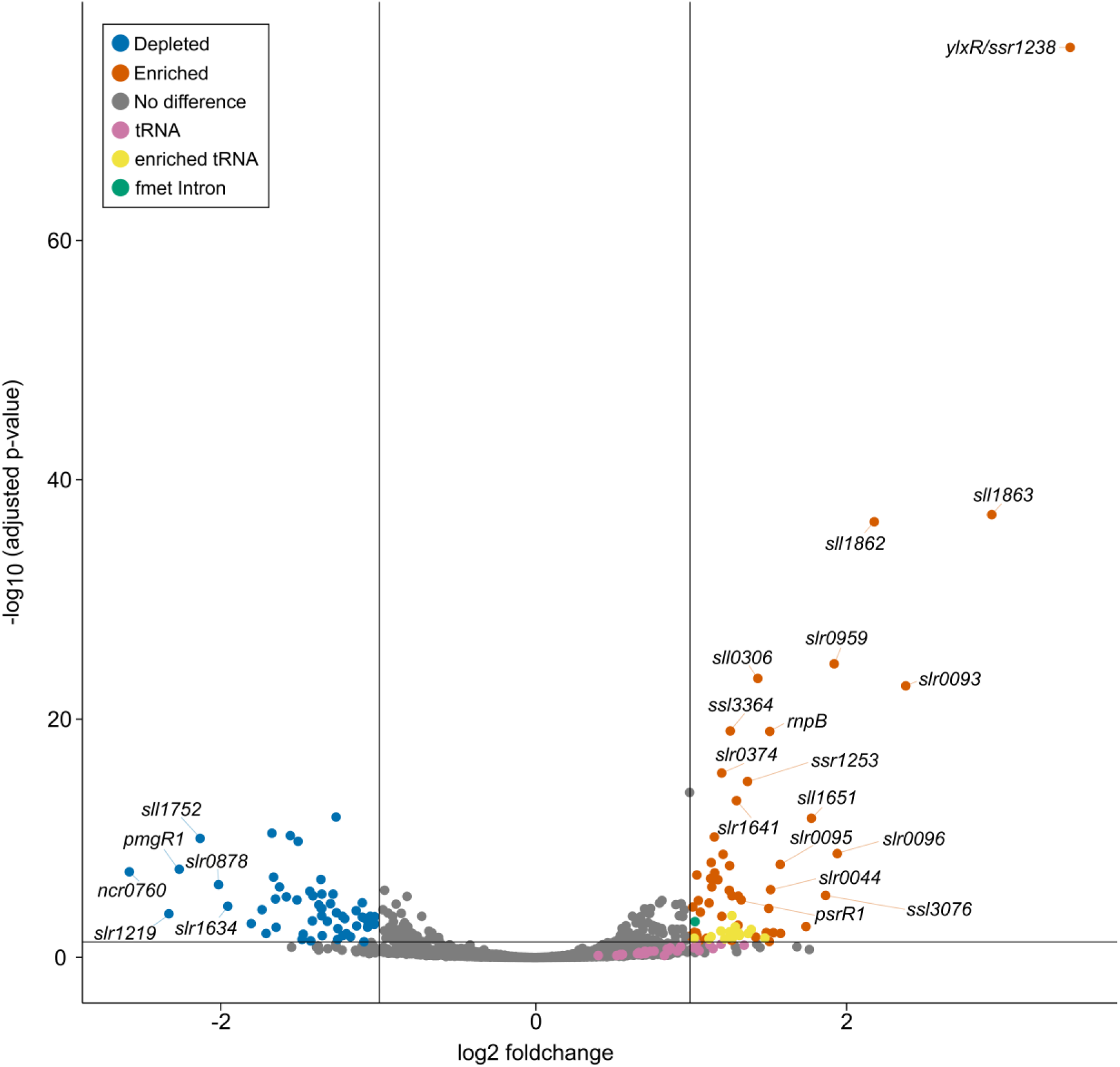
RNA-seq analysis of up- and down-regulated genes upon overexpression of YlxR/Ssr1238. Volcano plot showing transcripts upregulated in OE-Ssr1238 compared to OE-sfGFP. The red dots indicate upregulated mRNAs and non-coding RNAs (*rnpB*, RNase P RNA; *psrR1*, photosynthesis regulatory RNA 1), yellow and magenta dots indicate upregulated tRNAs and a green dot the tRNA_i_^fmet^ intron fragment. Blue dots indicate down-regulated transcripts (*pmgR1*, photomixotrophic growth RNA 1). See ***SI Appendix* Table S5** for full details.

Consistent with its intentional overexpression, *ssr1238* (log_2_FC of +3.446) was the most upregulated gene, followed by *sll1863*, *sll1862*, and *slr0093* encoding one of the seven DnaJ homologs, with log_2_FCs between +2.1 and 2.9. These genes have often been implicated in stress responses [41–43], and Sll1863, a protein of unknown function, was previously identified as the most induced protein under salt stress [44]. Along these lines, we also noticed an upregulation of *ggpR/ssl3076* (log_2_FC of +1.872), the regulator mediating the salt-induced activation of glucosylglycerol synthesis in *Synechocystis* 6803 [45]. To test for possible phenotypic effects, we spotted aliquots of cultures overexpressing *ssr1238* under 6 different conditions (***SI Appendix* Fig. S5**). Although differences in growth were almost undetectable under standard growth conditions or with the addition of 10 mM glucose, the overexpressing strains had a clear growth disadvantage under cold stress and at an elevated level of illumination. Finally, *rnpB*, the gene from which the RNA component of RNase P is transcribed, was found to be upregulated (log_2_FC of +1.512), indicating a possible stabilizing effect of YlxR/Ssr1238 overexpression. Intriguingly, the largest group of upregulated genes in the Ssr1238 overexpressor encompassed tRNA genes. In fact, we found that all 44 tRNA species present in *Synechocystis* 6803 were slightly more abundant in the OE- Ssr1238 strain, 16 of which had a significant log_2_FC ≥1. One of these tRNA genes contains a self-splicing group I intron in *Synechocystis* 6803 (***SI Appendix* Fig. S6**). The intron is located within the gene for the N-formylmethionine initiator-tRNA (tRNA_i_^fmet^). The complex gene arrangement in this genomic region is often misannotated, although this intron was first described 30 years ago [46]. Relative to the tRNA structure, the intron is inserted within the anticodon loop. At the DNA level, the intron contains the protein-coding gene *slr0915* encoding the DNA double-strand homing endonuclease I-Ssp6803I [47]. Furthermore, the tRNA_i_^fmet^ precursor transcript extends into the neighboring gene *slr0917* (*bioF*) encoding 8-amino-7-oxononanoate synthase. The entire segment, including the tRNA exons 1 and 2, the intron containing *slr0915,* and the downstream gene slr0917, is transcribed from a single transcription

start site 4 nt upstream of tRNA exon 1, thus constituting a single contiguous transcription unit (TU) of 1,816 nt, classified as TU2935 in the genome-wide identification of transcription start sites [5] (***SI Appendix* Fig. S6**). Here, we found log_2_FCs for the different parts of this transcript as follows: tRNA-6803t34-gene (tRNA_i_^fmet^), +1.269; tRNA_i_^fmet^-intron-*bioF* precursor, +1.027 and *bioF* alone, +1.025. Moreover, we noticed a specific slightly enriched intron segment in the crosslinked RNA (see ***SI Appendix* Fig. S6**). This segment is located downstream of gene *slr0915* and comprises parts of intron domains P7, P8 and P9, including the pseudoknot formed between P8 and P3.

We conclude that 24 h of YlxR/Ssr1238 overexpression resulted in some, although not dramatic, changes in the transcriptome. The observed upregulation of both RNase P and tRNAs suggests at a possible stabilizing effect of YlxR/Ssr1238 upon binding, which likely contributes secondarily to the higher tRNA levels. In addition, *ssr1238* overexpression led to further changes in the composition of the transcriptome and affected vitality, which was more pronounced under certain stress conditions.

### YlxR/Ssr1238 co-immunoprecipitates with proteins involved in RNA metabolism and nitrogen metabolism

To gain insight into the cellular processes that involve YlxR/Ssr1238, co-IP experiments were performed with strain P*_rha_*_*ssr1238-*3×FLAG. After 24 h induction, YlxR/Ssr1238-3×FLAG and bound interaction partners were immunoprecipitated from the lysate using anti-FLAG magnetic beads. In parallel, lysates from P*_rha_*_*sfGFP-*3×FLAG cultures were purified for control. Both experiments were performed in triplicate.

Western blots were used for quality control of the protein pull-down for both proteins (***SI Appendix* Fig. S7**). Signals of ∼30 kDa in the crude extract, flow-through and elution fractions were obtained for the sfGFP-3×FLAG samples, while signals of ∼13 kDa were obtained for the YlxR/Ssr1238-3×FLAG replicates. These signals were consistent with the expected molecular masses. Thus, the proteins were not lost during the preparation. The eluate fractions were analyzed by LC-MS/MS. A total of 1,010 proteins were detected in one or in both samples.

Statistical analysis indicated that a total of 234 proteins were enriched (152 significantly enriched) in the co-IP with YlxR/Ssr1238 compared to the sfGFP control (**Figure 5**). The complete list of detected proteins is shown in ***SI Appendix* Table S6**.

**Figure 5.**
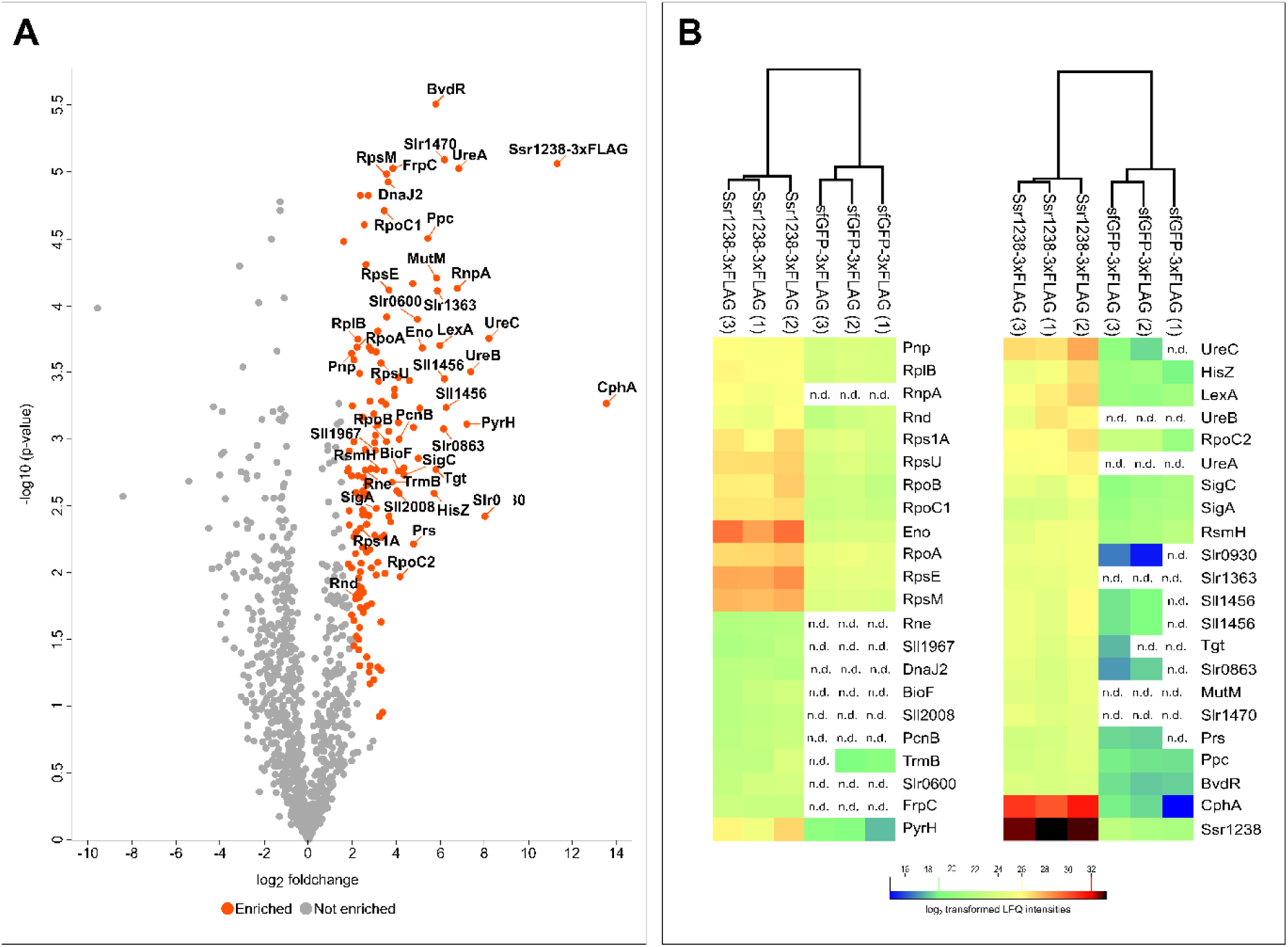
Co-IP experiments for the identification of protein interactors of YlxR/Ssr1238. **A.** Volcano plot based on a two-sample t test of enriched proteins using a false discovery rate (FDR) of 0.01 and a coefficient for variance minimization [86] s0 of 2. **B.** Hierarchical clustering of the most abundant proteins detected by mass spectrometry. The two most-enriched proteins are YlxR/Ssr1238 and cyanophycin synthetase (CphA [49]). For the complete list of detected proteins, see ***SI Appendix* Table S6,** for details of the data analyses ***SI Appendix* Tables S7 & S8**.

Despite its small size of 84 amino acids, 4 unique peptides were detected for YlxR/Ssr1238 and the associated log_2_FC of 10.99 indicated that the IP worked successfully. Among the 152 significantly enriched proteins, 24 were found exclusively in the YlxR/Ssr1238 co-IP and 190 were significantly enriched compared to their LFQ intensities in the sfGFP co-IP.

**Table 1** shows the 36 proteins that were enriched with a log_2_FC ≥4 in the YlxR/Ssr1238 samples. Of these, several are related to RNA metabolism, RnpA, the protein component of ribonuclease P, the queuine tRNA-ribosyltransferase Slr0713, Sll1253, a 924 amino acids protein with a nanoRNase/pAp phosphatase domain and a separate A-adding tRNA nucleotidyltransferase domain, and finally enolase, an enzyme that is considered to be a component of the RNA degradosome [48].

**Table 1.**
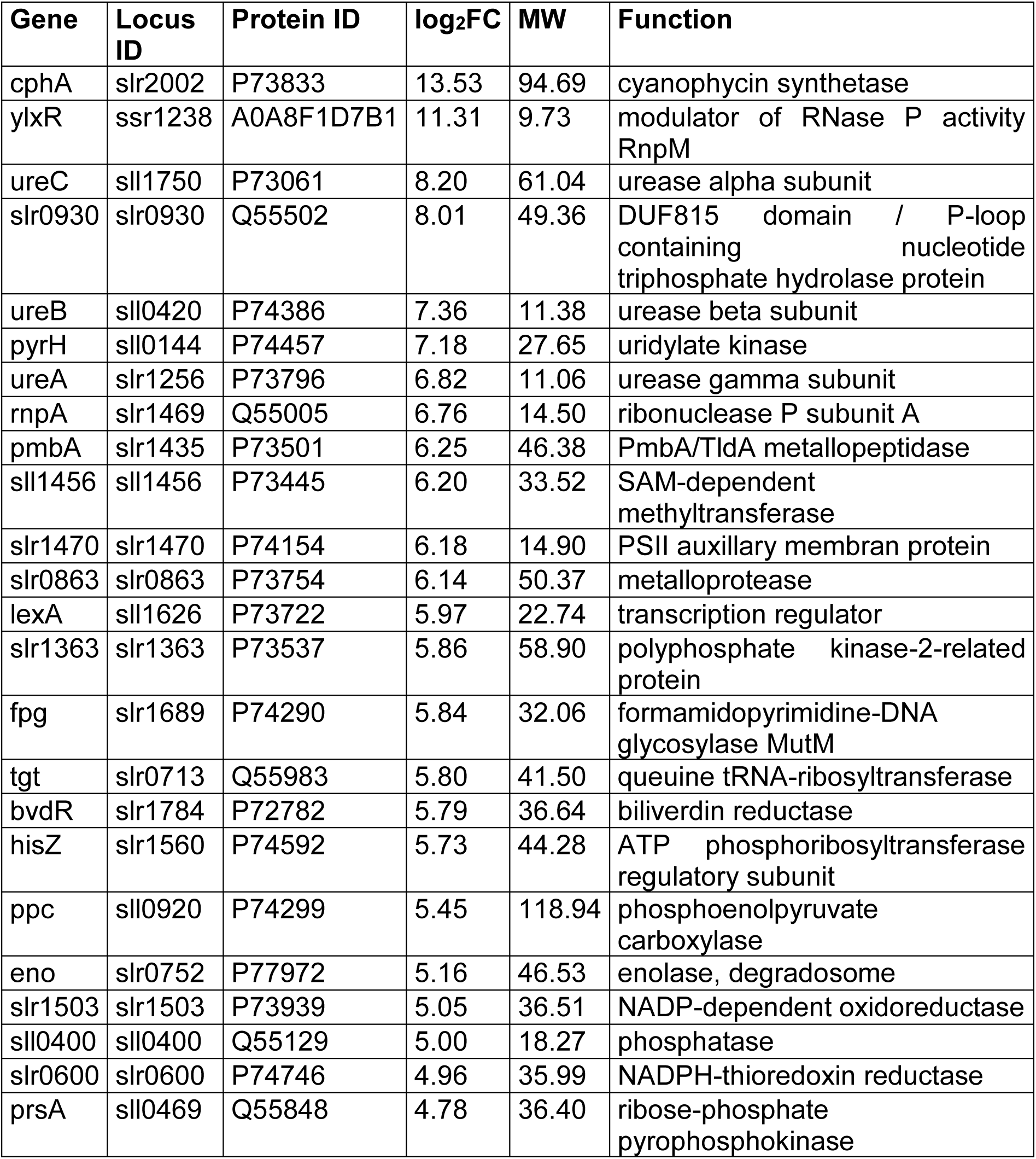
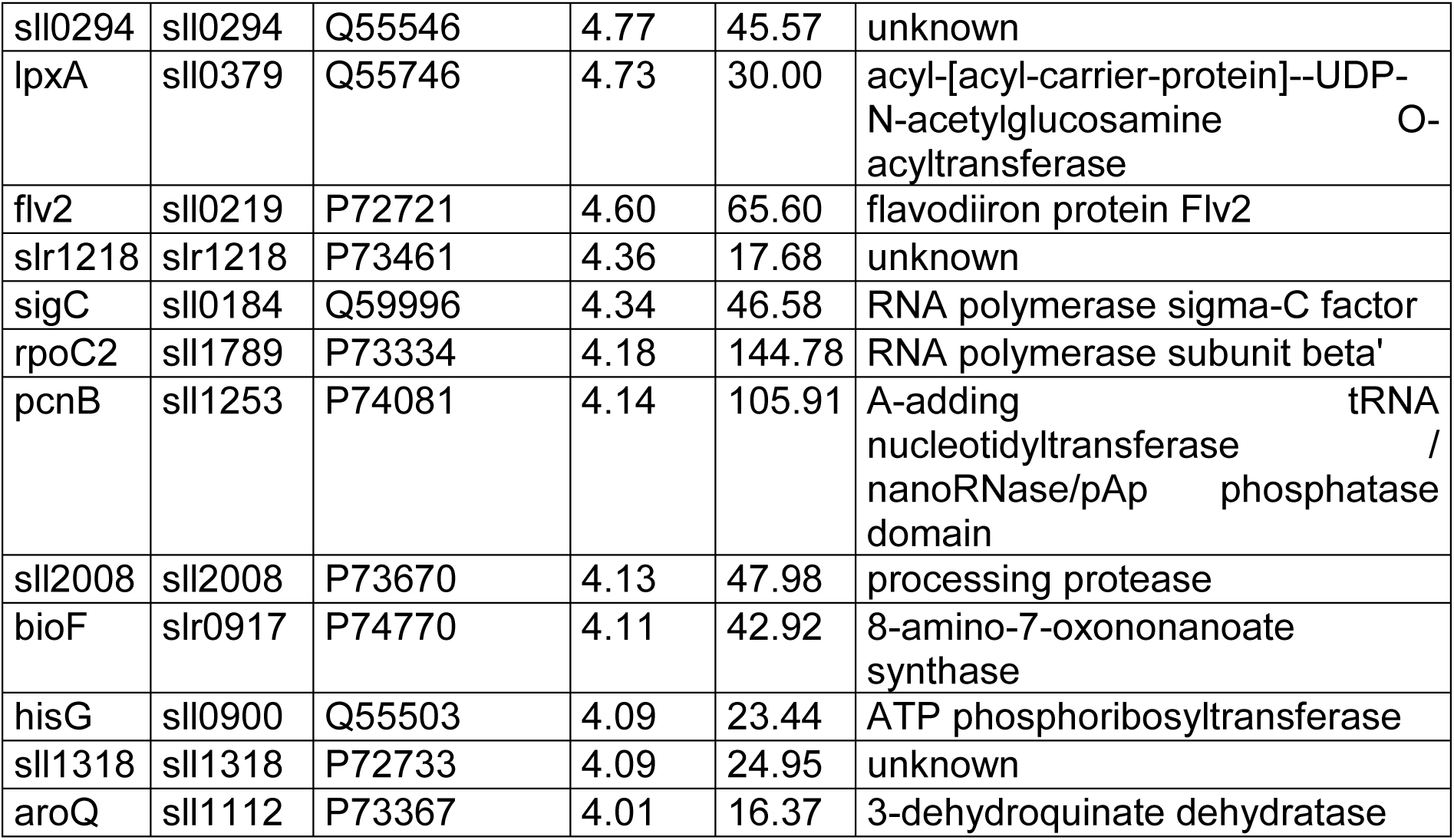
Most enriched proteins identified by MS analysis for FLAG affinity co-IP of samples containing 3×FLAG-tagged Ssr1238/YlxR versus 3×FLAG-tagged sfGFP (log_2_FC ≥4). Given is the gene name followed by the locus ID, the uniport ID, the log_2_FC value and the function. Boldface letters indicate proteins related to RNA metabolism. Acronyms: MW, molecular weight in kDa; PSII, photosystem II. The experiment was performed in biological triplicate.

A second functional category was formed by enriched proteins related to nitrogen metabolism. These proteins are cyanophycin synthetase (CphA) and all three urease subunits (UreA, UreB and UreC). CphA catalyzes the synthesis of the reserve polymer cyanophycin, an Asp-Arg polymer [49] and was with a log_2_FC of 10.99 the most enriched protein in the co-IP with YlxR/Ssr1238. Furthermore, the putative 8-amino-7- oxononanoate synthase BioF, encoded by *slr0917* in a co-transcript with tRNA_i_^fmet^ (***SI Appendix* Fig. S6**), was found. Finally, 3 out of 36 highly enriched proteins perform completely unknown functions (**Table 1**).

## Discussion

### YlxR/Ssr1238 as an RNA-binding protein in Synechocystis

YlxR/Ssr1238 is the cyanobacterial homolog of the better studied YlxR protein in Bacillus [25,26], recently renamed RpnM [28]. However, the relatively low global sequence identity and deviations in certain amino acid positions critical for RNA binding in RpnM (**Figure 1A**) questioned the role of YlxR/Ssr1238 as an RBP. Therefore, we here performed UV crosslinking and sequencing of transcripts that were potentially bound to it. We found two highly enriched transcripts, RNase P RNA and the 3’ segment of repA. The first result is consistent with findings for Bacillus [28] and indeed suggests a conserved function as a modulator of RNase P activity also in cyanobacteria. This interaction probably leads indirectly to the enrichment of RnpA, the protein component of RNase P, which is also bound to the RNase P RNA (**Figure 6**). Interestingly, we found some distinct differences in the transcriptomes of strains that either overexpressed YlxR/Ssr1238 or sfGFP (**Figure 4**). These differences included slightly elevated levels of tRNAs, caused by the likewise elevated RNase P level. The overexpression of YlxR/Ssr1238 may be directly related to this effect by protecting the RNase P RNA from degradation. In E. coli, overexpression of the RNase P RNA led to the more efficient processing of RNase P-dependent polycistronic tRNA operons [50].

**Figure 6.**
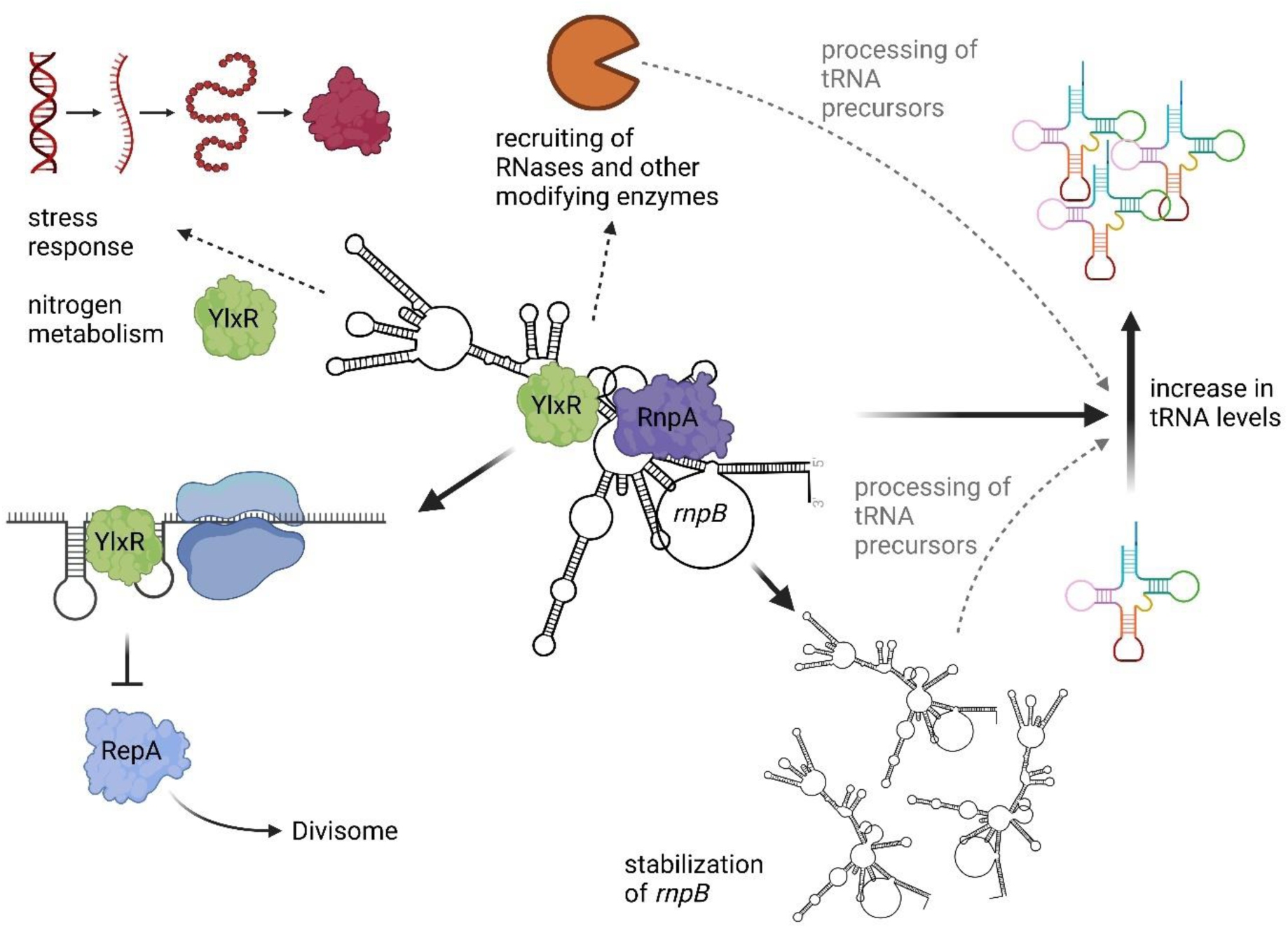
Proposed model of YlxR/Ssr1238 functions in the context of the cyanobacterial cell. Middle: YlxR/Ssr1238 binds to RNase P RNA (Figure 1), which also interacts with its cognate protein component, RnpA, explaining its high enrichment in the protein interactome (Figure 5). Overexpression of YlxR/Ssr1238 also resulted in higher RNase P RNA levels, presumably by stabilizing the RNA (lower right corner), and this likely contributed to the slightly elevated tRNA levels observed (Figure 4). The protein interactome data also suggested that the RnpA-RnpB-YlxR complex may act in close proximity to ribosomes and interact, directly or indirectly, with other key enzymes of RNA metabolism (Figure 5). These effects affected the composition of the transcriptome including the upregulation of stress-related mRNAs (upper left corner). Finally, we observed that YlxR/Ssr1238 recruited a stem-loop structure in the rare lipoprotein *repA* mRNA 3’ segment (Figure 3). This may prevent ribosomes from finishing translation (bottom, left corner). RepA is required for cell division, and once it is produced, it is likely to be recruited to the mid-cell division plane, together with FtsZ. Whether YlxR switches between the *repA* mRNA and the RNase P RNA or how it dissociates from the *repA* mRNA is a matter for further research. The figure was created with elements from Biorender.com.

However, the broader physiological consequences of YlxR/Ssr1238 overexpression indicated the activation of several stress-related genes. Previous work in *Bacillus* revealed that the manipulation of YlxR expression affected the regulation of more than 400 genes and the response to glucose [25]. Interestingly, there is a parallel in the work performed here, as YlxR/Ssr1238 overexpression led to opposite effects on two sRNA regulators that are involved in the regulation of mixotrophic versus photosynthetic lifestyles. While PsrR1, an sRNA regulator of several photosynthetic genes [7], showed an increased transcript level in OE-Ssr1238, the level of PmgR1, a regulator of mixotrophy [9], was decreased (**Figure 4, *SI Appendix* Table S5**).

However, the second highly enriched transcript by UV crosslinking pointed in a different direction. The enriched *repA* 3’ segment contains a strongly folded secondary structure that is not just a rho-independent terminator, but likely serves as a recognition domain for YlxR/Ssr1238. Moreover, this secondary structure is conserved among several homologs from other *Synechocystis* cyanobacteria (**Figure 3C**). The bound RNA segment contained the *repA* stop codon (**Figure 3B**). We therefore speculate that binding of YlxR/Ssr1238 to this region interferes with the completion of *repA* translation. Rare lipoproteins such as RepA are part of the divisome and elongasome complexes [51] and function as lytic transglycosylases, enzymes that are critical for cell wall remodeling during cell division [40]. RlpA (in *Pseudomonas*) is present in higher amounts at the nascent septa of replicating bacteria while it is otherwise found at lower levels along the sidewall length of the bacterium [52]. Therefore, the needed higher levels required for cell division may result from resumption of *repA* translation when YlxR binding is released and possibly captured by RNase P RNA (**Figure 6**).

### Proteins interacting with YlxR/Ssr1238

Finally, co-IP revealed a remarkable set of proteins that interact directly or indirectly with YlxR/Ssr1238. First, the enrichment of ribosomal proteins suggested a close association with ribosomes. An unexpected category of enriched proteins were four proteins related to nitrogen metabolism, all three urease subunits as well as cyanophycin synthetase (CphA). Cyanophycin is a dynamic reserve polymer discovered 138 years ago [53]. The relationship between CphA and YlxR/Ssr1238 is unknown, but a cycling of CphA between a granule-bound active and an inactive form in the cytosol has been described [54]. It is therefore possible that YlxR/Ssr1238 is involved in the formation of the inactive form.

Consistent with the strong interaction of YlxR/Ssr1238 and RNase P RNA was the co-enrichment of RnpA, i.e. the protein subunit of RNase P, in the co-IP experiments. Furthermore, we noticed the enrichment of several enzymes related to RNA metabolism, such as the A-adding tRNA nucleotidyltransferase Sll1253, the queuine tRNA ribosyltransferase Tgt, the putative RNA methyltransferase Sll1967 and the tRNA (guanine-N(7)-)-methyltransferase TrmB (Sll1300) (**Figure 5**). This finding points at the existence of an interaction network of potential tRNA maturation complexes. Interestingly, a complex between a protein-only RNase P enzyme and the tRNA methyl transferases TRM1A and TRM1B has recently been described in the plant *Arabidopsis* [55]. These parallels may suggest that such complexes are more widely distributed in nature.

The co-enrichment of multiple enzymes involved in RNA modification and metabolism is furthermore consistent with reports that the *Bacillus subtilis* YlxR regulates TsaD, a component of the TsaEBD system for the synthesis of the tRNA threonylcarbamoyl adenosine (t6A) modification [26]. Increasing numbers of adaptors or modulators of RNase activities are being identified in bacteria. Well-known examples are the RraA regulator of RNase E activity [56] and the *E. coli* RNase adaptor protein RapZ [57]. Recently, an unrelated modulator of enzymatic activity has also been described for RNase E in cyanobacteria [58]. According to the work of Wicke et al. [28] and based on the results of the present study, YlxR/Ssr1238 is another example of this category of regulatory proteins, but it primarily interacts with RNase P.

Another interesting co-enrichment was observed with RNase E (Rne, **Figure 5**), a central enzyme in the maturation and degradation of mRNAs in cyanobacteria [48]. The co-enrichment of YlxR/Ssr1238 with RNase E is remarkable because RNase E is a central enzyme in mRNA maturation and degradation, while RNase P is best known for its crucial role in the maturation of tRNAs. However, there are several examples where both enzymes cooperate in the maturation of tRNA precursors in *E. coli* [59,60], or in the generation of the 3’ ends of three different proline tRNAs [61,62]. Recently, RNase E was also reported to be involved in the maturation of several tRNAs *in vivo* in *Synechocystis*, the most striking example being its involvement in in the accurate 5’ maturation of the glutamyl-tRNA tRNA^Glu^_UUC_ [63]. YlxR/Ssr1238 is a candidate protein to be physically involved in the crosstalk between both ribonucleases.

Another difference to the RpnM proteins in *Bacillus* are two cysteine pairs that are strikingly conserved in the cyanobacterial YlxR/Ssr1238 homologs and appear arranged on the protein side facing away from the RNA-binding interface (**Figure 1C**), possibly coordinating an Fe-S cluster. This possibility should be addressed in future work.

The results presented here are consistent with findings for the *Bacillus* YlxR homolog as an RNase P modulator [28], but extend its proposed role to the phylum cyanobacteria and suggest a broader function in the integration of tRNA maturation, modification, protein synthesis, nitrogen metabolism and cell division.

## Materials and Methods

### Culturing conditions

*Synechocystis* 6803 PCC-M [64] wild type and mutant strains were cultivated in copper-free BG11 supplemented with 20 mM N-[Tris-(hydroxymethyl)-methyl]-2- aminoethanesulfonic acid pH 7.5 under continuous white light of 50 µmol photons m^-2^ s^-1^ at 30°C. Mutant strains containing pVZ322-derived plasmids were cultivated in the presence of 2.5 µg/mL gentamycin.

### Construction of mutant and tagged strains

Here we constructed *Synechocystis* 6803 derivative strains that host the self-replicating pVZ322 plasmid [65] containing the *ssr1238*-3×FLAG or *sfGFP*-3×FLAG genes under control of a rhamnose-inducible promoter [31,66]. An intermediate cloning vector was prepared using the pUC19 plasmid as backbone. The sequence of the rhamnose construct starting at the *ilvBN* terminator and ending with the *rhaBAD* cassette of pCK355 [66] was amplified using Q5 polymerase (NEB), adding overlaps for AQUA cloning into pUC19 [67]. The ECK120034435 terminator of pCK351 [66] was introduced downstream of the *yfp* gene to prevent readthrough. The sequence of the terminator was amplified using the Q5 polymerase adding overlaps for assembly into the cloned pUC19-P_Rha_-YFP plasmid. The resulting plasmid was then subjected to inverse PCR to add the *ooP* terminator from *E. coli* downstream of the *rhaBAD* cassette and the 3*×*FLAG-tag downstream and in reading frame with the *yfp* gene. The corresponding PCR products were subjected to *Dpn*I digestion (Thermo Fisher Scientific), 5’ phosphorylation by T4 polynucleotide kinase (Thermo Fisher Scientific), self-ligation by T4 DNA ligase (Thermo Fisher Scientific) and heat shock transformation into chemically competent DH5α *E. coli* cells. The *yfp* gene was then replaced with either *ssr1238* or *sfgfp*. For this, the pUC19-P_Rha_-YFP-3*×*FLAG plasmid was inverse amplified by PCR (Q5 polymerase, NEB), omitting the *yfp* gene. The *ssr1238* sequence was amplified from *Synechocystis* genomic DNA and the *sfgfp* sequence was amplified from plasmid pXG10-SF [68]. Overlaps to the rhamnose promoter sequence and the 3*×*FLAG-tag were added to both sequences via the respective primers. After assembly, plasmids pUC19-P_Rha_-*ssr1238*-3*×*FLAG and pUC19-P_Rha_-*sfGFP*-3*×*FLAG were obtained. The multi-host vector pVZ322 [65] was linearized using the restriction enzymes *Xba*I and *Xho*I (Thermo Fisher Scientific). Expression cassettes of *ssr1238* and *sfgfp* were amplified by PCR, adding overlaps to the linearized pVZ322. Throughout the cloning process, PCR products were subjected to a *Dpn*I digestion prior to the assembly to remove template DNA. Unless otherwise indicated, assembly was performed using the AQUA cloning protocol [67]. Transformations were performed by heat shock into chemically competent *E. coli* DH5α cells. After each assembly and transformation step, colony PCRs (GOTAQ, NEB) were performed on positive *E. coli* clones to verify for correct transformation. PCR products for assembly were purified using the NucleoSpin Gel and PCR Clean- up Kit (Macherey-Nagel), and plasmids from positive clones were extracted using the NucleoSpin Plasmid Kit (Macherey-Nagel). Throughout the whole cloning process, intermediate and final plasmids were verified by Sanger sequencing (GATC, Eurofins). All primers are listed in ***SI Appendix* Table S1** and the constructed plasmids in ***SI Appendix* Table S2**.

### Crosslinked immunoprecipitation and sequencing (CLIP-Seq)

Ssr1238-3*×*FLAG and sfGFP-3*×*FLAG expression was induced in 250 mL *Synechocystis* cultures with 0.6 mg/mL rhamnose at an OD_750_ of approximately 0.6-0.7. After 24 h of induction, the cultures were transferred to a plastic tray measuring 21 cm x 14.5 cm x 5.5 cm and crosslinked three times using a UV Stratalinker 2400 (Stratagene) at 0.45 J/cm^-2^. Then, 25 mL culture aliquots were collected to prepare total RNA samples as controls for the corresponding samples and the comparison by RNA-seq. These aliquots were collected by vacuum filtration through hydrophilic polyethersulfone filters (Pall Supor®-800, 0.8 μm) and then snap frozen in liquid nitrogen. Next, 1 mL of phenol, glycerol and Triton X-100 (PGTX) buffer [69] was added and RNA was extracted by incubating the samples at 65°C for 15 min. The samples were then washed and RNA precipitation was carried out as previously described [70]. The cultures volumes remaining after UV crosslinking and withdrawal of 25 mL aliquots for RNA-seq of total cellular RNA were centrifuged at 4,000 x g for 20 min at 4°C to harvest the cells. The cell pellets were then resuspended in 1 mL of lysis buffer (20 mM Tris/HCl pH 8.0, 1 mM MgCl_2_, 150 mM KCl, 1 mM DTT) with 1 U of RNase inhibitor per sample (RiboLock, Thermo Fisher Scientific). Cells were mechanically disrupted using a pre-chilled Precellys 24 (Bertin Instruments) with 200 µL glass beads (Ø 0.1-0.25 mm, Retsch). Samples were spun at 3,000 x g for 2 min at 4°C to remove intact cells and glass beads. Any remaining cell debris was removed via centrifugation at 20,000 x g for 1h at 4°C. Purified lysates were co-immunoprecipitated by incubation with 50 µL packed bead volume of Anti-FLAG M2 magnetic beads (Sigma) for 1 h at 4°C by slightly rotating. The DynaMag™-2 magnetic rack (Thermo Fisher Scientific) was used for flow-through removal and washing. Beads were washed twice with FLAG buffer, then twice with high salt FLAG buffer (50 mM HEPES/NaOH pH 7, 5 mM MgCl_2_, 25 mM CaCl_2_, 1 M NaCl, 10% glycerol, 0.1% Tween20), and then washed two more times with FLAG buffer. For all washes, a volume corresponding to the 20-fold volume of the packed beads was used, and each wash step was carried out for 3 min at 4°C. For subsequent Western blotting, 1/10 bead volumes were kept after the last washing step for immunodetection using the FLAG M2 monoclonal antiserum as described previously [71]. Remaining beads were subjected to proteinase K digestion (1% SDS, 10 mM Tris-HCl pH 7.5, 100 µg/mL proteinase K) for 30 min at 50°C. Then, 200 µL PGTX [69] was added and the samples were boiled for 15 min at 65°C to extract protein-bound RNA. After adding 140 µL chloroform/isoamyl alcohol (24:1), the samples were agitated and incubated for 10 min at room temperature. To obtain phase separation, the samples were centrifugated in a swing-out rotor at 3,260 x g for 3 min at room temperature. The upper aqueous phase was transferred to an RNase-free tube and mixed in a 1:1 ratio with 100% EtOH. Mixtures were further cleaned using the clean and concentrator kit (Zymo Research). RNA was eluted in 15 µL RNase-free ddH2O. Construction of cDNA libraries and sequence analysis were performed by a commercial provider (Vertis Biotechnologie AG Freising, Germany). The RNA samples were fragmented using ultrasound (4 pulses of 30 s each at 4°C). First, an oligonucleotide adapter was ligated to the 3’ end of the RNA molecules. First-strand cDNA synthesis was performed using M-MLV reverse transcriptase and a 3’ primer complementary to the adapter sequence. The first-strand cDNA was purified and the 5’ Illumina TruSeq sequencing adapter was ligated to the 3’ end of the antisense cDNA. The resulting cDNA was amplified by 13 cycles of PCR (15 to 18 for the negative control), to about 10-20 ng/μl using a high-fidelity DNA polymerase and introducing TruSeq barcode sequences as part of the 5’ and 3’ TruSeq sequencing adapters. The cDNA was purified using the Agencourt AMPure XP kit (Beckman Coulter Genomics) and analyzed by capillary electrophoresis. For Illumina NextSeq sequencing, the samples were pooled in approximately equimolar amounts. The cDNA pool was size-fractionated in the range of 200 – 600 bp using a preparative agarose gel and sequenced on an Illumina NextSeq 500 system with a read length of 75 bp.

Raw reads of the RNA-seq libraries were analyzed using the galaxy platform. The galaxy workflow can be accessed and reproduced at the following link: https://usegalaxy.eu/u/luisa_hemm/w/ylxr-3xflag-clip. In short, read quality was check with FastQC in the beginning. After the quality check, reads were mapped to the *Synechocystis* chromosome (GenBank accession NC_000911.1) and the 4 large plasmids pSYSA, pSYSG, pSYSM and pSYSX (NC_005230.1, NC_005231.1, NC_005229.1, NC_005232.1) using BWA-MEM [72] with default parameters. Resulting data were filtered using the BAM filter [73] to remove unmapped reads as well as reads smaller than 20 nucleotides.

The annotation gff-file was based on the reference GenBank annotations amended by several additional genes based on transcriptomic analyses [5]. htseq-count [74] (mode union, nonunique all, minaqual 0) was used to count the reads per transcript. Only transcripts with ≥ 10 reads in at least three samples were subjected to differential expression analysis with deseq2 [75]. To visualize coverage of the reads, bam files were transformed into wiggle files using RSeQC [76] and summarizing the reads from the triplicate analyses for each position.

### Protein co-IP and analyses by LC-MS/MS

Expression of Ssr1238-3*×*FLAG as well as sfGFP-3*×*FLAG was induced in *Synechocystis* cultures (250 mL) at an OD_750_ of ∼0.5 to 0.8 with 0.6 mg/mL rhamnose. Cells were collected by centrifugation at 4,000 × *g* and 4 °C for 20 min after 24 h of induction at an OD_750_ of ∼0.6 to 0.9. The cell pellets were resuspended in approximately 1.5 mL of FLAG-MS buffer (50 mM HEPES/NaOH pH 7, 5 mM MgCl_2_, 25 mM CaCl_2_, 150 mM NaCl) containing protease inhibitor (c0mplete, Roche), mixed with 200 µL of glass beads (Ø 0.1-0.25 mm, Retsch) and lysed mechanically in a prechilled Precellys 24 (Bertin Instruments). To remove glass beads and intact cells, samples were centrifuged for 2 min at 3,000 × *g*. Supernatants were collected for further processing. To solubilize membrane proteins, the lysates were incubated for 45 min in the presence of 2 % n-dodecyl β-D-maltoside in the dark at 4°C. Afterwards, cell debris was removed by centrifugation (21,000 × *g*, 4°C, 1 h). Co-IP was performed by incubating 75 µL packed volume of Anti-FLAG M2 magnetic beads (Sigma Aldrich) with the cleared lysate (1h, 4°C, rotating). Afterwards, the supernatant was removed using the DynaMag™-2 Magnet rack (Thermo Fisher Scientific) and the beads were washed 6 times with FLAG-MS buffer (20x packed bead volume). Western blots were performed as described previously [71] using 1/10 vol of beads and FLAG M2 monoclonal antiserum (Sigma) to verify protein pull down.

Remaining proteins were further processed according to the manufacturer’s protocol for S-Trap micro filters (Protifi), using the provided buffers. Briefly, 23 µL of the 1x lysis buffer (5% SDS, 50 mM TEAB pH 8.5) was added to the beads and incubated at 95°C for 10 min to elute the proteins. Proteins were reduced by incubation with reduction buffer (end concentration 5 mM Tris(2-carboxyethyl)phosphine (TCEP)) for 15 min at 55°C and alkylated by incubation with alkylation buffer (end concentration 20 mM methyl methanethiosulfonate (MMTS)) for 10 min at room temperature in the dark. Afterwards, a final concentration of 1.2% phosphoric acid and then six volumes of binding buffer (90% methanol; 100 mM TEAB; pH 7.55) were added. After gentle mixing, the protein solution was loaded onto an S-Trap filter and spun 3 times at 10,000 × *g* for 30 s. The filters were washed three times using 150 μL of binding buffer. Sequencing-grade trypsin (Promega, 1:25 enzyme:protein ratio) diluted in 20 µL digestion buffer (50 mM TEAB) was added to the filter and proteins were digested at 47 °C for 2 h. To elute peptides, three step-wise buffers were applied: a) 40 μL 50 mM TEAB, b) 40 µl 0.2% formic acid in H_2_O, and c) 50% acetonitrile and 0.2% formic acid in H_2_O. At each step, elution was carried out by centrifugation (10,000 *x* g for 1 min). The peptide solutions were combined and dried in a SpeedVac (DNA 120, Savant).

Thereafter, the peptide concentration was measured using the BCA assay (Thermo Scientific). For LC-MS/MS measurements 800 ng of peptides were analyzed on a Q- Exactive Plus mass spectrometer (Thermo Scientific, San Jose, CA) coupled to an EASY-nLCTM 1000 UHPLC system (Thermo Scientific). The column setup consisted of an Acclaim™ PepMap™ 100 C18 column (Thermo Fisher Scientific, Cat. No. 164946) and a 200 cm µPac GEN1 analytical column (PharmaFluidics, 55250315018210) coupled to a Nanospray FlexTM ion source (Thermo Scientific, ES071) and a fused silica emitter (MS Wil, TIP1002005-5).

For peptide separation, a linear gradient of increasing buffer B (0.1% formic acid in 80% acetonitrile, Fluka) was applied, ranging from 5 to 50% buffer B over the first 80 min and from 50 to 100% buffer B in the subsequent 40 min (120 min separating gradient length). Peptides were analyzed in data dependent acquisition mode (DDA). Survey scans were performed at 70,000 resolution, an Automatic Gain Control (AGC) target of 3e6 and a maximum injection time of 50 ms followed by targeting the top 10 precursor ions for fragmentation scans at 17,500 resolution with 1.6 m/z isolation windows, an NCE of 30 and a dynamic exclusion time of 35 s. For all MS2 scans the intensity threshold was set to 1e5, the AGC to 1e4 and the maximum injection time to 80 ms.

Raw data were processed and analyzed with MaxQuant (Version 1.6.17.0) with the built-in Andromeda peptide search engine [77]. The false discovery rate (FDR) at both the protein and peptide level was set to 1%. Two missed cleavage sites were allowed, no variable modifications were allowed, and carbamidomethylation of cysteines was set as fixed modification. The ‘match between runs’ option was selected. For label free quantification the MaxLFQ algorithm was applied using the standard settings.

Only unique peptides were used for quantification. The database “Synechocystis sp. (strain PCC 6803 / Kazusa)” was downloaded from https://www.uniprot.org/proteomes/UP000001425 on Jun 9^th^, 2022. The proteome raw data acquired by MS were deposited at the ProteomeXchange Consortium (http://proteomecentral.proteomexchange.org) via the PRIDE partner repository [78] under the identifier PXD047746.

Data were normalized at the peptide level by equalizing the medians using the MSstats package (v. 3.20.3) in R (v. 4.0.4) [79]. Subsequently, protein LFQ intensities were log_2_ transformed. *P*-values were adjusted using the Benjamini-Hochberg procedure.

The intensities were compared using LFQ (label-free quantification) values [80] with Perseus (version 2.0.11 [81]). Contaminants and reverse sequences were removed from the matrix and LFQ intensities were log_2_-transformed using Mstats [82]. Before t-test and visualization using a volcano plot, the missing values were replaced by imputation with the normal distribution for each column separately (default settings). For hierarchical clustering (default parameters), only proteins with three valid values in at least one declared group (YlxR_3×FLAG or sGFP_3×FLAG) were considered.

## Data availability

The CLIP-seq and RNA-seq data have been deposited in the SRA database under the BioProject accession number PRJNA1055971. Mass spectrometry raw data have been deposited at the ProteomeXchange Consortium [83] under the accession number PXD047746. Furthermore, all mass spectrometry proteomics datasets used and/or analyzed during this study are available online at the MassIVE repository (http://massive.ucsd.edu/; dataset identifier: MSV000093645.

## Acknowledgments

We thank Janina Nandy for assistance in the construction of plasmids and Marcus Ziemann for help with the visualization of RNA read coverage.

## Author Contributions

WRH and MR designed the project and WRH secured funding. ST and OS carried out MS-based proteomic analyses. JG contributed to the bioinformatic analyses, AK created and analyzed the protein models. All other experiments and analyses were performed by LH and AM. LH and WRH wrote the manuscript with input from all authors.

