## Supplementary material for "Interactors and effects of overexpressing YlxR/RpnM, a conserved RNA binding protein in cyanobacteria": SI Appendix

### *SI Appendix:* Supplementary data

### Datasets

**Dataset 1.** Multiple sequence alignment of homologs of the *Synechocystis* 6803 YlxR/Ssr1238 protein (ID WP_010871325.1) within the phylum cyanobacteria. See separate pdf file.

### Supplementary Tables

**Table S1**. Desoxyoligonucleotide primers used in this study.

| Name | Sequence | Description |
|---|---|---|
| pUC19_fwd | tcgcgcgtttcggtgatg | Linearization of the empty pUC19 backbone |
| pUC19_rev | gacgaaagggcctcgtgatac | |
| Rha_YFP_fwd | aggcgtatcacgaggccctttcgtcaagacccccgcaccgaaa | Amplification of the rhamnose promoter cassette from plasmid pCK355 [1] |
| Rha_YFP_rev | tcaccgtcatcaccgaaacgcgcgattattgcagaaagccatcccg | |
| YFP_terminator_fwd | acgctgaaaagcgtcttttttcgttttggtccaaagccacgttgtgtctc | Amplification of terminator sequence from plasmid pCK351 [1] |
| YFP_terminator_rev | ctcttttctggaatttggtaccgagctgcagttacttgtacagctcgtc | |
| RhaBADpos_tooP_fwd | ccgggcgttttttatttcgcgcgtttcggtgatg | Primer for inverse PCR to add the *ooP* terminator behind the rhamnose regulatory cassette |
| RhaBADpos_tooP_rev | cggcaaccgagcgaattattgcagaaagccatcccg | |
| YFP_3xFLAG_fwd | catgatattgattataaagatgatgatgataaatagctgcagctcggtaccaaa | Primer pair for inverse PCR to introduce the 3xFLAG-tag sequence behind the *yfp* gene |
| YFP_3xFLAG_rev | atctttataatcgccatcatgatctttataatccatcttgtacagctcgtcca | |
| pUC19_Rha_seq1_fwd | ataaacaaataggggttccgcg | Sequencing primer used for checking of the intermediate pUC19 plasmid |
| pUC19_Rha_YFPseq_fwd | acaacatcgaggacggcag | |
| pUC19_Rha_seq3_fwd | aaatctctgatgttacattgcacaag | |
| pUC19_Rha_seq4_fwd | ttgacgacatcaggaggccag | |
| pUC19_Rha_seq_rev | ttgtctgctcccggcatc | |
| pUC19_Rha_3xFLAG_fwd | atggattataaagatcatgatgg | Linearization of the pUC19-Rha-YFP-3xFLAG plasmid removing *yfp* |
| pUC19_Rha_3xFLAG_rev | ttctacctcctttgtatattataaac | |
| ssr1238_3xFLAG_fwd | aatatacaaaggaggtagaaatggcccctggataccgtc | Aplification of the ssr1238 gene sequence from the genome of Synechocystis adding overlaps to the rhamnose promoter as well as the 3xFLAG-tag |
| ssr1238_3xFLAG_rev | tcatgatctttataatccatgggagtggtgagaagacg | |
| GFP_RBS_ol_fwd | gtttataatatacaaaggaggtagaaatgagcaaaggagaagaacttttc | Amplification of the *sfgfp* sequence out of pGX10 [2] adding overlaps to the rhamnose promoter as well as the 3xFLAG-tag |
| GFP_tooP_ol_rev | ctggaatttggtaccgagctgcagctatttgtagagctcatccatg | |
| pVZ322_seq fwd | ataccatgctcagaaaagg | Sequencing primer for the pVZ322 constructs |
| pVZ322_seq rev | caaagccacgttgtgtctc | |

**Table S2.** Plasmids used in this study.

| Name | Resistance | Description |
|---|---|---|
| pUC19 | Amp | *E. coli* cloning vector |
| pCK351 | Km | Plasmid harboring the rhamnose promoter cassette with the ECK120034435 terminator [1] |
| pCK355 | Km | Plasmid harboring the rhamnose promoter cassette with the *ilvBN* terminator [1] |
| pUC19-PRha-YFP-3xFLAG-3xFLAG | Amp, Km | pUC19 plasmid harboring the rhamnose promoter cassette with *yfp* fused to the 3xFLAG-tag sequence. Constructed in this study |
| pUC19-PRha-ssr1238-3xFLAG | Amp, Km | pUC19 plasmid harboring the rhamnose promoter cassette with *ssr1238* fused to the 3xFLAG-tag sequence. Constructed in this study |
| pUC19-PRha-sfGFP-3xFLAG | Amp, Km | pUC19 plasmid harboring the rhamnose promoter cassette with *sfgfp* fused to the 3xFLAG-tag sequence. Constructed in this study |
| pVZ322-PRha-ssr1238-3xFLAG | Km, Genta | pVZ322 plasmid harboring the rhamnose promoter cassette with *ssr1238* fused to the 3xFLAG-tag sequence. Constructed in this study |
| pVZ322-PRha-sfGFP-3xFLAG | Km, Genta | pVZ322 plasmid harboring the rhamnose promoter cassette with *sfgfp* fused to the 3xFLAG-tag sequence. Constructed in this study |

**Table S3.** List of transcripts enriched in XL-Ssr1238 RNA compared to XL-sfGFP ($log_2FC \geq 1$, $p_{adj} \leq 0.1$).

See separate Excel file.

**Table S4.** List of transcripts enriched in XL-Ssr1238 RNA compared to OE-Ssr1238 ($log_2FC \geq 1$, $p_{adj} \leq 0.1$).

See separate Excel file.

**Table S5**. List of transcripts differentially regulated in OE-Ssr1238 RNA compared to OE-sfGFP ($log_2FC \geq 1$, $p_{adj} \leq 0.1$).

See separate Excel file.

**Table S6.** Complete list of detected proteins in coimmunoprecipitation experiments. See separate Excel file.

**Table S7**. Details of data analysis - Two-sided t-test for volcano plot in **Figure 5A** (Perseus v2.0.11 [3]).

| | | |
|---|---|---|
| Presets | log2 transformed LFQ intensities | |
| | imputation of missing values | |
| | Replace missing values from normal distribution | |
| | Width | 0.3 |
| | Down shift | 1.8 |
| | Mode | Separately for each column |
| **Grouping** | First group (right) | Ssr1238-3xFLAG |
| | Second group (left) | sfGFP-3xFLAG |
| **Test** | t-test | |
| **Side** | both | |
| **Number of randomizations** | 250 | |
| **Preserve grouping in randomizations** | None | |
| **FDR** | 0.05 | |
| **S0** | 2 | |

**Table S8**. Details of data analysis - hierarchical clustering in **Figure 5B** (Perseus v2.0.11 [3]).

| | | |
|---|---|---|
| **Presets** | log2 transformed LFQ intensities | |
| | 44 most interesting proteins were selected for clustering | |
| | No imputation of missing values | |
| **Rows** | | |
| Distance | Euclidean | |
| Linkage | Average | |
| Constraint | None | |
| Preprocess with k-means | Yes | |
| | Number of clusters | 300 |
| | Maximal number of Iterations | 10 |
| | Number of restarts | 1 |
| **Columns** | | |
| Distance | Euclidean | |
| Linkage | Average | |
| Constraint | None | |
| Preprocess with k-means | Yes | |
| | Number of clusters | 300 |
| | Maximal number of Iterations | 10 |
| | Number of restarts | 1 |

### Supplementary Figures

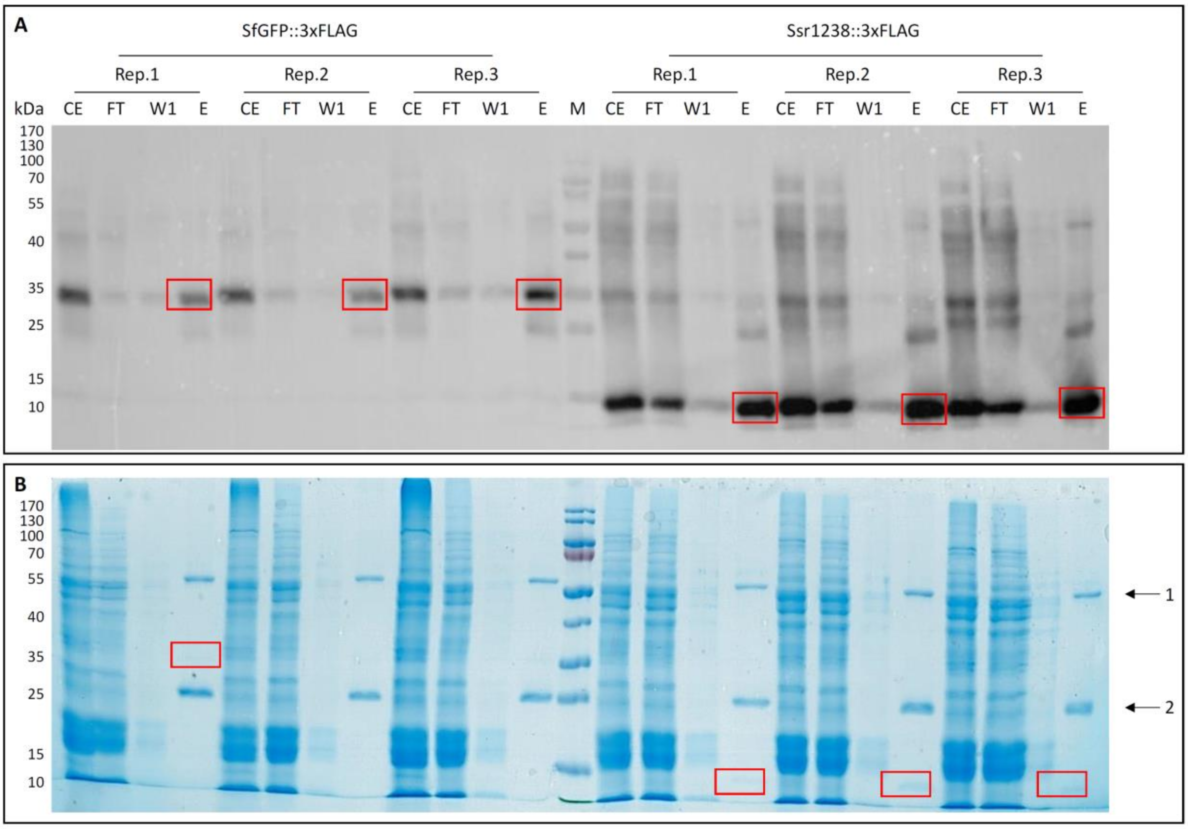

**Figure S1. SDS-PAGE of the crosslinked immunoprecipitates from the strains containing the pVZ322 derivative plasmids for rhamnose-induced expression of either Ssr1238-3xFLAG or sfGFP-3xFLAG.**

**A.** Western blot. **B.** SDS-PAGE of the cross-linked immunoprecipitation of strains containing a pVZ322 plasmid harboring either $P_{rha}$-*ssr1238*-3×FLAG or $P_{rha}$-*sfGFP*-3×FLAG from which upon induction either Ssr1238-3×FLAG or sfGFP-3×FLAG was expressed. The experiment was performed in technical triplicates (Rep. 1-3). Signals indicating the areas collected from the eluate fractions for RNA preparation are boxed in red. Marker (M): PageRuler (Thermo Fischer Scientific), CE: crude extract, FT: flow through, W1: wash fraction one, E: elution fraction. The arrows in panel B mark the two chains of the antibody (1, heavy chain; 2, light chain). The anti-FLAG beads were loaded directly onto the gel, therefore the anti-FLAG antibody chains are visible in the SDS gel. In panel A, the ANTI-FLAG M2-Peroxidase (HRP) antiserum (Sigma Aldrich) was used at a titer of 1:10000.

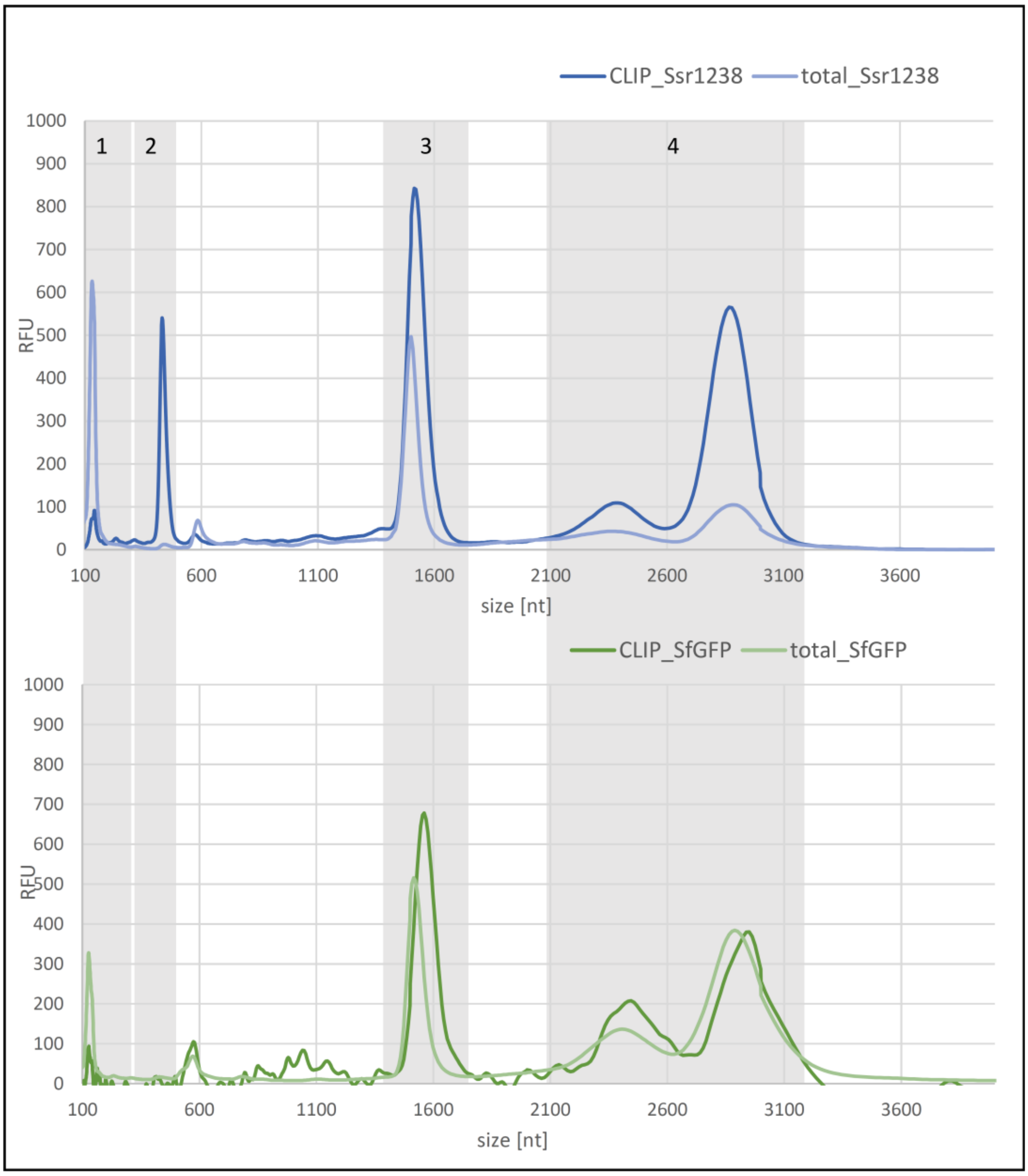

**Figure S2. Fragment analyzer run of UV cross-linked and total RNA of strains P$_{rha}$-*ssr1238*-3×FLAG and P$_{rha}$-*sfGFP*-3×FLAG.**

Electropherogram of cross-linked and total RNA from strains P$_{rha}$-*ssr1238*-3×FLAG and P$_{rha}$-*sfGFP*-3×FLAG. The experiment was done in technical triplicates, the graph shows the average of the samples. The electropherogram was normalized to 1 ng of RNA, the average of the three replicates is displayed. The labelled boxes indicate the following transcript types: 1, tRNA and 5S rRNA; 2, potential interaction partner; 3, 16S rRNA; 4, 23S rRNA. For all samples 2.5 ng were loaded, total RNA samples from both strains were used as controls. RFU, relative fluorescence units; M, molecular mass marker (HS RNA ladder (Agilent)).

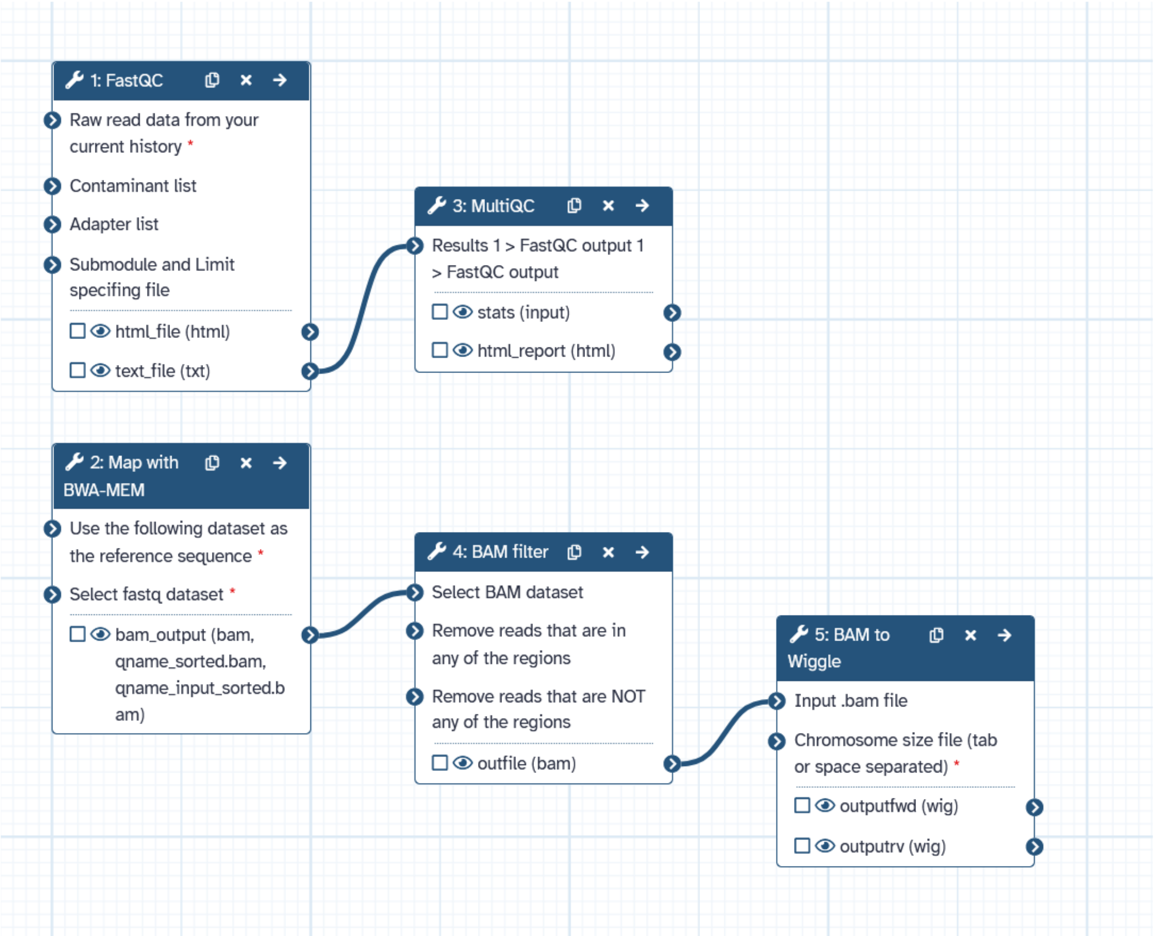

**Figure S3. Bioinformatic workflow used for the analysis of the CLIP-Seq data.** The workflow of the analysis is available also online, here: https://usegalaxy.eu/u/luisa_hemm/w/clip-seq-ssr1238-3xflag.

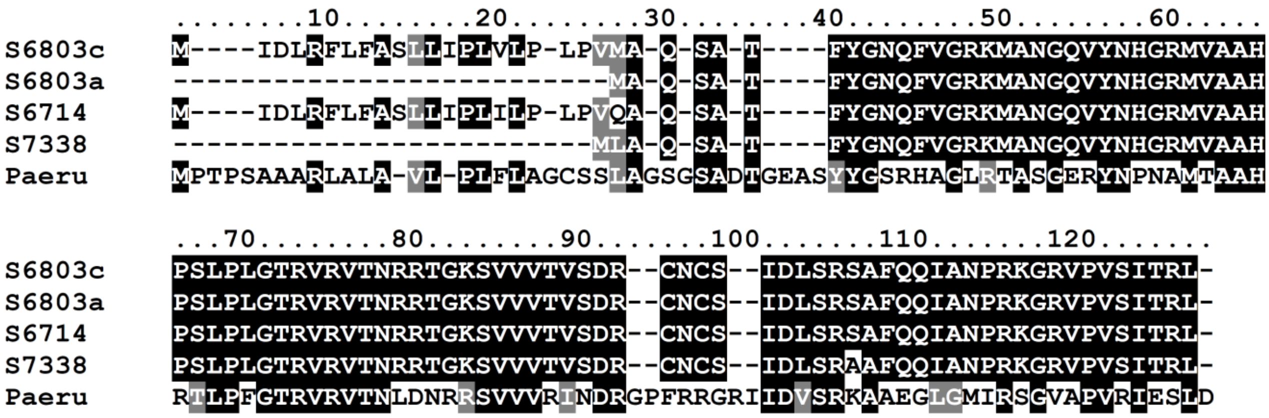

**Figure S4. Comparison of the annotated *ssl3177* ORF in *Synechocystis* 6803 (S6803a), here corrected (S6803c) as discussed in the text, with homologs in strains *Synechocystis* PCC 6714 and PCC 7338/7339 and with the functionally characterized homolog from *Pseudomonas aeruginosa* [4].** The alignment was obtained using Clustal X [5] as implemented in Bioedit v7.2.6 and shaded using Boxshade [6] v3.3 accessed at https://junli.netlify.app/apps/boxshade/.

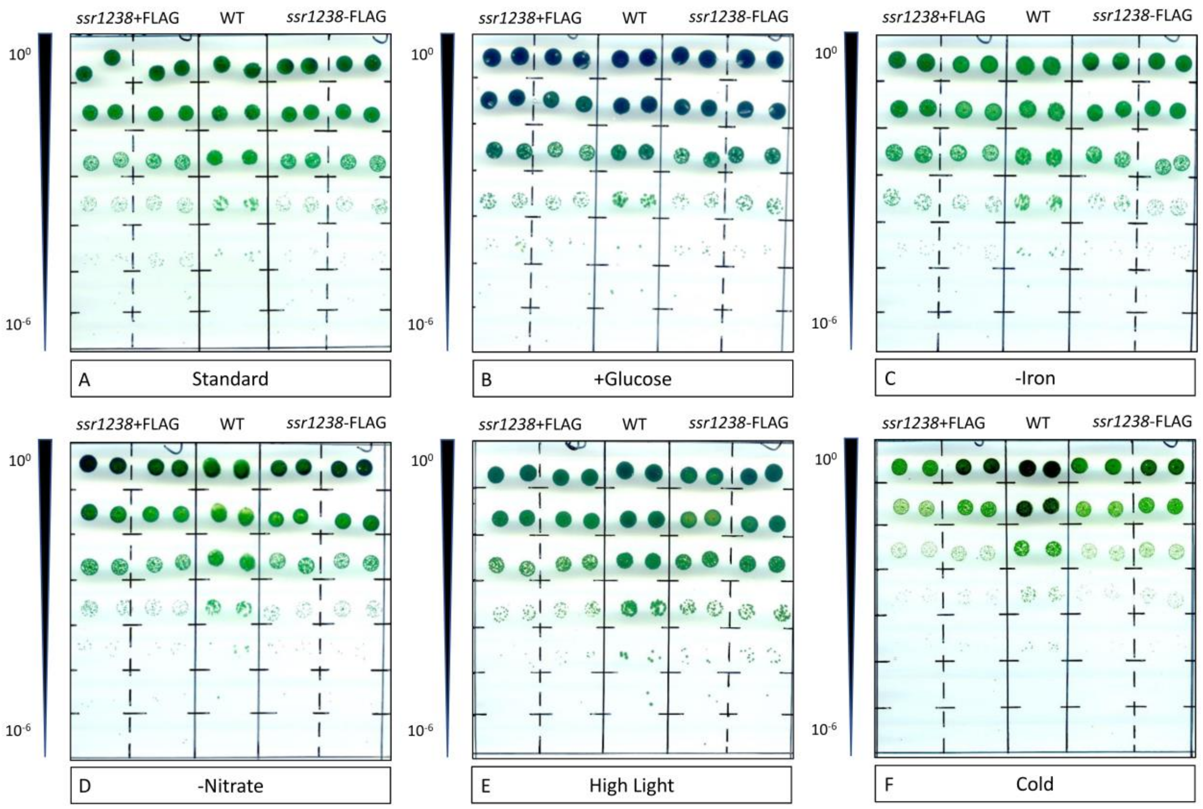

**Figure S5. Test for phenotypic effects of *ssr1238* overexpression under different growth conditions.** Five µL aliquots of *Synechocystis* 6803 cultures at an $OD_{750}$ of 0.2 were spotted in quadruplicates on BG11 plates for a strain overexpressing the native *ssr1238* gene (*ssr1238*-FLAG) or with an added 3×FLAG epitope tag (*ssr1238*+FLAG). For comparison, the parental wild-type (WT) strain was spotted in duplicates. Every row represents a 10x dilution, from $10^0$ to $10^{-6}$. Standard growth conditions were employed alongside elevated light treatment (75 µmol quanta × $m^{-2}$ $s^{-1}$), cold temperature (20°C), with 10 mM of added glucose, or using medium lacking nitrate or iron to impose nutrient stress. Rhamnose (0.6 mg/mL) was present in all the plates to induce the $P_{rha}$ promoter.

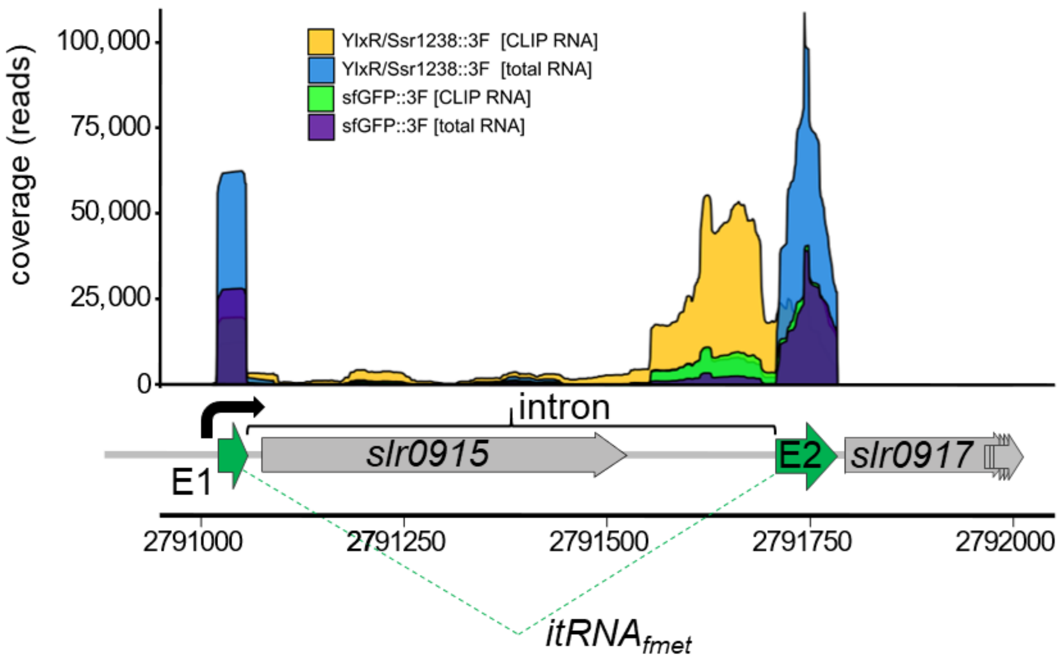

**Figure S6. Accumulation and selective enrichment of a group I intron fragment upon *ssr1238* overexpression and UV crosslinking.** The transcriptional start site (bend arrow) of the entire transcriptional unit is located at position 2,791,015 of the chromosome according to GenBank file NC_000911.1 [7], 4 nt upstream of the 5' end of the mature tRNA$_i^{fmet}$. The tRNA exons 1 and 2 (E1 and E2) are colored green. The corrected start codon of *slr0917/bioF* begins at position 2,791,820.

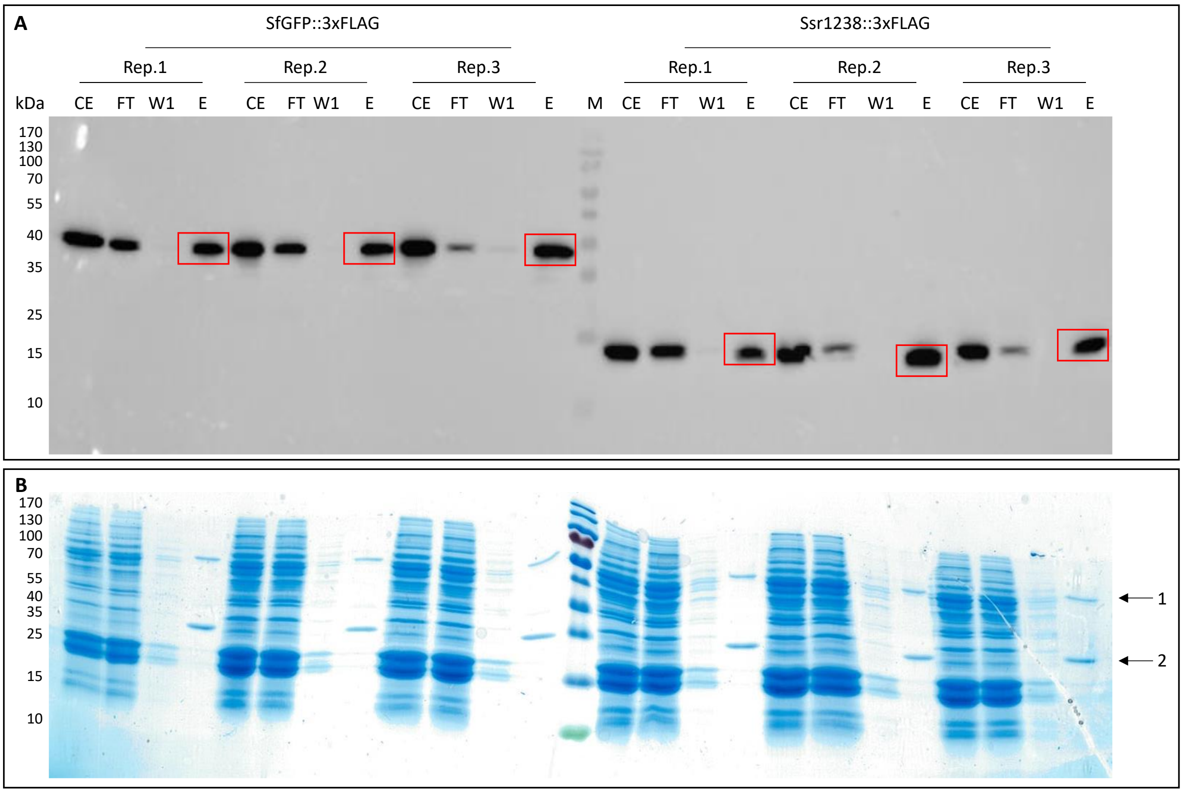

**Figure S7. Western blots as quality control of the co-immunoprecipitation of Ssr1238-3×FLAG and sfGFP-3×FLAG for mass spectrometry.** Western Blot (top) and SDS PAGE (bottom) of a protein pull-down of strains containing a pVZ322 plasmid harboring $P_{rha}$-*ssr1238*-3×FLAG or $P_{rha}$-*sfGFP*-3×FLAG from which upon induction either Ssr1238-3×FLAG or sfGFP-3×FLAG was expressed. The experiment was performed in technical triplicates (Rep. 1-3). The proteins were separated by 15 % SDS-PAGE. Relevant signals in the eluate fractions are boxed in red. Arrows in panel B mark the heavy and light chain of the antibody. Marker (M): PageRuler (Thermo Fischer scientific), antibody: ANTI-FLAG M2-peroxidase (HRP) (Sigma Aldrich), CE: crude extract, FT: flow through, W1: wash fraction one, E: elution fraction.
