## Supplementary material for "Interactors and effects of overexpressing YlxR/RpnM, a conserved RNA binding protein in cyanobacteria": Dataset 1

**Dataset 1. Homologs of the *Synechocystis* 6803 YlxR/Ssr1238 protein (ID WP_010871325.1) within the phylum cyanobacteria.**

```
                             10        20        30        40        50        60        70        80        90       100
                     ....|....|....|....|....|....|....|....|....|....|....|....|....|....|....|....|....|....|....|....|
ref|WP_010871325.1|  -----------------------MAP-------------------GYRRCLSCRKVGDRRQFWRIVRV-----Y-P---------SRT-----VQ-LD--
ref|WP_190597250.1|  -----------------------MAP-------------------GYRRCLSCRKVGDRRQFWRIVRV-----Y-P---------SRT-----VQ-LD--
ref|WP_028947476.1|  -----------------------MAP-------------------GYRRCLSCRKVGDRRQFWRIVRV-----Y-P---------SRT-----VQ-LD--
gb|MEB3228520.1|     --------MP--PG--------------------------------YRRCLSCRKVAHRREFWRIVRV-----H-P---------SR---T--VQ-LDHG
gb|MEB3311284.1|     ---------------------------------------M---KPNYRRCISCRKVAPKEEFWRIVRV-----Y-P---------SRQ-----VQ-LD--
gb|MEB3122126.1|     -----------------MKSHL---------RQ-KKP--------GDRRCISCRKLAPRNEFWRIVRV-----Y-P---------SRQ-----VQ-LD--
ref|WP_075904775.1|  --------------------------------M--KP--------NYRRCLSCRKVGPKQEFWRIVRV-----Y-P---------SRK-----VQ-LD--
gb|NEQ20425.1|       ---------ME-------------------------PN--------YRRCLSCRKVAHKETFWRIVRV-----Y-P---------SR---Q--VQ-LDEG
gb|UXE61785.1|       -----------------------------M-----KP--------QHRRCLSCRKVAPKNEFWRIVRV-----Y-P---------SRQ-----IQ-LD--
ref|WP_293043112.1|  --------------------------------M--KP--------NYRRCLSCRKVAPKQEFWRIVRV-----Y-P---------SRK-----VQ-LD--
gb|NEO94735.1|       --------------------------------M--KP--------NYRRCLSCRKVAPKQEFWRIVRV-----Y-P---------SRK-----VQ-LD--
ref|WP_293076746.1|  --------------------------------M--KP--------NYRRCLSCRKVAPKQEFWRIVRV-----Y-P---------SRK-----VQ-LD--
ref|WP_071107114.1|  --------------------------------M--KP--------NYRRCLSCRKVAPKQEFWRIVRV-----Y-P---------SRK-----VQ-LD--
ref|WP_044492698.1|  --------------------------------M--KP--------NYRRCLSCRKVAPKQEFWRIVRV-----Y-P---------SRK-----VQ-LD--
ref|WP_293017770.1|  --------------------------------M--KP--------NYRRCLSCRKVAPKQEFWRIVRV-----Y-P---------SRK-----VQ-LD--
ref|WP_293114070.1|  --------------------------------M--KP--------NYRRCLSCRKVAPKQEFWRIVRV-----Y-P---------SRK-----VQ-LD--
gb|PZV27550.1|       -----------------MKSHL---------RQ-KKP--------GDRCCISCRKLAPRNEFWRIVRV-----Y-P---------SRQ-----VQ-LD--
ref|WP_070395889.1|  --------------------------------M--KP--------NYRRCLSCRKVAPKQEFWRIVRV-----Y-P---------SRK-----VQ-LD--
gb|NEQ64070.1|       --------------------------------M--KP--------NYRRCLSCRKVAPKQAFWRVVRV-----Y-P---------SRK-----VQ-LD--
gb|MEB3190993.1|     -----------------------------M-----KS--------QHRRCLSCRKVAPKNEFWRIVRV-----Y-P---------SRQ-----IQ-LD--
gb|MBW4545396.1|     ---------ME-------------------------PN--------YRRCLSCRKVAHKETFWRIVRV-----Y-P---------SR---Q--VQ-LDEG
gb|MBE9166779.1|     --------------------------------M----K------PNYRRCISCYKIAPKASFWRVVRV-----Y-L---------SHQ-----I------
gb|NES99769.1|       --------------------------------M-Q-P--------NYRRCLSCRRVGLKQEFWRIVRV-----F-P---------SRK-----VQ-LD--
gb|NER21363.1|       --------------------------------M--KP--------NYRRCLSCRKVAPKQAFWRVVRV-----Y-P---------SRK-----VQ-LD--
ref|WP_096621468.1|  --------------------------------M-K-P--------NYRRCISCRKVGLKQEFWRIVRV-----F-P---------SGQ-----VQ-LD--
ref|WP_192221722.1|  ---------ME-------------------------PN--------YRRCLSCRKVAHKETFWRIVRV-----Y-P---------SR---Q--VQ-LDEG
gb|NES19959.1|       --------------------------------M--KP--------NYRRCLSCRQVAPKQSFWRVVRV-----Y-P---------SRK-----VQ-LD--
ref|WP_193875631.1|  -----------------------------M-----KP--------NYRRCISCRRVAPKPEFWRIVRL-----F-P---------DRT-----VQ-LD--
gb|NEP56036.1|       --------------------------------M--KP--------NYRRCLSCRKVAPKPAFWRVVRV-----Y-P---------SRK-----VQ-LD--
gb|NET56888.1|       --------------------------------M--KP--------NYRRCLSCRQVGPKQAFWRIVRV-----Y-P---------SRK-----VQ-LD--
ref|WP_272066750.1|  --------------------------------M--KP--------NFRRCVSCRAIAPKEALWRVVRV-----Y-P---------SRT-----VQ-LD--
ref|WP_008308507.1|  --------------------------------M--E--------PNYRCCISCRRIAPKAQFWRVVRL-----Y-P---------TRI-----AQ-LD--
gb|PSR15992.1|       -----------------MQP-------------------------NYRRCVSCRRVGPKAEFWRLVRVYP---------------NRT-----VQ-LD--
gb|PSN10559.1|       -----------------MQP-------------------------NYRRCVSCRRVGPKAEFWRLVRVYP---------------DRT-----VQ-LD--
gb|MBD0344432.1|     ------------------------------------------MEPNYRRCLSCRTVAPKEAFWRVVRV-----Y-P---------SRQ-----VQ-LD--
gb|NEP16756.1|       -----------------MK-----------------P--------GHRRCVSCRKVAPKSDFWRIVRV-----F-P---------SRT-----VQ-LD--
ref|WP_080808818.1|  --------------------------------M--KP--------NLRRCISCRKVAPKSALWRIVRV-----F-P---------SRT-----VQ-LD--
ref|WP_015954383.1|  -----------------------MKP-------------------NYRRCVSCGQVALKDSFWRIVRV-----Y-P---------SRK-----LQ-LD--
ref|WP_290221549.1|  -----------------------M-----------KP--------NYRRCASCRKVAPKEDFWRIVRV-----H-P---------SQT-----VQ-LD--
gb|MBJ7899262.1|     ---------MP-PG--------------------------------YRRCLTCRRTAPREEFWRVVRV-----H-P---------SR---E--IQ-LDSG
ref|WP_094348539.1|  --------------------------------M--KP--------NYRRCISCRKVGSKDQFWRIVRV-----F-P---------SGK-----VQ-LD--
gb|TVQ19707.1|       --------------------------------M-K-P--------NDRRCISCRKVDQKQYFWRIVRV-----Y-P---------SRT-----VQ-LD--
ref|WP_300417529.1|  --------------------------------M--KP--------NYRRCISCRKVGSKDQFWRIVRV-----F-P---------SGK-----VQ-LD--
ref|WP_110985927.1|  --------------------------------M--EP--------NYRRCICCRKVAPKQEFWRVVRV-----H-P---------DRS-----VQ-LD--
ref|WP_114081474.1|  --------------------------------M--KP--------NYRRCISCRKVGSKDEFWRIVRV-----F-P---------SGK-----VQ-LD--
gb|MCA1994153.1|     --------------------------------M--KP--------NYRRCLSCRKVAPKEAFWRIVRV-----H-P---------SRQ-----VQ-LD--
ref|WP_190490988.1|  --------------------------------M--QP--------NYRRCVSCRKVAPKQSFWRIVRV-----Y-P---------SRQ-----LQ-LD--
```

```
ref|WP_272127684.1|    ---------------------------------------M---KPNLRRCLACRQVAPKSSFWRIVRV-----H-P---------SRT-----VQ-LD--
ref|WP_015183318.1|    ---------------------------------------M---EPNYRRCLSCRKVAPKKAFWRVVRL-----Y-P---------SRQ-----VQ-LD--
ref|WP_190712634.1|    --------------------------------M-GESQAK---QPNARRCISCRKVAPKDAFWRVVRV-----Y-P---------SRT-----VQ-LD--
gb|NJM61055.1|         ---------------------------------------M---KPNYRRCASCRKIAPKEAFWRAVRV-----Y-P---------SHQ-----VQ-LD--
gb|MDJ0704997.1|       -----------------------MKP-------------------NYRRCISCRKIGHKQAFWRIVRV-----H-P---------SRA-----VQ-LN--
ref|WP_292849343.1|    --------------------------------M--KP--------NYRRCISCRKVGSKDEFWRIVRV-----F-P---------SGK-----VQ-LD--
ref|WP_169155198.1|    --------------------------------M--KP--------NYRRCISCRKIGLKQEFWRIVRV-----F-P---------SGQ-----VQ-LD--
tpg|HSM83622.1|        -----------------MQP-------------------------NYRRCVSCRRVAPKADFWRVVRV-----Y-P---------DRT-----VR-LDYG
gb|MEB3274626.1|       ------------------------------MPM--PP--------NYRRCISCRKVAPKAEFWRVVRL-----A-Q---------DRS-----IQ-LD--
ref|WP_324983291.1|    --------------------------------M--KP--------NYRRCISCRKVGSKDQFWRIVRV-----F-P---------SGK-----VQ-LD--
ref|WP_292866613.1|    --------------------------------M--QP--------NYRRCISCRKVGSKDEFWRIVRV-----F-P---------SGK-----VQ-LD--
ref|WP_022606905.1|    --------------------------------M----P------PNYRRCVSCRRSGPKASFWRVVRV-----F-P---------SQT-----V------
ref|WP_292789808.1|    --------------------------------M--KP--------NYRRCISCRKVGLKDEFWRIVRV-----F-P---------SGK-----VQ-LD--
ref|WP_096570286.1|    --------------------------------M-K-P--------NYRRCISCRRVGQKQEFWRIVRV-----F-P---------SRK-----VQ-LD--
ref|WP_012595979.1|    ------------------------------------MK------PNYRRCISCRKVAPKQAFWRIVRV-----Y-P---------SQE-----VQ-LD--
ref|WP_324629747.1|    --------------------------------M--KQN--------YRRCISCRQVAPKNAFWRVVRV-----Y-P---------SHQ-----VQ-LD--
ref|WP_190731269.1|    --------------------------------M--KP--------NYRRCVSCRKVGSKDEFWRIVRV-----F-P---------SGK-----VQ-LD--
tpg|HAA30330.1|        --------------------------------M--EP--------NYRRCLSCRTVAPKQSFWRVVRV-----Y-P---------SRQ-----VQ-LD--
ref|WP_298906330.1|    --------------------------------M--KP--------NYRRCISCRKVGSKDEFWRIVRV-----F-P---------SGK-----VQ-LD--
dbj|BAZ68765.1|        -----------------------M--------M--KP--------NYRRCISCRKVGIKNEFWRIVRV-----F-P---------SRQ-----VQ-LD--
gb|MBW4433734.1|       --------------------------------M--KP--------NYRRCISCRKVGIKNEFWRIVRV-----F-P---------SRQ-----VQ-LD--
ref|WP_181927846.1|    --------------------------------M--KP--------NYRRCISCRKVGSKDEFWRIVRV-----F-P---------SGK-----VQ-LD--
ref|WP_012412015.1|    --------------------------------M--KP--------NYRRCISCRKVGSKDEFWRIVRV-----F-P---------SGK-----VQ-LD--
ref|WP_322677349.1|    --------------------------------M--KP--------NYRRCISCRKVGSKDEFWRIVRV-----F-P---------SGK-----VQ-LD--
ref|WP_015139046.1|    --------------------------------M--KP--------NYRRCVSCRKVGLKEEFWRIVRV-----F-P---------SGK-----VQ-LD--
ref|WP_104905836.1|    --------------------------------M--KP--------NYRRCISCRKVGSKDEFWRIVRV-----F-P---------SGK-----VQ-LD--
ref|WP_206814010.1|    ---------------------------------------M---PSNIRRCLSCRKIAPKTSFWRIVRV-----Y-P---------SGQ-----VQ-LD--
gb|MDJ0601942.1|       -----------------------MKP-------------------NYRRCISCRRVAPKEAFWRIVRV-----Y-P---------SRQ-----VQ-LD--
ref|WP_292717392.1|    --------------------------------M--KP--------NYRRCISCRKVGSKDEFWRIVRV-----F-P---------SGK-----VQ-LD--
ref|WP_179066568.1|    --------------------------------M--KP--------NYRRCISCRKVGSKDEFWRIVRV-----F-P---------SGK-----VQ-LD--
ref|WP_322722179.1|    --------------------------------M--KP--------NYRRCISCRKVGSKDEFWRIVRV-----F-P---------SGK-----VQ-LD--
ref|WP_185581622.1|    --------------------------------M--KP--------NYRRCISCRKVGSKDEFWRIVRV-----F-P---------SGK-----VQ-LD--
ref|WP_292822895.1|    --------------------------------M--KP--------NYRRCISCRKVGSKDEFWRIVRV-----F-P---------SGK-----VQ-LD--
ref|WP_196512348.1|    --------------------------------M--KP--------NYRRCISCRKVGSKDEFWRIVRV-----F-P---------SGK-----VQ-LD--
gb|OKH54711.1|         --------------------------------M-K-P--------NYRRCLSCRKIGLKEEFWRIVRV-----F-P---------SGQ-----VQ-LD--
ref|WP_292785937.1|    --------------------------------M--QP--------NYRRCISCRKVGSKDEFWRIVRV-----F-P---------SGK-----VQ-LD--
gb|TAE56869.1|         --------------------------------M-K-P--------NYRRCLSCRRVGLKQEFWRIVRV-----F-P---------SRK-----VQ-LD--
gb|PZV08645.1|         --------------------------------M--KP--------SYRRCVACRKVALKVDFWRVVRV-----Y-P---------SQV-----VQ-LD--
ref|WP_292751094.1|    --------------------------------M--KP--------NYRRCISCRKVGSKDEFWRIVRV-----F-P---------SGK-----VQ-LD--
ref|WP_073597769.1|    ------------------------------------------MQKNYRRCISCRKVAPKESFWRIVRV-----Y-P---------SRQ-----VQ-LD--
ref|WP_073595002.1|    --------------------------------M-K-P--------NYRRCISCRKVDLKSNFWRIVRV-----Y-P---------SRQ-----VQ-LD--
gb|MCG8363753.1|       -----------------------MKP-------------------NYRRCASCRKVAHKQAFWRIVRV-----H-P---------SRA-----VQ-LN--
gb|PSO56092.1|         --------------------------------M-KPNY---------RRCVSCRQVAKKESFWRIVRV-----H-P---------SGQ-----VQ-LD--
gb|MBF2088208.1|       --------------------------------M-K-P--------NDRRCISCRKVDQKEYFWRIVRV-----Y-P---------SRT-----VQ-LD--
gb|NES80898.1|         --------------------------------M----------KPNYRRCLSCRKVAPKESFWRIVRV-----Y-P---------SQQ-----VH-LD--
gb|PSO96923.1|         --------------------------------M-KPNY---------RRCVSCRQVAKKESFWRIVRV-----H-P---------SGQ-----VQ-LD--
dbj|BAZ39493.1|        -----------------------------MK-P--N----------YRRCISCRKIGLKQEFWRIVRV-----F-P---------SRQ-----VQ-LD--
ref|WP_322731565.1|    --------------------------------M--KP--------NYRRCISCRKVGSKDEFWRIVRV-----F-P---------SGK-----VQ-LD--
ref|WP_193978934.1|    --------------------------------M-K-P--------NYRRCISCRKVDLKSNFWRIVRV-----Y-P---------SRQ-----VQ-LD--
ref|WP_300507896.1|    -----------------------MKP-------------------NYRRCLSCGRVAPKEEFWRIVRV-----Y-P---------SRQ-----VQ-LD--
gb|TVQ45794.1|         ------------------------------------------MKPNYRRCLSCRQIAPKESFWRIIKV-----Y-P---------SQE-----VK-LD--
gb|MBW4655594.1|       ------------------------------------------MNPNHRRCASCRTIAPKENFWRIVRV-----Y-P---------SRT-----IQ-LD--
ref|WP_204152300.1|    --------------------------------M-K-P--------NDRRCISCRKVDQKEYFWRIVRV-----Y-P---------SRT-----VQ-LD--
```

```
ref|WP_292734233.1|   --------------------------------M--KP--------NYRRCISCRKVGSKDQFWRIVRV-----F-P---------DGK-----VQ-LD--
gb|PSN19469.1|        --------------------------------M--EP--------NYRRCISCRRIGHKSEFWRIVRL-----A-D---------SRQ-----VQ-LD--
ref|WP_094331183.1|   --------------------------------M--KP--------NYRRCISCRKVGSKDEFWRIVRV-----F-P---------SGK-----VQ-LD--
gb|MDW8178835.1|      --------------------------MP--------PG--------WRRCLSCRTVGPREQFWRVVRL-----H-P---------SH---Q--VV-LDQG
ref|WP_223049825.1|   --------MEP--------------------------N--------YRRCLSCRKVAHRSEFWRIVRV-----Y-P---------SQ---T--IE-LDQG
dbj|BDA66563.1|       --------------------------------M-K-P--------NYRRCLSCRKIGLKEEFWRIVRV-----F-P---------SGQ-----VQ-LD--
ref|WP_194027457.1|   ---------MK-------------------------PN--------YRRCISCRKVASKEFFWRIVRV-----Y-P---------SR---K--VQ-LDQG
gb|NEO98984.1|        -----------------------MQP-------------------NYRRCLSCRIVLPKEAFWRVIRV-----Y-P---------SRK-----VQ-LD--
gb|MDZ8004527.1|      --------------------------------M--KP--------NYRRCISCRKLGLKEEFWRIVRV-----F-P---------SGT-----VQ-LD--
gb|KAB8333374.1|      --------------------------------M--KP--------NYRRCISCRKIGLKQEFWRIVRV-----F-P---------SGQ-----VQ-LD--
gb|MCY7392750.1|      --------------------------------M--KP--------GYRRCVACRKAAPKVVFWRVVRV-----Y-P---------SQS-----VQ-LD--
gb|MBE9078818.1|      -----------------------MKP-------------------NYRRCMSCRRIDHRQNFWRIVRV-----H-P---------SRQ-----VQ-LD--
ref|WP_322669753.1|   --------------------------------M--KP--------NYRRCISCRKVSSKDQFWRIVRV-----F-P---------SGK-----VQ-LD--
ref|WP_100898624.1|   --------------------------------M--KP--------NYRRCISCRKVGSKDEFWRIVRV-----F-P---------SGK-----VQ-LD--
ref|WP_229551764.1|   --------------------------------M--KP--------NYRRCISCRKVGSKDQFWRIVRV-----F-P---------SGK-----VQ-LD--
gb|MBD0304031.1|      --------------------------------M--KP--------NYRRCISCRKVGSKDEFWRIVRV-----F-P---------SGK-----VQ-LD--
ref|WP_169218276.1|   --------------------------------M--KP--------NYRRCISCRKIGLKQEFWRIVRV-----F-P---------SGQ-----VQ-LD--
gb|NEO32345.1|        -----------------------MKP-------------------NYRRCLSCRTILPKEAFWRVVRV-----H-P---------SRN-----VQ-LD--
ref|WP_190814021.1|   ------------------------------------------MEPNYRRCVSCRKVAPKSAFWRIVRV-----Y-P---------SQQ-----LQ-LD--
ref|WP_196527153.1|   --------------------------------M--KP--------NYRRCISCRKVGSKDEFWRIVRV-----F-P---------SGK-----VQ-LD--
ref|WP_323196799.1|   --------------------------------M--KP--------NYRRCISCRKVGLKEDFWRIVRV-----F-P---------SGK-----VQ-LD--
gb|PSO55897.1|        --------------------------------M-KPNY---------RRCVSCRQVAKKESFWRIVRV-----H-P---------SGQ-----VQ-LD--
ref|WP_229482756.1|   --------------------------------M--KP--------NYRRCISCRKVGSKDEFWRIVRV-----F-P---------SGK-----VQ-LD--
ref|WP_292709822.1|   --------------------------------M--KP--------NYRRCISCRKVGSKDEFWRIVRV-----F-P---------SGK-----VQ-LD--
ref|WP_069073458.1|   --------------------------------M--KP--------NYRRCISCRKVGSKDEFWRIVRV-----F-P---------SGK-----VQ-LD--
dbj|BAU66516.1|       --------------------------------M--QP--------NYRRCLSCRKIAPKNSFLRIVRI-----Y-P---------SYQ-----IK-LE--
ref|WP_229454557.1|   --------------------------------M--KP--------NYRRCISCRKVGSKDEFWRIVRV-----F-P---------SGK-----VQ-LD--
ref|WP_322697626.1|   --------------------------------M--KP--------NYRRCISCRKVSSKDQFWRIVRV-----F-P---------SGK-----VQ-LD--
ref|WP_273766710.1|   --------------------------------M--KP--------NYRRCISCRRVGSKDEFWRIVRV-----F-P---------SGK-----VQ-LD--
gb|MBE7381418.1|      -----------------------MKP-------------------NYRRCISCRTIAPKEAFWRIVRV-----Y-P---------TRT-----VQ-LE--
gb|RCJ24275.1|        --------------------------------M--KP--------NYRRCISCRKLGLKEEFWRIVRV-----F-P---------SGK-----VQ-LD--
ref|WP_015187990.1|   --------------------------------M-K-P--------NYRRCISCRRVALKHEFWRIVRV-----Y-P---------SGE-----LQ-LD--
gb|NET06949.1|        -----------------------MKP-------------------NYRRCLSCRTILPKEAFWRVVRV-----H-P---------SRN-----VQ-LD--
ref|WP_224409244.1|   --------------------------------M----------KTNYRRCLSCRRVGPKDSFWRIVRV-----Y-P---------SRQ-----VQ-LD--
ref|WP_155748928.1|   --------------------------------M-K-P--------NYRRCISCRKVGLKQEFWRIVRV-----F-P---------SGQ-----VQ-LD--
ref|WP_229547107.1|   --------------------------------M--KP--------NYRRCISCRKVGSKDEFWRIVRV-----F-P---------SGK-----VQ-LD--
gb|MDW8401950.1|      --------------------------MP--------PG--------WRRCLSCRTVGPREQFWRVVRL-----H-P---------SH---R--VV-LDQG
ref|WP_107668609.1|   --M---------------------KP-------------------NYRRCISCRRVAPKEAFWRIVRV-----Y-P---------SRQ-----VQ-LD--
ref|WP_008232964.1|   --------------------------------M--KA--------NYRRCISCRKIGLKQEFWRVIRV-----F-P---------SRQ-----VQ-LD--
gb|MEC4893587.1|      --------------------------------M----------KTNYRRCLSCRRVGPKDSFWRIVRV-----Y-P---------SRQ-----VQ-LD--
ref|WP_109009916.1|   --------------------------------M--KP--------NYRRCISCRKVGSKDEFWRIVRV-----F-P---------SGK-----VQ-LD--
gb|MEC4816215.1|      --------------------------------M-K-P--------NYRRCISCRKVGLKQEFWRIVRV-----F-P---------SGQ-----VQ-LD--
gb|MBW4486969.1|      -----------------------M-----------KP--------NYRRCTSCRKVAPKEDFWRIVRV-----H-P---------SQT-----VQ-LD--
tpg|HZG41009.1|       ---------------------------------------M---QPNYRRCISCRRVGPKTEFWRLVRV-----Y-P---------DRT-----VQ-LG--
ref|WP_322643731.1|   --------------------------------M--KP--------NYRRCISCRRVGSKDEFWRIVRV-----F-P---------SGK-----VQ-LD--
ref|WP_118168122.1|   --------------------------------M--KP--------NYRRCISCRKVGSKDEFWRIVRV-----F-P---------SGK-----VQ-LD--
ref|WP_194042652.1|   --------------------------------M--KP--------NYRRCISCRKVGSKDEFWRIVRV-----F-P---------DGK-----VQ-LD--
ref|WP_086688430.1|   --------------------------------M--KP--------NYRRCISCRQVALREEFWRIVRV-----F-P---------SGT-----VQ-LD--
gb|MEB3161329.1|      ------------------MP----------------PG--------FRRCLSCRKVAHRREFWRIVRV-----Y-P---------SR------TVQLDHG
gb|MBF2099090.1|      -----------------------M-----------RP--------GYRRCLSCRRLAPRQEFWRVVRL-----Q-G---------THQ-----VQ-LD--
gb|MBE8968595.1|      --------------------------------M--KP--------NYRRCISCRKVGSKDEFWRIVRV-----F-P---------DGK-----VQ-LD--
ref|WP_152589415.1|   --------------------------------M--KP--------NYRRCISCRKVGSKDEFWRIVRV-----F-P---------SGK-----VQ-LD--
gb|MCT7960255.1|      --------------------------------M--KP--------NYRRCISCRKVAPKEAFWRIVKL-----Y-P---------SDQ-----VQ-LD--
```

```
gb|MBW4576792.1|      -----------------------------------ME-------PNYRRCVSCRQLAPKEVFWRIVRV-----Y-P---------SQQ-----VQ-LD--
ref|WP_233748074.1|   --------------------------------M--EP--------NYRRCIACRKIAHKREFWRIVRV-----H-P---------SQQ-----VQ-IN--
gb|MDJ0730709.1|      -----------------------MKP-------------------NYRRCLSCGRVAPKEAFWRIVRV-----Y-P---------SRQ-----VQ-LD--
gb|NEO25536.1|        ---------------------------------------M---KPNFRRCLACRRVAPKSSFWRIVRV-----H-P---------SRK-----VQ-LD--
ref|WP_190559635.1|   --------------------------------M--KP--------NYRRCISCRRVGSKEEFWRIVRI-----F-P---------SGK-----VQ-LD--
ref|WP_190600959.1|   --------------------------------M--KP--------NYRRCISCRQVGSKEEFWRIVRV-----F-P---------SGK-----VQ-LD--
ref|WP_289796853.1|   --------------------------------M--KP--------NYRRCISCRKVGSKDEFWRIVRV-----F-P---------DGK-----VQ-LD--
ref|WP_194057957.1|   -----------------MPP-------------------------NYRRCVSCRRVGPKTEFWRLVRVHP---------------DRT-----VQ-LG--
ref|WP_016866716.1|   --------------------------------M-K-P--------NYRRCISCRKVGLKNEFWRIVRV-----F-P---------SGQ-----VQ-LD--
gb|MCS6814375.1|      -----------------------M-----------EP--------NYRRCVSCRRVALKADFWRVVRV-----Y-P---------SQA-----VQ-LD--
tpg|HBB35302.1|       ------------------------------------------MEPNYRRCLSCRKVAPKQAFWRVVRV-----H-P---------SRQ-----VK-LD--
gb|MDJ0715684.1|      --------------------------------M--KP--------NYRRCISCRQVAPKESFWRIVRV-----Y-T---------SAE-----VQ-LD--
gb|NEP12133.1|        --------------------------------M--KP--------NYRRCLSCRQVAPKEGFWRVVRV-----Y-P---------SRK-----VQ-LD--
gb|NJP17789.1|        ------------------------------------------MTKNYRRCISCRKIASKETFWRIVRV-----Y-P---------SRQ-----VQ-LD--
gb|KAB8319469.1|      --------------------------------M-K-P--------NYRRCISCRQVGLKQEFWRIVRV-----F-P---------SGQ-----VQ-LD--
ref|WP_190413342.1|   ------------------------------------------MEPNYRRCVSCRQVAPKSAFWRIVRV-----Y-P---------SQQ-----LQ-LD--
gb|MBR8834127.1|      --------------------------------M-K-P--------NYRRCISCRKVNPKQEFWRIVRV-----F-P---------SGQ-----VQ-LD--
gb|MCC5607216.1|      --------------------------------M--KP--------NYRRCISCRKVGSKDEFWRIVRV-----F-P---------SGK-----VQ-LD--
ref|WP_168729992.1|   --------------------------------M--KP--------NYRRCISCRRVGTKTEFWRIVRL-----F-P---------SGK-----VE-LD--
ref|WP_280654349.1|   --------------------------------M--KP--------NYRRCISCRQVGLKQDFWRIVRV-----F-P---------SGK-----VQ-LD--
gb|MBD2016732.1|      ---------------------------------------M---EPNYRRCLSCRLVAPKKAFWRVVRL-----Y-P---------SRQ-----VQ-LD--
ref|WP_099069649.1|   --------------------------------M--KP--------NYRRCISCRKLGLKEEFWRIVRV-----F-P---------SGK-----VQ-LD--
ref|WP_190533136.1|   ------------------------------------------MEPNYRRCVSCRQVAPKSAFWRIVRV-----Y-P---------SQQ-----LQ-LD--
gb|MCU0541201.1|      MP-----------------DARYPIPDTRCPMPH-TPQQM---KPNYRRCASCRKIAPKEAFWRAVRV-----Y-P---------SHQ-----VQ-LD--
ref|WP_193872423.1|   --------------------------------M--KP--------NYRRCLSCRKVAPKQAFWRVVRV-----Y-P---------SRQ-----VQ-LD--
ref|WP_061545594.1|   --------------------------------M--KP--------NYRRCISCRRVGTKTEFWRIVRL-----F-P---------SGK-----VE-LD--
ref|WP_102179423.1|   --------------------------------M-K-P--------NYRRCISCRKVGLKNEFWRIVRV-----F-P---------SGQ-----VQ-LD--
tpg|HLP90034.1|       --------------------------------M-K-P--------NYRRCISCRQVSPKEEFWRIVRV-----F-P---------SGK-----VQ-LD--
ref|WP_190956491.1|   --------------------------------M--KP--------NYRRCISCRKLGLKEEFWRIVRV-----F-P---------SGK-----VQ-LD--
ref|WP_102175048.1|   --------------------------------M-K-P--------NYRRCISCRKVGLKNEFWRIVRV-----F-P---------SGQ-----VQ-LD--
gb|NJO42002.1|        --------------------------------M--LP--------NYRRCVSCRKIAPKEEFWRVVRV-----F-P---------SQT-----VQ-LD--
ref|WP_006275620.1|   --------------------------------M--KP--------NYRRCISCRRVGTKTEFWRIVRL-----F-P---------SGK-----VE-LD--
gb|MBU7582629.1|      --------------------------------M--KP--------NYRRCISCRKVGLKQDFWRIVRV-----F-P---------SGK-----VQ-LD--
gb|MCG6133583.1|      --------------------------------M--KP--------NYRRCISCRKVGLKEDFWRIVRV-----F-P---------SGK-----VQ-LD--
ref|WP_016872536.1|   --------------------------------M--KP--------NYRRCLSCRKVGLKNEFWRIVRV-----F-G---------SGQ-----VQ-LD--
ref|WP_015117518.1|   --------------------------------M-K-P--------NWRRCISCRKVSLKSEFWRIVRV-----F-P---------SGK-----VQ-LD--
tpg|HBE20201.1|       --------------------------------M----------KPNYRRCLSCRQVGLKQVFWRVVRV-----Y-P---------SGK-----VQ-LD--
gb|MBW4500045.1|      --------------------------------M-K-P--------NYRRCMSCRQVGLKQEFWRIVRV-----F-P---------SGQ-----VQ-LD--
ref|WP_103135735.1|   -----------------------M-----------KP--------NYRRCVSCRKAGLKEEFWRIVRV-----F-P---------SGN-----VQ-LN--
ref|WP_105221837.1|   --------------------------------M-K-P--------NYRRCISCRKVALKHEFWQIVRV-----Y-P---------SGE-----LQ-LD--
ref|WP_190434246.1|   -----------------------M-----------KP--------NYRRCASCRKLASKQDFWRIVRV-----H-S---------SQT-----VQ-LD--
ref|WP_283760063.1|   --------------------------------M-K-P--------NQRRCLSCRRVDSKEAFWRIVRV-----Y-P---------DRT-----IQ-LD--
gb|MDZ8167634.1|      --------------------------------M--KP--------NYRRCISCRKLGLKEEFWRIVRV-----F-P---------SGK-----VQ-LD--
ref|WP_193879029.1|   --------------------------------M--KP--------NYRRCISCRKVGLKQDFWRIVRV-----F-P---------SGK-----VQ-LD--
ref|WP_190591144.1|   --------------------------------M--KP--------NYRRCISCRKVGLKQDFWRIIRV-----F-P---------SGK-----VQ-LD--
gb|MDY6781740.1|      ---------------------------------------M---KPNYRRCISCRKVAPKENFWRIVRD-----S-S---------SKT-----VQ-LD--
gb|MDZ8106853.1|      --------------------------------M--KP--------NYRRCISCRKLGLKEEFWRIVRV-----F-P---------SGK-----VQ-LD--
ref|WP_013325296.1|   ---------------------------------------M---KPNTRRCVSCRQIATKDNFLRIVRV-----Y-P---------SRQ-----VQ-LD--
ref|WP_171573341.1|   --------------------------------M--KP--------SYRRCVACRKAALKVDFWRIVRV-----H-P---------SQE-----VH-LD--
ref|WP_190938724.1|   --------------------------------M--KP--------NSRRCISCRKVGSKDQFWRIIRV-----F-P---------SGK-----VQ-LD--
ref|WP_086768880.1|   --------------------------------M--KP--------NYRRCISCRQLALREEFWRIVRV-----F-P---------SGT-----VQ-LD--
ref|WP_251960204.1|   --------------------------------M--KP--------NYRRCISCRKVGSKDEFWRIVRV-----F-P---------DGK-----VQ-LD--
ref|WP_009545653.1|   ------------------------------------------MKPNYRRCISCRRVAPKEAFWRIVRV-----Y-P---------SRQ-----VQ-LD--
```

```
ref|WP_127085004.1|   ---------MK-PN--------------------------------YRRCLSCRKIGLKEEFWRIVRV-----F-P---------SG---Q--VQ-LNEG
tpg|HIK08066.1|       -----------------------M-----------TP--------NYRRCVSCRKVGLKEDFWRIVRV-----F-P---------SGN-----VQ-LN--
gb|MDZ7979263.1|      --------------------------------M--KP--------NYRRCISCRQVSLKEEFWRIVRV-----F-P---------SGT-----VQ-LD--
gb|MCC5635819.1|      --------------------------------M--KP--------NYRRCISCRKVGLKQDFWRIVRV-----F-P---------SGK-----VQ-LD--
ref|WP_239731788.1|   --------------------------------M--KP--------NYRRCISCRKVGLKEDFWRIVRV-----F-P---------SGK-----VQ-LD--
ref|WP_193897322.1|   --------------------------------M--KP--------NYRRCISCRKVGLKQDFWRIIRV-----F-P---------SGK-----VQ-LD--
ref|WP_187706878.1|   --------------------------------M--KP--------NYRRCISCRRVGTKTEFWRIVRL-----F-P---------SGK-----VE-LD--
ref|WP_214438763.1|   --------------------------------M--KP--------NYRRCISCRKVGLREDFWRIVRV-----F-P---------SGK-----VQ-LD--
ref|WP_073632737.1|   --------------------------------M-K-P--------NYRRCMSCRQVGLKQEFWRIVRV-----F-P---------SGQ-----VQ-LD--
gb|MDZ8241182.1|      --------------------------------M--KP--------NYRRCISCRKLGLKEEFWRIVRV-----F-P---------SGK-----VQ-LD--
ref|WP_073548075.1|   --------------------------------M-K-P--------NYRRCISCRRVALKHEFWRIVRV-----Y-P---------SGE-----LQ-LD--
gb|MBD2163199.1|      --------------------------------M--KP--------NYRRCISCRQVGLKEEFWRIVRV-----F-P---------SGK-----VQ-LD--
gb|MBO1346139.1|      --------------------------------M----------LPNYRRCISCRKVASKEAFWRIVRV-----Y-P---------SGQ-----LQ-LD--
gb|MBW4600386.1|      --------------------------------M-K-P--------NYRRCLSCRKIGLKEDFWRIVRV-----F-P---------SGQ-----VQ-LD--
gb|MBC6453437.1|      --------------------------------M----------LPNYRRCISCRKVASKEAFWRIVRV-----Y-P---------SGQ-----LQ-LD--
ref|WP_289789294.1|   --------------------------------M--KP--------NHRRCLSCRKVGLKNEFWRIVRV-----F-E---------SGQ-----VQ-LD--
ref|WP_167722165.1|   --------------------------------M--KP--------NYRRCISCRQVGLKEEFWRIVRV-----F-P---------SGK-----VQ-LD--
ref|WP_095719821.1|   --------------------------------M-E-P--------NYRRCISCRKVGLKQEFWRIVRV-----F-P---------SGQ-----VQ-LD--
ref|WP_096644639.1|   --------------------------------M--KP--------NYRRCISCRRVALKEEFWRIVRV-----F-E---------SRT-----VQ-LD--
gb|NJM70351.1|        --------------------------------M--KP--------HYRRCISCRKIGLKQEFWRIVRV-----F-P---------SGQ-----VQ-LD--
ref|WP_082127362.1|   --------------------------------M-E-P--------NYRRCISCRKIGLKHEFWRVVRV-----F-P---------SMQ-----VQ-LD--
ref|WP_073610176.1|   --------------------------------M--QP--------NYRRCVSCRRVGPKADFWRVVRV-----Y-P---------DRS-----VR-LS--
gb|MDZ8028666.1|      --------------------------------M--KP--------NYRRCISCRKLGLKEEFWRIVRV-----F-P---------SGK-----VQ-LD--
gb|MDZ8019070.1|      --------------------------------M--KP--------NYRRCISCRKLGLKEEFWRIVRV-----F-P---------SGK-----VQ-LD--
ref|WP_017742689.1|   --------------------------------M--KP--------NYRRCISCRKVGLKQEFWRIVRV-----F-P---------SGQ-----VQ-LD--
ref|WP_045868281.1|   --------------------------------M--KP--------NYRRCVSCRQVGLKEEFWRIVRV-----F-P---------SGK-----VQ-LD--
ref|WP_208344959.1|   --------------------------------M----------KPNQRRCISCRKVSTKQEFWRIVRV-----F-P---------SGQ-----VQ-LD--
gb|MBW4560223.1|      --------------------------------M--KP--------NYRRCISCRQVGLKEEFWRIVRV-----F-P---------SGK-----VQ-LD--
gb|PSB49245.1|        ------------------------------------------MKPNYRCCASCRKIADKEAFWRVVRV-----Y-P---------SHQ-----VQ-LD--
ref|WP_038087798.1|   --------------------------------M-K-P--------NYRRCISCRQVGLKQEFWRIVRV-----F-P---------SGQ-----VQ-LD--
ref|WP_190427456.1|   ------------------------------------------MEPNYRRCVSCRQVAPKSAFWRIVRV-----Y-P---------SQQ-----LQ-LD--
ref|WP_261224160.1|   --------------------------------M-K-P--------NLRCCVSCRRIADKKEFWRIVRV-----Y-P---------SHQ-----VQ-LD--
gb|ARV61523.1|        --------------------------------M--KL--------NYRRCISCRKVGLKQEFWRIVRV-----F-P---------SGQ-----VQ-LD--
gb|MCU0523589.1|      -----------------------MHP-------------------NYRRCVSCRKVDHKSAFWRVVRV-----F-P---------SQE-----VQ-LD--
ref|WP_089091493.1|   --------------------------------M--KP--------NYRRCISCRKVGLKQDFWRIVRV-----F-P---------SGK-----VQ-LD--
gb|MBF2080108.1|      -----------------MQP-------------------------NYRRCVSCRRVAPKAEFWRIVRVHP---------------SRT-----VQ-LD--
ref|WP_268609846.1|   --------------------------------M----------PPHYRRCVSCRRVAPKQSFWRIVRV-----F-S---------SGT-----VQ-LD--
gb|MBC6418410.1|      --------------------------------M--KTN--------YRRCISCRKVGAKESFWRIVRV-----Y-L---------SRE-----V------
ref|WP_200324434.1|   --------------------------------M---K-------PNYRRCISCRRVGLKEEFWRIVRV-----F-P---------SGK-----VQ-LE--
ref|WP_015202566.1|   --------------------------------M-E-P--------NFRRCVSCRQAAPKVEFWRIVRV-----Y-P---------SQQ-----VQ-LD--
ref|WP_194023660.1|   ---------------------------------------M---QPNYRRCASCRRVGPKAEFWRLVRV-----Y-P---------DRT-----VC-LG--
dbj|BBD59214.1|       --------------------------------M--KP--------NYRRCISCRKVGLKQDFWRIVRV-----F-P---------SGK-----VQ-LD--
ref|WP_096579096.1|   --------------------------------M--KP--------NYRRCISCRKVGLKQDFWRIVRV-----F-P---------SGK-----VQ-LD--
ref|WP_315785635.1|   --------------------------------M-K-P--------NYRRCISCRKVGLKNEFWRITRV-----F-P---------SGQ-----VQ-LD--
gb|MDP8965164.1|      ------------------------------------------MEPNYRRCLICRKVAPKAAFWRVVRV-----Y-P---------SRQ-----VQ-LD--
ref|WP_015194852.1|   --------------------------------M--QP--------NYRRCLSCRKIAPKNSFLRIVRI-----Y-P---------SHQ-----IK-LE--
ref|WP_006528507.1|   --------------------------------M----------KTNYRRCLSCRCVGPKESFWRIVRV-----Y-P---------STE-----VQ-LD--
gb|MBC6423737.1|      --------------------------------M----------LPNYRRCISCRKVASKEAFWRIVRV-----Y-P---------SGQ-----LQ-LD--
gb|MBW4517440.1|      ------------------------------------------MLPNYRRCISCRKISPKEGFWRIVRV-----F-P---------SHI-----VQ-LE--
ref|WP_102221303.1|   --------------------------------M-K-P--------NYRRCISCRKVGLKNEFWRIVRV-----F-P---------SGQ-----VQ-LD--
gb|MBE9051640.1|      --------------------------------M--KP--------NYRRCISCRKVGLKEDFWRIVRV-----F-P---------SGK-----VQ-LD--
gb|MEB3268310.1|      -------------------M---------------EP--------NYRRCVSCRTVAPKHQFWRIVRH-----H-D---------TGT-----VQ-LD--
dbj|BAY27136.1|       --------------------------------M--KP--------NYRRCISCRRVALKEEFWRIVRV-----F-E---------SRT-----VQ-LD--
```

```
ref|WP_190744460.1|   --------------------------------M--KP--------NYRRCISCRRVALKEEFWRIVRV-----F-E---------SRT-----VQ-LD--
gb|MCS6791070.1|      --------------------------------M-KKP--------NQRRCISCYKVADKKEFWRVIRL-----Y-P---------SHQ-----VQ-LD--
ref|WP_015144681.1|   ------------------------------------------MQKNYRRCISCRKVAPKESFWRIVRV-----Y-P---------SRQ-----VQ-LD--
tpg|HYW19585.1|       --------------------------------M--KP--------NYRRCISCRKVGLKEDFWGIVRV-----F-P---------SGK-----VQ-LD--
gb|MCJ8278778.1|      -----------------------M-----------KP--------NWRRCISCRKVSLKHEFWRIVRV-----F-P---------SRK-----VQ-LN--
gb|RMF20653.1|        --------------------------------M--KP--------NYRRCISCRRIAPKVDFWRVVRV-----Y-P---------TQT-----VQ-LD--
gb|NRB09489.1|        --------------------------------M--KA--------NYRRCISCRKIGFKQEFWRIVRV-----F-P---------SGQ-----VQ-LD--
gb|MDJ0736158.1|      --------------------------------M--EV--------NYRRCISCRRVGLKQEFWRIVRV-----F-P---------SGQ-----VQ-LD--
gb|NJL87205.1|        --------------------------------M--KP--------NYRRCISCRQIAPRENLWRVVRV-----H-P---------THQ-----VQ-LD--
ref|WP_261207255.1|   --------------------------------M--KP--------NYRRCISCRKVAPKEAFWRIVKL-----Y-P---------SDQ-----VQ-LD--
ref|WP_008274651.1|   ------------------------------------MPEKSVIKPNYRRCISCRRVAPKEAFWRIVRV-----Y-P---------SRQ-----VQ-LD--
gb|MDJ0845508.1|      --M---------------------KP-------------------NYRRCISCGRIAPKEAFWRIVRV-----Y-P---------SRQ-----VQ-LD--
ref|WP_062247903.1|   --------------------------------M-K-P--------NYRRCISCHKVGLKNEFWRIVRV-----F-P---------SGQ-----VQ-LD--
gb|MDF5707201.1|      --------------------------------M--KP--------NYRRCISCRKLGLKDEFWRIVRV-----F-P---------SGK-----VQ-LD--
ref|WP_190555052.1|   -----------------------M-----------KP--------NYRRCASCRKLAPKEDFWRIVRV-----H-S---------SQT-----VQ-LD--
gb|TAD81818.1|        ------------------------------------------MKPNYRCCASCRKIATKEAFWRVVRV-----Y-P---------SHQ-----VQ-LD--
gb|MBF2035119.1|      ---------------------------------------M---KPNLRCCVSCRKIADKEDFWRIVRV-----Y-S---------TRT-----VQ-LD--
gb|MCY7283860.1|      ------------------------------------------MLPNYRRCISCRKISPKEGFWRIVRV-----F-P---------SQA-----VQ-LE--
ref|WP_106289357.1|   ---------------------------------------M---SKNYRRCISCRHLAVKKSFWRIVRV-----Y-P---------SYQ-----VQ-LD--
ref|WP_200989746.1|   --------------------------------M--KP--------NYRRCISCRKVGLKEDFWRIVRV-----F-P---------SGK-----VQ-LD--
ref|WP_026079831.1|   -------------------M---------------KP--------NVRRCVACRKVAPKTSFWRIVRL-----Y-P---------SQT-----VQ-LD--
ref|WP_196521436.1|   --------------------------------M--KP--------NYRRCISCRKVALKDEFWRIVRV-----F-P---------SGK-----VQ-LD--
gb|NJK27659.1|        --------------------------------M----------EPNQRRCISCRRIGSKDSFWRVVRV-----Y-P---------SHQ-----VQ-LD--
ref|WP_297080144.1|   -----------------------------MP----PP--------NYRRCLSCRRLAPKTEFWRIVRV-----Y-P---------SQT-----IQ-LD--
gb|NJL10525.1|        --------------------------------M--KP--------NYRCCISCRKIALKQEFWRVVRV-----F-S---------SGQ-----VQ-LD--
ref|WP_124976615.1|   --------------------------------M-K-P--------NYRRCLSCRCLAPKEAFWRIVRV-----S-P---------SRQ-----VQ-LD--
gb|MBW4673665.1|      --------------------------------M--KP--------NYRRCISCRKLGLKEEFWRIVRV-----F-P---------SGK-----VQ-LD--
ref|WP_261198198.1|   --------------------------------M--EP--------NYRRCISCRKVAPKEAFWRIVKL-----Y-P---------SDQ-----VQ-LD--
ref|WP_015128246.1|   --------------------------------M--KP--------NYRRCISCRRVGLKEEFWRIVRV-----F-P---------SGK-----VQ-LD--
gb|TAF21607.1|        --------------------------------M--KP--------NYRRCLSCRRVGLKQEFWRIVRV-----F-P---------SRK-----VQ-LD--
ref|WP_009458837.1|   --------------------------------M-K-P--------NYRRCISCRKVGLKNEFWRIVRV-----F-P---------SGQ-----VQ-LD--
ref|WP_190980837.1|   --------------------------------M--KP--------NYRRCISCREVGLKQDFWRIVRV-----F-P---------SGK-----VQ-LD--
ref|WP_283761350.1|   ---------MK-------------------------PN--------QRRCVSCRKVDRKQAFWRIVRV-----Y-P---------DD---E--IQ-LDEG
gb|MBU6184880.1|      --------------------------------M----G------IHYRRCISCRKVAPKDFFWRVVRL-----Y-P---------SNR-----V------
ref|WP_193197802.1|   --------------------------------M-K-P--------NYRRCTSCRKVGLKEEFWRIVRV-----F-P---------SGK-----VQ-LD--
ref|WP_015114806.1|   --------------------------------M--KP--------NYRRCISCRKVGLKQDFWRIVRV-----F-P---------SGK-----VQ-LD--
ref|WP_011318669.1|   -----------------------M-----------QP--------NYRRCISCRKVGLKQDFWRIVRV-----F-P---------SGK-----VQ-LN--
gb|MBC6477661.1|      --------------------------------M----------LPNFRRCISCRKVASKEAFWRIVRV-----Y-P---------SGQ-----LQ-LD--
gb|NDJ22983.1|        --------------------------------M-K-P--------NYRRCTSCRKVGLKEEFWRIVRV-----F-P---------SGK-----VQ-LD--
gb|RAM50821.1|        --------------------------------M--KP--------NYRRCISCRKVGLKNEFWRIVRV-----F-P---------SGQ-----VQ-LD--
gb|MCZ0898804.1|      ------------------------------------------MKPNYRCCASCRKIAPKEAFWRVVRL-----Y-P---------SHQ-----VQ-LD--
gb|MBW4554427.1|      --------------------------------M--KP--------NYRRCISCRKVGLKDDFWRIVRV-----F-P---------SGK-----VQ-LD--
ref|WP_179047786.1|   --------------------------------M--QP--------NYRRCISCRKVGLKQDFWRIVRV-----F-P---------SGK-----VQ-LN--
ref|WP_071599862.1|   --------------------------------M-K-P--------NNRRCISCRKVGLKQEFWRIVRV-----F-P---------SGQ-----VQ-LD--
ref|WP_207087020.1|   --------------------------------M--KP--------NYRRCISCRKVAPKEAFWRIVKL-----Y-P---------SDQ-----VQ-LD--
gb|MDZ8042175.1|      --------------------------------M--KP--------NYRRCISCRKLGLKEEFWRIVRV-----F-P---------SGK-----VQ-LD--
ref|WP_102207255.1|   --------------------------------M-K-P--------NYRRCISCRKVDLKNEFWRIVRV-----F-P---------SGQ-----VQ-LD--
ref|WP_066384474.1|   --------------------------------M--KP--------NYRRCVSCRKVGLKKDFWRIVRV-----F-P---------SGE-----VQ-LN--
gb|PSB20742.1|        --------------------------------M--PS--------NYRRCVSCRKIAPKDSFWRVVRV-----Y-P---------SGT-----VQ-LD--
ref|WP_190697022.1|   --------------------------------M--KP--------NYRRCLSCRKVALKEDFWRIVRV-----F-P---------SGK-----VQ-LD--
gb|MCT7955770.1|      --------------------------------M--KP--------NYRRCISCRKVAPKEAFWRIVKL-----Y-P---------SDQ-----VQ-LD--
ref|WP_254010714.1|   --------------------------------M----------EPNYRRCLSCRKVAPKQAFWRVVRV-----Y-P---------SGE-----VQ-LD--
gb|NBD15363.1|        -----------------------MKK-------------------NYRRCLSCRRIAPKSEFLRIVRV-----H-P---------TKT-----IQ-LD--
```

```
ref|WP_283755094.1|  --------------------------------M--KP--------NQRRCVSCRKVDDKEAFWRIVRV-----Y-P---------DHH-----IQ-LD--
gb|MDJ0534388.1|     -----------------MK-----------------P--------NYRRCVSCRQILPKEQLWRIVRV-----H-P---------SHS-----IV-LD--
gb|MBW4482298.1|     --------------------------------M--QPN--------YRRCISCRRVGPKAEFWRVVRV-----Y-P---------DRT-----VR-LG--
gb|MBD3880571.1|     --------------------------------M--SS--------NYRRCVSCRRVAPKTDFWRVVRV-----F-P---------TRT-----IQ-LD--
ref|WP_271731773.1|  --------------------------------M--KP--------NYRRCISCRQVGLKEDFWRVVRV-----F-P---------SGK-----VQ-LD--
gb|MBW4634383.1|     --------------------------------M-K-P--------NYRRCISCRKVGLKQEFWRVVHV-----F-P---------SGQ-----VQ-LD--
ref|WP_193999244.1|  --------------------------------M--KP--------NYRRCTSCRKVGLKEEFWRIVRV-----F-P---------SGK-----VQ-LD--
gb|NJN72043.1|       --------------------------------M--E--------PNLRRCVVCRQIAPKQIFWRIVRV-----Y-P---------SQT-----VQ-LD--
ref|WP_016950665.1|  --------------------------------M--KP--------NYRRCISCRRVGLKEEFWRIVRV-----F-P---------SGK-----VQ-LD--
ref|WP_191758834.1|  --------------------------------M-K-P--------NYRRCTSCRKVGLKEEFWRIVRV-----F-P---------SGK-----VQ-LD--
gb|MBD2237840.1|     --------------------------------M--KP--------NYRRCVSCRQVGLKEEFWRIVRV-----F-P---------SGK-----VQ-LD--
ref|WP_283767879.1|  ---------ME------------------------PN---------QRRCVSCRKVDRKQAFWRIVRV-----Y-P---------DD---E--IQ-LDEG
ref|WP_172358412.1|  -----------------------------MS----PP--------NYRRCLSCRRLAPKTEFWRIVRV-----Y-P---------SQT-----IQ-LD--
gb|MCP6758672.1|     --------------------------------M--KP--------NYRRCISCRKVDLKNEFWRIVRV-----F-P---------SGQ-----VQ-LD--
ref|WP_015228055.1|  --M---------------------EK-------------------NYRRCLSCRRIAPKEEFLRIVRV-----H-P---------SRK-----IQ-LD--
ref|WP_015210357.1|  --------------------------------M-K-P--------NYRRCISCRRVGFKEEFWRIVRV-----F-P---------SGK-----VQ-LN--
ref|WP_104545615.1|  --------------------------------M-K-P--------NYRRCISCRRVALKHEFWRIVRV-----Y-P---------SGE-----LQ-LD--
ref|WP_237991081.1|  -----------------------M-----------KP--------NYRSCVSCRKVGLKEEFWRIVRV-----F-P---------SGK-----VQ-LN--
ref|WP_190882003.1|  --------------------------------M--KP--------NYRRCISCRRLGLKEEFWRIVRV-----F-P---------SGK-----VQ-LD--
gb|MBF2073651.1|     ------------------------------------------MLPNYRRCISCRRVAHKSAFWRVVRV-----F-P---------SQQ-----VQ-LD--
ref|WP_323312969.1|  --------------------------------M-K-P--------NYRRCISCRRIGLKEEFWRIVRV-----F-P---------SGK-----VQ-LD--
gb|MBF2001555.1|     ------------------------------------------MLPNYRRCISCRRVAHKSAFWRVVRV-----F-P---------SQQ-----VQ-LD--
ref|WP_193967039.1|  -----------------MTP-------------------------NYRRCVSCRRVAPKEAFWRIVRVYD---------------DRS-----VC-LD--
ref|WP_190462952.1|  --------------------------------M--KP--------NYRRCISCRRVGLKEEFWRIVRV-----F-P---------SGK-----VQ-LN--
gb|MBV6625219.1|     --------------------------------M-K-P--------NWRRCISCRKVSLKSEFWRIVRV-----F-P---------SGK-----VQ-LD--
ref|WP_265262465.1|  -------------------M---------------KP--------NVRRCVACREVAPKTSLWRIVRL-----Y-P---------SQT-----VQ-LD--
ref|WP_281485672.1|  --------------------------------M--KP--------NYRRCISCRKVGLKQDFWRIVRV-----F-P---------SGK-----VQ-LD--
ref|WP_127053480.1|  --------------------------------M--KP--------NYRRCISCRRVGLKEEFWRIVRV-----F-P---------SGK-----VQ-LD--
tpg|HAC62310.1|      -----------------------------MK-P--N----------YRRCISCRKLALKQDFWRIVRL-----Y-P---------SYE-----VK-LD--
ref|WP_277772165.1|  -----------------------------MS----PP--------NYRRCLSCRRLAPKTEFWRIVRV-----Y-P---------SQT-----IQ-LD--
ref|WP_190570712.1|  -----------------------M-----------QP--------NYRRCISCRKVGLKQDFWRIVRV-----F-P---------SGK-----VQ-LN--
tpg|HIK13958.1|      -----------------MKP-------------------------NYRRCVSCRKVGPKSEFWRLVRQHP---------------SHT-----IQ-LD--
ref|WP_169264435.1|  --------------------------------M--KP--------NYRRCISCRKTRLKQEFWRIVRV-----F-P---------SGQ-----VQ-LD--
ref|WP_106455349.1|  -----------------------------------MP-------INIRRCISCHKIAPKTSFWRIVRV-----Y-P---------SGQ-----VQ-LD--
ref|WP_044206893.1|  ---------------------------------------M---EPNYRRCLSCRRVAQKQALWRVVRI-----H-P---------SRQ-----VK-LD--
ref|WP_071188254.1|  --------------------------------M--KP--------NYRRCISCRRVGLKEEFWRIVRV-----F-P---------SGK-----VQ-LD--
gb|MEC4803202.1|     --------------------------------M--KP--------NLRRCVSCRRLASKQDFWRIVRV-----H-P---------SRQ-----VQ-LD--
ref|WP_015135874.1|  -------------------------------MV--KT--------NHRRCVACRKLAPKEYFWRVVRV-----Y-P---------SQA-----VQ-LD--
gb|NER81563.1|       -----------------------MKP-------------------NHRRCVSCRAIAPKEAFWRIVRV-----Y-P---------TQT-----VQ-LD--
ref|WP_247215999.1|  --------------------------MP-------PG---------TRRCLSCRTLAPRERFWRVVRL-----F-P---------TH---Q--VV-LDEG
dbj|BAY28476.1|      --------------------------------M--KP--------NYRRCVSCRQVALKEEFWRIVRV-----F-P---------SGK-----VQ-LD--
gb|MCL6749841.1|     ---------------------------------------------NYRRCISCRKVGSKDEFWRIVRV-----F-P---------SGK-----VQ-LD--
ref|WP_190685510.1|  --------------------------------M--KP--------NYRRCISCRKVALKQDFWRIIRV-----F-P---------SGK-----VQ-LD--
gb|MBF2065700.1|     --------------------------------M---K-------PNYRRCLSCRKIALKQEFWRIVRV-----F-P---------SGQ-----VQ-LD--
ref|WP_010997971.1|  -----------------------M-----------QP--------NYRRCISCRKVGLKQDFWRIVRV-----F-P---------SGK-----VQ-LN--
gb|MBD2409277.1|     --------------------------------M--KP--------NYRRCISCRQLGLKEEFWRIVRV-----F-P---------SGK-----VQ-LD--
ref|WP_006196598.1|  --------------------------------M--EP--------NYRRCISCRKVGLKEDFWRIVRV-----F-P---------SGK-----VQ-LD--
ref|WP_088239620.1|  --------------------------------M---K-------PNFRRCISCRRIGIKQEFWRIVRV-----F-P---------SGQ-----VQ-LD--
gb|MBP5972342.1|     --------------------------------M--KP--------NYRRCISCRKTRLKQEFWRIVRV-----F-P---------SGQ-----VQ-LD--
ref|WP_193997393.1|  --------------------------------M--KP--------NYRRCISCRKVGLKEDFWRIVRV-----F-P---------SGK-----VQ-LD--
tpg|HLO50431.1|      MSFELHPQNSGRAQKFRPMITLFPLPSSFFLLP--SSFQM---KTNYRRCISCRKLGPKEAFWRIVRL-----Y-P---------CHQ-----VQ-LD--
ref|WP_146296814.1|  --------------------------------M--KKN--------YRRCISCRRVAPKEEFLRVVRT-----H-P---------SRT-----VK-LN--
gb|MBW4616862.1|     --------------------------------M-K-P--------NYRRCISCRQVGLKEEFWRIVRV-----F-P---------SGK-----VQ-LE--
```

```
ref|WP_323360326.1|   --------------------------------M--KP--------NYRRCLSCRRVGLKEEFWRIIRV-----F-P---------SGK-----VQ-LD--
ref|WP_254564844.1|   --------------------------------M--KP--------NYRRCISCRKVAPKEAFWRIVKL-----Y-P---------SDR-----VQ-LD--
gb|MDZ7956785.1|      --------------------------------M--KQ--------NYRRCISCRKVGLKEEFWRIVRV-----F-P---------SGK-----VQ-LD--
gb|NJK29495.1|        -----------------MQP-------------------------NYRRCISCRQVAPKSEFWRIVRVYP---------------DHR-----VQ-LD--
ref|WP_190965760.1|   --------------------------------M--KP--------NYRRCISCRKVGLKQDFWRIVRV-----F-P---------SGK-----VQ-LY--
ref|WP_261235470.1|   MAKDAPGGVDPASVWGKSDSSRRKDKPTHRHLM--EP--------NYRRCISCRKVAPKEAFWRIVKL-----Y-P---------SDQ-----VQ-LD--
ref|WP_261891940.1|   --------------------------------M--QP--------NYRRCISCRQVAPKSELWRIVRV-----Y-P---------DHQ-----IQ-LD--
ref|WP_190435367.1|   --------------------------------M-KPNY---------RRCVSCRKVALKSDFWRVVRL-----Y-P---------SGK-----LQ-LD--
ref|WP_190465083.1|   --------------------------------M-E-P--------NYRRCISCRRVAPKQAFWRIVRV-----Y-P---------SRQ-----VE-LD--
ref|WP_299492025.1|   --------------------------------M--QP--------NYRRCISCRQVAPKSEFWRIVRV-----Y-P---------DHH-----IQ-LD--
gb|MDF2387367.1|      --------------------------------M--KP--------NYRRCISCRKVSLKEDFWRIVRV-----F-P---------SGN-----VQ-LD--
ref|WP_265235232.1|   ------------------------------------------MKLNYRRCCASCRKIGTKEAFWRVVRL-----Y-P---------SHQ-----VQ-LD--
ref|WP_293148638.1|   ------------------------------------------MKPNYRRCCTSCRKIAPKEAFWRVVRL-----Y-P---------SHQ-----VQ-LD--
ref|WP_015151790.1|   --------------------------------M--KP--------NYRRCISCRKVAPKEAFWRIVKL-----Y-P---------SDQ-----VQ-LE--
ref|WP_271764063.1|   --------------------------------M--KP--------NYRRCISCRQVGWKEDFWRVVRV-----F-P---------SGR-----VQ-LD--
ref|WP_190473724.1|   -----------------------M-----------QP--------NYRRCISCRKVGLKQDFWRIVRV-----F-P---------SGN-----VQ-LN--
gb|MEB3313608.1|      --------------------------------M--PP--------NYRRCISCRQVAPKSSLWRVVRR-----H-P---------DGT-----VQ-LD--
gb|TAG88461.1|        ------------------------------------------MKPNYRRCCASCRKIAPKEAFWRVVRL-----Y-P---------SHQ-----VQ-LD--
gb|MBF2015645.1|      --------------------------------M-K-P--------NWRRCISCRKVSLKQEFWRIVRV-----F-P---------SGK-----VQ-LD--
gb|KOP28074.1|        --------------------------------M--KP--------NYRRCISCRKVGLKNEFWRIVRV-----F-S---------SGQ-----VQ-LD--
ref|WP_190516861.1|   --------------------------------M--QP--------NYRRCISCRQVAPKADFWRVVRV-----Y-P---------DRS-----VR-LG--
dbj|BAY09723.1|       --------------------------------M--KP--------NYRRCISCRQVALKEEFWRIVRV-----F-G---------SGT-----VQ-LD--
ref|WP_300634909.1|   -----------------------MKP-------------------NYRRCISCRKVGLKEEFWRIVRV-----F-S---------SRQ-----VQ-LD--
gb|TVQ05455.1|        -----------------MET-------------------------NHRRCVSCRQIAPKAEFWRVVRVYP---------------DRS-----IR-LD--
ref|WP_026732821.1|   --------------------------------M-K-P--------NYRRCISCRKVGLKNEFWRIVRV-----F-G---------SGQ-----VQ-LD--
ref|WP_318699068.1|   ---------MK-------------------------PN--------QRRCVSCRKVDRKEAFWRIVRV-----Y-P---------DR---E--IQ-LDWG
ref|WP_015195815.1|   --------------------------------M---K-------PNYRRCNSCRKVGLKHEFWRIVRV-----F-P---------SGQ-----VQ-LD--
gb|MEB3178814.1|      --------------------------------M----------KPNYRRCISCRRVGLKQEFWRIVRV-----F-P---------SGQ-----VQ-LD--
ref|WP_096832064.1|   ------------------------------------------MKPNYRRCCASCRKIAPKEAFWRVVRL-----Y-P---------SHQ-----VQ-LD--
gb|MBD2774873.1|      --------------------------------M-K-P--------NYRRCISCRKVGLKQEFWRVVRV-----YGE---------SGQ-----LQ-LD--
ref|WP_224341437.1|   --------------------------------M-E-P--------NYRRCVSCRRLAHRSEFWRIVRV-----H-P---------TQA-----VA-LD--
ref|WP_035987080.1|   --------------------------------M-K-P--------NYRRCVSCRRVGPKSDFWRLVRL-----Y-P---------DRT-----VG-LE--
ref|WP_084783000.1|   -----------------------------MP----PP--------NYRRCLSCRRLAPKTEFWRIVRV-----Y-P---------SQT-----IQ-LD--
gb|MBV9387306.1|      --------------------------------M--KP--------NYRRCISCRKVALKQDFWQIVRA-----Y-P---------SGQ-----VQ-LD--
ref|WP_017652092.1|   --------------------------------M--KP--------NYRRCISCRQTALKQEFWRIVRV-----F-P---------SGK-----VQ-LD--
ref|WP_322663056.1|   --------------------------------M--KP--------NYRRCISCRKVSSKEDFWRIVRV-----F-P---------SGK-----VQ-LD--
ref|WP_054468962.1|   --------------------------MK--------P-------N-YRRCVSCRQVFAKHELWRVVRV-------Y----P----SH---Q--VQ-LDR-
gb|MDJ0660438.1|      --M---------------------KP-------------------NYRRCISCGCVAPKGAFWRIVRV-----Y-P---------SCQ-----VQ-LD--
gb|MDY6802921.1|      --------------------------------M----------KPNYRRCAICRKVAHKEAFWRVVRL-----S-S---------SHM-----VQ-LD--
ref|WP_280651647.1|   --------------------------------M--KP--------NYRRCISCRKVSLKEDFWRIVRV-----F-P---------SGK-----VQ-LD--
gb|MBD1868348.1|      --------------------------------M-L-P--------NYRRCVSCRRVALKSGFWRIVRV-----H-P---------SGT-----VE-LD--
tpg|HAG85663.1|       ---------------------------------------M---EPNYRRCLSCHKVAPKTAFWRIVRV-----H-P---------SRQ-----VQ-LD--
ref|WP_071593325.1|   --------------------------------M--KP--------NMRRCISCRKVADKEEFWRVVRV-----C-P---------SHQ-----LQ-LD--
gb|TVQ51200.1|        --M---------------------PP-------------------NHRRCLACRRIGTKSEFWRIVRL-----H-P---------TGT-----VQ-LD--
ref|WP_190487963.1|   --------------------------------M-Q-P--------NYRRCASCRKVAPKEEFLRIVRV-----Y-P---------NRE-----IR-LD--
gb|MEC4984856.1|      --------------------------------M----------KINYRRCLSCRRVAPKESFWRIVRV-----Y-P---------SRQ-----VQ-LD--
ref|WP_190426410.1|   --------------------------------M-KPNY---------RRCVSCRQVALKSDFWRVVRL-----Y-P---------SGK-----LQ-LD--
ref|WP_220611023.1|   --------------------------------M--KP--------NYRRCISCRRVRPKEEFWRIVRI-----F-P---------SGK-----VQ-LD--
ref|WP_190391042.1|   --------------------------------M-K-P--------NYRRCISCRQVNLKQEFWRIVRV-----F-P---------SGK-----VQ-LD--
ref|WP_212664564.1|   --------------------------------M----------QPNHRRCISCRHIAPKSEFWRIVRV-----Y-P---------DHQ-----IQ-LD--
gb|NJR53419.1|        -----------------MQP-------------------------NYRRCISCRQVAPKSEFWRIVRVYP---------------DHR-----VQ-LD--
ref|WP_318728151.1|   ---------ME-------------------------PN--------QRRCVSCRKVDRKEAFWRIVRV-----Y-P---------DS---K--IQ-LDRG
gb|TAE00837.1|        --------------------------------M----------KPNYRRCCASCRKIATKEAFWRVVRV-----Y-P---------SHQ-----VQ-LD--
```

```
gb|NJR38792.1|         -----------------------------------------MLPNHRCCISCRKVSPKEVFWRIVRV-----F-P---------SQA-----VQ-LD--
dbj|BAY85099.1|        --------MK-PN--------------------------------WRRCISCRKVSLKHEFWRIVRV-----F-P---------EG---K--VQ-LNQG
gb|NJK66758.1|         -------------------------------M----------KPNYRCCASCRKIAPKEAFWRVVRL-----Y-P---------SHQ-----VQ-LD--
ref|WP_169616303.1|    ----------------MKP-------------------------NYRRCVSCRRVAPKADFWRVVRVY------P---------GGE-----VR-LG--
dbj|BAZ53121.1|        ----------------------MLTWGHDRPM--KP--------NYRRCISCRKVSLKEDFWRIVRV-----F-P---------SGN-----VQ-LD--
ref|WP_144864508.1|    -----------------------------------------MKKNYRRCVSCRHVASKEYFWRIVRT-----Y-P---------DRK-----IQ-LD--
gb|MBW4669271.1|       -------------------------------M--KT--------NYRRCISCRKIGLKQEFWRIVRV-----F-P---------SGQ-----VQ-LN--
ref|WP_193933574.1|    -------------------------------M-K-P--------NYRRCIGCRRVALKHEFWRIVRV-----Y-P---------SGE-----LQ-LD--
gb|MBD2363150.1|       ----------------------M-----------KP--------NYRRCVSCRKVGLKEEFWRIVRV-----F-P---------SGK-----VQ-LN--
ref|WP_271941877.1|    --------ME-------------------------PN--------QRRCVSCRKVDRKEAFWRIVRV-----Y-P---------DS---K--IQ-LDRG
ref|WP_190409163.1|    ----------------------M-----------QP--------NYRRCISCRKVGLKPDFWRIVRV-----F-P---------SGK-----VQ-LN--
ref|WP_198126109.1|    -------------------------------M--KP--------NYRRCISCRKVSLKEDFWRIVRV-----F-P---------SGN-----VQ-LD--
ref|WP_137906635.1|    -------------------------------M-K-P--------NYRRCISCRQVNLKQEFWRIVRV-----F-P---------SGK-----VQ-LD--
ref|WP_275521808.1|    ----------------------GVTNLYSP-----QKQM---KPNYRRCISCRKLGPKEAFWRLVRL-----Y-P---------CHQ-----VQ-LD--
gb|MBW4594787.1|       -------------------------------M--EP--------NYRRCISCRKTRLKQEFWRIVRV-----F-P---------SGQ-----VQ-LN--
gb|MDJ0678327.1|       ----------------------------M-----KP--------NYRRCISCRQILPKEKLWRIVKV-----H-P---------DQK-----IV-LD--
gb|QSJ17240.1|         -------------------------------M--KP--------NYRRCISCRRVSLKQDFWRIVRV-----F-P---------SGN-----VQ-LD--
gb|NEO84548.1|         -------------------------------M--LK--------NYRRCVACRKVALKQEFWRIVRL-----H-P---------SRT-----VQ-LD--
gb|OCQ94358.1|         -----------------------------------------MKPNHRRCVSCRKVGPKEAFWRLVRL-----Y-P---------SHQ-----VQ-LD--
ref|WP_012168150.1|    -------------------------------M--QP--------NYRLCISCRQVAPKSELWRIVRV-----Y-P---------DHQ-----IQ-LD--
gb|PSO47335.1|         -------------------------------M--KKN--------NRRCISCRRIAPKEKLWRVVRV-----H-S---------SQT-----VQ-LD--
ref|WP_193847003.1|    -------------------------------M-K-P--------NYRRCISCRQVNLKQEFWRIVRV-----F-P---------SGK-----VQ-LD--
dbj|BAZ90194.1|        -------------------------------M--KP--------NYRRCISCRRVGTKTEFWRIVRL-----F-P---------SGK-----VE-LD--
tpg|HAT12760.1|        -----------------------------------------MKPNYRCCASCRKIALKEAFWRVVRV-----Y-P---------SHQ-----VQ-LD--
gb|MCY7384728.1|       -----------------------------------------MKPNYRCCASCRKIALKEAFWRVVRV-----Y-P---------SHQ-----VQ-LD--
ref|WP_015226430.1|    ----------------------------M-----KT--------NYRRCLSCRRIAPKSEFLRIIRV-----H-P---------SKT-----IQ-LD--
ref|WP_236507116.1|    -----------------------------------------MKPNYRCCASCRKIAPKEAFWRVVRL-----Y-P---------SHQ-----VQ-LD--
ref|WP_012164444.1|    -------------------------------M--QPN--------HRRCISCRHVAPKSEFWRIVRV-----Y-P---------DHQ-----IQ-LD--
gb|TAF06084.1|         -------------------------------M--KP--------NYRRCISCRRGGQKEEFWRIVRV-----F-P---------SGK-----VQ-LD--
gb|NJL79274.1|         -------------------------------M-K-P--------NWRRCISCRKVSLKQEFWRIVRV-----F-P---------SGK-----VQ-LD--
ref|WP_315863139.1|    --------------------------------------M---AVRQRRCVACGRVADRSQFWRIVRC-----W-P---------DQT-----VQ-LD--
gb|MBD2100510.1|       ------------------------------------MRS---KPNYRRCLSCRRIDHRSSFWRIVRS-----Y-P---------SRK-----VQ-LE--
ref|WP_028082426.1|    -------------------------------M-K-P--------NYRRCISCRQVNLKQEFWRIVRV-----F-P---------SGK-----VQ-LD--
gb|MCL6433641.1|       -------------------------------M----------EPNYRRCLSCRRIAHKSELWRIVRV-----Y-P---------TGA-----IA-LD--
ref|WP_193925389.1|    -------------------------------M-K-P--------NWRRCISCRKVSLKQEFWRIVRV-----F-P---------SGK-----VQ-LD--
gb|PSB25882.1|         -----------------------------------------MKPNYRCCASCRKIALKEAFWRVVRV-----Y-P---------SHQ-----VQ-LD--
ref|WP_194064403.1|    -----------------------------------------MKPNYRCCASCRKIAPKEAFWRVVRV-----Y-P---------SHQ-----VQ-LD--
ref|WP_170189095.1|    --M--------------------KP-------------------NIRRCISCGKVAHKAAFWRIVRV-----H-P---------SHQ-----LQ-LD--
gb|MBW4686284.1|       -------------------------------M-K-P--------NYRRCTSCRKVGLKEEFWRIVRV-----F-G---------SGK-----VQ-LD--
ref|WP_271796938.1|    -------------------------------M-K-P--------NYRRCISCRQVNLKQEFWRIVRV-----F-P---------SGK-----VQ-LD--
ref|WP_071515609.1|    -------------------------------M--K--------SNYRRCISCRRVGPKESFWRVVRV-----Y-P---------SHQ-----VQ-LD--
gb|TVP65994.1|         ----------------MKP-------------------------NYRRCVSCRRVAPKADFWRVVRVY------P---------GGE-----VR-LG--
gb|MBI1240017.1|       -------------------------------M--KP--------NYRRCISCRKVSLKEDFWRIVRV-----F-P---------SGK-----VQ-LD--
ref|WP_252659375.1|    --M--------------------KP-------------------NLRRCISCGKVAPKAAFWRIVRV-----H-P---------SHQ-----LQ-LD--
gb|EKQ70938.1|         ----------------------------M-----EP--------NIRRCVACRKAAPKSEFWRIVRL-----H-P---------SHA-----IV-LD--
gb|NJR17659.1|         -------------------------------M-K-P--------NYRRCVSCRKVGLKQEFWRIVRL-----F-P---------SGQ-----VQ-LD--
ref|WP_168492396.1|    -------------------------------M-K-P--------NYRRCISCRQVNLKQEFWRIVRV-----F-P---------SGK-----VQ-LD--
ref|WP_010471496.1|    -------------------------------M----------QPNHRRCISCRHIAPKSEFWRIVRV-----Y-P---------DHQ-----IQ-LD--
ref|WP_313949248.1|    ---------------------------------MAPLRS---KPNYRRCLSCRRIDHRSSFWRIVRS-----Y-P---------SRK-----VQ-LE--
ref|WP_138499475.1|    -------------------------------M--KP--------NYRRCISCRKLGLKDEFWRIVRV-----F-P---------SGK-----VQ-LD--
tpg|HLO85610.1|        -------------------------------M-K-P--------NYRRCISCRLLRPKEEFWRIVRV-----F-P---------SGK-----VQ-LD--
ref|WP_190625876.1|    -------------------------------M--QP--------NYRRCVSCRRAAPKAEFWRVVRV-----Y-P---------NRL-----VH-LG--
ref|WP_250121821.1|    --------MK-PN--------------------------------YRRCISCRRVALRQEFWRIVRV-----H-P---------FG---K--LQ-LDRG
```

```
ref|WP_044105921.1|  --------------------------------M----------QPNYRRCVSCRKIAPKQTFWRIVRV-----H-S---------SRQ-----VQ-LD--
gb|MBW4519521.1|     --------------------------------M----------QPNYRRCLSCRKVAPKEAFLRLVRI-----H-P---------SHT-----VV-LD--
dbj|BAZ29108.1|      --------------------------------M-K-P--------NYRRCISCRKASLKEEFWRIVRV-----F-P---------SGK-----VQ-LN--
ref|WP_071992767.1|  --------------------------------M--KP--------NYRRCISCRQVNLKQEFWRIVRV-----F-P---------SGK-----VQ-LD--
ref|WP_013190105.1|  --------------------------------M---K-------PNYRRCISCRRLGLREEFWRIVRL-----F-P---------SGK-----VQ-LD--
ref|WP_015214913.1|  --------------------------------M-K-P--------NYRRCISCRRIELKEEFWRIVRV-----F-P---------SGK-----VQ-LD--
gb|MDJ0796006.1|     --------------------------------M--KA--------NYRRCISCRRIGIKQEFWRIVRV-----F-P---------SGQ-----VQ-LD--
gb|MDY6901569.1|     --------------------------------M-K-P--------NWRRCISCRKVSLKHEFWRIVRV-----F-P---------EGK-----VQ-LN--
gb|MBW4650502.1|     ------------------------------------------MEPNNRRCVSCRQVAPKAAFWRIVRV-----Y-P---------SGL-----VQ-LD--
ref|WP_214431970.1|  --------------------------------M--KP--------NYRRCISCRKVSPKEDFWRIVRV-----F-P---------EGK-----VQ-LD--
ref|WP_007355211.1|  ---------------------------------------M---KPNYRRCISCRKLGPKEAFWRLVRL-----Y-P---------CHQ-----VQ-LD--
ref|WP_163668145.1|  ---------MR-PN--------------------------------YRRCVSCRKAAHRDQFLRVVRT-----Y-P---------TG---T--IQ-LNYG
gb|MBW4642782.1|     --------------------------------M--EP--------NYRRCISCRQVRLKEEFWRIVRV-----F-P---------SGK-----VQ-LD--
gb|MCM0590902.1|     --------------------------------M--KP--------NYRRCISCRQVGLKEEFWRIVRV-----F-G---------SGK-----VQ-LD--
gb|RMF69880.1|       -----------------------------ML-----P--------NYRRCLSCRKTAPKAEFWRVVRV-----Y-P---------SQA-----IQ-LD--
ref|WP_199250663.1|  ---------------------------------------M---KPNYRRCVSCRQVAPKEAFWRVVRI-----H-P---------GHQ-----VQ-LD--
tpg|HIK55694.1|      -----------------MP-----------------P--------NYRRCVVCRKIDHKSAFWRIVRL-----F-P---------SRK-----IQ-LD--
gb|MBD0267699.1|     --------------------------------M-Q-P--------NYRRCASCRKVAPKEEFLRIVRV-----Y-P---------NRE-----IR-LD--
ref|WP_009344612.1|  --------------------------------M--KP--------NYRRCISCRRVGRKTEFWRIIRL-----F-P---------SGK-----VE-LD--
ref|WP_190700710.1|  --------------------------------M--PP--------NYRRCISCRRVAPKADFWRVVRV-----Y-P---------DRS-----VH-LG--
ref|WP_168646496.1|  --------------------------------M--KP--------NYRRCISCRQVNLKQEFWRIVRV-----F-P---------SGK-----VQ-LD--
gb|MEB3164071.1|     --------------------------------M----------PPHYRRCISCRKVAPKAEFWRVVRV-----A-G---------DRV-----VQ-LD--
gb|NJL35745.1|       --------------------------------M--LP--------NYRRCVSCRKVAPKSDFWRVVRV-----H-P---------SGT-----VE-LD--
gb|KPQ38479.1|       --M---------------------KP-------------------NIRRCISCGKVAPKAAFWRIVRV-----H-P---------SHQ-----LQ-LD--
ref|WP_190679736.1|  --------------------------------M-L-P--------NYRRCVSCRRVALKSGFWRIVRV-----H-P---------SGT-----VE-LD--
ref|WP_159789286.1|  --M---------------------KP-------------------NIRRCISCGKVAHKAAFWRIVRV-----H-P---------SHQ-----LQ-LD--
ref|WP_190641945.1|  --------------------------------M-L-P--------NYRRCVSCRRVALKSGFWRIVRV-----H-P---------SGT-----VE-LD--
ref|WP_015083448.1|  --------------------------------M--KP--------NYRRCISCRQVNLKQEFWRIVRV-----F-P---------SGK-----VQ-LD--
ref|WP_281153046.1|  --------------------------------M----------QPNYRRCISCRKIAPKQTFWRIVRV-----H-S---------SRQ-----VQ-LD--
ref|WP_198807873.1|  --------------------------------M--APN--------DRRCVSCRRVAPKAEFWRVVRL-----A-H---------STT-----VC-LD--
ref|WP_072033681.1|  --------------------------------M--KP--------NYRRCISCRQVNLKQEFWRIVRV-----F-P---------SGK-----VQ-LD--
tpg|HIK31517.1|      --------------------------------M--KP--------NYRRCVSCRQVAPKETFWRVVRL-----H-P---------SHQ-----LQ-LD--
ref|WP_293246005.1|  ------------------------------------------MKPNYRCCASCRKVAPKEAFWRVVRL-----Y-P---------SHQ-----VQ-LD--
gb|MBD1833872.1|     --------------------------------M-KPNY---------RRCVSCRKVALKSDFWRVVRL-----Y-P---------LGK-----LQ-LD--
ref|WP_277865451.1|  --------------------------------M--PP--------HTRRCLACRKLGPKSEFWRVVRI-----F-P---------SGE-----VV-LD--
ref|WP_168505662.1|  --------------------------------M-K-P--------NYRRCISCRQVNLKQEFWRIVRV-----F-P---------SGK-----VQ-LD--
gb|NEQ95601.1|       --M---------------------EQ-------------------NHRRCISCRKMAHKREFWRIVRQ-----Y-P---------SHN-----IQ-LD--
gb|MBW4459453.1|     --------------------------------M--QP--------NYRRCVSCRRVAPKADFWRVVRV-----Y-P---------DRV-----VR-LG--
gb|MDJ0617177.1|     --------------------------------M--KA--------NYRRCISCRRIGIKQEFWRIVRV-----F-P---------SGQ-----VQ-LD--
gb|MCL1475321.1|     --------------------------------M--EP--------NYRRCISCRQVAPKQALWRVVRV-----Y-P---------SRQ-----VQ-LD--
ref|WP_272035636.1|  ---SKGLGEEGKKRKSLSLFPFFPLPSSLAPSPFFTQKQM---KPNYRRCISCRKLGPKEAFWRLVRL-----Y-P---------CHQ-----VQ-LD--
ref|WP_106367757.1|  -----------------------------M-----KP--------NYRRCISCRRAALKQEFWRIIRV-----N-P---------SGQ-----LQ-LD--
ref|WP_096670737.1|  --------------------------------M--KP--------NYRRCISCRQVNLKQEFWRIVRV-----F-P---------SGK-----VQ-LD--
ref|WP_099701299.1|  --------------------------------M-K-P--------NYRRCIGCRRAALKHEFWRIVRV-----Y-P---------SGE-----LQ-LD--
ref|WP_224085747.1|  -----------------------M-----------QP--------NYRRCISCRKVGLKQDFWRIVRV-----F-P---------SGN-----VQ-LN--
gb|TAE05943.1|       ------------------------------------------MKPNYRCCTSCRKIAPKEAFWRVVRL-----Y-P---------SHQ-----VQ-LD--
ref|WP_067774615.1|  -----------------------M-----------QP--------NYRRCISCRKVRLKQDFWRIVRV-----F-P---------SGN-----VQ-LN--
ref|WP_015178990.1|  ------------------------------------------MKPNYRCCASCRKIAPKEAFWRVVRL-----Y-P---------SHQ-----VQ-LD--
gb|MCC5897105.1|     --M---------------------KP-------------------NIRRCISCGKVAHKAAFWRIVRV-----H-P---------SHQ-----LQ-LD--
gb|NER36886.1|       ------------------------------------------MKPNYRRCISCRKVAPKEAFWRAVRV-----Y-P---------SHQ-----VQ-LD--
gb|MEB3252158.1|     --------------------------------M----------PPNYRRCISCRQVAPRSCFWRVVRR-----H-P---------DGT-----VH-LD--
ref|WP_230402764.1|  --------MTP-PR----QRTTQKHKMP--------PG--------TRRCLSCRTLAPREQFWRVVRL-----F-P---------SH---Q--VV-LDQG
ref|WP_099532894.1|  ---------MP-PN--------------------------------YRRCLICRRVAHRSEFWRIVRV-----A-H---------SG---S--VQ-LDQG
```

```
ref|WP_190534658.1|   ---------MP-PN--------------------------------YRRCLICRRVAHRSEFWRIVRV-----A-H---------SG---S--VQ-LDQG
gb|NJN88730.1|        --------------------------------M--QP--------NQRRCTVCRSIAPKSAFWRIVRV-----Y-P---------SHT-----VQ-LD--
ref|WP_193975681.1|   ------------------------------------------MKPNYRCCASCRKIAPKEAFWRVVRL-----Y-P---------SHQ-----VE-LD--
ref|WP_053538412.1|   --------------------------------M--KP--------NYRRCISCRQVNLKQEFWRIVRV-----F-P---------SGK-----VQ-LD--
ref|WP_072620812.1|   -----------------------MHQ-------------------NYRRCLACRTCGPKVIFWRIVRL-----H-P---------TRT-----VQ-LD--
gb|MDD1421071.1|      --------------------------------M--KP--------NYRRCISCRQVNLKQEFWRIVRV-----F-P---------SGK-----VQ-LD--
ref|WP_271948199.1|   --MNPFI--------------------------------------NYRRCISCRLFAPKESFWRIVRL-----H-P---------SRQ-----IQ-LD--
gb|NES70347.1|        ---------------------------------------M---EPNYRCCVSCRKVALKSEFWRVVRL-----Y-S---------SKE-----VQ-LD--
gb|NBO30352.1|        --------------------------------M----G------IHYRRCISCRKVAPKDFFWRVVRL-----Y-P---------LNR-----V------
gb|MBD1828535.1|      ------------------------------------------MKPNYRCCASCRKIAPKEAFWRVVRL-----Y-P---------SHQ-----VQ-LD--
gb|PZV12745.1|        --------------------------------M--QT--------NHRRCVSCRRVAPKAEFWRVVRV-----Y-P---------DRQ-----VC-LD--
gb|PZO41402.1|        --------------------------------M--QT--------NHRRCVSCRRVAPKAEFWRVVRV-----Y-P---------DRQ-----VC-LD--
ref|WP_015172566.1|   ---------------------------------------M---QPNCRRCISCRTAGPKVAFWRVVRV-----Y-P---------SQA-----IQ-LD--
ref|WP_190754133.1|   --------------------------------M--QP--------NYRRCVSCRRIAPKADFWRVVRV-----Y-P---------DRV-----VR-LG--
gb|UNU24151.1|        ------------------------------------------MKPNYRCCASCRKIAPKEAFWRVVRL-----Y-P---------SHQ-----VQ-LD--
gb|MBS3030443.1|      --------------------------------M--KP--------NYRRCISCRQVNLKQEFWRIVRV-----F-P---------SGK-----VQ-LD--
gb|MBI4782862.1|      --------------------------------M-R-P--------NQRRCVSCRKIAPKADFWRIVRV-----F-P---------SRT-----VQ-LE--
gb|MDJ0724125.1|      --------------------------------M-Q-P--------NYRRCLGCGKIATKASFWRIVRV-----Y-P---------SSK-----VQ-LD--
gb|MBD1815077.1|      ------------------------------------------MKPNYRCCASCRKIAPKEAFWRVVRL-----Y-P---------SHQ-----VQ-LD--
gb|EGK88241.1|        ------------------------------------------MKPNYRCCASCRKIAPKEAFWRVVRL-----Y-P---------SHQ-----VQ-LD--
ref|WP_072207768.1|   -----------------------MKP-------------------NYRRCISCRKVGLKEEFWRIVRV-----F-S---------SGQ-----VQ-LD--
gb|MCG9884819.1|      -----------------------MAPP------------------NHRRCISCRRLGHRTEFWRIIRL-----H-P---------SHQ-----VQ-LD--
ref|WP_190528272.1|   ---------MP-PN--------------------------------YRRCLICRRVAHRSEFWRIVRV-----A-H---------SG---S--VQ-LDQG
ref|WP_011057771.1|   --------------------------------------MM---SVPQRRCVSCGRVADRSEFWRIVRC-----W-P---------DQK-----VQ-LD--
ref|WP_264321713.1|   --------------------------------M--KP--------NYRRCVSCRKVDDRDSFWRVVRL-----Y-P---------SHQ-----LQ-LD--
ref|WP_029634746.1|   -----------------------------MK-K--N----------YRRCISCRQVKLKEEFWRIVRV-----F-P---------SRQ-----VQ-LN--
ref|WP_026099823.1|   ---------------------------------------M---VPNTRRCLSCRKVAPKAEFWRVVRR-----A-G---------DRQ-----VQ-LD--
gb|MBW4568874.1|      -----------------------------MK-K--N----------YRRCISCRQVKLKEEFWRIVRV-----F-P---------SRQ-----VQ-LN--
gb|MDY7022637.1|      ------------------------------------------MLPNYRRCVACRQVAPKAAFWRVVRL-----Y-S---------SHQ-----VQ-LD--
gb|MBD1213631.1|      --------------------------------M-K-P--------NYRRCISCRQVNLKQEFWRIVRV-----F-P---------SGK-----VQ-LD--
ref|WP_071527241.1|   -----------------MQP-------------------------NYRRCVSCRRVGPKAEFWRLVRVYP---------------DRT-----VH-LG--
gb|PZU98241.1|        --------------------------------M--QPN--------YRRCVSCRQVGPKAEFWRVVRA-----Y-P---------DGT-----VR-VG--
tpg|HIK45324.1|       --------------------------------M--KP--------NYRRCVSCRRVGPKQEFWRLVRL-----A-G---------GTT-----VC-LD--
gb|OBQ35981.1|        --------------------------------M-K-P--------NYRRCISCRQVNLKQEFWRIVRV-----F-P---------SGK-----VQ-LD--
gb|NCR11955.1|        --------------------------------M--NTF------VNYRRCISCRLFAPKESFWRIVRL-----H-P---------SGQ-----IQ-LD--
ref|WP_228383259.1|   -------------------------------MS-RLP-------QGYRRCIACRRLAHRQVFWRVIRL-----S-S---------TGV-----IA-LD--
gb|MBW4611665.1|      -----------------------------MK-K--N----------YRRCISCRQVKLKEEFWRIVRV-----F-P---------SRQ-----VQ-LN--
ref|WP_028091624.1|   --------------------------------M-K-P--------NYRRCISCRQVNLKQEFWRIVRV-----F-P---------SGK-----VQ-LD--
gb|MEB3290003.1|      -----------------------------MA-----P--------NDRRCVSCRRVAPKSEFWRVVRL-----A-H---------STT-----IG-ID--
gb|MBD0262109.1|      -----------------------------MK-K--N----------YRRCISCRQVKLKEEFWRIVRV-----F-P---------SRQ-----VQ-LN--
ref|WP_035737147.1|   -----------------MVD-------------------------HLRRCVTCRRLAAKQEFWRLVRVYP---------------SHQ-----VE-LD--
ref|WP_024124972.1|   -------------------------------MM----A------VPQRRCVACGRVADRSQFWRIVRC-----W-P---------DQK-----V------
gb|OKH17997.1|        -------------------------------MT--QT--------NYRRCVACRKLAPKQSFWRVVRV-----Y-P---------SQT-----VQ-LD--
tpg|HBK98803.1|       --------------------------------M----------KPNYRCCACCRKIALKEAFWRVVRL-----Y-P---------SHQ-----VQ-LD--
ref|WP_006512183.1|   -----------------------------M-----KI--------NYRRCISCKKTASKEEFWRIVRL-----Q-G---------SCE-----II-LD--
ref|WP_072044931.1|   --------------------------------M----------KPNYRRCISCRRLALKQEFWQIVRL-----Y-P---------SGK-----LQ-LD--
ref|WP_023068840.1|   ------------------------------------------MQPNYRRCIACRQVARKETFWRVVRL-----Y-S---------SHQ-----VQ-LD--
gb|MBF2027289.1|      -----------------------------M-----EP--------NVRRCVACRKTAPKAEFWRIVRI-----H-G---------FHA-----IV-LD--
ref|WP_272116663.1|   --MVEFRVEEE-----NV-----LHQ-------------------NYRRCLACRTCGPKVIFWRIVRL-----H-P---------TRT-----VQ-LD--
ref|WP_193960470.1|   --MNPFV--------------------------------------NYRRCISCRLFAPKESFWRIVRL-----H-P---------SGQ-----IQ-LD--
gb|RFP56458.1|        ---------MP-PN--------------------------------YRRCLICRRVAHRSEFWRIVRV-----A-H---------SG---S--VQ-LDQG
ref|WP_125732142.1|   --MNTFV--------------------------------------NYRRCISCRLFAPKESFWRIVRL-----H-P---------SGQ-----IQ-LD--
ref|WP_006621525.1|   -----------------MVD-------------------------HLRRCVTCRRLAAKQEFWRLVRVYP---------------SHQ-----VE-LD--
```

```
ref|WP_190386203.1|   --------------------------------M--KP--------NYRRCISCRQVNLKQEFWRIVRV-----F-P---------SGK-----VQ-LD--
ref|WP_163518983.1|   -----------------------------MK-K--N----------YRRCISCRQVKLKEEFWRIVRV-----F-P---------SRQ-----VQ-LN--
dbj|BAI90404.1|       -----------MGGYKIMVD-------------------------HLRRCVTCRRLAAKQEFWRLVRVYP---------------SHQ-----VE-LD--
ref|WP_190574353.1|   --------------------------------M--EP--------NYRRCVSCRKIAPKTDFWRVVRT-----F-P---------DQA-----VQ-VD--
gb|MDJ0508716.1|      --M---------------------KP-------------------NIRRCISCRCVAPKEAFWRIVRV-----S-S---------SRK-----VQ-LD--
gb|TVR11480.1|        --M---------------------KP-------------------NIRRCISCGKVAHKAAFWRIVRV-----H-S---------SHQ-----LQ-LD--
ref|WP_310454240.1|   --------------------------------------MM---AVPQRRCVSCGRLADRSEFWRIVRC-----W-P---------DQK-----VQ-LD--
gb|MBC6434993.1|      ------------------------------------------MLPNYRRCISCRKVSPKEGFWRIVRV-----F-P---------SQA-----VQ-LG--
ref|WP_002798549.1|   --MNTFV--------------------------------------NYRRCISCRLFAPKESFWRIVRL-----H-P---------SGQ-----IQ-LD--
dbj|BAZ16285.1|       --------------------------------M---K-------PNYRRCISCRKIGLKQEFWRIVRV-----G-P---------SGQ-----VQ-LD--
gb|MBW4493842.1|      --------------------------------M--KP--------NMRRCISCRKVAGKEEFWRVVRV-----C-P---------SHQ-----LQ-LD--
gb|EKV02414.1|        ---------MR-PN--------------------------------YRRCVSCRKAAHRDQFLRVVRT-----Y-P---------TG---T--IQ-LNYG
ref|WP_190665968.1|   --------------------------------M-E-T--------NYRRCISCRRVAHKNDLWRIVRV-----F-P---------SHQ-----LQ-LD--
ref|WP_089128587.1|   -----------------------------MK-K--N----------YRRCISCRQVKLKEEFWRIVRV-----F-P---------SRQ-----VQ-LN--
ref|WP_293126645.1|   ------------------------------------------MKPNYRCCASCRKIAPKEAFWRVVRV-----Y-P---------SHQ-----VQ-LD--
gb|MEB3224903.1|      --------------------------------M--EP--------NLRRCVACRKLAPKQSLWRVVRV-----Y-P---------SQQ-----VQ-LD--
gb|NET30621.1|        -----------------------------M-----KP--------NYRRCISCRVVAAKEVFWRIVRV-----H-P---------TRT-----VQ-LD--
gb|QSF50481.1|        ---------------------------------------M---AVPQRRCVSCGRLADRSEFWRIVRC-----W-P---------DQK-----VQ-LD--
ref|WP_149818242.1|   --------------------------------------MM---AVPQRRCVSCGRLADRSEFWRIVRC-----W-P---------DQK-----VQ-LD--
ref|WP_263749268.1|   -----------------------------MK-K--N----------YRRCISCRQVKLKEEFWRIVRV-----F-P---------CRQ-----VQ-LN--
ref|WP_073069683.1|   --------------------------------M--EP--------NYRRCISCRRTAPKQEFWRVVRS-----H-S---------SNT-----IA-LD--
ref|WP_271954606.1|   --MNPFV--------------------------------------NYRRCISCRLFAPKESFWRIVRL-----H-P---------SGQ-----IQ-LD--
tpg|HIK12112.1|       ---------------------------------------M---KPNYRRCVSCRQVAPKEALWRVVRI-----H-P---------GHQ-----VQ-LD--
gb|MBW4506515.1|      --------------------------------M--EP--------NRRRCISCRKVSPKQDFWRVVRV-----F-S---------TGQ-----VQ-LD--
ref|WP_287684335.1|   --MNTFV--------------------------------------NYRRCISCRLFAPKESFWRIVRL-----H-P---------SGQ-----IQ-LD--
ref|WP_225875273.1|   ------------------MVLPLGDRPPKKHMT--QT--------NYRRCVACRKLAPKQSFWRVVRV-----Y-P---------SQT-----VQ-LD--
gb|TRV14446.1|        --MNPFV--------------------------------------NYRRCISCRLFAPKESFWRIVRL-----H-P---------SGQ-----IQ-LD--
gb|MBW4477482.1|      -----------------------------MK-K--N----------YRRCISCRQVKLKEEFWRIVRV-----F-P---------CRQ-----VQ-LN--
gb|NJK40366.1|        --------------------------------M-D-L--------NYRRCLSCRKVAPKSEFWRIVRV-----H-P---------THI-----VQ-LD--
gb|TRU22031.1|        --MNPFV--------------------------------------NYRRCISCRLFAPKESFWRIVRL-----H-P---------SGQ-----IQ-LD--
ref|WP_002768359.1|   --MNPFV--------------------------------------NYRRCISCRLFAPKESFWRIVRL-----H-P---------SGQ-----IQ-LD--
gb|MDJ0546172.1|      --MNTFV--------------------------------------NYRRCISCRLFAPKESFWRIVRL-----H-P---------SGQ-----IQ-LD--
ref|WP_287078697.1|   --MNTFV--------------------------------------NYRRCISCRLFAPKESFWRIVRL-----H-P---------SGQ-----IQ-LD--
ref|WP_226583963.1|   --------------------------------M--KP--------NFRRCVSCRQVAPKEDFWRLVRA-----Y-P---------SGQ-----VL-LD--
gb|MBL1176422.1|      -----------------------------M-----KP--------NYRCCVGCRKIAHKSTLWRIVRL-----Y-P---------SHQ-----VQ-LD--
ref|WP_159293949.1|   --MNPFV--------------------------------------NYRRCISCRLFAPKESFWRIVRL-----H-P---------SGQ-----IQ-LD--
ref|WP_071776908.1|   --------MAG-KN--------------------------------HRLCVSCRQTAHRDQLWRVVRT-----Y-P---------HR---K--VQ-LDYG
ref|WP_190792225.1|   -----------------MP-----------------P--------NHRRCISCRRVAPREEFWRVVRS-----F-P---------SQI-----VM-LD--
ref|WP_271991901.1|   --MNPFL--------------------------------------NYRRCISCRLFAPKESFWRIVRL-----H-P---------SGQ-----IQ-LD--
ref|WP_110577867.1|   --MNPFV--------------------------------------NYRRCISCRLFAPKESFWRIVRL-----H-P---------SGQ-----IQ-LD--
ref|WP_286826366.1|   --MNTFV--------------------------------------NYRRCISCRLFAPKESFWRIVRL-----H-P---------SGQ-----IQ-LD--
gb|NCR41277.1|        --MNPFV--------------------------------------NYRRCISCRLFAPKESFWRIVRL-----H-P---------SGQ-----IQ-LD--
gb|NCR55568.1|        --MNPFV--------------------------------------NYRRCISCRLFAPKESFWRIVRL-----H-P---------SGQ-----IQ-LD--
ref|WP_190367604.1|   --------------------------------M--KP--------NYRRCVSCRQVKLKQDFWRVVRI-----F-P---------SGN-----VE-LD--
ref|WP_150976468.1|   --MNPFL--------------------------------------NYRRCISCRLFAPKESFWRIVRL-----H-P---------SGQ-----IQ-LD--
ref|WP_149986802.1|   --MNPFV--------------------------------------NYRRCISCRIFAPKESFWRIVRL-----H-P---------SGQ-----IQ-LD--
gb|MDX2096196.1|      -----------------MK-----------------P--------NYRRCVSCQKIAPRTEFWRIVRL-----H-P---------SHE-----LQ-LD--
ref|WP_002738851.1|   --MNPFV--------------------------------------NYRRCISCRLFAPKESFWRIVRL-----H-P---------SGQ-----IQ-LD--
ref|WP_194012660.1|   ------------------------------------------MKPNYRCCASCRKIAPKEAFWRVVRV-----Y-P---------SHQ-----VQ-LD--
ref|WP_072721595.1|   --M-----------------------K------------------NQRRCVACRQLAGKETFWRFVRL-----Y-P---------SHQ-----VQ-LD--
ref|WP_108935646.1|   --MNPFV--------------------------------------NYRRCISCRLFAPKESFWRIVRL-----H-P---------SGQ-----IQ-LD--
ref|WP_272080703.1|   --MNPFV--------------------------------------NYRRCISCRLFAPKESFWRIVRL-----H-P---------SGQ-----IQ-LD--
gb|OUC16623.1|        --------------------------------M--QP--------NYRRCVSCRKAAPKADFWRIVRT-----F-P---------DQA-----VQ-LD--
```

```
ref|WP_271926239.1|  --MNPFV--------------------------------------NYRRCISCRLFAPKESFWRIVRL-----H-P---------SGQ-----IQ-LD--
gb|MBW4467673.1|     --------------------------------M-P-Q--------NYRRCVSCRKLDHRNSFWRIVRV-----F-P---------SQQ-----LQ-LD--
gb|NCQ85689.1|       --MNPFV--------------------------------------NYRRCISCRLFAPKESFWRIVRL-----H-P---------SGQ-----IQ-LD--
ref|WP_286626612.1|  --------------------------------------MM---AVPQRRCVSCGRLADRSDFWRIVRC-----W-P---------DQK-----VQ-LD--
ref|WP_009782890.1|  ------------------------------------------MQPNYRRCIACRQVAPKETFWRVVRV-----Y-S---------SHQ-----VQ-LD--
ref|WP_181496891.1|  ---------------------------------------M---AVPQRRCVACGRLADRSEFWRIVRC-----W-P---------DQT-----VQ-LD--
gb|AFY59850.1|       ------------------------------------------MPPHTRRCLSCRCLGHKSEFWRVVRLA------Q---------SQT-----VV-LD--
gb|MBC7883588.1|     --MN----------------------------Q-------------ERRCVSCRRVGPKTSFWRIVRI-----H-P---------SGQ-----IQ-LD--
ref|WP_009631737.1|  -----------------------------M-----KP--------NYRRCISCRRVALKQEFWRIIRV-----N-P---------SGQ-----LQ-LD--
gb|MBW4539957.1|     --------------------------------M--L--------PNYRRCISCRKVAPKEAFLRVVRL-----Y-P---------SHE-----IA-LN--
ref|WP_287969625.1|  --MNTFV--------------------------------------NYRRCISCRLFAPKESFWRIVRL-----H-P---------SGQ-----IQ-LD--
ref|WP_024970683.1|  --MNPFV--------------------------------------NYRRCISCRLFAPKESFWRIVRL-----H-P---------SGQ-----IQ-LD--
gb|MBX2866220.1|     --M---------------------QP-------------------NYRRCVSCRKIAHRNQLLRVVRT-----Y-P---------DAA-----VK-VN--
gb|NJK61661.1|       ---------ML------------------------------------RRCVSCRTLAHRSQFWRIVRD-----A-E---------TQ---T--VQ-LDVG
ref|WP_310743590.1|  --------------------------------------MM---AVPQRRCVSCGRLADRSEFWRIVRC-----W-P---------DQK-----LQ-LD--
ref|WP_119259560.1|  --------------------------------M----------QPNYRRCISCRKIAPKQTFWRIVRV-----H-S---------SRQ-----VQ-LD--
ref|WP_172191971.1|  ------------------------------------------MKSNYRRCCASCRKIAPKEAFWRVVRL-----Y-P---------SHQ-----VQ-LD--
ref|WP_322876998.1|  --------------------------MP--------PH--------TRRCLSCRCLGHKSEFWRVVRL---------A--Q----SQ---T--VV-LDQ-
ref|WP_310744398.1|  ---------------------------------------M---VVSQRRCVACGRLADRSEFWRIVRC-----W-P---------DQK-----VQ-LD--
ref|WP_088891762.1|  -----------------MP-----------------P--------NHRRCISCRRVAPKQEFWRVVRS-----F-P---------AQI-----VI-LD--
ref|WP_151695992.1|  --MNPFV--------------------------------------NYRRCVSCRLFAPKESFWRIVRL-----H-P---------SGQ-----IQ-LD--
ref|WP_267384513.1|  --------------------------------M----------QPNYRRCISCRKIAPKQTFWRIVRV-----H-S---------SRQ-----VQ-LD--
gb|NJO80776.1|       --------------------------------M--LP--------NHRRCISCRKVDHKSAFWRVVRV-----F-S---------SHQ-----VQ-LD--
gb|NEO34688.1|       --M---------------------KK-------------------NYRRCIGCRRVAHRSEFWRIVRQ-----Y-P---------SHT-----VQ-ID--
ref|WP_190359885.1|  --MNPFV--------------------------------------NYRRCISCRLFAPKESFWRIVRL-----H-P---------SGQ-----IE-LD--
ref|WP_308256896.1|  --------MNN-PN----KH----------------KN--------YRRCISCHRLDERKKFWRIVRT-----Y-P---------EK---K--VI-LDQG
gb|MDS3859689.1|     MK---PQPICCSER----KLTNSRLTMP--------PH--------TRRCLSCRCLGHKSEFWRVVRL---------A--Q----SQ---T--VV-LDQ-
gb|MBR8831668.1|     --------------------------------M--ET--------NYRRCPICRRTAPKGEFWRIVRV-----Y-P---------SKQ-----VQ-LD--
ref|WP_317267730.1|  -----------------MVN-------------------------HLRRCVTCRRLAAKQEFWRLVRVYP---------------SHQ-----VE-LD--
gb|REJ59451.1|       --MNPFV--------------------------------------NYRRCISCRLFAPKESFWRIVRL-----H-P---------SGQ-----IE-LD--
gb|MBP0000506.1|     --------------------------------M--KPH--------LRRCISCGKVAPKDAFWRVVRV-----H-P---------SHQ-----LQ-LD--
ref|WP_297050161.1|  ---------------------------------------M---ALPQRRCVSCGRLADRSEFWRIVRC-----W-P---------DQK-----VQ-LD--
tpg|HAZ43175.1|      --------------------------------M--KP--------NFRRCVSCRNVAPKEDFWRLVRA-----Y-P---------SGQ-----VL-LD--
gb|MBW4619585.1|     ---------ME-------------------------KN--------YRRCASCRLVALKSAFWRIVKV-----H-P---------SQ---Q--VQ-LDGG
ref|WP_298613741.1|  ---------------------------------------M---ANPQRRCVACGRLADRSEFWRVVRC-----W-P---------DQK-----VQ-LD--
gb|MCL1465298.1|     --------------------------------M--EP--------NYRRCISCRQVAPKQALLRVVRV-----Y-P---------SRQ-----VQ-LD--
ref|WP_298975829.1|  ---------------------------------------M---ANPQRRCVACGRLADRSEFWRVVRC-----W-P---------DQK-----VQ-LD--
ref|WP_168571252.1|  --------------------------------M--KV--------NYRRCISCRRVAPKQEFCRVVRV-----H-P---------SHT-----VQ-LD--
ref|WP_287729185.1|  --MSPFV--------------------------------------NYRRCISCRLFAPKESFWRIVRL-----H-P---------SGQ-----IQ-LD--
ref|WP_190764830.1|  --------------------------------M--PI--------NDRRCVSCRKLASKSEFWRVVRE-----H-P---------SGT-----VR-LD--
gb|NJL20540.1|       --------------------------------M--PI--------NDRRCVSCRKLAPKSEFWRIVRE-----H-P---------SGT-----VR-LD--
gb|MDA0866786.1|     --------------------------------M--QP--------NYRRCVSCREIAHKQDLWRIVRV-----H-D---------TQT-----VQ-LD--
gb|MDX2230996.1|     -----------------------------M-----KP--------NLRRCVVCRKIAPKPEFWRIVRL-----H-P---------SHA-----IA-LD--
gb|MBE9043893.1|     --------------------------------M--K-Q------KNYRRCISCRRVAPKKTFWRVVRL-----A-S---------SHE-----IQ-LD--
tpg|HAO10320.1|      ------------------------------------------MK-NQRRCVACRQLGAKESFWRFVRL-----Y-P---------SHQ-----VQ-LD--
ref|WP_317034091.1|  -----------------------------MQ-----P--------NHRCCVSCRRVAPKAEFWRVVRV-----Y-P---------DRT-----VV-LG--
gb|MBW4678534.1|     --------------------------------M-E-T--------NYRRCISCRRVAHKNDLWRIVRV-----F-P---------SHQ-----LQ-LD--
gb|MDJ0570890.1|     --------------------------------M---NQ------INYRRCISCRRVASKASFWRVVRL-----A-P---------NHE-----I------
ref|WP_287659929.1|  --MNPFV--------------------------------------NYRRCISCRLFAPKESFWRIVRL-----H-P---------SGQ-----IQ-LD--
ref|WP_190774570.1|  --------------------------------M-ERDY---------RRCVGCRQVAHKRYLWRVVRV-----F-P---------SRS-----VQ-LG--
ref|WP_310800080.1|  ---------------------------------------M---VVSQRRCVACGRLADRSEFWRIVRC-----W-P---------DHR-----VQ-LD--
ref|WP_002801176.1|  --MNPFV--------------------------------------NYRRCISCRLFAPKESFWRIVRL-----H-P---------SGQ-----IQ-LD--
ref|WP_190450280.1|  --M-------E---------------P------------------NYRCCISCRKIATKQAFWRIVRV-----A-P---------SQADSPPYVQ-LD--
```

```
gb|MBR8827281.1|      -----------------M-----------------KQ--------NYRRCISCRQIAPKDSFWRLVKL-----H-P---------SGQ-----VE-LD--
gb|MDC0831746.1|      --------------------------------M--KPN--------VRRCISCGKVAPKDAFWRVVRV-----H-P---------SHQ-----LQ-LD--
ref|WP_007306607.1|   -----------------------MKT-------------------NYRPCLSCGCIAPKQALWRIVRV-----Y-P---------SRQ-----VQ-LD--
gb|AJD57472.1|        -----------------------------------MP-------QGYRRCIACRRLAHRQEFWRVIRL-----A-S---------TGA-----IA-LD--
ref|WP_079676917.1|   ------------------------------------------MK-NQRRCVACRQLGAKESFWRFVRL-----Y-P---------SHQ-----VQ-LD--
ref|WP_082901633.1|   --------------------------------M--KPN--------VRRCISCGKVAPKDAFWRVVRV-----H-P---------SHQ-----LQ-LD--
ref|WP_190341424.1|   -----------------------------ME-----P--------NYRRCLGCGKIAHKSEFWRVVRT-----H-P---------SHA-----IE-LD--
ref|WP_149974369.1|   --MNPFV--------------------------------------NYRRCISCRLFAPKESFWRIVRL-----H-P---------SGQ-----IQ-LD--
ref|WP_019500197.1|   --------MAP-KL--------------------------------YRRCICCQRLAPREEFWRVVRL-----H-GDN--N----AT---N--IQ-LDRG
gb|MBV5261761.1|      --------------------------------M--EP--------NLRRCVACRKLAPKQTLWRVVRV-----Y-P---------SQQ-----VQ-LD--
ref|WP_324706802.1|   -----------------MPS-------------------------NLRRCVTCRQLAPKETFWRLVRVHP---------------SHR-----VQ-LD--
ref|WP_293094764.1|   -----------------------MKQ-------------------NYRRCVSCKKLAPKEEFWRVVRI-----K-Q---------SDT-----IR-LD--
ref|WP_287453144.1|   --------------------------------M--LP--------NYRRCICCRKVAPKAEFLRIVRL-----H-P---------SHE-----IA-ID--
ref|WP_083622039.1|   --M-----------------------K------------------NQRRCVACRQLAGKETFWRFVRL-----Y-P---------SHQ-----VQ-LD--
gb|NJP09111.1|        -----------------------------M-----EP--------NCRRCIACRRVAHKKDFWRIVRV-----H-P---------SHS-----VI-LD--
gb|MBW4420472.1|      --------------------------------M--L--------PNYRRCISCRKVAPKEAFLRVVRL-----Y-P---------SHE-----IA-LN--
ref|WP_159249505.1|   --MNPFV--------------------------------------NYRRCISCRLFAPKESFWRIVRL-----H-P---------SGQ-----IQ-LD--
ref|WP_083622686.1|   ------------------------------------------MK-NQRRCVACRRLGAKESFWRFVRL-----Y-P---------SHQ-----VQ-LD--
gb|MDX2215915.1|      -----------------ME-----------------V--------NYRRCVSCRKVAPKASFWRIVKV-----H-P---------AQN-----IC-LD--
ref|WP_046662795.1|   --MNPFV--------------------------------------NYRRCISCRLFAPKESFWRIVRL-----H-P---------SGQ-----IQ-LD--
ref|WP_316429393.1|   ------------------------------------------MPPHYRRCISCRRIAHKSAFWRVVRV-----F-P---------SHQ-----VQ-LD--
ref|WP_040054730.1|   --------MK--------------------------AN--------YRRCVSCRCTNHKKTFWRIVKV-----Y-L---------SQ---K--VQ-LDKG
gb|MBF2048868.1|      ------------------------------------------MPPHYRRCISCRRIAHKSAFWRVVRV-----F-P---------SHQ-----VQ-LD--
ref|WP_190504524.1|   -----------------ME-----------------I--------NYRRCVSCRRVALKSEFWRVVKT-----Y-P---------SSS-----IC-LD--
gb|MBW4692942.1|      -----------------M-----------------KP--------NDRRCVSCRKVAPKAEFWRIVRVQSAQRH-S---------PPT-----VQ-LD--
gb|MDR9403844.1|      --------------------------------M--EK--------NYRRCLSCRRIAPKTEFIRIIRV-----H-P---------SKR-----IQ-LD--
ref|WP_008203458.1|   --MNPFV--------------------------------------NYRRCISCRLFAPKESFWRIVRL-----H-P---------SGQ-----IQ-LD--
ref|WP_190759246.1|   --------------------------------M-E-T--------NYRRCISCRRVAHKNDLWRIVRV-----F-P---------SHQ-----LQ-LD--
tpg|HAX74477.1|       --------------------------------M--KP--------NFRRCVSCRKVAPKEDFWRIVRA-----D-P---------SGQ-----VV-LD--
gb|MCD8489703.1|      --------------------------------M--VV-------KNYRRCISCRKLAPKDAFWRVVRV-----H-P---------EGQ-----IQ-FD--
ref|WP_069966942.1|   ------------------------------------M-------KNYRRCISCRKLAPKDAFWRVVRV-----H-P---------EGQ-----IQ-FD--
ref|WP_204141672.1|   -----------------------------M-----KP--------NHRRCVSCRKIAPKSALWRVVRQ-----Y-G---------AGT-----VQ-VN--
ref|WP_141724342.1|   --------------------------------M--VV-------KNYRRCISCRKLAPKDAFWRVVRV-----H-P---------EGQ-----IQ-FD--
ref|WP_011244384.1|   -------------------------------MS-RLP-------QGYRRCIACRRLAHRQEFWRVIRL-----A-S---------TGA-----IA-LD--
gb|NEQ33015.1|        --------------------------------M--EP--------NYRRCISCRQVAHRDNLWRIVRV-----H-P---------THQ-----VQ-LD--
gb|MBW4699545.1|      -----------------M-----SNPAAVNPKQ------------NQRRCVSCRRTGDRESFWRIVRV-----H-P---------GWQ-----IQ-LN--
ref|WP_168535677.1|   --------------------------------M-K-P--------NYRRCISCRQVNLKQEFWRIVRV-----F-P---------SGK-----VQ-LD--
ref|WP_200669078.1|   --------------------------------M--KV--------NYRRCISCRRVAPKQEFCRVVRV-----H-P---------SHT-----VQ-LD--
ref|WP_250683588.1|   --------------------------------------MM---AVLQRRCVACGRLADRSEFWRIVRC-----W-P---------DQK-----VQ-LD--
ref|WP_024544928.1|   --------------------------------M--EP--------NLRRCVACRKLAPKESLWRVIRV-----Y-P---------SQQ-----VQ-LD--
gb|MBE9010997.1|      -----------------------M-----------PS--------NIRRCLSCRKVASKEEFLRVVRL-----Y-P---------SHE-----IS-ID--
ref|WP_193855116.1|   --------MAE-KN--------------------------------HRLCVSCRQTAHRNHLWRVVRT-----Y-P---------HR---K--IQ-LDYG
gb|NJN19948.1|        --------------------------------M--QP--------NYRRCVSCREIAHKRDLWRIVRV-----H-D---------TQA-----IQ-LD--
gb|MDJ0515540.1|      -----------------------MEQ-------------------NYRRCVSCKKLAPKEEFWRVVRI-----K-Q---------SDT-----IR-LD--
gb|MDY7006698.1|      -----------------------MKP-------------------NYRRCVSCKKLAPKEEFWRVIRL-----K-K---------SST-----IR-LD--
ref|WP_190649792.1|   --------------------------------M--LP--------NYRRCICCRKVAPKAEFLRIVRL-----H-P---------SHE-----IA-IN--
ref|WP_036044569.1|   --------------------------------M--LP--------NYRRCICCRKVAPKAEFLRIVRL-----H-P---------SHE-----IA-IN--
ref|WP_058999772.1|   --------------------------------M--PS--------NLRRCLTCRKIAPKTEFLRVVRL-----Y-P---------SHE-----IS-ID--
gb|MDJ0898873.1|      --M---------------------KI-------------------NYRRCISCKKTASKEEFWRVVRI-----P-N---------SHE-----IT-LA--
gb|MBW4665085.1|      -----------------------------M-----KP--------NHRRCISCRRVALKQEFWRIIRV-----N-T---------SGQ-----LQ-LD--
gb|MDJ0556426.1|      ---------MK-PN--------------------------------YRRCVSCKKLASKEDFWRVVRL-----S-K---------SS---T--VK-LDEG
gb|MBC8121277.1|      --MS----------------------------K------------NERRCVSCLRVGPKDSFWRMVRL-----H-P---------SWQ-----IQ-LN--
ref|WP_017287872.1|   ------------------MRSLKLRSHPVLFFM--LP--------NYRRCICCRKVAPKAEFLRIVRL-----H-P---------SHE-----IA-IN--
```

```
ref|WP_190376153.1|   -----------------------M-----------PP--------NYRRCLSCRKVALKEEFLRVVRL-----Y-P---------SHE-----IA-IN--
gb|MBC7972109.1|      -----------------M-----------------QP--------NYRRCVSCRKVALKQEFWRVVRVHSDVGQ-S---------TPV-----VQ-LD--
gb|MDY6937449.1|      --------------------------------M--KP--------KYRRCVSCRQIADRETLWRVVRL-----Y-G---------SHQ-----LQ-LD--
ref|WP_106237396.1|   --------------------------------M---NQ------INYRRCISCRKKIAPKQSFWRVVRL-----A-S---------SCQ-----I------
gb|MBD1842922.1|      -----------------------M-----------PP--------NHRRCLSCRKVAPKEEFLRVVRL-----Y-P---------SHE-----IS-ID--
ref|WP_323217923.1|   ------------------------------------------MSPNCRRCVACRQAAPKETFWRVVRV-----Y-S---------SHQ-----VQ-LD--
gb|NEQ45539.1|        --------------------------------M----------KPNHRRCVSCRAIAPKQGLWRAVRQ-----H-D---------TGI-----VQ-LD--
gb|MCY6491807.1|      --------------------------------M--LP--------NYRRCICCRKVAPKAEFLRIVRL-----H-P---------SHE-----IA-IN--
ref|WP_012954106.1|   --------MK--------------------------VN--------YRRCISCRCINHKDKFWRIVKV-----Y-L---------SQ---T--VQ-LDKG
ref|WP_065713394.1|   --------------------------------M--EP--------NLRRCVACRKLAPKQSLWRVVRV-----Y-P---------SQQ-----VQ-LD--
dbj|BAC90592.1|       --MSSYNRRSPPSRIRLI-----SNPAAVNPKK------------NQRRCVSCRRTGERESFWRIVRV-----H-P---------GWQ-----IQ-LN--
ref|WP_012306383.1|   --------------------------------M--EP--------NLRRCVACRKLAPKQSLWRVVRV-----Y-P---------SQQ-----VQ-LD--
gb|MBW4471632.1|      -----------------MKP-------------------------NDRRCVSCRKVASKQEFWRVVRVHSAQQL-S---------PPV-----VQ-LDHG
ref|WP_326498313.1|   ------------------MRSLKLRSHPVLFFM--LP--------NYRRCICCRKVAPKAEFLRIVRL-----H-P---------SHE-----IA-IN--
ref|WP_071782506.1|   ------------------------------------------MK-NQRRCVACRRLGAKESFWRFVRL-----Y-S---------SDQ-----VQ-LD--
tpg|HIK25275.1|       ---------------------------------------M---TLSQRRCVSCGRLAERSEFWRVVRC-----W-P---------DQR-----VQ-LD--
gb|NJK36543.1|        --------------------------------M----------PSNFRRCVVCRKLAPKQAFWRVVRD-----F-S---------SHQ-----VQ-LD--
gb|NEP78655.1|        -----------------------MKP-------------------NYRRCVSCKKLAPKEDFWRVIRL-----K-K---------SST-----IR-LD--
gb|NCO76359.1|        -----------------ML-------------R-KNK--------NHRRCISCNRIADRQQFWRIIRN-----Y-P---------EHQ-----IT-LD--
ref|WP_017293473.1|   -----------------------MKQPCPK-HK------------NYRRCISCHLIGDRSLFWRIVKN-----Y-P---------DHN-----IT-LD--
ref|WP_104387117.1|   --------------------------------M-K-P--------NYRRCISCRQVNLKQEFWRIVRV-----F-P---------SGN-----VQ-LD--
gb|TAD79778.1|        ------------------------------------------MPPNYRRCLICRRTAHRSEFWRIVRL-----A-N---------SGQ-----VQ-LN--
ref|WP_066345584.1|   --------MKN-LN----KQ----------------KN--------YRRCISCHKVGERKEFLRIVRV-----Y-P---------EQ---K--VS-LDQG
gb|MCL2931732.1|      -----------------MK-----------------P--------NYRRCVSCKKLSPKEEFWRVVRS-----K-K---------SDT-----IR-LD--
ref|WP_046277840.1|   ------------------------------------------MPPNCRRCVACRQAAPKETFWRVVRV-----Y-S---------SHQ-----VQ-LD--
gb|NJK54848.1|        --------------------------------M---NQ------INYRRCVSCRKIAPKQSFWRVVRL-----A-S---------SCQ-----I------
gb|MDX2244890.1|      --------------------------------M--QP--------NIRRCVVCRKLAPKQEFWRVVRV-----H-P---------SHA-----IV-LD--
ref|WP_215607325.1|   ---------MQ-QN--------------------------------YRRCVSCRQVTHRKNLLRVVRT-----Y-P---------TG---V--VQ-LNKG
ref|WP_315862757.1|   --------------------------------M--AP--------NDRRCVSCRRVAPKSAFLRIVRL-----H-P---------THT-----VQ-LD--
gb|MCL2923276.1|      -----------------MK-----------------P--------NYRRCVSCKKLSPKEEFWRVVRS-----K-K---------SAT-----IS-LD--
gb|NJL46250.1|        --------------------------------M--PPN--------YRRCVSCRRVAPRQQFWRVVRS-----S-A---------DGT-----TA-VD--
gb|MBW4534770.1|      --------------------------------M---NQ------INYRQCVSCRKVAPKQSFWRVVRV-----A-S---------SYA-----V------
ref|WP_015165576.1|   ---------MA--P----------------------KD--------YRRCISCRRLDHRDHFWRVVRS-----R-D---------SSGQIS--VQ-LDQG
gb|NJN60647.1|        --M------SQ---------------K------------------NDRRCVSCGEIGAKSSFWRIVRV-----H-P---------SHQ-----LQ-LD--
ref|WP_190497765.1|   --------------------------------M--PPN--------DRRCVSCRRVAPKSDFWRVVRL-----A-K---------TTT-----VC-LD--
gb|TAF56653.1|        --M------SN---------------K------------------HDRRCISCGEIGPKSSFWRIVRV-----H-P---------SHQ-----LQ-LD--
ref|WP_300873631.1|   ---------------------------------------M---TLPQRRCVSCGRLAERSEFWRVVRC-----W-P---------DQR-----VQ-LD--
gb|MBW4581861.1|      ---------------------------------------M---KPNYRRCVSCRKVALKEEFWRIVRL-----S-SPQAHT----SSL-----VE-LD--
ref|WP_193935386.1|   --------------------------------M-K-P--------NYRRCISCRQVNLKQEFWRIVRV-----F-P---------SGN-----VQ-LD--
ref|WP_124145084.1|   -----------------------MKP-------------------NYRRCVSCKKLAPKEDFWRVIRI-----K-K---------SST-----IR-LD--
gb|NJK99433.1|        MENCLIGSIAGG-----------------RGAS--PTH------VNLRRCVVCRHLAPKATFWRIVRV-----Y-P---------SLA-----VQ-LD--
gb|WRH67481.1|        ------------------------------------------MK-NQRRCVACRQLGEKESFWRFVRL-----H-P---------SHR-----VQ-LD--
ref|WP_071789028.1|   ---------MP-IN--------------------------------HRRCISCRKVVHRSQLLRVVRT-----H-P---------TG---T--VQ-FNKG
ref|WP_072041293.1|   --------------------------------M--KP--------NHRRCVSCRAIAPKPNLWRVVRQ-----H-D---------TGT-----VQ-LD--
gb|MBD1821041.1|      -----------------------M-----------PL--------NYRRCLSCRKVAPKEEFLRVVRL-----Y-P---------SHE-----IA-IN--
ref|WP_293079557.1|   -----------------------MKP-------------------NYRRCVSCKKLAPKEDFLRVVRL-----K-K---------SST-----IR-LD--
ref|WP_310422503.1|   ---------MT-KD--------------------------------YRRCVSCRKVAHRSEFWRLVRS-----H-P---------SS---Q--II-IDRG
gb|MDJ0591465.1|      --------------------------------M--K-Q------KNYRRCVSCRRVGPKQSFWRVVRL-----A-S---------NHQ-----VE-LE--
ref|WP_023172451.1|   -------------------------------MK------------HQRRCISCRRLDDRASFWRIVRV-----H-P---------GWQ-----IQ-LN--
ref|WP_036533928.1|   -----------------------------MT----TP--------NTRRCISCGKMDLKSAFWRVVRV-----H-P---------SQM-----VQ-LD--
ref|WP_162422901.1|   -----------------MP-----------------P--------NYRRCISCRKAAPKEEFLRVVRL-----H-P---------SHD-----IV-LN--
dbj|GGA05509.1|       M-------SKVLIKQKVK-----MKP-------------------NYRRCVSCKKLAPKEDFLRIVRL-----K-K---------SST-----IR-LD--
ref|WP_066120908.1|   --------MKN-PN----KQ----------------TN--------YRRCISCHLINDRTNFWRIVKN-----H-P---------DN---N--IT-LDVG
```

```
gb|UFP97105.1|          -----------------M-----SNPAAVNPKK------------NQRRCVSCRRTDARESFWRIVRV-----H-P---------GWQ-----IQ-LN--
ref|WP_019504603.1|     --------------------------------M---NQ------INYRRCISCRQLAPKKSLWRVIRL-----A-H---------THK-----I------
dbj|BAU14687.1|         -----------------------M-----------PS--------NLRRCLSCRKVAPKEEFLRVVRL-----F-P---------SHE-----IS-ID--
gb|MCY7272735.1|        --------------------------------M--EP--------NYRRCVSCRRSAPKYEFWRVVRS-----H-S---------SGT-----IV-LD--
ref|WP_202219931.1|     -----------------------MKP-------------------NYRRCVSCKKLAPKEDFLRIVRL-----K-K---------SST-----IR-LD--
ref|WP_310488256.1|     --MNELKPNSEGQRAKTV-----KHK-------------------DYRRCVSCRKVAARSEFWRVVKC-----H-P---------TNT-----IA-IDLG
gb|MBE9099686.1|        --------------------------------M--DK--------NYRRCISCRKVAPKSDFWRIVRV-----H-P---------SQE-----LQ-LD--
gb|MBU6228775.1|        --------------------------------M----------LSNYRRCISCRRLAPRSEFFRVVRS-----H-P---------YST-----IE-LN--
gb|NKB17092.1|          --------MAP-KL--------------------------------YRRCICCQRFAPREEFWRVVRV-----R-TEE--L----HT---L--IQ-LDRG
gb|NES69150.1|          -----------------------MKQ-------------------NYRRCVSCKKLAPKEDFWRVIRL-----K-K---------SST-----IR-LD--
gb|MBW4553259.1|        --MT---------------------K-------------------NYRRCISCRRLSLKTDFLRVVRV-----Y-P---------AQT-----IT-LN--
gb|PSP19040.1|          --------------------------------M-AESY---------RRCIGCRQLAPKSALWRIVRT-----H-P---------SGE-----VR-LD--
ref|WP_106312640.1|     --M---------------------LK-------------------DWRRCVCCRKVAHKKEFWRVVKA-----H-P---------SNL-----IK-IDLG
ref|WP_204104579.1|     --------------------------------M--KPN--------LRRCISCRKIAPKNAFWRIVRQTGGDVN-S---------KQT-----VQ-LD--
gb|MCU0552418.1|        --------------------------------M--IP--------NHRRCVCCRKVAPKAEFLRIVRL-----H-P---------SHK-----IA-IA--
gb|MBW4442872.1|        -----------------------M-----------QT--------CHRRCLSCRKVAPKSEFLRIVRL-----Y-P---------SHE-----IS-IE--
ref|WP_106254952.1|     -----------------------------M-----KP--------NYRRCISCRKVALKQEFWRLVRV-----S-S---------TQEGSP-LVV-LD--
emb|CAA9550821.1|       ------------------------------------------MELNHRRCVSCRTLRPKQFFWRVIRD-----H-S---------SGK-----VR-LD--
tpg|HCF29536.1|         --------------------------------M--KP--------NYRRCISCSRIGLKSEFWRIVRL-----N-C---------SGQ-----LQ-LD--
gb|MCA1903264.1|        --------------------------------M--APV-------GFRRCIVCRKLADRGEFWRIVRE-----------------GTT-----VK-LD--
gb|NEQ23162.1|          --------------------------------M--PI--------NDRRCVSCRRLAPKSEFWRIVRD-----C-S---------SGA-----VR-LD--
ref|WP_293133712.1|     -----------------MK-----------------P--------NYRRCISCKKVAPKEEFLRVVRS-----K-K---------SDS-----IK-LD--
ref|WP_015157060.1|     --------------------------------M---K-------PNYRRCISCGRVALKSEFWRIVRD-----R-D---------SGE-----LQ-LD--
ref|WP_121971475.1|     -----------------------MAE------K------------NYRLCISCRQTAHRDTLWRVVRT-----F-P---------DHQ-----VQ-LD--
dbj|BAZ46326.1|         --------------------------------M--K-P------INYRCCISCRRVAPKISFWRVVRL-----A-A---------NHQ-----IR-LD--
gb|MBW4527443.1|        -----------------------M-----------QT--------CHRRCLSCRKVAPKSEFLRIVRL-----Y-P---------SHE-----IS-IK--
gb|NEN88813.1|          -----------------------MKP-------------------NYRRCVSCKKLAPKEDFLRVIRL-----K-K---------SST-----IK-LD--
gb|MDX2273513.1|        --MFTSAQSPR---------------Q------------------PLRCCVSCRLLAPRSQFWRVVRV-----A-S---------SQQ-----VQ-LD--
ref|WP_293062802.1|     -----------------------MKP-------------------NYRRCVSCKKLAPKEDFLRVIRL-----K-K---------SST-----IK-LD--
ref|WP_218081389.1|     --------MTQAPH----VA----------------EN--------LRICLACRRKSGRDEFWRVVRV-----W-P---------SH---E--VQ-LDQG
ref|WP_015168983.1|     --------------------------MK--------P-------N-LRRCLSCQKVDLKQEFWRVVRV-----------------DS---A--VK-LDV-
ref|WP_287129737.1|     --------M---PE----PI----------------IQ--------TRSCLACRRKDGRESFWRVVRV-----W-P---------SH---Q--VQ-LDWG
gb|MCL2927522.1|        -----------------MK-----------------P--------NYRRCVSCKKLAPKEEFWRVVRS-----K-K---------SDT-----IE-LD--
ref|WP_192153424.1|     --------------------------------M---K-------PNYRRCISCGRVALKSEFWRIVRD-----R-D---------SGE-----LQ-LD--
gb|NJK69321.1|          --------------------------------M----------KPNYRRCASCRKIAPKEAFWRAVRV-----Y-P---------SHQ-----VQ-LD--
gb|MBP0016486.1|        --------------------------------M--QKN--------LRRCIACRKIAPKNHFWRIVRQANDTVH-P---------EQT-----VQ-LD--
gb|PZO20693.1|          -----------------------MAQ------K------------NHRLCISCRQTAHRDTLWRIVRT-----F-P---------SRH-----VS-LN--
gb|MCH9055185.1|        MSM--------------------GIP-------------------Q-RRCVACGRVANRTEFWRIVRR-----W-P---------DHA-----VK-VG--
ref|WP_099799890.1|     --M--------------------GIP-------------------Q-RRCVACGRVANRTEFWRIVRR-----W-P---------DHA-----VK-VG--
gb|MDJ0773676.1|        --------------------------------M---K-------PNYRRCISCRKVGLKEEFWRIVRV-----F-S---------SGQ-----VQ-LD--
gb|OWY68048.1|          --------------------------------M---K-------PNYRRCISCGRVALKSEFWRIVRD-----R-D---------SGE-----LQ-LD--
gb|MCX7595673.1|        --------------------------------M---K-------PNYRRCISCRKVGLKNEFWRIVRV-----F-E---------SGQ-----VQ-LD--
gb|NEQ50465.1|          --------------------------------M-QQNY---------RRCVSCRQVSHRNNLLRVVRT-----A-P---------TGV-----VT-FN--
gb|NJN30919.1|          --------------------------------M--RRA------TNVRCCVVCRRAGPKTSFWRVVRD-----H-G---------SGL-----VQ-LD--
gb|MDX2254087.1|        --------------------------MA--------L-------KLERRCVSCHCVAPRDRFWRIVRVPM---T-K----T----DG---E--RS-PDAQ
gb|AFY96277.1|          -----------------------------------------MLK-DWRRCICCRKVAHKNEFWRVVKA-----H-P---------DNL-----IT-ID--
ref|WP_229640858.1|     --------------------------------M--K-Q------KNYRRCVSCRRNRPKQSFWRVVRI-----A-L---------NHQ-----IE-ID--
ref|WP_068815920.1|     --------------------------------M--EP--------NYRRCVSCRRSAPKDEFWRVVKS-----H-S---------SGT-----IV-LD--
gb|MCY7336460.1|        --M---------------------VK-------------------DYRRCVSCRKLAPRNEFWRVVRS-----H-P---------SHT-----VT-IEL-
gb|MBS9769718.1|        -----------------MK-----------------P--------NYRRCVSCKKLAPKEEFWRVVRS-----K-K---------SDT-----IK-LD--
ref|WP_190352545.1|     -----------------------MDDVK--------PL-----CKLERRCVSCQSVAPRDRFWRVVRVPH---A-DHP--Q----QF---Q--VQ-LDV-
gb|PZV14006.1|          --------------------------------------MA---LKLHRRCVSCQRLSQRDDFWRVVRV-------SIAPID----NSSIEVATKH-----
gb|MDE5085023.1|        -----------------MK-----------------P--------NYRRCVSCKKLSPKEEFLRVIRS-----K-E---------SDT-----IK-LD--
```

```
ref|WP_293122704.1|   -----------------------MKP-------------------NYRRCVSCKKLAPKEDFLRVIRL-----K-K---------YST-----IK-LD--
ref|WP_319421161.1|   --------------------------------M---KQ------INYRRCVSCRRVAPKESFWRIVRL-----Q-R---------NHQ-----I------
ref|WP_295619447.1|   -----------------MNESNSRDSAQVEKIK-AAK--------NERRCVSCRKIAPKHEFWRIVKCH------P---------SKK-----IA-IDLS
gb|MCL2936457.1|      -----------------MK-----------------P--------NYRRCVSCKKLSPKEEFLRVIRS-----K-E---------SDT-----IK-LD--
gb|MDE5075942.1|      -----------------MK-----------------P--------NYRRCVSCKKLSPKEEFLRVIRS-----K-E---------SDT-----IK-LD--
ref|WP_293158219.1|   -----------------------MKP-------------------NYRRCVSCKKLAPKEDFLRVIRL-----K-K---------SST-----IR-LD--
gb|OLP17560.1|        -------------------MAELCASSEKPPKK--RP--------GYRRCISCGRIAPKHEFFRVVRV-----C-S---------SQE-----LQ-LN--
gb|NJM49111.1|        --------------------------------M--SQQ------TQLRCCVTCRRLAPKQTFWRVVRR-----Y-P---------SHE-----V------
ref|WP_244349359.1|   -----------------MS--G-GDPL--------------------RRCVACRQLFPRSQLWRLVRQFD---------------SHQ-----VL-LE--
ref|WP_232432138.1|   --------------------------------M-NLSGEGKVLK-DWRRCICCRKVAHKNEFWRVVKA-----H-P---------DNL-----IT-ID--
ref|WP_193989930.1|   ---------MQ-RN--------------------------------YRRCVSCRRVAHRDDLLRVVRT-----Y-P---------TG---A--VQ-CHYG
ref|WP_235280105.1|   -----------------MEPVS-RDPL--------------------RRCVACRQLFPRSQLWRLVRQFD---------------SHQ-----VL-LE--
ref|WP_166274652.1|   --------MVQ-VG--------------------------------VRRCISCRRLAHREQFLKVVRL-----S-C---------SG---Q--VV-IGMG
ref|WP_255131202.1|   ---------MT--------------------------T--------MRRCVSCRNLVDRQKLWRVVRL-----G-G---------GE---G--IQ-LDQG
ref|WP_264324473.1|   --------------------------------M--EDEAL---IKNHRRCVSCRKVAPKSDFWRIVRQ-----H-P---------GHQ-----LV-LD--
ref|WP_310410468.1|   -------------------MVKIAD----------RK--------DYRRCISCRKFAPKSEFWRVVKC-----H-P---------SKQ-----VA-ID--
ref|WP_259727590.1|   ---------MN--P-----A----------------PT--------LRRCVSCRQLRDRRLLWRVIRL-----P-G---------RE------VV-LDHG
ref|WP_115093230.1|   ---------MS-------ER----------------PV--------LRRCVACRQLLDRRQLWRVIRD-----Y-Q---------DG------VL-LDAG
gb|MAH58821.1|        ---------MT-------SK----------------PI--------LRRCVACRQLLDRRQFWRVIRD-----H-K---------DG------IL-LDSG
ref|WP_259703109.1|   ---------MN--P-----A----------------PT--------LRRCVSCRQLRDRRLLWRVIRL-----P-G---------RE------VV-LDHG
gb|NDC14930.1|        ---------MS--S-----K----------------PV--------LRRCVSCRELLDRQQLWRIVRL-----A-E---------GG------IG-LDQG
ref|WP_309742497.1|   -------------------MVKIAD----------RK--------DYRRCISCRKFAPKSEFWRVVKC-----H-P---------SKQ-----VA-ID--
ref|WP_036487578.1|   --------------------------------M--KP--------NYRCCISCRQLAPKETLIRVVRV-----H-P---------SYA-----VQ-LD--
gb|NJN76171.1|        --------------------------------M----K------PNYRRCISCRSVGVKEQFHRIVRE-----H-A---------TGE-----L------
gb|MBJ7364496.1|      ---------MT--------------------------T--------MRRCVSCRNFVDRQKLWRVVRL-----G-G---------GE---G--IQ-LDQG
gb|NBV69567.1|        -----------------------MATL--------------------RRCISCRSLVDRQHLWRVVRL-----A-G---------GAG-----VQ-LD--
gb|NJM97597.1|        --------------------------------M----ADK---N--IRRCVSCHRIAAKDAFWRVVRT-----F-P---------DHH-----VQ-LD--
gb|MCY7407389.1|      --------------------------------M--TI--------NQRCCISCRRLADKSEFWRVVRL-------N---------NRE-----IS-IN--
ref|WP_254933944.1|   -----------------------------MS-S--RP--------VLRRCVSCRELLDRQQLWRIIRL-------A---------EGG-----IG-LD--
ref|WP_017327771.1|   -----------------------------MPAS--QE--------PVRRCIACGKLAPRHEFFRVVRQ-----F-E---------TGQ-----VQ-LN--
ref|WP_309728133.1|   -----MKSNSEDRSVKTD-----KHK-------------------DYRRCVSCRKVAARQEFWRVIKC-----H-P---------TNA-----IA-INLG
gb|MCY7367819.1|      ---------MT-KD--------------------------------FRRCVSCRKIALRTEFWRVVKC-----H-P---------SN---Q--IA-IDLG
gb|MEC8605584.1|      --MS-------------------TQP-------------------TLRRCVSCRELFDRRQLLRIIRL-----A-G---------GAG-----VV-LD--
gb|MBW4662466.1|      --------------------------------M-Q-P--------NYRRCVSCRQVAPKADLLRIVRV-----F-P---------SRA-----VQ-LD--
ref|WP_190401439.1|   --------------------------------------MA---LKLHRRCVSCQCVARRDHFWRVVRL-------PIAVGN----DHDQNQHLNQ-----
ref|WP_161823453.1|   --------------------------------M--EP--------NLRRCVSCRRVAPKSAFWRVLRL-----A-H---------NGQ-----VQ-LD--
gb|MCS6943628.1|      --------------------------------M-E-KRIKQ---KNYRRCISCGLYAPKEQFWRVVRN-----Y-P---------DGH-----IT-LD--
ref|WP_255116400.1|   --------------------------------M-SGH-------AVLRRCVTCRQLLDRQQLWRVVRL-----K-D----------GG-----LS-LD--
gb|AFZ48415.1|        -----------------------------M--T--KK--------NYRRCVSCGLIAPKNHFCRVVRN-----F-P---------DHK-----IT-LD--
gb|NUN65110.1|        -----------------------------------MPLKL------YRRCVSCQCVAQRDNFWRVVRV-----PIAQNTAANLLMDKA-----KV-PDSN
ref|WP_115022254.1|   ---------MS-------ER----------------PV--------LRRCVACRQLLDRRQLWRVIRD-----H-Q---------DG------VL-LDAG
gb|MCX5962897.1|      --------------------------------M--TI--------NQRCCISCRRLADKSEFWRMVRL-------N---------NRE-----IS-IN--
gb|MBM5807801.1|      --------------------------------M--S---E---RSVPRRCVTCRQLLDRRQLWRVIRL-----A-----------DGG-----LA-LD--
gb|MBV2350088.1|      --------------------------------M--S---E---RSVLRRCVTCRELLDRRQLWRVIRL-----A-----------DGG-----LA-LD--
gb|MCG9892687.1|      --M---------------------EI-------------------NYRRCISCRATAPKPYFLRVVRV-----H-P---------TGE-----VR-LH--
gb|MBJ7492561.1|      -----------------------------MK-G--KP--------VLRRCVTCRELLDRQQLWRVVRL-------Q---------DGS-----LA-LD--
emb|CAI8160962.1|     ---------MT-------SR----------------PV--------LRRCVACRQLLDRRQLWRVIRD-----H-Q---------DG------VL-LDEG
gb|PSI02062.1|        -----------------------MATL--------------------RRCISCRSMVDRQHLWRVVRL-----A-G---------GAG-----VQ-LD--
gb|OON12454.1|        -----------------------MATL--------------------RRCISCRSMVDRQHLWRVVRL-----A-G---------GAG-----VQ-LD--
gb|NBO27849.1|        -----------------------MATL--------------------RRCISCRSLVDRQHLWRVVRL-----A-G---------GAG-----VQ-LD--
ref|WP_255008768.1|   -----------------------------MS-A--VP--------TLRRCVACREVLDRRQLWRVIRL-------A---------AGG-----LA-LD--
ref|WP_255098738.1|   -----------------------MATL--------------------RRCISCRSLVDRQHLWRVVRL-----A-G---------GAG-----VQ-LD--
ref|WP_255096693.1|   -----------------------MATL--------------------RRCISCRSLVDRQHLWRVVRL-----A-G---------GAG-----VQ-LD--
```

```
ref|WP_296444202.1|   --------------------------------M--S---E---RSVLRRCVTCRELLDRRQLWRVIRL-----A-----------DGG-----IA-LD--
ref|WP_211167718.1|   --------------------------------------MA---LKLHRRCVSCQCIAMRDNFWRVVRV-------PIA---------QADIHRTDGNPQQ
gb|NMF59328.1|        -------------------------------------------MKLHRRCVSCQCIAMRDNFWRVVRV-------PIA---------QADIHRTDGNPQQ
ref|WP_174235275.1|   --------------------------------------MA---LKLHRRCVSCQSVALRDSFWRIVRV-------PIAQDH----QILGDR-------DS
gb|MBM5799360.1|      --------------------------------M-SARP-------VLRRCVSCRQLLDRNELWRVIRL-----A-N---------GGG-----VQ-LD--
ref|WP_142655983.1|   --------------------------------------MA---LKLLRRCVSCQCIASRDTFWRVVRV-------PIA--S----TDENLIEAKSKHDQN
gb|MCP4972515.1|      ---------MN-------QR----------------PV--------LRRCVACRKLLNRQQLWRVIRD-----H-Q---------DG------VV-LDAG
ref|WP_043692418.1|   ---------MN-------ER----------------PI--------LRRCVACRQLLDRRQLWRVVRD-----H-R---------DG------VL-LDTG
ref|WP_011936384.1|   --MS-------------------AQP-------------------TLRRCVSCRELFDRRQLLRIIRL-----A-G---------GAG-----VV-MD--
gb|MCA6503255.1|      --------------------------------------MT---LKLHRRCVSCQCVAIRDTFWRVVRV-------PIA--S----TDENPNEAKAKHDQN
ref|WP_131456745.1|   ---------MN--I----DR----------------PI--------LRRCVACRQLLDRRQLWRVVRD-----H-Q---------DG------VL-LDAG
gb|MCB4394176.1|      ---------MS-------ER----------------PV--------LRRCVACRQLLDRRQLWRVIRD-----H-Q---------DG------VL-LDAG
gb|MCS5706208.1|      --------------------------------M-S---S----RTVLRRCVSCRELFDRTQLWRVIRL-----A-E---------G-G-----MA-LD--
gb|MEB3105445.1|      --------------------------------M-S---S----RAVLRRCVSCRELFDRTQLWRVIRL-----A-E---------G-G-----MA-LD--
gb|MCL1492198.1|      --------------------------------------MT---LKLHRRCVSCQCIAIRDTFWRVVRV-------PIA--S----TDENLIEAKAKHDQN
ref|WP_255087149.1|   ---------MN--A-----A----------------PT--------LRRCVSCRQLLDRRLLWRVIRK-----Q-G---------KD------VV-LDQG
ref|WP_269621837.1|   ---------MK-------HK----------------QI--------LRRCVTCRKLIDRKELWRVVRD-----Y-Q---------DG------VV-LDKG
ref|WP_310483930.1|   -----------------MNESNISNSERVIKTK-AAK--------NDRRCISCRKIAAKHEFLRAVKCH------P---------SKK-----IA-IDLG
ref|WP_156796781.1|   --------------------------------M-T-SGRP-----VLRRCVACRALMDRSQLWRVVRL-------A---------TGG-----LQ-LD--
gb|MDA0716923.1|      ---------MS--D-----R----------------AV--------LRRCVACRALLDRQQLWRVIRL-----A-E---------GG------IS-LDHG
tpg|HJN36909.1|       ---------MN-------DR----------------PI--------LRRCVACRALVDRRQLWRVIRD-----H-Q---------DG------LV-LDDG
gb|MCS5691210.1|      --------------------------------M-N-AEPR---RPVLRRCCACRRLADRREFWRVIRR-----A-D---------GDG-----LA-LD--
gb|NBQ18864.1|        -----------------------MATL--------------------RRCISCRSLVDRQHLWRVVRL-----A-G---------GAG-----VQ-LD--
ref|WP_255613922.1|   ----------------------------------------------MRRCVACRQLLDRRQLWRVIRD-----H-Q---------DG------VL-LDAG
ref|WP_201324463.1|   --------------------------------------MA---LKLHRRCVSCQCISIRDTFWRVVRV-------PIA--S----TDKNPNEANAKHDQN
ref|WP_011125798.1|   ---------MK-------RN----------------SV--------LRRCVACKKLVDRKQLWRVVRD-----Y-Q---------DG------IV-LDQG
gb|MBM5803857.1|      --------------------------------M-SAAP-------VLRRCVACRQLVDRNQLWRVIRL-----A-G----------GG-----LA-LD--
ref|WP_094591796.1|   --------------------------------M-SGK-------SVLRRCVTCRELLDRQQLWRVIRL-----A-N----------GG-----LA-LD--
ref|WP_197212874.1|   --------------------------------M-SRTVAA---RPVLRRCVACRVLRDRSELWRVIRL-----A-Q---------G-G-----LA-LD--
ref|WP_115120514.1|   ------------------------------------MNEP---RPVLRRCVACRKLLDRQQLWRVIRD-----H-R---------EG------VL-LD--
gb|MBM5785306.1|      --------------------------------M--S---E---RSVLRRCVTCRELLDRRQLWRVIRL-----A-----------DGG-----IA-LD--
ref|WP_038023507.1|   ------------------------------------MNEP---RPVLRRCVACRKLLDRQQLWRVIRD-----H-Q---------DG------VL-LD--
ref|WP_254938812.1|   -----------------------------MS-A--AP--------TLRRCVACRELLDRQQLWRVIRL-------A---------AGG-----LA-LD--
ref|WP_038553465.1|   ---------MS-------ER----------------PV--------LRRCVACRQLLDRRQLWRVIRD-----H-Q---------DG------VL-LDAG
gb|MDT7945133.1|      -----------------MP--G-ADPL--------------------RRCVGCRQLFPRSQLWRLVRQFD---------------SHQ-----VL-LD--
gb|MEB3326908.1|      ---------MT--C-----R----------------PV--------LRRCVSCRKLLDRQQLWRVIRL-----A-E---------GG------IC-LDQG
ref|WP_130129912.1|   ---------MK--T----PR----------------PV--------LRRCVACRQLLDRRQLWRVIRD-----H-Q---------DG------VL-LDVG
ref|WP_038001092.1|   ---------MK--T----PR----------------PV--------LRRCVACRQLLDRRQLWRVIRD-----H-R---------DG------VL-LDVG
ref|WP_258040662.1|   --------------------------------------MA---LKLNRRCVSCQSIAIRDTFWRVVRV-------PIA--S----TENKPNEANAKHDQV
gb|PSP26132.1|        -----------------------------------MK-------PNYRRCVSCRQVAKKESFWRIVRV-----H-P---------SGQ-----VQ-LD--
gb|MEB3304480.1|      ----------------------------------MAPQAR----PVLRRCVACRQLRDRQELFRVVRL-----W-A---------GGG-----LA-LD--
gb|NDC16240.1|        --------------------MSPG------------P--------TLRRCVACRELLDRQLLLRVIRL-----P-R---------AQG-----IV-LD--
tpg|HBC43141.1|       -------------------------------------------MKLHRRCVSCQCIAMRDNFWRVIRV-------PIK---------QSDIHRADGHPQQ
ref|WP_206339282.1|   ----------------------------------------------MRRCVACRQLLDRRQLWRVIRD-----H-Q---------DG------VL-LDAG
ref|WP_063406354.1|   ---------MN-------QR----------------PV--------LRRCVTCRQLLDRQQLWRVIRD-----H-Q---------EG------VV-LDEG
gb|NJM75400.1|        -----------------------M-----------EP--------NFRRCVSCRVVAHKNQLLRVVKE-----H-S---------TGK-----LE-VD--
ref|WP_259734545.1|   ---------MT--S-----S----------------PV--------LRRCVACRKLLDREQLWRVIRL-----A-E---------GG------IC-LDQG
gb|MBF2058462.1|      --MK-------------------KNK-------------------NWRRCLSCGHYAPKDDFIRVVRI-----F-P---------DHQ-----IA-IN--
ref|WP_197151498.1|   ---------MT--R-----S----------------PV--------LRRCVACRKLLDREQLWRVIRL-----A-E---------GG------IC-LDQG
ref|WP_254928579.1|   ---------MS--R-----R----------------PV--------LRRCVACRELLDRQQLWRVIRL-----A-E---------GG------IG-LDQG
ref|WP_323259706.1|   --------------------------------------MA---LKLHRRCVSCQCVAMRDTFWRVVRV-------PIA--------VRHPNEDKPNHNQV
gb|MBD2423618.1|      ---------MS--R-----R----------------PV--------LRRCVACRELLDRQQLWRVIRL-----A-E---------GG------IG-LDQG
gb|MBM5795377.1|      --------------------------------M--S---E---RSVLRRCVACRELLDRRQLWRVIRL-----A-----------DGG-----LA-LD--
```

```
ref|WP_281008927.1|    --------------------------------------MA---LKLHRRCVSCQCVAMRDTFWRVVRV-------PIA--------DRHPNEDKPNHNQV
gb|MCH2566529.1|       ---------MN-------QR----------------PV--------LRRCVTCRQLLDRQQLWRVIRD-----H-Q---------EG------VV-LDEG
gb|MEB3307264.1|       ---------MS--S-----R----------------PV--------LRRCVSCRQCFDRRNLWRVIRL-----A-G---------GG------IA-LDRG
tpg|HBH73361.1|        --------------------------------M-S----P---QPVMRRCVSCRQLCDRTLLWRVIRL-----A-A---------G-G-----IA-LD--
ref|WP_255146391.1|    --------------------------------M-AAS-------PTLRRCVSCRLLADRRQFLRVIRL-----A-G----------GG-----IA-LD--
ref|WP_271253996.1|    --------------------------------------MA---LKLHRRCVSCQCVALRDSFWRVVRV-------PIAQEY----QILGDR-------DS
gb|MEB3234896.1|       --------------------------------M-TAKV-------VLRRCVACRQLVDRDQLWRVIRL-----A-E----------GG-----IA-LD--
gb|MDA0886513.1|       ---------MT--S-----S----------------PV--------LRRCVACRKLLDREQLWRVIRL-----A-G---------GG------IC-LDQG
ref|WP_115019618.1|    ------------------------------------MNEP---RPVLRRCVACRELLDRQQLWRVIRD-----H-R---------EG------VL-LD--
ref|WP_011130895.1|    ---------MN-------QR----------------PV--------LRRCVTCRQLLDRQQLWRVIRD-----H-Q---------EG------VV-LDEG
gb|NDG24387.1|         --------------------------------M-SGR-------SVLRRCVTCRTLLDRQQLWRVIRQ-----A-E----------GG-----LA-LD--
gb|MBM5797070.1|       --------------------------------M-SAKV-------VLRRCVACRQLLDREQLWRVIRL-----A-E----------GG-----VV-LD--
gb|MCA6523574.1|       -------------------------------------------MKLHRRCVSCQCIAMRDNFWRVMRV-------PIK---------QSDIHRAGGHPQQ
gb|MCX5946264.1|       -----------------------------MN-G--KP--------VLRRCITCRELLDRQQLLRVIRL-------P---------DGS-----LA-LD--
gb|MAV11321.1|         -----------------------------MN-Q--KP--------ILRRCVACRQLLDRRLFWRVIRX-----H-K---------DG------VL-LD--
ref|WP_320673936.1|    ---------MN-------KS----------------TV--------LRRCVACRKLLDRQQLLRVIRD-----H-Q---------DG------VV-LTKG
gb|MCP9817453.1|       --------------------------------M-TGK-------PVLRRCVTCRELLDRQRLWRVIRL-----A-D----------GG-----LA-LD--
gb|NJK35889.1|         --------------------------------M-----------KGDRVCVSCRRVGPRESFWRVVRL-------P---------DGS-----IV-ID--
gb|NJL99010.1|         --------------------------------M--SSS------DPLRQCISCRMFAPRSQFWRVVR--------L---------SSE-----A--PQPD
ref|WP_193800382.1|    -----------------------------MP-P--KN--------Y-RRCVSCGLISSKHDFIRVVRN-----F-P---------SHH-----IT-IN--
ref|WP_063403536.1|    ---------MN-------QR----------------PV--------LRRCVTCRQLLDRQQLWRVIRD-----H-Q---------EG------VV-LDEG
gb|RMD73144.1|         --------MRK-PN----KI----------------KN--------YRRCISCQKIDHKSNFFRVVKN-----N-T---------QQ---I--IT-LDQG
gb|MCY7333721.1|       --------------------------------------MA---LKLHRRCVSCQCIALRDRFWRIVRV-------PIAPVS----DHDSKITPKN-----
gb|MDG2329318.1|       ---------MS-------AK----------------PI--------LRRCVACRQLQDRRHLWRVIRD-----H-K---------DG------VL-LDSG
gb|MCX5941757.1|       -----------------------------MN-G--KP--------VLRRCVTCRELLDRQQLLRVIRL-------P---------DGS-----LA-LD--
tpg|HJN33659.1|        ---------MN-------QR----------------PV--------LRRCVTCRQLLDRQQLWRVIRD-----H-Q---------EG------VV-LDEG
tpg|HIK19498.1|        -----------------MP--G-ADPL--------------------RRCVGCRQLFPRSQLWRLVRQFD---------------SHQ-----VL-LD--
tpg|HCX54226.1|        -----------------------------MN-E--RP--------ILRRCVACRQLLDRRQLWRVVRD-----H-R---------DG------VL-LD--
ref|WP_063399518.1|    ---------MN-------QR----------------PV--------LRRCVTCRQLLDRQQLWRVIRD-----H-Q---------EG------VV-LDEG
ref|WP_011359406.1|    ---------MS-------AK----------------PI--------LRRCVACRQLQDRRHLWRVIRD-----H-K---------DG------VL-LDFG
gb|MEB3185113.1|       --------------------------------M-R---C----RAVQRRCVSCRELFDRTQLWRVIRL-----A-D---------G-G-----VA-LD--
ref|WP_186470267.1|    ---------------MND-----VRP-------------------VLRRCVACRELLDRSLLWRVIRD-----H-Q---------NGV-----LL-DQ--
ref|WP_009628060.1|    --------------------------------------MA---LKLHRRCVSCQCIALRDNFWRVVRV-------PIAPEQ----NDQDDRDDRK--NIY
gb|MBE67324.1|         ---------MS-------AK----------------PV--------LRRCVACRQLQDRRHLWRVIRD-----H-K---------DG------VL-LDFG
ref|WP_272159772.1|    ---------MN-------AK----------------PI--------LRRCVACRQLQDRRHLWRVIRD-----H-Q---------DG------VL-LDFG
ref|WP_063419582.1|    ---------MN-------QR----------------PV--------LRRCVTCRQLLDRQQLWRVIRD-----H-Q---------EG------VV-LDAG
gb|MEB3240836.1|       --MS-------------------SKP-------------------TLRRCVSCRQLFDRDQLLRVIRL-----P-E---------GQG-----LS-LE--
gb|NBW63894.1|         --------------------------------M-SER-------AVLRRCVSCRQLLDRRQLWRVIRL-----A-G----------GG-----MA-LD--
gb|MBM5788852.1|       --------------------------------M-SGR-------SVLRRCVTCRSLLDRQQLWRVIRL-----A-D----------GG-----LA-LD--
gb|MEB3262778.1|       --------------------------------M-S----S---RPVLRRCVSCRELFDRSQLWRVIRL-----A-E---------G-G-----LA-LD--
ref|WP_296365315.1|    --------------------------------M--S---E---RSVLRRCVTCRQLLDRQQLWRVIRL-----A-----------DGG-----LA-LD--
gb|PZU97808.1|         -------------------------------------------MKLHRRCVSCQRIAQRDEFWRLVLV-------PIAQDS----AENELIAALGKPRRE
gb|MAN18293.1|         ---------------MNN-----VRP-------------------VLRRCVACRELLDRSLLWRVIRD-----H-Q---------NGV-----LL-DQ--
ref|WP_186583364.1|    ---------MK--P----QR----------------PI--------LRRCVACRQLLDRSLLWRIVRD-----Y-R---------DG------VL-LDHG
gb|MCS5699361.1|       --------------------------------M-N-AEPR---RPVLRRCCACRRLADRREFWRVIRR-----A-D---------GSG-----LS-LD--
ref|WP_186570491.1|    ---------MS-------ER----------------PV--------LRRCVACRQLLDRRQLWRVIRD-----H-Q---------DG------VL-LDAG
ref|WP_186501173.1|    ---------------MND-----VRP-------------------VLRRCVACRELLDRSLLWRVIRD-----H-Q---------NGV-----LL-DQ--
gb|MBL6794407.1|       ---------MS-------AK----------------PI--------LRRCVACRQLQDRRHLWRVIRD-----H-K---------DG------VL-LDLG
gb|MBL6880295.1|       ---------MS-------AK----------------PI--------LRRCVACRQLQDRRHLWRVIRD-----H-K---------DG------VL-LDLG
gb|MDB4653764.1|       ---------MS-------AN----------------PI--------LRRCVACRQLQDRRHLWRVIRD-----H-K---------DG------VL-LDLG
gb|RPF82291.1|         ---------MN-------DR----------------PI--------LRRCVACRQLLDRRLLWRVVRD-----Y-R---------DG------VL-LEEG
gb|MCH1457301.1|       ---------MS-------AK----------------PI--------LRRCVACRQLQDRRHLWRVIRD-----H-K---------DG------VL-LDLG
gb|MED5164851.1|       ---------MN-------QR----------------PV--------LRRCVTCRQLLDRQQLWRVIRD-----H-Q---------EG------VV-LEEG
```

```
gb|PZO43371.1|          --------------------------------------MA---LKLHRRCVSCQCIAMRDTFWRVVRV-------PIAANE----KYLTDKSLNE--SNV
ref|WP_028952666.1|     ---------MN-------DR----------------PI--------LRRCVACRQLLDRRLLWRVVRD-----Y-R---------DG------VL-LEEG
gb|MAI96565.1|          ---------MS-------AR----------------PI--------LRRCVACRQLLDRRQLWRVIRD-----H-Q---------DG------VL-LDRG
ref|WP_037988505.1|     ---------MS-------AK----------------PI--------LRRCVACRQLLDRRHLWRVIRD-----H-K---------DG------VL-LDLG
ref|WP_029553103.1|     --------------------------------M--S---E---RSVLRRCVTCRELLDRRLLWRVIRL-----A-----------DGG-----LA-LD--
ref|WP_106502372.1|     ---------MS--R-----R----------------AV--------LRRCVACRALLDRQQLWRVIRL-----A-E---------GG------LG-LDQG
dbj|GDX71743.1|         --------------------------------M-SGR-------SVLRRCVTCRTLLDRQQLWRVIRL-----A-H----------GG-----LA-LD--
ref|WP_036911320.1|     ---------MN-------QR----------------PV--------LRRCVTCRQLLDRQQLWRVIRD-----H-Q---------EG------VV-LDEG
ref|WP_011825091.1|     ---------MN-------QR----------------PV--------LRSCVTCRQLLDRQQLWRVIRD-----H-Q---------EG------VV-LDEG
gb|MBL6803472.1|        ---------MN-------PR----------------PV--------LRRCVACRQLQDRRSLWRVIRD-----H-K---------DG------VR-VDIG
ref|WP_048017238.1|     ---------MT--V-----R----------------AV--------QRRCVACRQLLDRSRLWRVIRL-----A-Y---------GG------IT-LDQG
gb|MAV12819.1|          ---------MN-------PR----------------PV--------LRRCVACRQLQDRRSLWRVIRD-----H-K---------DG------VR-VDIG
ref|WP_115010044.1|     ---------MS-------DR----------------PV--------LRRCVACRQLLDRRQLWRVIRD-----H-Q---------DG------VL-LDKG
ref|WP_006851947.1|     ---------MS-------DR----------------PV--------LRRCVACRQLLDRRQLWRVIRD-----H-Q---------DG------VL-LDEG
gb|NDG75793.1|          --------------------------------M--S---A---RPVLRRCVTCRQLLDRRQLWRVIRL-----S-----------EGG-----MA-LD--
gb|MEC8441465.1|        ---------MN-------QR----------------PV--------LRRCVACRQLQDRRSLWRVIRD-----H-Q---------DG------VR-VDVG
ref|WP_069789292.1|     -----------------------------MP-P--KN--------Y-RRCVSCGLLAPKNYFYQVVRN-----F-P---------HHQ-----IT-IN--
gb|WRL41094.1|          --------MRK-PN----KI----------------KN--------YRRCISCQKIAHKSNFFRVVKN-----N-A---------QQ---I--IT-VDQG
ref|WP_320002208.1|     --------MRK-SN----KI----------------KN--------YRRCISCQKIDHKSNFFRVVKN-----N-A---------QQ---I--IT-VDQG
ref|WP_205909688.1|     ----------------------------------------------MRRCVTCRQLLDRQQLWRVIRD-----H-Q---------EG------VV-LDAG
ref|WP_011432759.1|     -----------------MP--G-ADPL--------------------RRCVGCRQLFPRSQLWRLVRQFD---------------SHQ-----VL-LG--
gb|MDA7432491.1|        ---------MS-------AK----------------PI--------LRRCVACRQLQDRRHLWRVIRD-----H-K---------DG------VL-LDLG
gb|MEB3202259.1|        ---------MT--G-----R----------------PV--------LRRCVACRELLDRQQLWRVIRL-----A-E---------GG------IG-LDQG
gb|OIP77665.1|          ---------------------------------------------MHRRCVSCQRIAQRDEFWRLVLVPI---A-KDI--S----EN---E--LI-APIS
gb|MEB3159801.1|        ------------------------------------MNEP---RPVLRRCIACRQLLDRQQLWRVIRD-----H-Q---------NG------VL-VD--
gb|MED5263527.1|        ---------MN-------QR----------------PV--------LRRCVTCRQLLDRQQLWRVIRD-----H-Q---------EG------VV-LDEG
ref|WP_036919232.1|     ---------MT-------QN----------------RV--------LRRCVACRQILDRKQLWRVIRD-----C-Q---------DG------VV-LDKG
gb|MBM5790812.1|        --------------------------------M--S---S---KAVLRRCVTCRELLDRQQLWRVIRL-----A-----------EGG-----MA-LD--
gb|MED5384943.1|        ---------MN-------DR----------------PI--------LRRCVACRQLLDRRLLWRVVRD-----H-R---------DG------VL-LEQG
ref|WP_199310374.1|     --------------------------------------MA---LKLHRRCVSCQCIAMRDTFWRVVRV-------PIT--------DQHPKEDKPNHNQV
gb|MBD2316621.1|        -------------------------------------------MKLHRRCVSCQCIAMRDTFWRVVRV-------PIT--------DQHPKEDKPNHNQV
ref|WP_038546212.1|     ---------MN-------DR----------------PI--------LRRCVACRQLLDRRLLWRVVRD-----H-R---------DG------VL-LEQG
ref|WP_322771268.1|     --------------------------------M--S---E---RSVLRRCVTCRQLLDRKQLWRVIRL-----A-----------DGG-----LA-LD--
gb|WVL01204.1|          --------MRK-LN----KI----------------KN--------YRRCISCQKIDHKNNFFRVVKN-----N-A---------QQ---I--IT-VDQG
ref|WP_015218854.1|     --------MRK-PN----KI----------------KN--------YRRCISCQKIDHKSNFFRVVKN-----N-A---------QQ---I--IT-VDQG
ref|WP_055075296.1|     --------------------------------------MA---LKLHRRCVSCQCIALRDNFWRVVRV-------PIAPEQ----NDQDDRNDRE--NVH
gb|MEB3276547.1|        --------------------------------M-TARV-------VLRRCVACRQLLDREQLWRVIRL-----A-E----------GG-----IV-LD--
gb|MCB4428669.1|        ---------MS-------DR----------------PV--------LRRCVACRQLLDRRQLWRVIRD-----Y-Q---------DG------VL-LDEG
ref|WP_094510206.1|     --------------------------------M-ATP-------PTLRRCVSCRLLADRDQFLRVIRL-----A-G----------GG-----IA-LD--
gb|MDP6171750.1|        ---------MN-------QR----------------PV--------LRRCVSCRQLLNRQQLWRVIRD-----H-Q---------EG------VV-LDQG
ref|WP_223805489.1|     --------------------------------------MA---LKLHRRCVSCQCIALRDNFWRVVRV-------PIAPDQ----NDQDDRNDRE--NVH
ref|WP_011365029.1|     ---------MS-------DR----------------PV--------LRRCVACRQLLDRRQLWRVIRD-----H-Q---------DG------VR-LDEG
ref|WP_214339661.1|     --------------------------------M--S---E---RSVLRRCVTCRQLLDRQQLWRVIRL-----A-----------DGG-----LA-LD--
ref|WP_255099870.1|     --------------------------------M-APP-------PTLRRCVSCRLLADRDQFLRVIRL-----A-G----------GG-----IA-LD--
gb|TYQ31904.1|          -------------------------------------------MKLHRRCVSCQCIALRDNFWRVVRV-------PIAPDQ----NDQDDRNDRE--NVH
ref|WP_320666922.1|     ---------MS-------QR----------------PV--------LRRCVACRKLLDRKQLLKVTKD-----Y-Q---------DG------IV-LDGG
gb|MEB3297319.1|        ---------MS--S-----R----------------PV--------LRRCVSCRQLLDRTHLWRVIRL-----A-G---------GG------IA-LDQG
ref|WP_198953901.1|     ---------MT--P-----R----------------IV--------LRRCVACRALLDRQQLWRVIRL-----A-E---------GG------IG-LDQG
ref|WP_186595138.1|     ---------MN--P----QR----------------PI--------LRRCVACRQLLDRSLLWRIVRD-----Y-R---------DG------VL-LDDG
ref|WP_186493755.1|     ---------MS-------DR----------------PV--------LRRCVACRQLLDRRQLWRVIRD-----H-Q---------DG------VR-LDEG
gb|MEC7393122.1|        ---------MS-------DR----------------PV--------LRRCVACRQLLDRRQLWRVIRD-----H-Q---------DG------VR-LDEG
gb|QNJ16179.1|          ---------MN-------QR----------------PI--------LRRCVACRQLLDRRSLWRVIRD-----H-Q---------DG------VR-VDRG
gb|QNI91005.1|          ---------MN-------QR----------------PI--------LRRCVACRQLLDRRSLWRVIRD-----H-Q---------DG------VR-VDRG
```

```
gb|NJK59609.1|        --------------------------------M--AA-------KDVRRCVSCRRLAPRQEFWRVVRL-----A-D---------SRR-----VV-WD--
ref|WP_259728879.1|   --------------------------------M-ATS-------PTLRLCVSCRLLADRDQFLRVIRL-----A-G----------GG-----IA-LD--
gb|AUC59923.1|        -----------------------------MP-P--KN--------Y-RRCVSCGLLAPKNYFSQVVRN-----F-P---------HHQ-----IT-IN--
ref|WP_185186942.1|   -----------------------------MK-Q--QP--------VLRRCVTCRALLDRQQLLRVIRL-------A---------EGG-----MA-LD--
ref|WP_255599781.1|   ----------------------------------------------MRRCVACRQLLDRRQLWRVIRD-----Y-Q---------DG------VL-LDEG
ref|WP_012196167.1|   --------------------------------M-NQK-------TVLRRCVACRKLIDRQQLWRVTRD-----H-Q----------GG-----VV-LD--
ref|WP_115080914.1|   ---------MS-------DR----------------PV--------LRRCVACRQLLDRRQLWRVIRD-----H-Q---------DG------VL-LDQG
gb|MBR75694.1|        ---------MN-------QR----------------PI--------LRRCVACRQLLDRRSLWRVIRD-----H-Q---------DG------VR-VDGG
gb|RNC90972.1|        ---------MS-------QR----------------PI--------LRRCVACRQLLDRRSLWRVIRD-----H-Q---------DG------VR-VDGG
gb|MCB4389873.1|      ---------MS-------DR----------------PV--------LRRCVACRQLLDRRQLWRVIRD-----H-Q---------DG------VL-LDEG
ref|WP_303534944.1|   ---------MS-------DR----------------PV--------LRRCVACRQLLDRRQLWRVIRD-----H-Q---------DG------VL-LDEG
ref|WP_186495900.1|   ---------MS-------DR----------------PV--------LRRCVACRQLLDRRQLWRVIRD-----H-Q---------DG------VL-LDKG
ref|WP_320676133.1|   ---------M--------QP----------------TV--------LRRCVACRKLIDRKQLLRVIRD-----H-Q---------DG------VV-LDRG
ref|WP_011933969.1|   ---------MK--P----QR----------------PI--------LRRCVACRLLLDRSLLWRIVRD-----H-R---------DG------VL-LDHG
ref|WP_011127466.1|   ---------MS-------QH----------------PI--------LRRCVACRQLLDRRSLWRVIRD-----H-Q---------DG------VR-VDGG
gb|MCX5948231.1|      --------MSA-PN----RR----------------PV--------LRRCVACRNLRDRQLLWRLIRQ-----A-D---------GS------IA-LDQG
gb|MCS6960746.1|      --M--------------------APP-------------------LFRRCVICRRLAHRDEFWRIVRE-----------------GNT-----VK-LD--
ref|WP_217901569.1|   --------------------------------------MA---LKLHRRCVSCQCVAMRDTFWRVVRV-------PIA--------DQHHNEDKPKHDQV
gb|OYQ64342.1|        -------------------------------------------MKLHRRCVSCQCVAMRDTFWRVVRV-------PIA--------DQHHNEDKPKHDQV
ref|WP_190398881.1|   --------------------------------------MA---LKLHRRCVSCQCVALRDNFWRVVRV-------PIALHA----NQSANQSANQQKSNS
ref|WP_114989435.1|   ---------MS-------DR----------------PV--------LRRCVACRQLLDRRQLWRVIRD-----H-Q---------DG------VL-LDKG
gb|MCB4407378.1|      ---------MS-------DR----------------PV--------LRRCVACRQLLDRRQLWRVIRD-----H-Q---------DG------VL-LDEG
ref|WP_186479837.1|   ---------------MND-----ARP-------------------VLRRCVACRELLDRSLLWRVIRD-----H-R---------DGV-----LL-DQ--
ref|WP_247910210.1|   -----------------------------MK-Q--QS--------VLRRCVTCRALLDRQQLLRVIRL-------A---------EGG-----MA-LD--
dbj|GCE64220.1|       -----------------------------MS-K--AI--------VMRRCIACKKLLSRQHLWRVVRD-----H-Q---------NG------IQ-LD--
ref|WP_099812864.1|   -----------------MA--G-ADPL--------------------RRCVGCRQLFPRSQLWRLVRQYD---------------THQ-----VL-LD--
gb|MBT65681.1|        ---------MN--V----SR----------------PV--------LRRCVACRQLLDRSMLWRVIRD-----H-Q---------NG------VC-LDQG
gb|MEB3183095.1|      --------------------------------M-T---PR---RPVLRQCVACRRVADRGELLRVVRL-----A-----------GGG-----LA-LD--
ref|WP_254995692.1|   ---------MT--P-----R----------------IV--------LRRCVACRALLDRQQLWRVIRL-----A-E---------GG------IG-LDQG
ref|WP_150884534.1|   ---------------MM------ARP-------------------VLRRCAACRLLADRSTLWRVIRD-----H-E---------DGV-----VL-DQ--
ref|WP_006043237.1|   ---------MK--P----QR----------------PI--------LRRCVACRQLLDRNLLWRIVRD-----H-R---------DG------VL-LEQG
gb|NDC34872.1|        --------------------------------M-S----S---RPVLRRCVACRELLDRAQLWRVVRL-----A-T---------G-G-----IQ-LD--
ref|WP_255613750.1|   ----------------------------------------------MRRCVACRQLLDRRQLWRVIRD-----H-Q---------DG------VL-LDEG
gb|OUT75469.1|        --MN-------------------SKA-------------------TLRRCVSCRQLHDRCELLRVIRL-----P-G---------GQG-----LA-LD--
gb|NCG15537.1|        ---------MN--V----SR----------------PV--------LRRCVACRELLDRSMLLRVIRD-----H-Q---------EG------VL-LDQG
gb|TGG79053.1|        --------------------------------M-SAR-------PVLRRCVACRCLRDRDQLWRVIRL-----A-V----------GG-----VA-LD--
ref|WP_115131857.1|   ---------MS-------DR----------------PV--------LRRCVACRQLLDRRQLWRVIRD-----H-Q---------DG------VL-LDKG
ref|WP_284500670.1|   ----------------------------------------------MRRCVACRQLLDRRQLWRVIRD-----H-Q---------DG------VL-LDKG
ref|WP_255616060.1|   ----------------------------------------------MRRCVACRQLLDRRQLWRVIRD-----H-Q---------DG------VL-LDEG
gb|RCL52291.1|        --MS-------------------SNV-------------------TLRRCVSCRQLHDRCKLLRVIRL-----P-S---------GQG-----LV-LD--
tpg|HAN46371.1|       --------------------------------M--AP-------KDHRRCVSCRRLAPRTEFWRVVRL-----A-G---------GRE-----VV-WD--
ref|WP_186587596.1|   ---------MN-------QR----------------PI--------LRRCVACRQLLDRSSLWRVIRD-----H-Q---------DG------VR-VDGG
ref|WP_011431137.1|   -----------------MA--G-ADPL--------------------RRCVGCRQLFPRSQLWRLVRQYD---------------THQ-----VL-LD--
ref|WP_186498867.1|   ---------MN-------QR----------------PI--------LRRCVACRQLLDRRSLWRVIRD-----H-Q---------DG------VR-VDRG
ref|WP_259720650.1|   -----------------------------MM-A--RP--------VLRRCVACGELFNREQLWRVIRL-------A---------QGG-----IG-LD--
gb|MDP6195789.1|      ---------MN-------QR----------------PV--------LRRCIACRQLLNRQQLWRVIRD-----H-Q---------EG------VV-LDQG
gb|RZO05221.1|        ---------MN-------QR----------------PI--------LRRCVACRQLLDRSSLWRVIRD-----H-Q---------DG------VR-VDGG
ref|WP_269603352.1|   ---------MS-------QS----------------PI--------LRRCVACKKVLDRKYFLKVTRD-----F-Q---------NG------VV-LSGG
ref|WP_036900583.1|   --------------------------------M-NQR-------TVLRRCVACRTLFDRQKLWKVTRD-----F-Q----------DG-----IV-LD--
ref|WP_048347195.1|   ---------MN--V----SR----------------PV--------LRRCVACRQLLDRSMLLRVIRD-----H-Q---------EG------VL-LDQG
ref|WP_197156911.1|   -----------------------------MM-A--RP--------VLRRCVACGELFNREQLWRVIRL-------A---------QGG-----IG-LD--
gb|MCX5969416.1|      --------MSA-RI----RR----------------PV--------LRRCVACRSLQDRQLLWRLIRQ-----A-D---------GS------IA-LDLG
ref|WP_259735584.1|   --------------------------------M--SVG------VVLRRCVSCRERQDRRLLWRIIRP-----F-D---------GKG-----VV-LEGP
```

```
emb|CAK6691010.1|     ------------------------------M-A--RP--------VLRRCVACGELFNREQLWRVIRL-------A---------QGG-----IG-LD--
ref|WP_067095672.1|   --------------------------------M--NT--S---RPVLRRCVACRELFDRSILWRVIRD-----H-----------RDG-----VL-LD--
ref|WP_269608866.1|   ---------MS-------QN----------------PI--------LRRCVACKKVLDRKYFLKVTRD-----F-Q---------NG------VV-LSGG
gb|MBU6251331.1|      --------------------------------M-S----S---RPVLRRCVACRQLLDRAQLWRVVRL-----S-T---------G-G-----VQ-LD--
ref|WP_115125622.1|   --------------------------------M--NT--S---RPVLRRCVACRELFDRSILWRVIRD-----H-----------RDG-----VL-LD--
gb|MAB54535.1|        ---------MN-------SR----------------PI--------LRRCVACRQLMDRNLLWRVIRD-----H-R---------DG------VL-LDSG
gb|MBM5813821.1|      ---------MT--P-----R----------------IV--------LRRCVACRALLDRQLLWRVIRL-----A-E---------GG------IG-LDQG
ref|WP_114994692.1|   --------------------------------M--ST--S---RPVLRRCVACRELFDRSILWRVIRD-----H-----------RDG-----VL-LD--
gb|MEB3351632.1|      --------------------------------M-S---NH---RPVLRRCVACRTLADRLDLWRLVRL-----A-D---------GTG-----VV-LD--
ref|WP_186538343.1|   --------------------------------M--NI--S---RPVLRRCVACRELFDRSILWRVIRD-----H-----------RDG-----VL-LD--
ref|WP_038654169.1|   ---------MS-------QN----------------PI--------LRRCVACKKVLDRKYFLKVTRD-----F-Q---------NG------VV-FSGG
ref|WP_011620279.1|   ---------MN--V----SR----------------PV--------LRRCVACRQLLDRSMLLRVIRD-----H-Q---------EG------VL-LDQG
ref|WP_186589231.1|   ---------MN--V----SR----------------PV--------LRRCVACRELLDRSMLLRVIRD-----H-Q---------EG------VL-LDQG
ref|WP_074162604.1|   --------------------------------M--NI--S---RPVLRRCVACRELFDRSILWRVIRD-----H-----------RDG-----VL-LD--
gb|MAD68593.1|        --------------------------------M--NT--S---RPVLRRCVACRELFDRSLLWRVIRD-----H-----------QDG-----VL-LD--
ref|WP_110861757.1|   ---------MS-------QT----------------PI--------LRRCVACKKVLDRKYFLKVTRD-----F-Q---------NG------VV-FSDG
ref|WP_322782688.1|   ----------------------------------------------MRRCVSCRERQDRRRLWRIIRQ-----F-----------GDG-----VV-LEGT
ref|WP_197162564.1|   --------------------------------M--SGG------VVLRRCVSCRERQDRRRLWRIIRQ-----F-----------GDG-----VV-LEGT
gb|MAF41027.1|        --------------------------------M--NA--S---RPVLRRCVACRELFDRSILWRVIRD-----H-----------RDG-----VL-LD--
ref|WP_186523834.1|   ---------MN--D----SR----------------PV--------LRRCVACRELLDRSTLLRVIRD-----H-Q---------EG------VL-LDQG
gb|PZO47380.1|        -----------------------MAE------K------------NHRRCISCRQTAHRNTLWRIVRT-----F-P---------DHQ-----IQ-LD--
gb|MEB3350462.1|      ---------MN--A----PR----------------AV--------LRRCVACRALRDRRDLWRVIRG-----A-N---------GA------LA-LDVG
ref|WP_254968188.1|   -----------------------------MG-G--KP--------VLRRCVACRRLSDRSLLWRVVRL-------A---------GGG-----VQ-LD--
ref|WP_011295369.1|   ---------MS-------QN----------------PI--------LRRCIACKKVLDRKYFLKVTRD-----F-Q---------NG------VV-FSGG
ref|WP_029626136.1|   -----------------------------MK-Q--QP--------VLRRCVTCRALLDRQHLLRVIRL-------A---------EGG-----MA-LD--
gb|MAK15572.1|        ---------MN--V----SR----------------PV--------LRRCVACRQLLDRSKLLRVIRD-----H-Q---------EG------VL-LDQG
ref|WP_186490350.1|   --------------------------------M--NT--S---RPVLRRCVACRELFDRSLLWRVIRD-----H-----------RDG-----VL-LD--
gb|MEB3169508.1|      --------MVR--S----MR----------------LV--------QRRCAACRRLADRRLLWRIIRP-----A-G---------GGP-----LL-LDAG
ref|WP_255141823.1|   ----------------------------------------------MRRCVSCRERQDRRLLWRIVRQ-----Y-----------GDG-----VV-LEGA
ref|WP_036905622.1|   ---------MS-------QT----------------PI--------LRRCVACKKVLDRKYFLKVTRD-----F-Q---------NG------VV-FSGG
gb|EAQ76225.1|        ----------------------------------------------MRRCVSCRERQDRRLLWRIIRQ-----Y-----------GDG-----VV-LEGA
ref|WP_011824427.1|   ---------MS-------QP----------------PI--------LRRCVACKKVLDRKYFLKVTRD-----F-Q---------NG------VV-FSGG
gb|MDP7995305.1|      ---------MN--V----SR----------------PV--------LRRCVACRELLDRSMLLRVIRD-----H-Q---------EG------VL-IDQG
gb|MCX5957055.1|      ----------M--P----PR----------------PV--------LRRCVSCRMLVDRLQLWRVIRL-----A-D---------GG------LG-LDGG
ref|WP_037979888.1|   --------------------------------M--SGG------VVMRRCVSCRERQDRRLLWRIIRQ-----Y-----------GDG-----VV-LEGA
ref|WP_106220260.1|   --------------------------------M-TSPAGR----PVLRRCVSCRRLCDRTQLWRVVRQ-------A---------DGT-----VS-LD--
ref|WP_006854377.1|   ---------MN--V----SR----------------PV--------LRRCVACRQLLDRSLLLRVIRD-----H-Q---------DG------VL-LDQG
ref|WP_228007088.1|   ----------------------------------------------MRRCVACRVLADRRGLWRVVRL-------A---------SGG-----LQ-LD--
ref|WP_271488669.1|   ---------MN--V----SR----------------PV--------LRRCVACRELLDRSMLLRVIRD-----H-Q---------EG------VL-LDQG
ref|WP_186516396.1|   ---------MN--I----SR----------------PV--------LRRCVACRELLDRSMLLRVIRD-----H-Q---------EG------VL-LDQG
ref|WP_322775930.1|   ----------------------------------------------MRRCVSCRERQDRRLLWRIIRQ-----Y-----------GDG-----VV-LEGA
ref|WP_257473614.1|   ----------------------------------------------MRRCVACKKVLDRKYFLKVTRD-----F-Q---------NG------VV-FSDG
gb|QNI69741.1|        -----------------------------------------M---VLRRCVACRVLADRRGLWRVVRL-------A---------SGG-----LQ-LD--
gb|MDA7433454.1|      ---------MN--A----SR----------------PV--------LRRCVACRQLLDRSMLLRVIRD-----H-Q---------EG------VL-FDQG
ref|WP_269623649.1|   ---------MS-------ES----------------PI--------LRRCVACKKVLDRKCLLKVTKD-----F-Q---------NG------VV-LSGG
gb|MAR07732.1|        ---------MT-------QR----------------AI--------LRRCVACRQLLDRSLLWRVVRD-----H-K---------DG------VL-LDKG
ref|WP_254954616.1|   --------------------------------M-TSPARR----PVLRRCVSCRRLCDRSQLWRVVRQ-------A---------DGT-----VS-LD--
ref|WP_286160945.1|   ---------------------------------------------MLRRCVSCRRLCDRTQLWRVVRQ-------A---------DGT-----VS-LD--
ref|WP_269611454.1|   ---------MS-------QS----------------PI--------LRRCVACKKVLDRKFLLKVTKD-----F-Q---------NG------VV-LSGG
gb|MEB3173427.1|      ---------------------------------------------MLRRCVSCRQLVDRDQLWRVIRL-----A-G----------GG-----LA-LD--
ref|WP_225875770.1|   --------------------------------M-SPSSAPPV---VLRRCVACRVLADRRGLWRVVRL-------A---------SGG-----LQ-LD--
gb|MBD2718795.1|      --------------------------------M-TAPTGR----PVLRRCVSCRRLCDRTQLWRVVRQ-------A---------DGT-----VS-LD--
ref|WP_254992200.1|   --------------------------------M-TSNARR----PVLRRCVSCRRLCDRSQLWRVVRQ-------A---------DGS-----VS-LD--
```

```
gb|TVS05339.1|        --------------------------------M-GER-------PVMRRCVACRQLCDRRQLWRVIHL-----A-G----------GG-----VA-LD--
gb|QNI50382.1|        --------------------------------------------------MACRQLLDRRSLWRVIRD-----H-Q---------DG------VR-VDGG
gb|QBE69991.1|        -------------------------------------------------------MLDRRQLWRVIRD-----H-Q---------DG------VL-LDVG
gb|MBM5821121.1|      --------------------------------M-TPPTGR----PVLRRCVSCRRLCDRTQLWRVVRQ-------A---------DGT-----VS-LD--
gb|EAQ68383.1|        -------------------------------------------------------MLDRRQLWRVIRD-----H-R---------DG------VL-LDVG
ref|WP_254963331.1|   --------------------------------M-TSPARR----PVLRRCVSCRRLCDRSQLWRVVRQ-------A---------DGT-----VS-LD--
ref|WP_186544284.1|   ---------MN--D----SR----------------PV--------LRRCVACRQLLDRSLLLRVIRD-----H-Q---------DG------VL-FDQG
gb|MEB3171165.1|      --------------------------------M--SVQGG---RVVERRCVACRRLADRRELWRVIRL-----A-----------EGG-----LA-LD--
gb|MBW4530560.1|      --------------------------------M-TPPARR----PVLRRCVSCRRLCDRNQLWRVVRQ-------A---------DGS-----VS-LD--
ref|WP_254957545.1|   --------------------------------M-TSNARR----PVLRRCVSCRRLCDRSQLWRVVRQ-------A---------DGS-----VS-LD--
gb|EAU74800.1|        -------------------------------------------------------MLDRQQLWRVIRD-----H-Q---------DG------VL-LD--
ref|WP_015110535.1|   --------------------------------M-TSPAGR----PVLRRCVSCRRLCDRTQLWRVVRQ-------S---------DGT-----VS-LD--
gb|QNJ13310.1|        --------------------------------------------------MACRQLLDRRSLWKVIRD-----H-Q---------DG------VR-VDGG
ref|WP_254944301.1|   --------------------------------M-TAPARR----PVLRRCVSCRRLCDRSQLWRVVRR-------A---------DGS-----VS-LD--
gb|MBD2549645.1|      ----------------------------------------------MRRCVSCRRLCDRTQLWRVVRQ-------A---------DGT-----VV-LD--
gb|MCT0206289.1|      ----------------------------------------------MRRCVSCRQLCDRTQLWRVVRQ-------A---------DGS-----VS-LD--
ref|WP_159820065.1|   --------------------------------M-TSPAGR----PVLRRCVSCRRLCDRTLLWRVVRQ-------A---------DGT-----VS-LD--
ref|WP_286194360.1|   ----------------------------------------------MRRCVSCRRLCDRSQLWRVVRQ-------A---------DGS-----VS-LD--
ref|WP_254980322.1|   -----------------------------MT-E--RP--------VMRRCVACRQLLDRSQLWRVIRL-----A-G---------GG------LG-LD--
ref|WP_158467192.1|   ---------MS-------QN----------------PI--------LRRCVACKKVLDRKCLLKVTRD-----F-Q---------NG------VV-LCGG
gb|QVL53487.1|        -----------------------MAGG--------RP--------VLRRCVACRALRDRRELWRVVRL-------A---------GGG-----VV-LD--
gb|KAF0652084.1|      ---------------------------------------------MLRRCVSCRRLCDRTLLWRVVRQ-------A---------DGT-----VS-LD--
ref|WP_255092431.1|   --------------------------------M--SGG------VVLRRCVSCRERRDRSLLWRIIRQ-----Y-----------GDG-----VV-LEGA
gb|MEB3361199.1|      --------MVH--S----PR----------------LV--------QRRCAACRRLADRRLLWRIIRP-----A-G---------GGA-----LL-LDMG
ref|WP_198949670.1|   --------------------------------M--SGE------VVLRRCVSCRERRDRSLLWRIIRQ-----Y-----------GDG-----VV-LEGP
ref|WP_323356953.1|   --------------------------------M-TSPARR----PVLRRCVSCRRLCDRSLLWRVVRQ-------A---------DGS-----VS-LD--
gb|MEB3266211.1|      ---------MS--Q----PR----------------PV--------LRRCVACRALRDRRELWRVIRG-----T-G---------GE------VW-LDEG
ref|WP_255103146.1|   --------------------------------M--SGG------VVLRRCVSCRERRDRSLLWRIIRQ-----Y-----------GDG-----VV-LAGA
ref|WP_255110457.1|   --------------------------------M--SGG------VVLRRCVSCRERRDRRLLWRIIRQ-----S-----------SDG-----VV-LDGA
ref|WP_257473432.1|   ----------------------------------------------MRRCVACKKVLDRKCLLKVTRD-----F-Q---------NG------VV-LCGG
gb|MBM5793042.1|      --------------------------------------------------MACRELLDRRQLWRVIRL-----A-----------DGG-----LA-LD--
gb|MCX5932016.1|      --------------------------------M--SGG------VVLRRCVSCRERRDRRLLWRIIRQ-----S-----------SDG-----VV-LDGA
gb|MEB3351395.1|      ----------------------------------MRPQAR----PVLRCCVACRQTKDRQELWRVVRL-----A-G---------GGG-----LT-LD--
gb|MBE9154035.1|      --------------------------------------------------MACRVLADRRGLWRVVRL-------A---------SGG-----LQ-LD--
gb|MBM5825515.1|      -----------------------------MG-G--RP--------VLRRCVACRERLDRQHLWRVIRL-----A-E----------GG-----IA-LD--
gb|MCU0530038.1|      -----------------------------MA-G--KP--------VLRRCVACRQLSDRTRLWRVVRL-------A---------GGG-----VQ-LD--
gb|MEB3156001.1|      ----------------------------------MPSRGR----PVLRRCVACRQLKDRADLWRLVRL-------A---------SGG-----LA-LD--
gb|MBM5816260.1|      ----------------------------------MPSRGR----PVLRRCVACRQLKDRADLWRLVRL-------A---------SGG-----LA-LD--
gb|MCF8131934.1|      --------------------------------M--TWNEP---RPVLRRCVSCRELHDRRHLWRLIRQ-----A-G---------DGS-----VT-LD--
gb|MDM7936623.1|      --------------------------------M--SGG------VVLRRCVSCRERRDRQLLWRIIRQ-----S-----------SDG-----VV-LDGA
gb|MEB3260791.1|      --------------------------------M-PSRGR-----PVLRRCVACRQLKDRVDLWRVVRL-----A-G----------GG-----LA-LD--
gb|MEB3353798.1|      -----------------------------MG-G--RP--------VLRRCVACRELCDRSRLWRVIRL-----A-G----------GG-----IG-LE--
gb|MEB3334793.1|      ----------------------------------MSERGR----PVLRRCVACRQLKDRRDLWRLVRL-------A---------EGG-----LA-LD--
tpg|HYP03616.1|       --------------------------------M----TAR---RPVLRRCVACRRLVDREALWRVIR------L-G---------GGG-----IG-LD--
ref|WP_221629786.1|   ----------------------------------------------MRRCVSCRELRDRRLLCRIIRQ-----N-----------RDG-----VV-LEVA
gb|MBC1260749.1|      --------------------------------M--SER------IVLRRCVSCRELRDRRLLCRIIRQ-----N-----------RDG-----VV-LEVA
gb|MEB3176206.1|      --MS-------------------SKP-------------------TLRRCVSCRQLRDRRLLLRVIRL-----P-E---------GQG-----LS-LD--
ref|WP_259738767.1|   --------------------------------M--SER------IVLRRCVSCRELRDRRLLCRIIRQ-----N-----------RDG-----VV-LEVA
gb|MEB3195012.1|      ----------------------------------MSSQGK----PVMRRCVACRQLRNRLELWRVVRL-----P-A---------GGG-----LA-LD--
gb|MEB3257996.1|      ----------------------------------MPPQGR----PVLRRCVACRQLRDRQELWRVVRL-------A---------EGG-----LA-MD--
gb|MBO6974532.1|      ------------------------------MIQ-KTP--------VMRICISCRKIFDRKYLFKITKDYK---------------KGI-----MF-QK--
gb|MEB3257676.1|      ----------------------------------MPPQGK----PVLRRCVACRQLRNRLELWRVIRL-----P-A---------KGG-----LA-LD--
gb|MEB3165236.1|      --------MSA-RS----PR----------------PV--------LRRCCACRTLADRSGLLRVIRL-----A-D---------GA---G--LA-LDAG
```

```
gb|MEB3349663.1|     -----------------------M-GG--------RP--------VLRRCVACRVLVDRAELWRIVRL-------A---------AGG-----IA-LD--
ref|WP_209040626.1|  ------------------------------MIQ-QTP--------VMRICISCRKTYDRKDLFKITKDHK---------------QGI-----MI-QK--
gb|MCH9714882.1|     -----------------------MAGG---------P--------VLRRCVACRTLCDRQLLWRIIRQ-------A---------QGG-----IG-LD--
ref|WP_245156268.1|  ----------------------------------------------MRICISCRKTYDRKDLFKITKDHK---------------QGI-----MI-QK--
gb|MDX1977370.1|     --------------------------------M-SSRLKTK----LERRCIACGSLLTKSQLWRLVRM-------P---------SGK-----VE-VDLG
gb|QNI77496.1|       ------------------------------------------------------------MLLRVIRD-----H-Q---------EG------VL-LDQG
gb|EAU72346.1|       ----------------------------------------------------------------MIRD-----H-K---------DG------VL-LDLG
gb|MBL6801298.1|     ----------------------------------------------------------------MVRD-----Y-R---------DG------VL-LDDG
ref|WP_082303561.1|  ------------------------------MTP-QTP--------VMRICISCRKTYDRKHLIKITKDHK---------------LGI-----LF-QK--
gb|MCX5959459.1|     ------------------------------------------------------MLVDRLQLWRVIRL-----A-D---------GG------LG-LDGG
gb|QNI89774.1|       --------------------------------------------------------------MRVIRD-----H-Q---------DG------VL-FDQG
gb|QNJ32650.1|       ----------------------------------------------------------------MIRD-----H-Q---------EG------VL-LDQG
tpg|HAS93740.1|      ------------------------------------M--------VDRRCVSCWQVKNRDELIKITAK-----N-D---------DGK-----VI-VN--
gb|MBE7709406.1|     -------------------------------MV--LV--------VERKCVGCGIIKDRENLIKITAQ-----N-S---------YCD-----II-VN--
tpg|HBH17627.1|      -----------------MKDRN-KKNMKKRKLN-KKG--------IIRKCIGCGELKERSDLIKITKTYD---------------TYE-----VV-VAPS
gb|MEB3319688.1|     --------------------------------------MAEGGRPVLRRCVACRSLEDRRRLWRIIRLA----------------GGG-----IA-LD--

                             110       120       130       140       150       160       170       180       190       200
                     ....|....|....|....|....|....|....|....|....|....|....|....|....|....|....|....|....|....|....|....|
ref|WP_010871325.1|  ------------HG----A----GRSAYLC-PTHDCLRKARH-KNLLGRAL-------RA-P-VPSHLFEQL-EAR----L-LTTP--------------
ref|WP_190597250.1|  ------------HG----M----GRSAYLC-PTHDCLRKARH-KNLLGRAL-------RA-P-VPSHLFEQL-EAR----L-LATP--------------
ref|WP_028947476.1|  ------------HG----M----GRSAYLC-PTHDCLRKARQ-KNTLGRAL-------RA-P-VPDHLFEQL-EAR----L-LATP--------------
gb|MEB3228520.1|     ------------------S----GRSAYVC-PTASCLAKARH-KNLLGRSL-------RS-P-VPEALFTQL-ESR----L-PPPL--------------
gb|MEB3311284.1|     ------------QG----I----GRSAYLC-PQPNCLQAARQ-KNRLGRAL-------KA-P-VPDTLFQLL-EER----L-AHL--------SPP----
gb|MEB3122126.1|     ------------EG----M----GRSAYLC-PQSSCLQAARH-KNRLGRAI-------KA-T-VPEQIFQDL-EER----L-ATTKS---QESV------
ref|WP_075904775.1|  ------------QG----M----GRSAYLC-PQLSCLAAAQK-KNRLGRSL-------RA-S-VPKNLYETL-WQR----L-ATMPS-------------
gb|NEQ20425.1|       ------------------M----GRSAYLC-PQSSCLVAAQK-KNRLGRSL-------RA-S-VPDHLYKTL-EQR----L-A-------TVANT-----
gb|UXE61785.1|       ------------QG----M----GRSAYLC-PQASCLQAARQ-KNRLGRAL-------KA-P-VPDAVYEQL-LAR------LVHS--------------
ref|WP_293043112.1|  ------------QG----M----GRSAYLC-PQLSCLAAAQK-KNRLGRSL-------RA-S-VPKNLYETL-WQR----L-ATMPS-------------
gb|NEO94735.1|       ------------QG----M----GRSAYLC-PQLSCLAAAQK-KNRLGRSL-------RA-S-VPKNLYETL-WQR----L-ATMPS-------------
ref|WP_293076746.1|  ------------QG----M----GRSAYLC-PQVSCLAAAQK-KNRLGRSL-------RA-S-VPKNLYETL-WQR----L-ATMPS-------------
ref|WP_071107114.1|  ------------QG----M----GRSAYLC-PQLSCLAAAQK-KNRLGRSL-------RA-S-VPKNLYETL-WQR----L-ATMPS-------------
ref|WP_044492698.1|  ------------QG----M----GRSAYLC-PQLSCLAAAQK-KNRLGRSL-------RA-S-VPKNLYETL-WQR----L-ATMPT-------------
ref|WP_293017770.1|  ------------QG----M----GRSAYLC-PQLSCLAAAQK-KNRLGRSL-------RA-S-VPKNLYETL-WQR----L-ATMPS-------------
ref|WP_293114070.1|  ------------QG----M----GRSAYLC-PQLSCLAAAQK-KNRLGRSL-------RA-S-VPKNLYETL-WQR----L-ATMPS-------------
gb|PZV27550.1|       ------------EG----M----GRSAYLC-PQSSCLQAARH-KNRLGRAI-------KA-T-VPEQIFQDL-EER----L-ATTKS---QESV------
ref|WP_070395889.1|  ------------QG----M----GRSAYLC-PQLSCLAAAQK-KNRLGRSL-------RA-S-VPKNLYDTL-WQR----L-ATMPS-------------
gb|NEQ64070.1|       ------------QG----M----GRSAYLC-PNESCLAIAQK-KNRLGRSL-------RA-S-VPDHLYETL-WQR----L-AATAD-----------SP
gb|MEB3190993.1|     ------------QG----M----GRSAYLC-PQASCLQAARQ-KNRLGRAL-------KA-P-VPDAVYEQL-LAR------LVHS--------------
gb|MBW4545396.1|     ------------------M----GRSAYLC-PQTSCLAAAQK-KNRLGRSL-------RA-S-VPDNLYKTL-EQR----L-A-------AVANT-----
gb|MBE9166779.1|     --------Q-LDSG----M----GRSVYIC-PQADCLKKARH-KNRLGRAL-------KA-N-VPEQIYQQL-QER----L-VT---------------S
gb|NES99769.1|       ------------QG----M----GRSAYIC-PTSSCLQAAQK-KNRLGRSL-------RA-T-VPETLYQTL-WQR----L-DQTNTQKKQI--------
gb|NER21363.1|       ------------QG----M----GRSAYLC-PNESCLAIAQK-KNRLGRSL-------RA-S-VPDHLYDTL-WQR----L-AATAD-----------SP
ref|WP_096621468.1|  ------------EG----M----GRSAYLC-PQADCLQIARK-KDRIGRSL-------RT-P-VPEAIYQTL-SER----L-TNTASSTSI---------
ref|WP_192221722.1|  ------------------M----GRSAYLC-PQTSCLAAAQK-KNRLGRSL-------RA-S-IPDNLYKTL-EQR----L-A-------AVANT-----
gb|NES19959.1|       ------------KG----M----GRSAYLC-PNESCLATAQK-KNRLGRSL-------RA-S-VPEHLYETL-WQR----L-AAVAD-----------TP
ref|WP_193875631.1|  ------------QG----A----GRAAYLC-PTADCLHLAQK-KNRLGRSL-------KA-A-VPNTIYEEL-GQR----L-SSTNT-------------
gb|NEP56036.1|       ------------QG----M----GRSAYLC-PNESCLAIAQK-KNRLGRSL-------RA-S-VPEHLYETL-RQR----L-AAIAD-----------TP
gb|NET56888.1|       ------------QG----M----GRSAYLC-PNESCLTTAQK-KNRLGRSL-------RA-S-VPECLYETL-WQR----L-AAMVN-----------TP
ref|WP_272066750.1|  ------------KG----M----GRSAYLC-PNANCLKAAQK-KNRLGRAL-------RA-P-VHESVYEQL-WQR----L-DTMPT-------------
ref|WP_008308507.1|  ------------TG----M----GRSAYLC-PTADCLRIAQK-KNRLGKAL-------RA-S-VPDYIYDQL-WAR----L-DS-------SQSKSAN--
gb|PSR15992.1|       ------------TG----I----GRSAYLC-PQGDCLRQAQK-KGRLGRSL-------KA-N-VPEELYQTL-WQR----L-ETAAA---KAPGA-----
gb|PSN10559.1|       ------------TG----M----GRSAYLC-PQADCLRQAQK-KGRLGRSL-------KA-N-VPEELYQTL-WQR----L-ETAAA---KAP-------
```

```
gb|MBD0344432.1|        ------------QG----M----GRSAYLC-PQASCLATAQK-KNRLGRSL-------HA-S-VPEQLYETL-WQR----L-ATLPTAENQQGEAKS---
gb|NEP16756.1|          ------------QG----A----GRSAYLC-PNATCLQMAQK-KDRLGRAL-------KA-Q-VPPDIYQTL-QQR----L-S-----------------
ref|WP_080808818.1|     ------------EG----M----GRSAYLC-PQVDCLRTAQK-KNRLGRAL-------KA-S-VPPSIYEEL-WQR----L-ESSSV-------------
ref|WP_015954383.1|     ------------YG----M----GRSAYLC-PCASCLKEARQ-KNRLGRAL-------RV-P-IPDYIYQAL-SER----L-TPTK--------------
ref|WP_290221549.1|     ------------QG----M----GRSAYLC-PEVQCLQAAQK-KNRLGRSL-------KA-S-IPAELYQTL-EQR----L-AKAE--------------
gb|MBJ7899262.1|        ------------------M----GRSCYLC-PTAACLQGARQ-KNRLARAL-------KA-P-APESVYQAL-AAR----L-LPQSD---SV--------
ref|WP_094348539.1|     ------------QG----M----GRSAYIC-PETSCLQAAQK-KNRLGRSL-------HA-S-VPEALYQSL-SQR----L-ARSN--------------
gb|TVQ19707.1|          ------------QG----M----GRSAYLC-PSSDCLRAAQR-KDRLG-------RSLKA-T-VPDTIYQVL-WQR----L-SHDTATQSFSTDRSPSSR
ref|WP_300417529.1|     ------------QG----M----GRSAYIC-PETSCLQAAQK-KNRLGRSL-------HA-S-VPETLYQSL-SQR----L-ARSN--------------
ref|WP_110985927.1|     ------------HG----M----GRSAYLC-AQADCLRLAQK-KNRLGRAL-------RA-S-VPDSIYQTL-QER----L-TEGDR-------------
ref|WP_114081474.1|     ------------QG----M----GRSAYIC-PQTSCLQAAQK-KNRLGRSL-------HA-S-VPETLYQSL-SQR----L-ASSN--------------
gb|MCA1994153.1|        ------------QG----M----GRSAYLC-PQASCLAAAQK-KNRLGRSL-------RA-S-VPEQVYETL-SQR----L-SA--M-------------
ref|WP_190490988.1|     ------------QG----M----GRSAYLC-PQANCLGAARK-KNRLGNTL-------KA-P-VPEELYQSL-WQR----L-AEKQDGGCESGGETFRLA
ref|WP_272127684.1|     ------------QG----M----GRSAYLC-PQADCLKLAQK-KNRIARSL-------KA-P-VPDTLYQTL-WQR----L-TDSP-----PPNSQTINT
ref|WP_015183318.1|     ------------QG----M----GRSAYLC-PQASCLAAAQK-KNRLGRSL-------RT-S-VPEKLYEAL-WQR----L-APKP----ATPTQPREGE
ref|WP_190712634.1|     ------------WG----M----GRSAYLC-ATESCLKAARH-KNRLGRAL-------KT-A-IPETIYQSL-ELR----L-GAG----ALSAPPKE---
gb|NJM61055.1|          ------------SG----M----GRSAYLC-PEAECLKAAQK-KNRLGRAL-------KA-P-VPEELYQTL-RQR----LAASESV---SQGREPDNNS
gb|MDJ0704997.1|        ------------EG----M----GRSAYLC-PQESCLRIGQK-KNRLGRAL-------KA-S-IPTELYDVL-WNR----L-KTADL---VESDSSLSG-
ref|WP_292849343.1|     ------------QG----M----GRSAYIC-PETSCLQAAQK-KNRLGRSL-------HA-S-VPETLYQSL-SQR----L-ARTN--------------
ref|WP_169155198.1|     ------------EG----M----GRSAYIC-PQHNCLLAAQK-KNRLGRAL-------RA-S-VPEALYHTL-WQR----L-H-----------------
tpg|HSM83622.1|         ------------------L----GRSAYLC-PQADCLRQAQK-KSRLGRAL-------KA-N-VPEELYQQL-WQR----L-GDSAP---AP-GLDRPAR
gb|MEB3274626.1|        ------------HG----M----GRSAYLC-PNLDCLQAAQK-KNRLGRAL-------RA-P-IPADLYPKL-SDR----L-S-----------------
ref|WP_324983291.1|     ------------QG----M----GRSAYIC-PQTSCLQAAQK-KNRLGRSL-------HA-S-VPETLYQSL-LQR----L-TRTN--------------
ref|WP_292866613.1|     ------------QG----M----GRSAYIC-PETSCLQAAQK-KNRLGRSL-------HA-S-VPEALYQSL-SQR----L-ARTN--------------
ref|WP_022606905.1|     --------Q-LDEG----I----GRSAYIC-PQAACLRISRR-KNRLGRAL-------KT-A-VPETIYEQL-SER----L-VA---------------R
ref|WP_292789808.1|     ------------QG----M----GRSAYIC-PETSCLQAAQK-KNRLGRSL-------QA-S-VPETLYQSL-SQR----L-VRSN--------------
ref|WP_096570286.1|     ------------QG----M----GRSAYIC-PQADCLQAAQK-KNRLGRSL-------HG-T-VPETLYQTL-WQR----L-SERNPPNQI---------
ref|WP_012595979.1|     ------------HG----M----GRSAYLC-PEKSCLDNATK-KNRLGRTL-------KA-S-VPPTIYQSL-WER----L-EELPP-------------
ref|WP_324629747.1|     ------------KG----M----GRSAYLC-PTASCLKAAQK-KNRLGRSL-------KA-P-VPPELYETL-WQR----L-SLMST---PDNQSNAIN-
ref|WP_190731269.1|     ------------QG----M----GRSAYIC-PQTSCLQAAQK-KNRLGRSL-------HA-S-VPETLYQSL-SQR----L-ARTN--------------
tpg|HAA30330.1|         ------------KG----M----GRSAYLC-PQASCLTAAQK-KNRLGRSL-------RT-S-IPEHVYETL-WQR----L-SVVTV-------------
ref|WP_298906330.1|     ------------QG----M----GRSAYIC-PETSCLQAAQK-KNRLGRSL-------HA-S-VPETLYQSL-SQR----L-ARTN--------------
dbj|BAZ68765.1|         ------------EG----M----GRSAYIC-PNMSCLSAAQK-KNRLGRSL-------HA-S-VPETLYKTL-LQR----L-AQSN--------------
gb|MBW4433734.1|        ------------EG----M----GRSAYIC-PNMSCLSAAQK-KNRLGRSL-------HA-S-VPETLYKTL-LQR----L-AQSN--------------
ref|WP_181927846.1|     ------------QG----M----GRSAYIC-PETSCLQAAQK-KNRLGRSL-------HA-S-VPETLYQSL-SQR----L-ARSN--------------
ref|WP_012412015.1|     ------------RG----M----GRSAYIC-PQTSCLQAAQK-KNRLGRSL-------HA-S-VPETLYQSL-SQR----L-ASSN--------------
ref|WP_322677349.1|     ------------QG----M----GRSAYIC-PEMSCLQAAQK-KNRLGRSL-------HA-S-VPETLYQSL-SQR----L-ARSN--------------
ref|WP_015139046.1|     ------------EG----M----GRSAYIC-PNHSCLQAAQK-KNRLGRSL-------RA-S-VPETLYQTL-WQH----L-AQKN--------------
ref|WP_104905836.1|     ------------QG----M----GRSAYIC-PETSCLQAAQK-KNRLGRSL-------HA-S-VPETLYQSL-SQR----L-ARSN--------------
ref|WP_206814010.1|     ------------QG----M----GRSAYLC-PTQNCLMNARQ-KNRLQRAL-------KA-P-IPNDLYQVL-QER----L-ERAT-----T--------
gb|MDJ0601942.1|        ------------QG----M----GRSAYLC-PQESCLTLASK-KNRLGRRL-------KA-P-IPDSIYQQL-WQR----L-SQFMA---DG--------
ref|WP_292717392.1|     ------------QG----M----GRSAYIC-PQTSCLQAAQK-KNRLGRSL-------HA-S-VPETLYQSL-SQR----L-TPSN--------------
ref|WP_179066568.1|     ------------QG----M----GRSAYIC-PETSCLQAAQK-KNRLGRSL-------HA-S-VPETLYQSL-SQR----L-ARIN--------------
ref|WP_322722179.1|     ------------QG----M----GRSAYIC-PETSCLQAAQK-KNRLGRSL-------HA-S-VPETLYQSL-SQR----L-APNN--------------
ref|WP_185581622.1|     ------------QG----M----GRSAYIC-PQTSCLQAAQK-KNRLGRSL-------HA-S-VPETLYQSL-SQR----L-PSSN--------------
ref|WP_292822895.1|     ------------QG----M----GRSAYIC-PETSCLQAAQK-KNRLGRSL-------HA-S-VPETLYQSL-SQR----L-APSN--------------
ref|WP_196512348.1|     ------------QG----M----GRSAYIC-PKTSCLQAAQK-KNRLGRSL-------HA-S-VPETLYQSL-SQR----L-ARSN--------------
gb|OKH54711.1|          ------------EG----M----GRSAYIC-PSLDCLGAAQK-KNRLGRAL-------RA-A-VSDEIYQSL-WQR----L-TVVQPTIVNIG-------
ref|WP_292785937.1|     ------------QG----M----GRSAYIC-PETSCLQAAQK-KNRLGRSL-------HA-S-VPEALYQSL-SQR----L-ARSN--------------
gb|TAE56869.1|          ------------QG----I----GRSAYIC-PKSSCLQAAQK-KDRLGRSL-------RE-S-VPETLYQTL-WQR----L-NQTNTQEKQI--------
gb|PZV08645.1|          ------------EG----M----GRSAYLC-PQTECLKAAQK-KNRLGKAV-------KA-V-ISDSLYEEL-EKR----L-YKQEI-------------
ref|WP_292751094.1|     ------------QG----M----GRSAYIC-PQTSCLQAAQK-KNRLGRSL-------HA-S-VPETLYQSL-SQR----L-APSN--------------
ref|WP_073597769.1|     ------------RG----M----GRSAYIC-PQASCLTAACQ-KNRLGRAL-------KA-P-IPDRIYESL-WER----L-ATV---------------
```

```
ref|WP_073595002.1|   ------------WG----M----GRSAYLC-PQATCLETAQK-KNRLGKTL-------RA-T-VPPELYQAL-WQR----L-SNQPEIANPTSTKQPTSE
gb|MCG8363753.1|      ------------EG----I----GRSVYLC-PQADCLRTGQK-KNRLGRSL-------KA-P-IPAELYDVL-WDR----L-KTAEL---AESESSLSG-
gb|PSO56092.1|        ------------SG----V----GRSAYLC-PNSSCVKAARR-KNRLGRSV-------KA-R-VPEQIYQTL-EER----L-ATR-------------T-
gb|MBF2088208.1|      ------------EG----M----GRSAYLC-PSSDCLKAAQR-KDRLG-------RSLKA-T-VPDTIYQGL-WQR----L-SHDTAAQSMSTDRSPQAR
gb|NES80898.1|        ------------QG----M----GRSAYLC-PQASCLQRAQA-KNRLGRSL-------KA-S-VPQQLYQKL-WER----L-AAKCT---TT--------
gb|PSO96923.1|        ------------SG----V----GRSAYLC-PNSSCVKAARR-KNRLGRSV-------KA-R-VPEQIYQTL-EER----L-ATR-------------T-
dbj|BAZ39493.1|       ------------EG----M----GRSAYIC-PQASCLTTAQK-KNRLGRSL-------HA-T-VPEALYQAL-WQR----L-ALPQQ-------------
ref|WP_322731565.1|   ------------QG----M----GRSAYIC-PETSCLQAAQK-KNRLGRSL-------HA-S-VPETLYQSL-SQR----L-ARSN--------------
ref|WP_193978934.1|   ------------EG----M----GRSAYLC-PQATCLETAQK-KNRLGKTL-------RA-T-VPPELYQAL-WQR----L-SNQPVQANPTTPKLSASD
ref|WP_300507896.1|   ------------QG----M----GRSAYLC-PQESCLTLAGK-KNRLGRRL-------KA-P-IPDNIYQQL-WQR----L-NKVLP---EQ----ELS-
gb|TVQ45794.1|        ------------QG----M----GRSAYLC-PQETCLRAAQQ-KNRLAKAL-------KT-S-VPAEIYQQL-WQR----L-ETN---------------
gb|MBW4655594.1|      ------------EG----A----GRSVYLC-PQASCLKAAQK-KDRLGRSL-------RA-P-VPPRIYEAL-WQR----I-ATG---------------
ref|WP_204152300.1|   ------------EG----M----GRSAYLC-PSSDCLKAAQR-KDRLG-------RSLKA-T-VPDTIYQGL-WQR----L-SHDTAAQSMGTDRSPQAR
ref|WP_292734233.1|   ------------QG----M----GRSAYIC-PETSCLQAAQK-KNRLGRSL-------HA-S-VPEALYQSL-SQR----L-ARSN--------------
gb|PSN19469.1|        ------------QG----M----GRSAYLC-PTEDCLKAAHK-KNRLGRML-------KS-P-IPESIYQEL-WQR----L-GS--------------KQ
ref|WP_094331183.1|   ------------RG----M----GRSAYIC-PQTSCLQAAQK-KNRLGRSL-------HA-S-VPETLYQSL-SQR----L-AISN--------------
gb|MDW8178835.1|      ------------------M----GRSAYLC-PKRECLQYAQQ-KNRLGKAL-------KA-P-VPPEIYAAL-WAR----L-NTAGP---APDNRPP---
ref|WP_223049825.1|   ------------------M----GRSAYLC-PQASCLQAAQR-KDRLGRVL-------KV-P-VPESIYNAL-WSR----L-AQGQE---GVKFDTENHK
dbj|BDA66563.1|       ------------EG----M----GRSAYIC-PSYDCLGAAQK-KNRLGRAL-------HA-G-VSDEIYQSL-WQR----L-TVVQSTIVNIG-------
ref|WP_194027457.1|   ------------------M----GRSAYLC-PQETCLRAAQQ-KKRLSRSL-------RT-T-VPDEVYQQL-WLR----L-R-------AIENN-----
gb|NEO98984.1|        ------------EG----M----GRSAYLC-PKASCLATAQK-KNRLGRAL-------RA-P-VPLSLYKTL-WER----L-AEESV---VGTSKLAEEQ
gb|MDZ8004527.1|      ------------RG----M----GRSAYIC-PQASCLQAAQK-KNRLGRSL-------HA-S-VPETLYQSL-SQR----L-AASN--------------
gb|KAB8333374.1|      ------------EG----M----GRSAYIC-PQQNCLLAAQK-KNRLGRAL-------RA-S-VPEALYHTL-RQR----L-H-----------------
gb|MCY7392750.1|      ------------EG----M----GRSAYLC-PQLECLKAAQK-KNRLGRAL-------KA-A-ISNSLYEEL-RGR----L-CK--P-------------
gb|MBE9078818.1|      ------------QG----M----GRSAYLC-PRADCLTAAKR-KNRLARVL-------KA-Q-VPETLYQTL-EQR----L-EQSEE---ASAGATSGSA
ref|WP_322669753.1|   ------------QG----M----GRSAYIC-PETSCLQAAQK-KNRLGRSL-------HA-S-VPEALYQSL-SQR----L-ARSN--------------
ref|WP_100898624.1|   ------------QG----M----GRSAYIC-PETSCLQAAQK-KNRLGRSL-------HA-S-VPETLYQSL-LQR----L-AQSN--------------
ref|WP_229551764.1|   ------------QG----M----GRSAYIC-PQTSCLQAAQK-KNRLGRSL-------HA-S-VPETLYQSL-LHR----L-AQSN--------------
gb|MBD0304031.1|      ------------QG----M----GRSAYIC-PQTSCLQAAQK-KNRLGRSL-------HA-S-VPETLYQSL-LQR----L-AKSN--------------
ref|WP_169218276.1|   ------------EG----M----GRSAYIC-PQQSCLLAAQK-KNRLGRAL-------RA-S-VPEALYHTL-LQR----L-YQSS--------------
gb|NEO32345.1|        ------------EG----M----GRSAYLC-PKASCLATAQK-KNRLGRVL-------RA-P-VPSSLYQTL-WER----L-VEASA---VGASRLAGEP
ref|WP_190814021.1|   ------------RG----M----GRSAYLC-PQESCLVAAQK-KNRLGRAL-------KV-P-VPEALYQTL-WQR----L-ATKLEGCDRSLLKSE---
ref|WP_196527153.1|   ------------EG----M----GRSAYIC-PQTSCLQAAQK-KNRLGRSL-------HA-S-VPETLYQSL-LQR----L-AQSN--------------
ref|WP_323196799.1|   ------------QG----M----GRSAYIC-PQMSCLQAAQK-KNRLGRSL-------HA-S-VPETLYEEL-WQR----L-AQSH--------------
gb|PSO55897.1|        ------------SG----L----GRSAYLC-PNSSCVKAARR-KNRLGRSV-------KA-R-VPEQIYQTL-EER----L-ATR-------------T-
ref|WP_229482756.1|   ------------QG----M----GRSAYIC-PQTSCLQAAQK-KNRLGRSL-------HA-S-VPETLYQSL-LQR----L-AQSN--------------
ref|WP_292709822.1|   ------------RG----M----GRSAYIC-PQTSCLQAAQK-KNRLGRSL-------HA-S-VPETLYQSL-LQR----L-ASSN--------------
ref|WP_069073458.1|   ------------QG----M----GRSAYIC-PEMSCLQAAQK-KNRLGRSL-------HA-S-VPETLYQSL-WQR----L-ARSN--------------
dbj|BAU66516.1|       ------------EG----M----GRSAYLC-PEEDCLLKARH-KNRLGRSL-------RA-N-IPEEIYRSL-AQR----L-QT----------------
ref|WP_229454557.1|   ------------QG----M----GRSAYIC-PQRSCLQAAQK-KNRLGRSL-------HA-S-VPETLYQSL-LQR----L-AQSN--------------
ref|WP_322697626.1|   ------------QG----M----GRSAYIC-PETSCLQAAQK-KNRLGRSL-------HA-S-VPETLYQSL-SQR----L-AQ----------------
ref|WP_273766710.1|   ------------QG----M----GRSAYIC-PETSCLQAAQK-KNRLGRSL-------HA-S-VPETLYQSL-SQR----L-ARTN--------------
gb|MBE7381418.1|      ------------EG----A----GRSAYLC-PNAACLKAAQK-KDRIGRSL-------RT-T-IPNGLYDQL-WQR----L-M--PS---SSNAEHE---
gb|RCJ24275.1|        ------------RG----M----GRSAYIC-PQASCLQGAQK-KNRLGRSL-------HA-S-VPETLFQSL-SQR----L-ASNN--------------
ref|WP_015187990.1|   ------------QG----M----GRSAYLC-PQAGCLQTAQK-KNLLRRSL-------RI-S-VPEAIYQAL-GQR----L-LTNPSVSETQKIETTLPI
gb|NET06949.1|        ------------EG----M----GRSAYLC-PKASCLATAQK-KNRLGRVL-------RA-P-VPSSLYQTL-WER----L-AEASA---VGASRLAGET
ref|WP_224409244.1|   ------------EG----M----GRSAYLC-PGESCLSRAQS-KNRLGRSL-------RA-Y-VPQEIYQSL-WQR----L-SSVTNKEEISAVPKNNCP
ref|WP_155748928.1|   ------------EG----M----GRSAYIC-PNQSCVAAAQK-KNKLGRAL-------QA-S-VPEPLYQLL-WQR----L-SQENNQNSTDN-------
ref|WP_229547107.1|   ------------QG----M----GRSAYIC-PETSCLQAAQK-KNRLGRSL-------HA-S-VPETLYQSL-LQR----L-APSN--------------
gb|MDW8401950.1|      ------------------M----GRSAYLC-PKRECLQYAQQ-KNRLGKAL-------KA-P-VPPEIYAAL-WAR----L-NTAGP---APDDRLP---
ref|WP_107668609.1|   ------------QG----M----GRSAYLC-PNKSCLTLAGQ-KNRLGRGL-------KA-S-VPDSIYQQL-WER----L-G--EL---VEDT------
ref|WP_008232964.1|   ------------EG----M----GRSAYIC-PTSNCLSVAQK-KNRLGRSL-------NT-L-VPKALYQLL-WQR----L-IQTG--------------
gb|MEC4893587.1|      ------------EG----M----GRSAYLC-PGESCLSRAQS-KNRLGRSL-------RA-Y-VPQEIYQSL-WQR----L-SSVTNKEAISAVPKNNCP
```

```
ref|WP_109009916.1|  ------------QG----M----GRSAYIC-PQTSCLQAAQK-KNRLGRSL-------HA-S-VPETLYQSL-LQR----L-AQCN--------------
gb|MEC4816215.1|     ------------QG----M----GRSAYIC-PHQSCLSVAQK-KNKLGRAL-------HA-S-VPETLYQTL-WQR----L-SQRNNQNQTQT-------
gb|MBW4486969.1|     ------------QG----M----GRSAYLC-PEVQCLQAAQK-KNRLGRSL-------KA-F-VSAELYQTL-EQR----L-AKADE-------------
tpg|HZG41009.1|      ------------EG----M----GRSAYLC-PQADCLRQAQK-KSRLGRAL-------KV-N-VPDEVYQAL-WQR----L-EPATD---PAAPVP----
ref|WP_322643731.1|  ------------WG----M----GRSAYIC-PQTSCLQTAQK-KNRLGRSL-------HA-S-VPETLYQSL-SQR----L-ASSN--------------
ref|WP_118168122.1|  ------------QG----M----GRSAYIC-PETSCLQAAQK-KNRLGRSL-------HA-S-VPETLYQSL-LHR----L-ASNN--------------
ref|WP_194042652.1|  ------------QG----M----GRSAYIC-PETSCLQAAQK-KNRLGRSL-------HA-S-VPEALYQSL-SQR----L-ARSN--------------
ref|WP_086688430.1|  ------------EG----M----GRSAYIC-PQTNCLQAAQK-KNRLGRSL-------HA-T-VPETLYQTL-WQR----L-AKSN--------------
gb|MEB3161329.1|     ------------------S----GRSAYLC-PTVDCLGKARH-KN-------------------------------------------------------
gb|MBF2099090.1|     ------------EG----Q----GRSAYLC-PTADCLKQAQQ-KNRLGRAL-------KA-P-VSPEIYAQL-WSR----L-SQ----------------
gb|MBE8968595.1|     ------------QG----I----GRSAYIC-PETSCLQAAQK-KNRLGRSL-------HA-S-VPEALYQSL-SQR----L-ARSN--------------
ref|WP_152589415.1|  ------------QG----M----GRSAYIC-PKTSCLQAAQK-KNRLGRSL-------HA-S-VPETLYQSL-LHR----L-ASNN--------------
gb|MCT7960255.1|     ------------TG----M----GRSAYLC-PEASCLKTAQK-KNRLGRSL-------KA-A-VPSELYSNL-WQR----L-PLSAQ---SEQ-------
gb|MBW4576792.1|     ------------QG----M----GRSAYLC-PQASCLEAARK-KNRLGRSL-------KA-S-VPAEIYQTL-SQR----L-AASSR---GDPPSYRMGN
ref|WP_233748074.1|  ------------QG----S----GRSAYIC-PQEDCLKTAQK-KNRLGRSL-------RC-A-VPDSIYREL-WER----L-QRRTS-------------
gb|MDJ0730709.1|     ------------QG----M----GRSAYLC-PQKSCLTLASQ-KNRLGRRL-------KA-P-IPDSIYQQL-WQR----L-SKFIP---DS--------
gb|NEO25536.1|       ------------QG----M----GRSAYLC-PRADCLKLAQK-KNRIARSL-------KA-P-VPDSLYQTL-WQR----L-AEQP-----TPNLPP---
ref|WP_190559635.1|  ------------QG----M----GRSAYIC-PETDCLQAAQK-KNRLGRSL-------HG-T-VPETLYQTL-WQR----L-TESN--------------
ref|WP_190600959.1|  ------------EG----M----GRSAYIC-PQASCLQVAQK-KNKLGRSL-------HA-T-VPEALYQTL-WQR----L-TNSN--------------
ref|WP_289796853.1|  ------------QG----M----GRSAYIC-PETSCLQAAQK-KNRLGRSL-------HA-S-VPEALYQSL-SQR----L-APSN--------------
ref|WP_194057957.1|  ------------DG----M----GRSAYLC-PQADCLRQAQK-KRRLGRAL-------KA-N-VPEELYQTL-WQR----L-ETAVA---DPGAGTT---
ref|WP_016866716.1|  ------------EG----M----GRSAYIC-PNMSCLSAAQK-KNKLGRSL-------HA-S-VPETLYQTL-WQR----L-SQSNLPNQIEL-------
gb|MCS6814375.1|     ------------QG----M----GRSAYLC-PQASCLKLAQQ-KNRLGRAL-------KA-S-IPPELYQTL-WQR----L-SQVAQ-------------
tpg|HBB35302.1|      ------------EG----I----GRSAYLC-PQASCLTAAQK-KNRLGRSL-------QA-S-VPEELYQTL-WQR----L-AALPSLGSAIEKSAH---
gb|MDJ0715684.1|     ------------RG----M----GRSAYLC-PQVSCLQAACR-KNRLGRAL-------KA-K-IPEHIYQSL-WAR----L-ANEPR---SKIGKSPNS-
gb|NEP12133.1|       ------------EG----M----GRSAYLC-RNESCLTTAQK-KNRLGRSL-------RA-S-VPEHLYKTL-WQR----L-AAIAD-----------RP
gb|NJP17789.1|       ------------RG----V----GRSAYIC-PEASCLKAACQ-KNRLGRSL-------KA-S-VPDFIYESL-WER----L-ANA---------------
gb|KAB8319469.1|     ------------EG----M----GRSAYIC-PQQSCLVAAQK-KNRLGRSL-------RA-S-VPEALYHKL-WQR----L-SLNNNQKQT---------
ref|WP_190413342.1|  ------------RG----M----GRSAYLC-PQESCLVAAQK-KNRLGRAL-------KV-P-VPEALYQTL-WQR----L-ATKLEGCDRSLSRSE---
gb|MBR8834127.1|     ------------QG----M----GRSAYIC-PQLNCLQAAQK-KNRLGRAL-------HA-S-VPEALYQTL-WQR----L-AQSNSQNQT---------
gb|MCC5607216.1|     ------------QG----M----GRSAYIC-PETSCLQAAQK-KNRLGRSL-------HA-S-VPETLYQSL-LHR----L-AQSN--------------
ref|WP_168729992.1|  ------------QG----M----GRSAYIC-PQADCLHKAQR-KNKLGRSL-------KG-T-VPETVYETL-WQR----L-SQEN--------------
ref|WP_280654349.1|  ------------QG----M----GRSAYIC-PQTGCLQAAQK-KNRLGRSL-------QA-S-VPEALYQTL-WQR----L-AQTN--------------
gb|MBD2016732.1|     ------------EG----M----GRSAYLC-PQPSCLAVAQK-KNRLGRSL-------RT-S-VPEKLYETL-WQR----L-AALP----AIPTQSGEGE
ref|WP_099069649.1|  ------------RG----M----GRSAYIC-PQASCLQAAQK-KNRLGRSL-------HA-S-VPETLYQSL-SQR----L-ASNN--------------
ref|WP_190533136.1|  ------------RG----M----GRSAYLC-PQESCLVAAQK-KNRLGRAL-------KV-P-VPEALYQTL-WQR----L-ATKLEGCDRSLPRSE---
gb|MCU0541201.1|     ------------SG----M----GRSAYLC-PEAECLKAAHK-KNRLGRAL-------KA-P-VPDELYQTL-RQR----LGASESV---LQGREPDNNS
ref|WP_193872423.1|  ------------EG----M----GRSAYLC-PQVSCLAAAQK-KNRLGRVL-------RT-L-VPEKVYQTL-WLR----L-EATSA-------------
ref|WP_061545594.1|  ------------QG----M----GRSAYIC-PQADCLHKAQR-KNKLGRSL-------KG-T-VPETVYETL-WQR----L-SQEN--------------
ref|WP_102179423.1|  ------------EG----M----GRSAYIC-PNMSCLSAAQK-KNKLGRSL-------HA-S-VPETLYQTL-WQR----L-SQSNLPNQIEH-------
tpg|HLP90034.1|      ------------QG----M----GRSAYIC-PKMSCLQTAQK-KNRLGRSL-------RA-T-VPETLYQAL-WQR----L-PQTNKPKQIELE------
ref|WP_190956491.1|  ------------RG----M----GRSAYIC-PQASCLQAAQK-KNRLGRSL-------HA-S-VPETLYQSL-SQR----L-ASNN--------------
ref|WP_102175048.1|  ------------EG----M----GRSAYIC-PKMSCLSAAQK-KNKLGRSL-------HA-S-VPETLYQTL-WQR----L-SQSNLPNQIEL-------
gb|NJO42002.1|       ------------QG----M----GRSAYLC-PNRACLQTAQK-KNRLGRSL-------KA-S-VLEEFYQVL-WQR----L-FETDA---ESQTR-----
ref|WP_006275620.1|  ------------QG----M----GRSAYIC-PQADCLHKAQR-KNKLGRSL-------KG-T-VPETVYETL-WQR----L-GQEN--------------
gb|MBU7582629.1|     ------------EG----M----GRSAYLC-PQASCLQAANK-KNRLGRSL-------QA-S-VPETLYQTL-WQR----L-APSHP-------------
gb|MCG6133583.1|     ------------QG----M----GRSAYIC-PQMSCLQAAQK-KNRLGRSL-------HA-S-VPETLYQTL-WQR----L-AQTH--------------
ref|WP_016872536.1|  ------------EG----M----GRSAYIC-PNPSCLSAAQK-KNRLGRSL-------RA-S-VPEALYQTL-WQR----L-VKSN--------------
ref|WP_015117518.1|  ------------RG----M----GRSAYIC-PNSTCLSLAQK-KNKLGRAL-------RA-P-VPEMLYKTL-WQQ----L-TIK---------------
tpg|HBE20201.1|      ------------VG----M----GRSAYIC-PQENCLRAAQK-KNRLGRAL-------RV-S-VSEELYQML-WQR----L-AAKAE---EPINAGSDP-
gb|MBW4500045.1|     ------------QG----M----GRSAYIC-PQQSCLVAAQK-KNKLGRSL-------RA-S-VPEALYHIL-WQR----L-SQSNNQKQT---------
ref|WP_103135735.1|  ------------EG----M----GRSAYIC-PQHSCLQAAQK-KNKLGRSL-------RA-S-VPETLYQTL-WHH----L-ALKNP-------------
ref|WP_105221837.1|  ------------QG----M----GRSAYLC-PQASCLQTAQK-KNLLRRSL-------RT-S-VPEAIYQAL-WQR----L-LTNQLVSD-QKIETTAPV
```

```
ref|WP_190434246.1|    ------------QG----M----GRSAYLC-PEVSCLQAAQK-KNRLGRSL-------KA-S-VPPDLYQTL-EQR----L-AKARA-------------
ref|WP_283760063.1|    ------------GG----M----GRSAYLC-PQADCLKAAEK-KNRLA-------KSLRA-T-VPPSIYETL-WSR----L-SH----------------
gb|MDZ8167634.1|       ------------RG----M----GRSAYIC-PQVSCLQAAQK-KNRLGRSL-------HA-S-VPETLYQSL-SQR----L-ASNN--------------
ref|WP_193879029.1|    ------------QG----M----GRSAYIC-PQASCLQAANK-KNKLGRSL-------QA-T-VPETLYQTL-WQR----L-APSHH-------------
ref|WP_190591144.1|    ------------EG----M----GRSAYLC-PQATCLQTAQK-KNRLGRSL-------QA-S-VPETLYQTL-WQR----L-APSHH-------------
gb|MDY6781740.1|       ------------RG----M----GRSAYLC-PTDCCLKAAQH-KNRLGRSL-------KA-P-IPEEIYLSL-RSR----R-NL------LPPTCPTLAK
gb|MDZ8106853.1|       ------------RG----M----GRSAYIC-PQASCLQAAQK-KNRLGRSL-------HA-S-VPETLYQSL-SQR----L-GSNN--------------
ref|WP_013325296.1|    ------------QG----M----GRSAYLC-PKASCLKEARH-KNRLSRAL-------KT-P-VPESIYQLL-SER----M-THSI-----LQQPPSE--
ref|WP_171573341.1|    ------------EG----M----GRSAYLC-PKIECLKAARK-KNRLGRAL-------KA-A-VPDSLYENL-EQR----F-S-----------------
ref|WP_190938724.1|    ------------QG----M----GRSAYIC-PEMSCLQAAQK-KNRLGRSL-------HA-S-VPETLYQSL-WQR----L-SGSN--------------
ref|WP_086768880.1|    ------------EG----I----GRSAYIC-PQTNCLQAAQK-KNRLGRSL-------HA-T-VPETLYQTL-WQR----L-AKSN--------------
ref|WP_251960204.1|    ------------QG----M----GRSAYIC-PETSCLQAAQK-KNRLGRSL-------HA-S-VPETLYQSL-WQR----L-CGSN--------------
ref|WP_009545653.1|    ------------QG----M----GRSAYLC-PNASCLTLASK-KNRLGRGL-------KT-S-IPERIYQQL-WKR----L-EKSVLDT-----------
ref|WP_127085004.1|    ------------------M----GRSAYIC-PSYDCLGAAQK-KNRLGRAL-------HA-A-VSDEIYQKL-LAT----V-DSGST---YNR-------
tpg|HIK08066.1|        ------------EG----M----GRSAYIC-PQQSCLQAAQK-KNKLGRSL-------RA-S-VPETLYQTL-WQH----L-ALNNP-------------
gb|MDZ7979263.1|       ------------EG----M----GRSAYIC-PETSCLQAAQK-KNRLGRSL-------HA-T-VPETLYQTL-WQR----L-AKSN--------------
gb|MCC5635819.1|       ------------EG----M----GRSAYIC-PQASCLQAANK-KNRLGRSL-------QA-S-VPETLYQTL-WQR----L-APSHQ-------------
ref|WP_239731788.1|    ------------QG----M----GRSAYIC-PQMSCLQAAQK-KNRLGRSL-------RA-S-VPETLYQTL-WQS----L-AQSH--------------
ref|WP_193897322.1|    ------------QG----M----GRSAYIC-PQASCLQAANK-KNRLGRSL-------QA-S-VPETLYQTL-WQR----L-APSHQ-------------
ref|WP_187706878.1|    ------------QG----M----GRSAYIC-PQADCLHKAQR-KNKLGRSL-------KG-I-VPETVYETL-WQR----L-GREN--------------
ref|WP_214438763.1|    ------------QG----M----GRSAYIC-PQAGCLQAAQK-KNRLGRSL-------HA-S-VPETLYQTL-WQR----L-AQSN--------------
ref|WP_073632737.1|    ------------EG----M----GRSAYIC-PQQSCLVAAQK-KNRLGRSL-------RA-S-VPEALYHIL-WQR----L-SLSNNQKQT---------
gb|MDZ8241182.1|       ------------RG----M----GRSAYIC-PQASCLQAAQK-KNRLGRSL-------HA-S-VPETLYQSL-SQR----L-AHNN--------------
ref|WP_073548075.1|    ------------QG----M----GRSAYLC-PQASCLQTAQK-KNLLRRSL-------RI-S-VPEAIYQAL-WQR----L-STNPSVSETQKIETTLPI
gb|MBD2163199.1|       ------------EG----M----GRSAYIC-PQTSCLQAAQK-KNRLGRSL-------HA-T-VPETLYQTL-WQR----L-AKSN--------------
gb|MBO1346139.1|       ------------LG----M----GRSAYLC-PEQECLKAAQK-KNRLGKAL-------RA-R-VPQEIYQTL-WER----L-GAVSP---GAASRGSATR
gb|MBW4600386.1|       ------------EG----M----GRSAYIC-PSLDCLGAAQK-KNRLGRAL-------HA-A-VSDEIYQSL-WQR----L-TVVQPTIVNIG-------
gb|MBC6453437.1|       ------------LG----M----GRSAYLC-PEQECLKAAQK-KNRLGKAL-------RA-R-VPQEIYQTL-WER----L-GAVSP---GSA-------
ref|WP_289789294.1|    ------------EG----M----GRSAYIC-PNPNCLSAAQK-KNRLGRSL-------RA-S-VPETLYQTL-WQR----L-VQSN--------------
ref|WP_167722165.1|    ------------EG----M----GRSAYIC-PQTSCLQAAQK-KNRLGRSL-------HA-T-VPETLYQTL-WQR----L-AKSN--------------
ref|WP_095719821.1|    ------------EG----M----GRSAYIC-PQASCLTAAQK-KNRLGRSL-------HA-T-VPEALYQSL-WQR----L-TLSNSPVEINYSRRYIHP
ref|WP_096644639.1|    ------------EG----M----GRSAYIC-PQTSCLQAAQK-KNRLGRSL-------HA-T-VPETLYQTL-WQR----L-AKSNH-----------QS
gb|NJM70351.1|         ------------EG----M----GRSAYIC-PQQSCFLAAQK-KNRLGRAL-------RA-S-VPEALYHTL-LQR----L-YQSS--------------
ref|WP_082127362.1|    ------------EG----M----GRSAYIC-PQASCLTMAQK-KNRLG-------RSLRT-A-VPETLYQSL-WQR----L-THSSLNESVS--------
ref|WP_073610176.1|    ------------DG----M----GRSAYLC-PQADCLRQAQK-KSRLGRAL-------KA-N-VPEEIYQNL-WNQ----L-ESKMA-------------
gb|MDZ8028666.1|       ------------RG----M----GRSAYIC-PQGSCLQAAQK-KNRLGRSL-------HA-S-VPETLYQSL-SQR----L-AHNN--------------
gb|MDZ8019070.1|       ------------RG----M----GRSAYIC-PQGSCLQAAQK-KNRLGRSL-------HA-S-VPQTLYQSL-SQR----L-AHNN--------------
ref|WP_017742689.1|    ------------EG----M----GRSAYIC-PNQSCVAAALK-KNKLGRAL-------HA-S-VPETLYQVL-WQR----L-SQGN--------------
ref|WP_045868281.1|    ------------EG----M----GRSAYIC-PQTSCLQAAQK-KNRLGRSL-------HA-T-VPETLYQTL-WQR----L-ANSN--------------
ref|WP_208344959.1|    ------------KG----M----GRSAYIC-PQQSCLQAAQK-KNRLGRSL-------HA-P-VPETLYQTL-WQR----L-AICQS---PGSG------
gb|MBW4560223.1|       ------------QG----M----GRSAYIC-PQTSCLQAAQK-KNRLGRSL-------HA-T-VPETLYQTL-WQR----L-AQSN--------------
gb|PSB49245.1|         ------------RG----L----GRSAYLC-QASQCLKAAQK-KNRLGRSL-------KA-P-VSNELYQTL-WQR----L-ATSD--LTSDC-------
ref|WP_038087798.1|    ------------EG----M----GRSAYIC-PSQSCVATAQK-KNKLGRAL-------HA-S-VPEPLYQLL-WQR----L-SQGNQNQPEN--------
ref|WP_190427456.1|    ------------RG----M----GRSAYLC-PQESCLVAAQK-KNRLGRVL-------KV-P-VPEALYQTL-WQR----L-ATKLEGCDRSLSRSE---
ref|WP_261224160.1|    ------------QG----E----GRSAYLC-PLLSCLQAAQK-KNRLGRTL-------KA-N-VPSEIYQSL-SQR----L-SAKQEPEIQGSAS-----
gb|ARV61523.1|         ------------EG----M----GRSAYIC-PQQSCLAAAQK-KNRLGRSL-------RA-S-VPEALYHIL-QQR----L-SQSH--------------
gb|MCU0523589.1|       ------------QG----M----GRSAYLC-PSQDCLQLAQK-KDRIGRSL-------RV-A-VPTALYQTL-GQR----L-S--PS---KQSSSAE---
ref|WP_089091493.1|    ------------QG----M----GRSAYIC-PEMSCLQAAQK-KNRLGRSL-------RA-S-VPETLYQTL-WQH----L-AQSH--------------
gb|MBF2080108.1|       ------------EG----A----GRSAYVC-PTATCLKLSQR-KDRLGRSL-------KA-A-VPATLYQAL-WHR----L-DHQEN---PHEPVQDEA-
ref|WP_268609846.1|    ------------QG----M----GRSSYLC-PTIECLQTAQK-KNRLGRAL-------KT-P-VPPEIYQTL-WHR----L-SSSTSLEKKELEE-----
gb|MBC6418410.1|       --------Q-LEKG----M----GRSAYIC-RSAECLRVARQ-KNKLGRSL-------KA-T-VPEHIYQSL-SLADVLTI-LP---------------R
ref|WP_200324434.1|    ------------QG----M----GRSAYIC-PQADCLQAAQK-KNKLGRSL-------HG-T-VPETLYQTL-WQR----L-TASN----S---------
ref|WP_015202566.1|    ------------QG----M----GRSAYLC-RQTACLEAAKK-KNRLGRSL-------KA-P-VPEHIYITL-WQR----L-ATS---------------
```

```
ref|WP_194023660.1|   ------------EG----M----GRSAYLC-PRADCLRQAQK-KGRLGRAL-------KA-N-VPEEMYQTL-WQR----L-EAAPE---RTAPENSDRV
dbj|BBD59214.1|       ------------EG----M----GRSAYIC-PQATCLQAANK-KNRLGRSL-------QA-S-VPEALYQTL-WQR----L-APSHH-------------
ref|WP_096579096.1|   ------------EG----M----GRSAYIC-PQATCLQAANK-KNRLGRSL-------QA-S-VPETLYQTL-WQR----L-APSHH-------------
ref|WP_315785635.1|   ------------EG----M----GRSAYIC-PNMSCLSAAQK-KNKLGRSL-------HA-S-VPETLYQTL-WQR----L-SQSNPPNQIEL-------
gb|MDP8965164.1|      ------------KG----M----GRSAYLC-PQASCLTAAQK-KNRLGRSL-------QA-A-VSEEVYETL-WQR----L-AAQPAAQEHKVEGEP---
ref|WP_015194852.1|   ------------EG----M----GRSAYLC-PEEDCLLKARH-KNRLGRSL-------RA-N-IPEEIYRSL-AQC----L-KT----------------
ref|WP_006528507.1|   ------------QG----Q----GRSAYLC-PQEDCLRIAQQ-KNRLGKAL-------KI-Y-VSEEIYQKL-WTR----L-SYYS--------------
gb|MBC6423737.1|      ------------LG----M----GRSAYLC-PEQECLKAAQK-KNRLGKAL-------RA-R-VPQEIYQTL-WER----L-GAVSP---GSA-------
gb|MBW4517440.1|      ------------EG----I----GRSAYLC-PQESCLQVAQK-KNRLGRSL-------KA-A-VPEELYQSL-WQR----L-SAPPTQCHLEKAKLE---
ref|WP_102221303.1|   ------------EG----M----GRSAYIC-PHMSCLSAAQK-KNKLGRSL-------HA-S-VPETLYQTL-WQR----L-SQSNLPNQIEH-------
gb|MBE9051640.1|      ------------QG----M----GRSAYIC-PQMSCLQAAQK-KNRLGRSL-------HA-S-VPETLYQTL-WQR----L-AHSH--------------
gb|MEB3268310.1|      ------------HG----M----GRSAYLC-PQADCLRAAQK-KNRLGRAL-------RT-A-VGDRLYDQL-WER----L-NATGS-------------
dbj|BAY27136.1|       ------------EG----M----GRSAYIC-PQTSCLQAAQK-KNRLGRSL-------HA-T-VPETLYQTL-WQR----L-AKSNN-----------HS
ref|WP_190744460.1|   ------------EG----M----GRSAYIC-PQTSCLQAAQK-KNRLGRSL-------HA-T-VPETLYQTL-WQR----L-TKSNH-----------HS
gb|MCS6791070.1|      ------------KG----M----GRSAYLC-PSASCLTAAQR-KNRLGRAL-------KA-T-VPIEIYEKL-WQR----L-SAA-AAATPTKS------
ref|WP_015144681.1|   ------------RG----M----GRSAYIC-PQASCLTAACQ-KNRLGRAL-------KV-S-IPDRIYESL-WER----L-ATV---------------
tpg|HYW19585.1|       ------------QG----M----GRSAYIC-PQMSCLQAAQK-KNRLGRSL-------RA-S-VPETLYQTL-WQR----L-AQSH--------------
gb|MCJ8278778.1|      ------------EG----M----GRSAYIC-PNQSCLSVAQK-KNKLGRAL-------RA-S-VPETLYTTL-WQQ----L-SRL---------------
gb|RMF20653.1|        ------------QG----M----GRSAYIC-PQISCLLQAQK-KNRLGKIL-------KA-K-IPETIYQTL-RAR----L-DIS---------------
gb|NRB09489.1|        ------------EG----M----GRSAYIC-PKSSCLSAAQK-KNRLGRSL-------HT-S-VPETVYQQL-WQR----L-AQTE--------------
gb|MDJ0736158.1|      ------------EG----M----GRSAYIC-PKASCLSAAGK-KNRLGRAL-------RA-S-VPETLYQTL-WQR----L-TQTN--------------
gb|NJL87205.1|        ------------QG----M----GRSAYLC-PQASCLQAAQK-KNRLGRSL-------RT-S-VPSTIYETL-ERR----L-E-----------------
ref|WP_261207255.1|   ------------TG----M----GRSAYLC-PEASCLKTAQK-KNRLGRSL-------KA-A-VPPELYSTL-GQR----L-PLSAQ---SEQPVGTLTP
ref|WP_008274651.1|   ------------QG----M----GRSAYLC-PNKSCLTLASQ-KNRLGRGL-------KA-S-VPDSIYQTL-WER----L-A-----------------
gb|MDJ0845508.1|      ------------QG----M----GRSAYLC-PKKSCLTLATQ-KNRLGRRL-------KA-S-IPETIYQQL-WER----L-G--KS---PSDP------
ref|WP_062247903.1|   ------------EG----M----GRSAYIC-PNMSCLSAAQK-KNKLGRSL-------HA-S-VPETLYQTL-WQR----L-SQSNLPNQIEL-------
gb|MDF5707201.1|      ------------RG----M----GRSAYIC-PQASCLQAAQR-KNRLGRSL-------HA-S-VPETLYQSL-SQR----L-ARSN--------------
ref|WP_190555052.1|   ------------QG----M----GRSAYLC-PEVSCLQAAQK-KNRLGRSL-------KA-S-VPPDLYQTL-EQR----L-AKARA-------------
gb|TAD81818.1|        ------------LG----M----GRSAYLC-PQAECLKAAQK-KNRLGRSL-------KA-P-VSDELYQTL-WQR----L-ATSE--CAG---------
gb|MBF2035119.1|      ------------HG----M----GRSAYLC-PTAACLQAAQK-KNRLARAL-------RV-T-VPETLYQVL-WQR----LAEDSAS---PRTPDPDATF
gb|MCY7283860.1|      ------------EG----I----GRSAYLC-PQESCLQAAQK-KNRLGRSL-------KA-P-VSEEIYQRL-WQR----L-SVPLTPCHLEEAKLE---
ref|WP_106289357.1|   ------------YG----M----GRSAYIC-PQESCLEAARK-KNRLGRSL-------KA-N-VPEAIYQKL-WQR----L-DTNQ----L-------LS
ref|WP_200989746.1|   ------------QG----M----GRSAYIC-PQMSCLQAAQK-KNRLGRSL-------HA-S-VPETLYQTL-WHR----L-AESH--------------
ref|WP_026079831.1|   ------------GG----M----GRSAYLC-PRADCLQAAQK-KNRLGRSL-------KT-Q-VPLTIYEEL-WQR----L-NAKIQ-------------
ref|WP_196521436.1|   ------------QG----M----GRSAYIC-PQASCLQAAQK-KNRLGRSL-------HA-S-VPETLYQSL-SQR----L-AHSN--------------
gb|NJK27659.1|        ------------RG----M----GRSAYLC-PQASCLKSAQK-KNRLGRVL-------KA-S-VPEEVYETL-WQR----L-NEGG--------------
ref|WP_297080144.1|   ------------YG----M----GRSAYLC-PTASCLQAAQR-KDRLGRSL-------KV-S-VPEAVYQTL-RQR----L--AVDS-------------
gb|NJL10525.1|        ------------EG----M----GRSAYIC-PSKSCLAMAQK-KNRLGRAI-------RA-S-VPETLYKTL-SSR----L-TVTST-------LQQ---
ref|WP_124976615.1|   ------------QG----M----GRSAYVC-PQKSCLRTATQ-KNRLGKIL-------KA-P-VPDSIYESL-WER----L-EHFSTQGQSHEKTP----
gb|MBW4673665.1|      ------------RG----M----GRSAYIC-PQASCLQAAQK-KNRLGRSL-------HA-S-VPETLYQSL-LQR----L-APSN--------------
ref|WP_261198198.1|   ------------TG----M----GRSAYLC-PEASCLKTAQK-KNRLGRSL-------KA-A-VPPELYSNL-WQR----L-PLSAQ---SEHRAATPT-
ref|WP_015128246.1|   ------------QG----M----GRSAYIC-PQTSCLQAAQK-KNRLGRSL-------HA-A-VPEALYQTL-WQR----L-AQSN--------------
gb|TAF21607.1|        ------------QG----I----GRSAYIC-PKSSCLQAAQK-KDRLGRSL-------RE-S-VPETLYQ------------------------------
ref|WP_009458837.1|   ------------EG----M----GRSAYIC-PHMGCLSAAQK-KNKLGRSL-------HA-S-VPETLYQTL-WQR----L-SQSNLPNQIEH-------
ref|WP_190980837.1|   ------------EG----M----GRSAYLC-PQVTCLQAAQK-KNRLGRSL-------QA-S-VPETLYQTL-WQR----L-APSHH-------------
ref|WP_283761350.1|   ------------------M----GRSAYLC-PQGDCLKAAQK-KNRLGRSL-------KA-S-VPPSVYERL-WSR----L-Q-------AVDDP-----
gb|MBU6184880.1|      --------Q-LDCG----I----GRSAYLC-PQKSCLLSAQR-KNRLGRSL-------KA-D-VPSEIFYQL-EQR----L-LP---------------L
ref|WP_193197802.1|   ------------MG----M----GRSAYIC-PQVSCLQAAQK-KNRLGRSL-------HA-S-VPEALYQTL-WQR----L-AQSNSQKKT---------
ref|WP_015114806.1|   ------------EG----M----GRSAYIC-PQVTCLQAANK-KNRLGRSL-------QA-S-VPEALYQTL-WQR----L-APSHH-------------
ref|WP_011318669.1|   ------------EG----M----GRSAYIC-PQMSCLQAAQK-KNRLGRSL-------QA-S-VPDTLYQKL-WQN----L-AQNDP-------------
gb|MBC6477661.1|      ------------LG----M----GRSAYLC-PEQECLKAAQK-KNRLGKAL-------RA-R-VPQEIYQTL-WER----L-GAVSP---GSA-------
gb|NDJ22983.1|        ------------MG----M----GRSAYIC-PQVSCLQAAQK-KNRLGRSL-------HA-S-VPEALYQTL-WQR----L-AQSNSQNKT---------
gb|RAM50821.1|        ------------EG----M----GRSAYIC-PNMSCLSTAQK-KNRLGRSL-------HA-S-VPETLYQTL-WQH----L-AQSN--------------
```

```
gb|MCZ0898804.1|      ------------HG----I----GRSAYLC-REDDCLKAAQK-KNRLGRSL-------KA-P-VSDELYQTL-WQR----L-ATSD--CTSY----S---
gb|MBW4554427.1|      ------------QG----M----GRSAYIC-PQMSCLQAAQK-KNRLGRSL-------HA-S-VPETLYQTL-WQR----L-AHSH--------------
ref|WP_179047786.1|   ------------QG----M----GRSAYIC-PQMSCLQAAQK-KNRLGRSL-------QA-T-VPETLYQTL-WQN----L-AQNN--------------
ref|WP_071599862.1|   ------------EG----M----GRSAYIC-PQQSCLLAAQK-KNRLGRSL-------RA-S-VPEALYHRL-WQR----L-SQSNNQKQT---------
ref|WP_207087020.1|   ------------TG----M----GRSAYLC-PEASCLKTAQK-KNRLGRSL-------KA-A-VPPELYSTL-GQR----L-PLIAQ---SDHPPATPTQ
gb|MDZ8042175.1|      ------------RG----M----GRSAYIC-PQASCLQAAQK-KNRLGRSL-------HA-S-VPETLYQSL-LQR----L-ARSN--------------
ref|WP_102207255.1|   ------------EG----M----GRSAYIC-PNMSCLSAAQK-KNKLGRSL-------HA-S-VPETLYQTL-WQR----L-SQSNLPNQIEL-------
ref|WP_066384474.1|   ------------QG----M----GRSAYIC-PKDSCLQAAQK-KNRLGRSL-------RA-T-VPETLYQTL-WQH----L-AQNN--------------
gb|PSB20742.1|        ------------QG----M----GRSAYLC-PNADCLQMAQK-KNRLGRSL-------KC-H-VPPNIYEAL-WQR----L-PVSVD-------------
ref|WP_190697022.1|   ------------QG----M----GRSAYIC-PQATCLQAAQK-KNKLGRSL-------QA-S-VPETLYQTL-WQR----L-APSHH-------------
gb|MCT7955770.1|      ------------TG----M----GRSAYLC-PEASCLKTAQK-KNRLGRSL-------KA-A-VPPELYSNL-WQR----L-PLSAQ---SEHRGATPTQ
ref|WP_254010714.1|   ------------RG----M----GRSAYIC-PQANCLTTAQK-KNRLGRAL-------RV-S-VSEELYQLL-WQR----L-AAMAE---EATE-GSSPE
gb|NBD15363.1|        ------------QG----M----GRSAYIC-PCASCLKSAQH-KNRLGRAL-------KV-K-IPANIYEQL-WER----I-N-QSS---PPSTFTSKR-
ref|WP_283755094.1|   ------------WG----M----GRSAYLC-PKADCLKAAQK-KNRLGRSL-------VA-P-VTPFVYEAL-WSR----L-KAVEA-----------DP
gb|MDJ0534388.1|      ------------EG----M----GRSAYIC-PQADCLQQARQ-KNRLGRSL-------KA-K-IPDQIYQSL-QQK----L-KINLE---P---------
gb|MBW4482298.1|      ------------EG----M----GRSAYLC-PQADCLRQAQK-KSRLGRAL-------KV-N-VPEEIYQRL-WHQ----L-EAAGV---K---SSAIA-
gb|MBD3880571.1|      ------------QG----M----GRSAYLC-PTVDCLQLAQK-KDRLGRSL-------KS-P-VSKDTYQTL-WQR----L-SILDK-------------
ref|WP_271731773.1|   ------------QG----M----GRSAYIC-PQMSCLQAAQK-KNRLGRSL-------HA-S-VPETLYQTL-WQR----L-GQTH--------------
gb|MBW4634383.1|      ------------EG----M----GRSAYIC-PQQSCLLAAQK-KNRLGRSL-------RA-S-VPEALYHTL-WQR----L-SQGNNQKQA---------
ref|WP_193999244.1|   ------------MG----M----GRSAYIC-PQPSCLQAAQK-KNRLGRSL-------HA-S-VPEALYQTL-WQR----L-AQNN--------------
gb|NJN72043.1|        ------------QG----M----GRSVYLC-QNHDCLAIAKR-KNSLGRSL-------KK-E-IPPEIHQTL-EAR----L-KAQIL-------------
ref|WP_016950665.1|   ------------QG----M----GRSAYIC-PQTSCLQAAQK-KNRLGRSL-------HG-T-VPETLYQTL-WQR----L-TENN--------------
ref|WP_191758834.1|   ------------MG----M----GRSAYIC-PQVSCLQAAQK-KNRLGRSL-------HA-S-VPEALYQTM-WQR----L-AQSNSQNKT---------
gb|MBD2237840.1|      ------------EG----M----GRSAYIC-PQTSCLQAAQK-KNRLGRSL-------HT-T-VPETLYQTL-WQR----L-ANSN--------------
ref|WP_283767879.1|   ------------------M----GRSAYLC-PQADCLKAAQK-KNRLGRSL-------KA-S-VAPSIYEKL-WSR----L-K-------EVDDP-----
ref|WP_172358412.1|   ------------YG----M----GRSAYLC-PTASCLQAAQR-KDRLGKSL-------KV-S-VPEAVYQTL-RQR----I--AVDS-------------
gb|MCP6758672.1|      ------------EG----M----GRSAYIC-PNMSCLSAAQK-KNRLGRSL-------HA-S-VPETLYQTL-WQR----L-AQSN--------------
ref|WP_015228055.1|   ------------QG----M----GRSAYVC-PCASCLKTAQH-KNRLGRAI-------KA-K-IPPTIYQEL-ENR----I-N--GR---ESSENLKSLV
ref|WP_015210357.1|   ------------EG----M----GRSAYIC-PQVNCLQAAQK-KNRLGRSL-------HA-A-VPEALYQTL-WQR----L-AQSNPQNQI---------
ref|WP_104545615.1|   ------------QG----T----GRSAYLC-PQASCLQTAQK-KNLLRRSL-------RI-S-VPEAIYQAL-WQR----L-STNQSVSEARKIETTLPL
ref|WP_237991081.1|   ------------QG----M----GRSAYIC-PQQSCLQIAQK-KNKLGRSL-------RA-S-VPETLYQTL-WQH----L-TPNNS-------------
ref|WP_190882003.1|   ------------QG----M----GRSAYIC-PQASCLQAAQK-KNRLGRSL-------HA-S-VPETLYQSL-LQR----L-APNN--------------
gb|MBF2073651.1|      ------------QG----M----GRSAYLC-PQASCLQIAQK-KNRLGRSL-------KA-V-IPEEVYHIL-WQR----L-STDPASANPTADRADRTM
ref|WP_323312969.1|   ------------QG----M----GRSAYIC-PQTSCLQVAQK-KNRLGRSL-------HG-T-VPEAVYQEL-SQR----L-AQSNTQKNHI--------
gb|MBF2001555.1|      ------------QG----M----GRSAYLC-PQASCLQIAQK-KNRLGRSL-------KA-V-IPEEVYHIL-WQR----L-STDPASANPTADRADRTM
ref|WP_193967039.1|   ------------RG----M----GRSVYLC-PQVDCLRQAHQ-KNRLGRAL-------KA-N-VPEEIYQTL-WQR----L-ETAE--------------
ref|WP_190462952.1|   ------------QG----M----GRSAYIC-PQVNCLQAAQK-KNRLGRSL-------HA-A-VPEALYQTL-WQR----L-AQSN--------------
gb|MBV6625219.1|      ------------RG----M----GRSAYIC-PNPSCLSLAQK-KNKLGRAL-------RT-P-VPETLYTTL-WQQ----L-TIK---------------
ref|WP_265262465.1|   ------------GG----M----GRSAYLC-PRADCLQAAQK-KNRLGRSL-------KT-Q-VPSTIYEEL-WQR----L-NAKIQ-------------
ref|WP_281485672.1|   ------------QG----M----GRSAYIC-PQMSCLQAAQK-KNRLGRSL-------HA-S-VPETLYQTL-WQH----L-AQSN--------------
ref|WP_127053480.1|   ------------QG----M----GRSAYIC-PQTSCLQAAQK-KNRLGRSL-------HG-T-VPETLYQTL-WLR----L-TQNN--------------
tpg|HAC62310.1|       ------------QG----M----GRSAYLC-PQESCLDNASK-KNRLGRVL-------KA-T-VPKSIYESL-WER----L-ALED--------------
ref|WP_277772165.1|   ------------YG----M----GRSAYLC-PTASCLQAAQR-KDRLGKSL-------KV-S-VPEAVYQTL-RQR----L--AVDS-------------
ref|WP_190570712.1|   ------------EG----M----GRSAYIC-PQMSCLQVAQK-KNRLGRSL-------QA-S-VPETLYQTL-WQN----L-AQNDP-------------
tpg|HIK13958.1|       ------------EG----M----GRSVYLC-PQAECLKAAQR-KDRLGRSL-------KT-S-VPQAIYERL-WQR----L-ATVEN---PKSSLT----
ref|WP_169264435.1|   ------------EG----M----GRSAYIC-PQQNCLLAAQK-KNRLGRAL-------RA-S-VPEALYHTL-WQR----L-Q-----------------
ref|WP_106455349.1|   ------------QG----M----GRSAYLC-PTDSCLKNALS-KNRLGRAL-------RS-P-IPQDLVQTL-RER----L-IN----------------
ref|WP_044206893.1|   ------------QG----M----GRSAYIC-PQASCLAAAQK-KNRLGRAL-------RA-S-VPDEIYQRL-WQR----L-EPCPW---SIQRSSD---
ref|WP_071188254.1|   ------------QG----M----GRSAYIC-PQTGCLQAAQK-KNRLGRSL-------HG-T-VPETLYQTL-WQR----L-TENN--------------
gb|MEC4803202.1|      ------------QG----M----GRSAYLC-PCDSCLAVAQK-KNRLGRSL-------RA-T-IPPAIYQQL-RSR----L-PSPDT-------------
ref|WP_015135874.1|   ------------EG----M----GRSAYLC-PTPDCLTIAQR-KNRLGRSL-------RK-Q-IPSEIYQQL-TSR----L-H-----------------
gb|NER81563.1|        ------------EG----M----GRSAYLC-PKVSCLKAAQK-KNRLGRSL-------RV-S-VPDDVYDRL-WQR----L-M--PS---STLSSNELFL
ref|WP_247215999.1|   ------------------M----GRSAYLC-PQQSCLQQAQH-KNRLGKVL-------KA-P-IPPEIYATL-WAR----L-TTPDP---VPDAHPP---
```

```
dbj|BAY28476.1|       ------------EG----M----GRSAYIC-PQTSCLQAAQK-KNRLGRSL-------HA-T-VPETLYQTL-WQR----L-ANSN--------------
gb|MCL6749841.1|      ------------QG----M----GRSAYIC-PQTSCLQAAQK-KNRLGRSL-------HA-S-VPETLYQSL-LQR----L-AQSN--------------
ref|WP_190685510.1|   ------------QG----M----GRSAYIC-PQATCLQAAQK-KNRLGRSL-------QA-S-VPETLYQTL-WQR----L-APSHH-------------
gb|MBF2065700.1|      ------------EG----M----GRSAYIC-PNTTCLGVAQK-KNRLGRVL-------HA-P-VSEELYQRL-WQR----L-AQAD----S---------
ref|WP_010997971.1|   ------------EG----M----GRSAYIC-PQMSCLQVAQK-KNRLGRSL-------QA-S-VPDTLYQTL-WQN----L-AQNDP-------------
gb|MBD2409277.1|      ------------RG----M----GRSAYIC-PQASCLQAAQK-KNRLGRSL-------HA-S-VPETLYQSL-LQR----L-APSN--------------
ref|WP_006196598.1|   ------------QG----M----GRSAYIC-QQMSCLQASQK-KNRLGRSL-------RA-S-VPETVYQTL-WQR----L-AQSY--------------
ref|WP_088239620.1|   ------------EG----M----GRSAYIC-PQPSCLSAAQK-KNRLGRSL-------HA-S-VPETLYEIL-WQR----L-AQYD----E---------
gb|MBP5972342.1|      ------------QG----M----GRSAYIC-PQQNCLLAAQK-KNRLGRAL-------RA-S-VPEALYHTL-WQR----L-H-----------------
ref|WP_193997393.1|   ------------QG----M----GRSAYIC-PQMSCLQAAQK-KNRLGRSL-------HA-S-VPEALHQTL-WQR----L-AQSH--------------
tpg|HLO50431.1|       ------------RG----M----GRSAYLC-PEADCLKAAQK-KNRLGRSL-------KA-S-VSNELYQTL-WQR----LATSPPA---SQ----PGSA
ref|WP_146296814.1|   ------------QG----M----GRSAYIC-PCENCLKTAQH-KNRLGRAL-------KA-Q-IPEEIYEQL-WEK----V-AQVST---PQP---STS-
gb|MBW4616862.1|      ------------MG----M----GRSAYIC-PRTSCLQLAQK-KNRLGRSL-------HA-S-VPEALYQTL-SQR----L-AQSNNQNKN---------
ref|WP_323360326.1|   ------------QG----M----GRSAYIC-PQTTCLQAAQK-KNRLGRSL-------HG-I-VPETLYQTL-WQR----L-TESN--------------
ref|WP_254564844.1|   ------------TG----M----GRSAYLC-PEASCLKMAQK-KNRLGRSL-------KA-A-VPPELYSTL-GQR----L-PLIDR---SDRPPANPTP
gb|MDZ7956785.1|      ------------EG----M----GRSAYIC-PQPSCLQAAQK-KNRLGRSL-------HA-T-VPEALYQTL-WQR----L-AKSN--------------
gb|NJK29495.1|        ------------EG----M----GRSVYLC-PQTDCLKTAQK-KNRLAKVL-------RA-K-VPDPVYQTL-WER----L-ETQEG---TLLKEQSEDT
ref|WP_190965760.1|   ------------EG----M----GRSAYLC-PQATCLQTAQK-KNRLGRSL-------QA-S-VPETLYQTL-WQR----L-TPSHH-------------
ref|WP_261235470.1|   ------------TG----M----GRSAYLC-PEASCLKTAQK-KNRLGRSL-------KA-A-VPPELYSNL-WQR----L-PLSAQ---SEHRAATPT-
ref|WP_261891940.1|   ------------QG----M----GRSAYLC-PQANCLKAAQK-KNRLAKAL-------RA-K-VPDSLYEFL-WER--LEL-PSRIF-------------
ref|WP_190435367.1|   ------------KG----M----GRSAYIC-PQESCLVAARK-KNKLGRSL-------KA-P-VSEELYQTL-WQR----L-TASQ----HERSLLLVS-
ref|WP_190465083.1|   ------------RG----E----GRSAYLC-PTQSCLQAAQK-KNRLGNAL-------KT-A-VPPEIYQAL-WQH----L-SNHQASINSKPRERIASD
ref|WP_299492025.1|   ------------QG----M----GRSAYLC-PQANCLKAAQK-KNRLAKVL-------RA-K-VPDSLYQSL-WER--LES-PSRIS-------------
gb|MDF2387367.1|      ------------QG----M----GRSAYIC-PQLSCLQAAQK-KNRLGRSL-------HA-T-VPEALYQTL-WQR----L-APSN--------------
ref|WP_265235232.1|   ------------RG----L----GRSAYLC-PASECLKAAQK-KNRLGRSL-------KA-P-VSDELYQTL-WQR----L-ATSD--CTSDL--------
ref|WP_293148638.1|   ------------RG----L----GRSAYLC-PASQCLKAAQK-KNRLGRSL-------KA-P-VSDELYQTL-WQR----L-ATSD--CTSD----S---
ref|WP_015151790.1|   ------------TG----M----GRSAYLC-PEASCLKTAQK-KNRLGRSL-------KA-A-VPPELYSTL-WQR----L-ALISQ---SDNPPA----
ref|WP_271764063.1|   ------------QG----M----GRSAYIC-PQMSCLQAAQK-KNRLGRSL-------HA-S-VPETLYQTL-WQR----L-GQAH--------------
ref|WP_190473724.1|   ------------EG----M----GRSAYIC-PQVSCLQAAQK-KNRLGRSL-------QA-S-VPETLYQTL-W-H----L-AQNDP-------------
gb|MEB3313608.1|      ------------QG----M----GRSAYLC-PRAECLRRAER-KNRLGRSL-------RA-T-IPTPLFAQL-WQR----L-PADPD---MPPELSQNLG
gb|TAG88461.1|        ------------LG----M----GRSAYLC-PQAECLKAAQK-KNRLGRSL-------KA-P-VSDELYQTL-WQR----L-ATSD--SADL--------
gb|MBF2015645.1|      ------------EG----M----GRSAYIC-PNPSCLSVAQK-KNRLGRAL-------RT-S-VPETLYTTL-WQR----L-P-----------------
gb|KOP28074.1|        ------------EG----M----GRSAYIC-PNMSCISVAQK-KNRLGRSL-------HA-S-VPETLYKTL-WQR----L-AQSN--------------
ref|WP_190516861.1|   ------------DG----M----GRSAYLC-PQADCLRQAQK-KSRLGRAL-------KV-N-VPEEVYQSL-WSR----L-EGKTA-------------
dbj|BAY09723.1|       ------------EG----M----GRSAYIC-PQTSCLQAAQK-KNRLGRSL-------HA-T-VPETLYQTL-WQR----L-AKSN--------------
ref|WP_300634909.1|   ------------RG----M----GRSAYIC-QNPNCLQAAGK-KNRLGRAL-------RV-S-VPDTVYQTL-WER----L-N-QSN---SHTKCS----
gb|TVQ05455.1|        ------------EG----M----GRSAYLC-PQTDCLRQAQK-KKRLSRAL-------KA-N-VPEAIYQSL-WQR----L-ETTAV---SENT------
ref|WP_026732821.1|   ------------EG----M----GRSAYIC-PQASCLSVAQK-KNKLGRSL-------HA-S-VPEALYQTL-WQR----L-SQSNLPNQIEL--------
ref|WP_318699068.1|   ------------------M----GRSAYLC-PQADCLKAAQK-KNRLGRSL-------RA-S-VAPLVYEAL-WSR----L-Q-------AVDNL-----
ref|WP_015195815.1|   ------------QG----M----GRSAYIC-QNQSCLTAAQK-KNRLGRSL-------HA-T-VPEALYQEL-WQR----L-AESQ----S---------
gb|MEB3178814.1|      ------------KG----M----GRSAYIC-PQASCLSAAQK-KNRLGRSL-------HA-S-LPDTIYETL-WQR----L-AAQTQ---PKTQPELNKV
ref|WP_096832064.1|   ------------LG----M----GRSAYLC-PQAECLKAAQK-KNRLGRSL-------KA-P-VADQLYQTL-WQR----L-ATSD--CADL--------
gb|MBD2774873.1|      ------------EG----M----GRSAYIC-PLESCLEAAQK-KNRLGRSL-------RA-S-VPSALYQTL-WQR----L-AR----------------
ref|WP_224341437.1|   ------------QG----M----GRSAYLC-PQSGCLQAAQR-KNRLGRVL-------KA-P-VPESIYQVL-WQR----L-TKSEASK-----------
ref|WP_035987080.1|   ------------TG----M----GRSAYLC-PRADCLQQAQK-KGRLGRAL-------KA-N-VPDGIYQTL-WQR----L-SAA-TVETGTDGEAV---
ref|WP_084783000.1|   ------------CG----M----GRSAYLC-PTASCLQAAQR-KDRLGRSL-------KV-S-VPEAVYQTL-RQR----L--AVDS-------------
gb|MBV9387306.1|      ------------QG----A----GRSAYLC-PQKSCLQAAQK-KNRLGRSL-------HT-A-VPEAVYQSL-WQR----L-ANQ----------SPK--
ref|WP_017652092.1|   ------------QG----M----GRSAYIC-PQMSCLQAAQK-KNRLGRSL-------HA-A-VPEALYQTL-WQR----L-AQSN--------------
ref|WP_322663056.1|   ------------RG----M----GRSAYIC-PQPGCLQAAQK-KNRLGRSL-------HA-S-VPETLYQTL-WQR----L-A-----------------
ref|WP_054468962.1|   -----------------GM----GRSAYLC-PQATCLQAAQK-KNRLGKAL-------RS-P-VPPELYQTL-WEK----I-PLMSS---TERTQDP---
gb|MDJ0660438.1|      ------------QG----M----GRSAYLC-PTKDCLTLATQ-KNRLGRRL-------KA-S-IPESIYQQL-WER----L-G--KS---TLDP------
gb|MDY6802921.1|      ------------KG----E----GRSAYLC-PQADCLKAAQK-KNRLGRSL-------KA-P-VNKEIYQAL-EER----L-AS--S---IVSGDRPETE
ref|WP_280651647.1|   ------------QG----M----GRSAYIC-PQTSCLQAAQK-KNRLGRSL-------HA-S-VPEALYQTL-CQR----L-AQPN--------------
```

```
gb|MBD1868348.1|      ------------QG----E----GRSVYLC-PQESCLRTAQK-KNRLGRSL-------KA-C-VPETIYQIL-LKR----L-LASSSPS-----------
tpg|HAG85663.1|       ------------QG----L----GRSAYLC-PQASCLAAAQK-KNRLGRSL-------HT-S-VPQELYQTL-WQR----L-DTLT----PEPTSVNSPE
ref|WP_071593325.1|   ------------TG----M----GRSAYLC-PQASCLSAAQK-KNRLGRAL-------KT-S-VPTEVYQKL-WQR----L-PS----------------
gb|TVQ51200.1|        ------------QG----M----GRSAYLC-PQVRCLQIAQK-KNRLGRSL-------KT-P-VPDAIYDHL-RQR----L-T--PQ---PHPTVNNPQD
ref|WP_190487963.1|   ------------QG----M----GRSAYLC-PHPDCLQAAQK-KNRLGRAL-------KA-A-IPEEIYQML-WQR----L-SSSGQRHAQDSASWH---
gb|MEC4984856.1|      ------------RG----M----GRSAYLC-PCETCLARAES-KNRLGRSL-------GA-Y-VPEEIYQRL-WQR----L-EVATEKEKNLASAKQ---
ref|WP_190426410.1|   ------------QG----M----GRSAYIC-PQESCLVAARK-KNKLGRSL-------KA-P-VSEELYQTL-WQR----L-TASQ----HERSLLLVS-
ref|WP_220611023.1|   ------------QG----M----GRSAYIC-PETDCLQAAQK-KNRLGRSL-------HG-T-VPETLYQTL-WQR----L-SERN--------------
ref|WP_190391042.1|   ------------QG----M----GRSAYIC-PQASCLQAAQK-KNRLGRSL-------RG-A-VPDTVYQTL-WQR----L-DHGNSQNHIQW-------
ref|WP_212664564.1|   ------------QG----M----GRSAYLC-PQEDCLKAAQK-KNRLAKVL-------RA-K-VPDSLYQSL-WDR----L-EVKS--------------
gb|NJR53419.1|        ------------EG----M----GRSVYLC-PQTDCLKTAQK-KNRLAKVL-------RA-K-VPDPVYQTL-WER----L-ETQEG---TLLKEQSEDT
ref|WP_318728151.1|   ------------------M----GRSAYLC-PQADCLKAAQK-KNRLGRSL-------RA-S-VAPSVYEAL-WSR----L-Q-------AVDNP-----
gb|TAE00837.1|        ------------LG----M----GRSAYLC-PQAECLKAAQK-KNRLGRSL-------KA-P-VSDELYQTL-WQR----L-AI-GS---PTCNYR----
gb|NJR38792.1|        ------------KG----M----GRSAYIC-PQESCLQAAQK-KNRLGRSL-------KA-T-VPEEIYQSL-WQR----L-SVPPTPCHLEKAKLE---
dbj|BAY85099.1|       ------------------M----GRSAYIC-PTNHCLSVAQK-KNKLGRAL-------RA-S-VPETLYTTL-WEQ----L-SSLDI---QK--------
gb|NJK66758.1|        ------------LG----M----GRSAYLC-PEAECLKAAQK-KNRLGRSL-------KA-P-VSDELYQTL-WQR----L-GT--S---DSASYRRKTD
ref|WP_169616303.1|   ------------EG----M----GRSAYLC-PTQDCLRAAQR-KNRLGRAL-------RA-K-VPDNVYKTL-GQW----L-EGRYD---ENMGLD--FE
dbj|BAZ53121.1|       ------------QG----M----GRSAYIC-PQLSCLQAAQK-KNRLGRSL-------HA-T-VPEALYQTL-WQR----L-APSN--------------
ref|WP_144864508.1|   ------------RG----M----GRSAYLC-PQIDCLVKAKQ-KKRLQKTL-------KT-A-VSEQIYQVL-ENR----L-ITQP----K---------
gb|MBW4669271.1|      ------------EG----M----GRSAYIC-PQTECLSAAQK-KNRLGRSL-------HA-S-VPETLYQVL-WQR----L-AQSN--------------
ref|WP_193933574.1|   ------------QG----M----GRSAYLC-PQASCLQTAQK-KNLLRRSL-------RT-S-VPEAMYQAL-WQR----L-STNQLVSN-QKIETTLPI
gb|MBD2363150.1|      ------------EG----M----GRSAYIC-LQQSCLQTAQK-KNKLGRSL-------RA-S-VPETLYQTL-WQH----L-PPNNP-------------
ref|WP_271941877.1|   ------------------M----GRSAYLC-PQADCLKAAQK-KNRLGRSL-------RA-S-VAPSVYEAL-WSR----L-Q-------AVDNP-----
ref|WP_190409163.1|   ------------EG----M----GRSAYIC-PQMSCLQAAQK-KNRLGRSL-------QA-S-VPDTLYQTL-WQN----L-AQNDP-------------
ref|WP_198126109.1|   ------------QG----M----GRSAYIC-PQLICLQAAQK-KNRLGRSL-------HA-T-VPEALYQTL-WQR----L-APSN--------------
ref|WP_137906635.1|   ------------QG----M----GRSAYIC-PQETCLQVAQK-KNRLGKSL-------RG-A-VPDAVYQTL-WQR----L-NHSNPQ----------DI
ref|WP_275521808.1|   ------------MG----M----GRSAYLC-PEAECLKAAQK-KNRLGRSL-------KA-S-VSNELYQTL-WQR----LATPSPA---SH----KLSA
gb|MBW4594787.1|      ------------EG----M----GRSAYIC-PQQNCLLAAQK-KNRLGRAL-------RA-S-VPEALYHTL-WQR----L-H-----------------
gb|MDJ0678327.1|      ------------SG----M----GRSAYLC-PTRDCLQKASQ-KRRLERSL-------KA-K-VPQQIYQRL-EAR----L-ESQNL-------------
gb|QSJ17240.1|        ------------QG----M----GRSAYIC-PQLSCLQAAQK-KNRLGRSL-------HA-A-VPETLYQTL-WQR----L-APSN--------------
gb|NEO84548.1|        ------------QG----M----GRSAYLC-PQLSCLRTAQK-KDRLRRSL-------KT-S-VPDEIYHQL-QER----L-T-----------------
gb|OCQ94358.1|        ------------RG----M----GRSVYLC-PEAECLKVAQK-KNRLGRSL-------KA-S-VSNEIYQTL-WQR----L-AT-------SSPASQ---
ref|WP_012168150.1|   ------------QG----M----GRSAYLC-PQANCLKAAQK-KNRLAKAL-------RA-K-VPDSLYESL-WER--LEL-PSRIF-------------
gb|PSO47335.1|        ------------QG----M----GRSAYIC-P-SSNCLKAAQYKNRLGRAL-------KA-K-VPPEIYEQL-WER----L-KRHSS---PSPTSASDS-
ref|WP_193847003.1|   ------------QG----M----GRSAYIC-PQETCLQVAQK-KNRLGKSL-------RG-A-VPDAVYQTL-WQR----L-NHSNPQ----------DL
dbj|BAZ90194.1|       ------------QG----M----GRSAYIC-PQAECLHKAQR-KNKLVRSL-------KG-A-VPEIVYETL-WQR----L-SQEN--------------
tpg|HAT12760.1|       ------------IG----M----GRSAYLC-PQAECLKAAQK-KNRLGRSL-------KA-P-VSDEQYQTL-WQR----L-ATSDSVCKEL--------
gb|MCY7384728.1|      ------------RG----L----GRSAYLC-QEAECLKAAQK-KNRLGRSL-------KA-P-VSNELYQTL-WQR----L-ATSD--LTSDC-------
ref|WP_015226430.1|   ------------QG----M----GRSAYVC-PCASCLKTAQH-KNRLGRAL-------KV-K-IPDDIYDQL-WER-------VKNV-------------
ref|WP_236507116.1|   ------------LG----M----GRSAYLC-PQAECLKAAQK-KNRLGRSL-------KA-P-VAEELYQTL-WQR----L-ATPD--CGDL--------
ref|WP_012164444.1|   ------------QG----M----GRSAYLC-PQADCLKAAQK-KNRLAKVL-------RA-K-VPDSLYQAL-WKR----L-EA----------------
gb|TAF06084.1|        ------------QG----M----GRSAYIC-PQTGCLQAAQK-KNRLGRSL-------HG-I-VPETLYQTL-WQR----L-TESN--------------
gb|NJL79274.1|        ------------EG----M----GRSAYIC-PNPSCLSVAHK-KNRLGRAL-------RT-S-VPETLYTTL-WQR----L-P-----------------
ref|WP_315863139.1|   ------------RG----M----GRSAYLC-PTAECLKVAKQ-KKRLARSL-------RC-P-IPEEIFTAL-AAR----L-DAHGN-------------
gb|MBD2100510.1|      -------------G----T----GRSAYLC-PQAECLAVARR-KDRLGRAL-------KV-P-VPDEIYQIL-QQR----L-GTLP-----STAKPGFHQ
ref|WP_028082426.1|   ------------QG----M----GRSAYIC-PQETCLQVAQK-KNRLGKSL-------RG-A-VPDAVYQTL-WQR----L-NHSNPQ----------DR
gb|MCL6433641.1|      ------------QG----I----GRSAYLC-PDASCLKSAQR-KDRLGRAL-------KT-P-VPEAIYQEL-WQR----L-AKGGE---NLVEMKGSPK
ref|WP_193925389.1|   ------------EG----M----GRSAYIC-PNPSCLTVAHK-KNRLGRAL-------RT-S-VPETLYTTL-WQR----L-P-----------------
gb|PSB25882.1|        ------------RG----L----GRSAYLC-QASQCLKAAQK-KNRLGRSL-------KA-P-VSNELYQTL-WQR----L-ATSD--LTSDC-------
ref|WP_194064403.1|   ------------SG----L----GRSAYLC-RASECLKAAQK-KNRLGRSL-------KA-P-VSDELYQRL-WQR----L-ATSD--CTFDCTSDS---
ref|WP_170189095.1|   ------------QG----M----GRSAYIC-PQADCLKTAQK-KNRLGRTL-------RA-P-VPPSVYDSL-WQR----L-A--QN---DHRTVNT---
gb|MBW4686284.1|      ------------MG----M----GRSAYIC-PQVSCLQAAQK-KNRLGRSL-------HA-S-VPEALYQTL-WQR----L-AQSNSQNKT---------
ref|WP_271796938.1|   ------------QG----M----GRSAYIC-PQETCLQVAQK-KNRLGKSL-------RG-A-VPDAVYQTL-WQR----L-NHSNPKVLFNGDNKNCDN
```

```
ref|WP_071515609.1|    ------------QG----M----GRSAYIC-PDPACLQAAQK-KNRLGKAL-------KA-K-VPPEIYPTL-WQR----L-QS-------SQNRFQTNA
gb|TVP65994.1|         ------------GG----M----GRSAYLC-PTQDCLRAAQR-KNRLGRAL-------RA-K-VPDNVYKTL-GQW----L-EGRYD---ENIGLD--FE
gb|MBI1240017.1|       ------------QG----M----GRSAYIC-PQTSCLQAAHK-KNRLGRSL-------HA-S-VPEALYQTL-CQR----L-AQPN--------------
ref|WP_252659375.1|    ------------QG----M----GRSAYIC-PQADCLKTAQK-KNRLGRTL-------RA-P-VPPSVYDSL-WQR----L-A--QN---DHRTVNT---
gb|EKQ70938.1|         ------------KG----I----GRSAYLC-PHESCLKIAQK-KNRLGKAL-------KA-P-VPESIYHEL-ETR-------LHHL-------------
gb|NJR17659.1|         ------------EG----M----GRSAYIC-PQASCLTAAQK-KNRLGRSL-------HA-T-VPEALYQSL-WRI----L-ALSDPPVEKN-SPRHTHP
ref|WP_168492396.1|    ------------QG----M----GRSAYIC-PQASCLQAAQK-KNRLGRSL-------HG-T-VPETLYQTL-WQR----L-NHSNPQ----------DR
ref|WP_010471496.1|    ------------QG----M----GRSAYLC-PQADCLKAAQK-KNRLAKVL-------RA-K-VPDSLYQAL-WER----L-EVQS--------------
ref|WP_313949248.1|    -------------G----T----GRSAYLC-PQAECLAVARR-KDRLGRAL-------KV-P-VPDEIYQIL-QQR----L-GTLP-----STAKPGFHQ
ref|WP_138499475.1|    ------------GG----M----GRSAYIC-PQVSCLQAAQR-KNRLGRSL-------HX-S-VPETLYQSL-LQR----L-ARSN--------------
tpg|HLO85610.1|        ------------QG----M----GRSAYIC-PTASCLQAAQK-KNRLGRSL-------HA-A-VPEALYQTL-WQR----L-NQSNSQTHIKLE------
ref|WP_190625876.1|    ------------DG----M----GRSAYLC-PQADCLRQAQK-KSRLGRAL-------KA-N-VPEDIYQSL-WSQ----L-ESKTA-------------
ref|WP_250121821.1|    ------------------I----GRSAYIC-PQASCLQVAHK-KNLLGRSL-------RT-S-VPEALYQTL-WQR----L-ATEMP---N--SK-----
ref|WP_044105921.1|    ------------KG----M----GRSVYIC-PQENCLYNASK-KNLLGKML-------RT-N-VPDSIYQTL-WSR----W-QILADTESVSAKPPNS--
gb|MBW4519521.1|       ------------RG----M----GRSAYLC-PEKNCLQMARK-KNKLDRAL-------KA-S-VPAEIHEML-WQR----I-SYTPQ-------------
dbj|BAZ29108.1|        ------------EG----M----GRSAYIC-PQASCLQAAQK-KNRLGRSL-------HA-A-VPEALYQTL-WQR----L-AQSNPQNQI---------
ref|WP_071992767.1|    ------------QG----M----GRSAYIC-PQESCLQVAQK-KNRLGRSL-------HG-T-VPDTVYQTL-WQR----L-NQGD--------------
ref|WP_013190105.1|    ------------EG----M----GRSAYIC-PQTSCLQAAQK-KNRLGRSL-------HG-A-VPETVYQTL-WQR----L-TVSN----T---------
ref|WP_015214913.1|    ------------QG----M----GRSAYIC-PQTSCLQVAQK-KNRLGRSL-------HG-T-VPEAVYQEL-NQR----L-AQSNTQKNHI--------
gb|MDJ0796006.1|       ------------EG----M----GRSAYIC-PQSSCLSAAQK-KNRLGRSL-------HT-S-VPETLYKTL-WQR----L-NQNN--------------
gb|MDY6901569.1|       ------------QG----M----GRSAYIC-PTPSCLSVAQK-KNKLGRAL-------RA-S-VPETLYTTL-WQQ----L-SCLDTQK-----------
gb|MBW4650502.1|       ------------EG----M----GRSAYLC-QNSSCLAAAQK-KNRLGRSL-------RA-S-VPEEIYKTL-WQR----L-AAMPVAENAQVATHSPKK
ref|WP_214431970.1|    ------------RG----M----GRSAYIC-PQLSCLQTAQK-KNRLGRSL-------HA-S-VPETLYQDL-WQR----L-A-----------------
ref|WP_007355211.1|    ------------MG----M----GRSAYLC-PEAECLKAAQK-KNRLGRSL-------KA-S-VSNEMYQTL-WQR----LATPSPP---SH----TLSA
ref|WP_163668145.1|    ------------------M----GRSAYLC-PQIDCLNSAHK-KNRLGRSL-------KA-P-VPEEIYQHL-HQL----I-KQKLA---SPR-------
gb|MBW4642782.1|       ------------QG----M----GRSAYIC-PQVSCLQAAQK-KNRLGRSL-------HA-A-VPEALYQTL-WQR----L-AQSN--------------
gb|MCM0590902.1|       ------------QG----M----GRSAYVC-PQASCLQAAGK-KNRLGRSL-------HA-T-VPETLYQTL-WQR----L-AQSN--------------
gb|RMF69880.1|         ------------KG----M----GRSVYLC-PQASCLQTAQR-KDRLGKSL-------RV-A-VPESVYQTL-WER----L-TSDNL-------------
ref|WP_199250663.1|    ------------AG----M----GRSAYLC-PQATCLKAAQK-KNRLGQAL-------RA-A-VPLELYQSL-WQR----L-SPAPF---ADPQCPK---
tpg|HIK55694.1|        ------------EG----M----GRSAYLC-PQESCLQAAQK-RDRLGRSL-------KA-P-VPPEIYQLL-YQR----L-SAGKS---ETPAGATAAT
gb|MBD0267699.1|       ------------QG----M----GRSAYLC-PHPECLQAAQK-KNRLGRAL-------KA-A-IPEEIYQML-WQR----L-SSSGQRHAQDSASWH---
ref|WP_009344612.1|    ------------QG----M----GRSAYIC-PQAECLHKAQK-KNKLVRSL-------KG-T-VPETVYETL-WQR----L-GQEN--------------
ref|WP_190700710.1|    ------------DG----M----GRSAYLC-PEADCLRQAQK-KSRLGRAL-------KV-N-VPEEVYQSL-WSQ----L-EGKTA-------------
ref|WP_168646496.1|    ------------QG----M----GRSAYIC-PQESCLQVAQK-KNRLGRSL-------HG-T-VPDTVYQAL-WQR----L-NHSD--------------
gb|MEB3164071.1|       ------------QG----M----GRSAYLC-PCASCLQAAQK-KNRLGRAL-------RV-P-VPPDLYTTL-WDR----L-SVNCS-------------
gb|NJL35745.1|         ------------QG----E----GRSVYLC-PRESCLRTAHK-KNRLGRSL-------KA-C-VPEGIYQTL-WQR----L-SVIDP-------------
gb|KPQ38479.1|         ------------RG----M----GRSAYIC-PQADCLKTAQK-KNRLGRTL-------RA-P-VPPSVYDSL-WQR----L-A--QN---DHRTVNT---
ref|WP_190679736.1|    ------------QG----E----GRSVYLC-PQESCLRTAQK-KNRLGRSL-------KA-C-VPETIYQIL-WKR----L-SASSSQS-----------
ref|WP_159789286.1|    ------------RG----M----GRSAYIC-PQADCLKTAQK-KNRLGRTL-------RA-P-VPPSVYDSL-WQR----L-A--QN---DHRTVNT---
ref|WP_190641945.1|    ------------QG----E----GRSVYLC-PQESCLRTAQK-KNRLGRSL-------KA-C-VPETIYQIL-WQR----L-SASSSQS-----------
ref|WP_015083448.1|    ------------QG----M----GRSAYIC-PQESCLQAAQK-KNRLGRSL-------HG-A-VPDTVYQTL-WQR----L-NHGD--------------
ref|WP_281153046.1|    ------------KG----M----GRSAYIC-PQENCLYNASK-KNRLGKML-------RT-N-VPDSIYQTL-WNR----W-QTLS-AESVSAKLPNS--
ref|WP_198807873.1|    ------------QG----M----GRSAYLC-PQADCLRIAQR-KNRLGRAL-------KA-P-IPDSLWPQL-WQR----L-EAGVS---P--LASK---
ref|WP_072033681.1|    ------------QG----M----GRSAYIC-PQESCLQLAQK-KNRLGRSL-------HG-T-VPDTVYQAL-WQR----L-NQGD--------------
tpg|HIK31517.1|        ------------KG----M----GRSAYIC-TTANCLRAAQK-KNRLGKAL-------KA-T-VPPDLYQTL-WER----L-SLTENGESD---------
ref|WP_293246005.1|    ------------RG----L----GRSAYLC-PASQCLKAAQK-KNRLGRSL-------KA-S-VSDELYQTL-WQR----L-ATSD--CTSD----S---
gb|MBD1833872.1|       ------------KG----M----GRSAYIC-PQESCLVAARK-KNKLGRSL-------KA-P-VSEELYQTL-WQR----L-TASQ----HERSLLLAS-
ref|WP_277865451.1|    ------------QG----M----GRSAYLC-PHPECLQKAEK-KNRLGRSL-------KA-K-IPAQIYQTL-KKR--------LD--------------
ref|WP_168505662.1|    ------------QG----M----GRSAYIC-PQEGCLQAAQK-KNRLGRSL-------HG-T-VPDTVYQTL-WQR----L-NYGNPQ----------DR
gb|NEQ95601.1|         ------------KG----M----GRSAYLC-PSVQCLKIARR-KKRIGRAL-------KA-P-VPQTVYDEL-AQR----L-G--ET---VV--------
gb|MBW4459453.1|       ------------TG----M----GRSAYLC-PQADCLRQAQK-KSRLGRAL-------KV-N-VPEEVYQSL-WSQ----L-EGKTA-------------
gb|MDJ0617177.1|       ------------EG----M----GRSAYIC-PQSGCLSAAQK-KNRLGRSL-------HT-S-VPETLYKTL-WQR----L-SQKN--------------
gb|MCL1475321.1|       ------------RG----E----GRSAYLC-PTEICLQAAQK-KNRLGHAL-------KT-A-VSPEIYQTL-WKR----L-ESPQV-------------
```

```
ref|WP_272035636.1|   ------------MG----M----GRSAYLC-PEAECLKAAQK-KNRLGRSL-------KA-S-VSNEMYQTL-WQR----LATPSPP---SH----TVSA
ref|WP_106367757.1|   ------------RG----M----GRSAYLC-PQVDCLQAAQK-KNRLGRSL-------RT-S-IPSGVYDTL-WQR----L-------------------
ref|WP_096670737.1|   ------------QG----M----GRSAYIC-PQESCLQVAQK-KNRLGRSL-------HG-T-VPDTVYQAL-WQR----L-NHGD--------------
ref|WP_099701299.1|   ------------QG----M----GRSAYLC-PQASCLQTAQK-KNLLRRSL-------RT-S-VPEAMYQAL-WQR----L-STNQLVSN-QKIETTLPI
ref|WP_224085747.1|   ------------EG----M----GRSAYIC-QQVSCLQAAQK-KNRLGRSL-------KA-S-VPDTLYQTL-W-H----L-AQNDP-------------
gb|TAE05943.1|        ------------RG----L----GRSAYLC-PASQCLKAAQK-KNRLGRSL-------KA-A-VSDELYQTL-WQR----L-ATSD--CTSD----S---
ref|WP_067774615.1|   ------------EG----M----GRSAYIC-PQMSCLQAAQK-KNRLGRSL-------KA-S-VPDTLYQTL-W-H----L-AQNDP-------------
ref|WP_015178990.1|   ------------RG----M----GRSAYLC-REAECLKAAQK-KNRLGRSL-------KA-P-VSDELYQKL-WQR----L-ATSD--CTSY----S---
gb|MCC5897105.1|      ------------RG----M----GRSAYIC-PQADCLKTAQK-KNRLGRTL-------RA-P-VPPSVYDSL-WQR----L-A--QN---DHRTVNTQRC
gb|NER36886.1|        ------------QG----M----GRSAYIC-PNATCLSAAQK-KNRLSKVL-------KA-A-VPREIYQTL-WQR----L-SEQS--------------
gb|MEB3252158.1|      ------------QG----M----GRSAYLC-PKVECLRRAER-KNHLGRSL-------RA-N-IPISLFAQL-WER----L-EVGSELGQNLSKT-----
ref|WP_230402764.1|   ------------------M----GRSAYLC-PRRECLQQAQQ-KNRLGKVL-------KT-P-IPPEIYATL-WAR----I-TIPDP---VPDTHLP---
ref|WP_099532894.1|   ------------------M----GRSAYLC-PCADCLRAAQR-KDRIGRSL-------KA-P-VPDPLYDLL-WQR----L-TAPVP---NPDSL-----
ref|WP_190534658.1|   ------------------M----GRSAYLC-PCADCLRAAQR-KDRIGRSL-------KA-P-VPDPLYDLL-WQR----L-TARVP---NPDSL-----
gb|NJN88730.1|        ------------QG----M----GRSAYLC-PTAACLQAAYK-KNRLGRSL-------KA-T-VPESIYQAL-WQR----L-AEADR---MEIVNSTPYN
ref|WP_193975681.1|   ------------SG----L----GRSAYLC-RVSECLKAAQK-KNRLGRTL-------KA-P-VSDELYQTL-WQR----L-ATSD--CTSD----S---
ref|WP_053538412.1|   ------------QG----M----GRSAYIC-PQESCLQLAQK-KNRLGRSL-------HG-T-VPDTVYQAL-WQR----L-NHGD--------------
ref|WP_072620812.1|   ------------EG----M----GRSAYLC-PTADCLQGAQK-KNRLGRSL-------KT-N-IPDAIYDAL-WER----L-N-QSS---SLADCPTVKN
gb|MDD1421071.1|      ------------QG----M----GRSAYIC-PQVSCLQAAQK-KNRLGRSL-------HG-T-VPDTVYQTL-WQR----L-NQGD--------------
ref|WP_271948199.1|   ------------QG----M----GRAAYIC-PQPNCLQLAKQ-KNRLGRAL-------KA-S-IPETLYESL-QER----L-GTAKQ---GGPNQD----
gb|NES70347.1|        ------------KG----M----GRSAYIC-PQAVCLKVAQK-KNRLGRAL-------KA-N-IQEHIYQVL-WQR----L-DSHS--------------
gb|NBO30352.1|        --------Q-LDCG----M----GRSAYLC-PQKSCLLSAQR-KNRLGRSL-------KA-D-VPPEIFYQL-EQR----L-LP---------------L
gb|MBD1828535.1|      ------------RG----M----GRSAYLC-REAECLKAAQK-KNRLGRSL-------KA-P-VSDELYQTL-WQR----L-ATSD--CTSG----S---
gb|PZV12745.1|        ------------QG----M----GRSAYLC-PQADCLRQAQK-KSRLGRAL-------KA-N-VPEAIYQSL-WQR----L-AEIEE---KEPTTITPCP
gb|PZO41402.1|        ------------QG----M----GRSAYLC-PQADCLRQAQK-KSRLGRAL-------KA-N-VPEAIYQSL-WQR----L-AEIEE---KEPTTITPGP
ref|WP_015172566.1|   ------------EG----M----GRSAYLC-RTSDCLQAAQK-KNRLQRSL-------RS-P-VPPGIYDVL-WER----L-HTQAA---LAANLPNALS
ref|WP_190754133.1|   ------------SG----M----GRSAYLC-PQADCLRQAQK-KSRLGRAL-------KV-N-VPEDVYQSL-WSQ----L-EGETA-------------
gb|UNU24151.1|        ------------RG----M----GRSAYLC-REAECLKAAQK-KNRLGRSL-------KA-P-VSDELYQTL-GQR----L-ATSD--CTSY----S---
gb|MBS3030443.1|      ------------QG----M----GRSAYIC-PQVSCLQAAQK-KNRLGRSL-------HG-T-VPDTVYQTL-WQR----L-NQGD--------------
gb|MBI4782862.1|      ------------KG----M----GRSVYLC-PQHSCLQLAQK-KDRLGRSL-------KA-V-VPPEIYQTL-WQH----L-SASDR--MIGSTMVESTA
gb|MDJ0724125.1|      ------------RG----M----GRSAYLC-PRASCLQAARK-KNRLGRNL-------KI-A-IPEEIYQIL-WDR----L-SPGNSESQ----------
gb|MBD1815077.1|      ------------RG----M----GRSAYLC-REAECLKAAQK-KNRLGRSL-------KA-P-VSDELYQTL-WQR----L-ATSD--CTSY----S---
gb|EGK88241.1|        ------------RG----M----GRSAYLC-REAECLKAAQK-KNRLGRSL-------KA-P-VSDELYQTL-WQR----L-ATSD--CTSY----S---
ref|WP_072207768.1|   ------------RG----M----GRSAYIC-QNPSCLQAAGK-KNRLGRAL-------RV-S-VPDTVYQTL-WER----L-N-QSN---SHTKCS----
gb|MCG9884819.1|      ------------RG----M----GRSVYLC-PTEACLNSAQR-KNRLSKVL-------KA-P-VPPDVYQSL-WNR----L-RTHP--------------
ref|WP_190528272.1|   ------------------M----GRSAYLC-PCADCLRAAQR-KDRIGRSL-------KS-P-VPAPLYDLL-WQR----L-TARVP---NPDSL-----
ref|WP_011057771.1|   ------------RG----M----GRSAYLC-PTAECLRVAKQ-KKRLARSL-------RC-P-IPEEIFTTL-AAR----L-DVHRN-------------
ref|WP_264321713.1|   ------------KG----M----GRSAYIC-AQAKCLKIAQK-KNRLGNAL-------KV-T-VPLQLYQTL-WQR----L-APREDGESSLGHP-----
ref|WP_029634746.1|   ------------EG----M----GRSAYIC-PQAGCLCAAQK-KNRLGRSL-------HA-S-VPETLYEAL-WQR----L-A-----------------
ref|WP_026099823.1|   ------------QG----M----GRSAYLC-PTASCLRSAQK-KNRLGRSL-------RV-P-IAETLYQTL-WQR----L-DSHP--------------
gb|MBW4568874.1|      ------------EG----M----GRSAYIC-PQASCLCAAKK-KNRLGRSL-------HA-S-VPEALYQTL-WQR----L-A-----------------
gb|MDY7022637.1|      ------------RG----M----GRSAYLC-RNLECLKAAQK-KNRLGRSL-------KA-A-IPEEIYQIL-EQR----L-SSE---------------
gb|MBD1213631.1|      ------------QG----M----GRSAYIC-PQETCLQVAQK-KNRLGKSL-------GG-A-VPDAVYQTL-WQR----L-NHSSPQ----------NR
ref|WP_071527241.1|   ------------TG----M----GRSAYLC-PRPDCLSAAQK-KKRLGRAL-------KA-S-VPDEVYQLL-WQC----L-EGTVV---EVSPASASVT
gb|PZU98241.1|        ------------EG----M----GRSAYLC-PQADCLRQAQK-KSRLGRAL-------KV-N-VPEEIYQRL-WHQ----L-EESKA---K---FGAIA-
tpg|HIK45324.1|       ------------QG----Q----GRSAYLC-PRADCLRLAQR-KNRLGRAL-------KA-S-LPETLWETL-WQR----L-GETSQ-------------
gb|OBQ35981.1|        ------------QG----M----GRSAYIC-PQETCLQVAQK-KNRLGKSL-------GG-A-VPDAVYQTL-WQR----L-NHSNPQ----------NR
gb|NCR11955.1|        ------------QG----M----GRSAYIC-PQPNCLQLARQ-KNRLGRAL-------KA-N-IPETLYESL-QER----L-GTA---------------
ref|WP_228383259.1|   ------------HG----E----GRSAYLC-PTADCLNQVRR-KNRLGRAL-------RA-S-IPEELFTVL-AER----L-TAATAPSASGEPDG----
gb|MBW4611665.1|      ------------QG----M----GRSAYIC-PQPSCLCAAQK-KNRLGRSL-------HA-S-VPEALYEAL-WQR----L-A-----------------
ref|WP_028091624.1|   ------------QG----M----GRSAYIC-PQETCLQVAQK-KNRLGKSL-------GG-A-VPDAVYQTL-WQR----L-NHSSPQ----------NP
gb|MEB3290003.1|      ------------QG----M----GRSAYLC-PRADCLRLAQR-KNRLGRSL-------RA-P-IPDSLWPQL-WQR----L-ELAAE-------------
gb|MBD0262109.1|      ------------EG----M----GRSAYIC-PQPSCLCAAQK-KNRLGRSL-------HA-S-VPEALYEAL-WQR----L-A-----------------
```

```
ref|WP_035737147.1|  ------------EG----M----GRSAYVC-QTAECLMVAQK-KNRLGRAL-------RA-P-VPEKVFQIL-WQR----L-ATDSQ---PDS-------
ref|WP_024124972.1|  --------Q-LDRG----M----GRSAYLC-PTAECLKVAQQ-KKRLARSL-------RC-P-IPEDIFTTL-AAR----L-AA---------------H
gb|OKH17997.1|       ------------RG----M----GRSAYLC-PNRDCLTLAQR-KQRLRRSL-------KK-N-IPTEIYQQL-ALR----L-N-----------------
tpg|HBK98803.1|      ------------IG----M----GRSAYLC-PEAECLKAAQK-KNRLGRSL-------KA-P-VSDEMYQTL-WQR----L-A-----------------
ref|WP_006512183.1|  ------------QG----I----GRSAYIC-PKSDCLQKAQR-KNLLKRAL-------RA-M-VPQGIYDLL-QQR----L-------------------
ref|WP_072044931.1|  ------------RG----M----GRSAYIC-PQVDCLQAAQK-KNRLGRSL-------RT-S-IPPAIYDTL-WQR----L-N-----------------
ref|WP_023068840.1|  ------------RG----M----GRSAYLC-RNAECLKIAQK-KNRLGRSL-------KA-A-IPEEIYQIL-KQR----L-PSE---------------
gb|MBF2027289.1|     ------------QG----M----GRSAYLC-PQANCLKTAQK-KNRLGKAL-------KA-P-VPESIYHEL-EAR-------LRQS-------------
ref|WP_272116663.1|  ------------EG----M----GRSAYLC-PTADCLQGAQK-KNRLGRSL-------KT-N-IPDAIYDAL-WER----L-N-QSS---SLADCPTVKN
ref|WP_193960470.1|  ------------QG----M----GRSAYIC-PQPNCLQLARQ-KNRLGRAL-------KA-N-IPETLYESL-QER----L-GTAKQ---GRSNQD----
gb|RFP56458.1|       ------------------M----GRSAYLC-PCADCLRAAQR-KDRIGRSL-------KA-P-VPDPFYDLL-WQR----L-TAPAP---NPDSL-----
ref|WP_125732142.1|  ------------QG----M----GRSAYIC-PQPNCLQLARQ-KNRLGRAL-------KA-N-IPETLYESL-QER----L-GTAKQ---GRPNQD----
ref|WP_006621525.1|  ------------QG----M----GRSAYVC-QTAECLMVAQK-KNRLGRAL-------RA-P-VPEKVFQIL-WQR----L-ATDSQ---PGG-------
ref|WP_190386203.1|  ------------QG----M----GRSAYIC-PQESCLQAAQK-KNRLGRSL-------HG-T-VPDTVYQTL-WQL----L-NHSD--------------
ref|WP_163518983.1|  ------------EG----M----GRSAYIC-PQASCLCAAQK-KNRLGRSL-------HA-S-VPEALYQTL-WQR----L-A-----------------
dbj|BAI90404.1|      ------------EG----M----GRSAYVC-QTAECLMVAQK-KNRLGRAL-------RA-P-VPEKVFQIL-WQR----L-ATDSQ---PDS-------
ref|WP_190574353.1|  ------------GG----M----GRSAYLC-PNATCLRDAER-KDRLGRSL-------KV-P-ISKTIYQTL-WSR----L-TEVST-------LSN---
gb|MDJ0508716.1|     ------------QG----M----GRSAYLC-PQESCLLIASK-KNRLGRSL-------KV-S-VPDSIYKQL-WQR----L-G--KS---FDDQ------
gb|TVR11480.1|       ------------QG----M----GRSAYIC-PQADCLKTAQK-KNRLGRTL-------RA-P-VPPSVYDSL-WQR----L-A--QN---DHRTVNT---
ref|WP_310454240.1|  ------------RG----M----GRSAYLC-PTAECLKVAKQ-KKRLARSL-------RC-P-IPEEIFTTL-AAR----L-DAHRN-------------
gb|MBC6434993.1|     ------------EG----M----GRSAYLC-PQESCLQAAQK-KDRLGRSL-------KA-T-VPEEIYRSL-WQC----L-SVPKTPCHLEGAELK---
ref|WP_002798549.1|  ------------QG----M----GRSAYIC-PQPNCLQLARQ-KNRLGRAL-------KA-N-IPETLYESL-QER----L-GTAWQ---GRPNQD----
dbj|BAZ16285.1|      ------------KG----M----GRSAYIC-PSVNCLGVAQK-KNRLGRAL-------HS-R-VSDEIYQEL-WQM----L-AQSL----P---------
gb|MBW4493842.1|     ------------GG----M----GRSAYLC-PQASCLSAAQK-KNRLGRAL-------KT-S-VPTEVYQTL-WQR----L-PS----------------
gb|EKV02414.1|       ------------------M----GRSAYLC-PQINCLNSAHK-KNRLGRSL-------KT-P-VPEEIYQHL-HQL----I-KQKLE---SIR-------
ref|WP_190665968.1|  ------------QG----M----GRSAYLC-SNESCLSAAQK-KNRLGRAL-------KA-P-VPIEIYQIL-WQR----L-SAM-----QADASKQLTA
ref|WP_089128587.1|  ------------EG----M----GRSAYIC-PQGSCLCAAQK-KNRLGRSL-------HT-S-VPEALYETL-WQR----L-A-----------------
ref|WP_293126645.1|  ------------RG----L----GRSVYLC-RESQCLKAAQK-KNRLGRSL-------KA-P-VSDELYQTL-WQR----L-ATSD--CTSD----S---
gb|MEB3224903.1|     ------------GG----M----GRAAYLC-PCHSCLELAQR-KDRLGRSL-------KK-S-VPSDIYQTL-KQR----L-DAQT--------------
gb|NET30621.1|       ------------QG----I----GRSAYLC-PQAECLRTAQK-KKRLERSL-------KA-S-VSQEVYAAL-WRR----L-------------------
gb|QSF50481.1|       ------------RG----M----GRSAYLC-PTAECLKVAKQ-KKRLARSL-------RC-P-IPEDIFTTL-AAR----L-DAHRN-------------
ref|WP_149818242.1|  ------------RG----M----GRSAYLC-PTAECLKVAKQ-KKRLARSL-------RC-P-IPEDIFTTL-AAR----L-DAHRN-------------
ref|WP_263749268.1|  ------------QG----M----GRSAYIC-PQASCLCAAQK-KNRLGRSL-------HA-S-VPETLYEAL-WQR----L-A-----------------
ref|WP_073069683.1|  ------------QG----M----GRSAYLC-PRSQCLKAAQK-KDRLSRVL-------KA-P-VSKQVYETL-SQR----L-LAETT-------------
ref|WP_271954606.1|  ------------QG----T----GRAAYIC-PQPNCLQLARQ-KNRLGRAL-------KA-N-IPETLYESL-QER----L-GTAKQ---GGPNQD----
tpg|HIK12112.1|      ------------VG----M----GRSAYLC-PQATCLKAAQK-KNRLGQAV-------RA-A-VPAELYQTL-WQR----L-SPAPS---QDHSAPNSQS
gb|MBW4506515.1|     ------------KG----M----GRSAYIC-PCQSCLVTAQK-KNRLGKAL-------RA-S-VPETLYKTL-SQR----L-AQGS--------------
ref|WP_287684335.1|  ------------QG----M----GRSAYMC-PQPNCLQLARQ-KNRLGRAL-------KA-N-IPETLYESL-QER----L-GPAWQ---GRPNQD----
ref|WP_225875273.1|  ------------RG----M----GRSAYLC-PNRDCLTLAQR-KQRLRRSL-------KK-N-IPTEIYQQL-ALR----L-N-----------------
gb|TRV14446.1|       ------------RG----M----GRSAYIC-PQPNCLQLARQ-KNRLGRAL-------KA-N-IPETLYESL-QER----L-GTAKQ---GRPNQD----
gb|MBW4477482.1|     ------------EG----M----GRSAYIC-PQASCLCAAQK-KNRLGRSL-------HA-S-VPETLYEAL-WQR----L-A-----------------
gb|NJK40366.1|       ------------QG----M----GRSAYLC-PQDICLKNAQR-KNRLAKVL-------KA-S-VPEAIYQEL-WHR----L-SAAGSIDILEDSSAG---
gb|TRU22031.1|       ------------QG----M----GRAAYIC-PQPNCLQLARQ-KNRLGRAL-------KA-N-IPETLYESL-QER----L-GTAKQ---GSPNQD----
ref|WP_002768359.1|  ------------QG----M----GRAAYIC-PQPNCLQLARQ-KNRLGRAL-------KA-N-IPETLYESL-QER----L-GTAKQ---GRSNQD----
gb|MDJ0546172.1|     ------------QG----M----GRAAYIC-PQPNCLQLARQ-KNRLGRAL-------KA-N-IPETLYESL-QER----L-GTAKQ---GRSNQD----
ref|WP_287078697.1|  ------------QG----M----GRSAYIC-PQPNCLQLARQ-KNRLGRAL-------KA-N-IPETLYESL-QER----L-GPARQ---GRPNQD----
ref|WP_226583963.1|  ------------QG----M----GRSAYLC-RDSRCLLAAQK-KDRLGRAL-------KA-P-VPPEIYQKL-WQR----L-STQPD-------------
gb|MBL1176422.1|     ------------TG----I----GRSAYLC-PQESCLQLAQK-KGRLGRVL-------KA-T-VPETIFQQL-RQR----L-------------------
ref|WP_159293949.1|  ------------QG----M----GRAAYIC-PQPNCLQLARQ-KNRLGRAL-------KA-N-IPETLYESL-QER----L-GTAKQ---LRPNQD----
ref|WP_071776908.1|  ------------------I----GRSAYLC-PNANCLSMAKK-KNGLAKAL-------RT-A-VPSEIYQIL-EAR----L-QSQSA---NITSE-----
ref|WP_190792225.1|  ------------QG----M----GRSAYLC-PDSACLQTAQR-KDRLGRAL-------KA-T-VSEEIYRTL-WQR----L-STTAT---IE--------
ref|WP_271991901.1|  ------------QG----M----GRAAYIC-PQPNCLQLARQ-KNRLGRAL-------KA-N-IPETLYESL-QKR----L-GTAKQ---GSPNQD----
ref|WP_110577867.1|  ------------QG----M----GRAAYIC-PQPNCLQLARQ-KNRLGRAL-------KA-N-IPETLYESL-QER----L-GTAKQ---GGPNQD----
```

```
ref|WP_286826366.1|   ------------QG----M----GRSAYIC-PQPNCLQLARQ-KNRLGRAL-------KA-N-IPETLYESL-QER----L-GPAWQ---GRPNQD----
gb|NCR41277.1|        ------------QG----M----GRAAYIC-PQPNCLQLARQ-KNRLGRAL-------KA-N-IPETLYESL-QER----L-GTVKQ---GRPNQD----
gb|NCR55568.1|        ------------QG----M----GRAAYIC-PQPNCLQLARQ-KNRLGRAL-------KA-N-IPGTLYESL-QER----L-GTAKQ---GGPNQD----
ref|WP_190367604.1|   ------------QG----M----GRSAYIC-PQESCLQAAQK-KNRLGRSL-------HS-T-VPDTLYQIL-WER----I-DPSDR-------------
ref|WP_150976468.1|   ------------QG----M----GRAAYIC-PQPNCLQLARQ-KNRLGRAL-------KA-N-IPETLYESL-QER----L-GTAKQ---GRPNQD----
ref|WP_149986802.1|   ------------QG----M----GRAAYIC-PQPNCLQLARQ-KNRLGRAL-------KA-N-IPETLYESL-QER----L-GTARQ---GRPNQD----
gb|MDX2096196.1|      ------------EG----M----GRSAYLC-PCEECLRIVQK-KNRLSKVL-------KV-P-VSPDLYQAL-WSR----L-PLS---------------
ref|WP_002738851.1|   ------------QG----M----GRAAYIC-PQPNCLQLARQ-KNRLGRAL-------KA-N-IPETLYESL-QER----L-GTAKQ---GRPNQD----
ref|WP_194012660.1|   ------------RG----F----GRSAYLC-RASECLKAAQK-KNRLGRSL-------KA-P-VSDELYQTL-WQL----L-DTSD--CTFD----S---
ref|WP_072721595.1|   ------------QG----M----GRSAYLC-PNPDCLRVAQK-KNRLGHAL-------RA-S-VPEELYQTL-WQR----L-ATPD--------------
ref|WP_108935646.1|   ------------QG----M----GRAAYIC-PQPNCLQLARQ-KNRLGRAL-------KA-N-IPETLYESL-QER----L-GTAGQ---LRPNQD----
ref|WP_272080703.1|   ------------QG----M----GRAAYIC-PQPNCLQLARQ-KNRLGRAL-------KA-N-IPETLYESL-QER----L-GTARQ---GRPNQD----
gb|OUC16623.1|        ------------KG----M----GRSAYLC-PNANCLREAER-KDRLSRSL-------RS-P-ISKDFYQML-WHR----L-AEP--------SR-----
ref|WP_271926239.1|   ------------QG----M----GRAAYIC-PQPNCLQLAKQ-KNRLGRAL-------KA-S-IPETLYESL-QER----L-GTAKQ---GGPNQD----
gb|MBW4467673.1|      ------------QG----M----GRSVYLC-PQADCLKLAQK-KDRLGRSL-------RL-T-VPESFYQVL-WLR----L-QSSNSPEAAASSPAVEPA
gb|NCQ85689.1|        ------------QG----M----GRSAYIC-PQPNCLQLARQ-KNRLGRAL-------KV-N-IPETLYESL-QER----L-GTAGQ---LRPNQD----
ref|WP_286626612.1|   ------------RG----M----GRSAYLC-PTAECLKVAKQ-KKRLARSL-------RC-P-IPEDIFTTL-AAR----L-DAHRN-------------
ref|WP_009782890.1|   ------------RG----M----GRSAYLC-RNAECLKIAQK-KNRLGRSL-------KA-A-IPEEIYHIL-KQR----L-PSQ---------------
ref|WP_181496891.1|   ------------RG----M----GRSAYLC-PTAACLKVAKQ-KKRLARSL-------RC-P-IPEDIFTTL-AAR----L-DAHHN-------------
gb|AFY59850.1|        ------------QG----Q----GRSAYLC-PQASCLHQARK-KNHLGRVL-------KA-P-IPAEIYTAL-ASRL-TTIREDESSSPLLERPSKLPKR
gb|MBC7883588.1|      ------------TG----I----GRSAYIC-PSDDCLRQAQR-KNRLGRAL-------KV-S-VDEALFGEL-RQR----F-EKVQM---L--PHVENAE
ref|WP_009631737.1|   ------------RG----M----GRSAYLC-PQVDCLLAAQK-KNRLGRSL-------RT-S-IPPGVYHTL-LQR----L-------------------
gb|MBW4539957.1|      ------------QG----M----GRSAYLC-PQSGCLQTARK-KDRLSRVL-------KA-P-VPESLYQAL-TQQ----L-ESLPT---VKCQRLGNDR
ref|WP_287969625.1|   ------------QG----M----GRSAYIC-PQPNCLQLARQ-KNRLGRTL-------KA-N-IPETLYESL-QER----L-GTAWQ---GRPNQD----
ref|WP_024970683.1|   ------------QG----M----GRAAYIC-PQPNCLQLARQ-KNRLGRAL-------KA-N-IPETLYESL-QER----I-GTAKQ---GRPNQD----
gb|MBX2866220.1|      ------------HG----M----GRSAYLC-PQLDCFSSAYK-KNRLGRSL-------KT-S-IPETIYEQL-QRQ----I-E--G--------------
gb|NJK61661.1|        ------------------M----GRSAYLC-PRPSCLQEARK-KNRLGRAL-------RT-S-VPEQIYETL-QHR----L-ASTAF---ELTSV-----
ref|WP_310743590.1|   ------------RG----M----GRSAYLC-PTAECLKVAKQ-KKRLARSL-------RC-P-IPEDIFTTL-AAR----L-DAHRN-------------
ref|WP_119259560.1|   ------------KG----M----GRSAYIC-PQESCLYNASK-KDRLGKML-------RT-N-VPDSIYQIL-WSR----W-QILS-AESVSAKLPNS--
ref|WP_172191971.1|   ------------RG----M----GRSAYLC-REGECLKAAQK-KNRLGRSL-------KA-P-VSDELYQTL-WQR----L-ATSD--CTSYSDR-----
ref|WP_322876998.1|   -----------------GQ----GRSAYLC-PQASCLHQAQK-KNHLGRVL-------KA-P-IPPEIYATL-HSR----L-VARPE---HPLTLPPPVS
ref|WP_310744398.1|   ------------RG----M----GRSAYLC-PTAECLKVAKQ-KKRLARSL-------RC-P-IPEDIFTTL-AAR----L-DAHRN-------------
ref|WP_088891762.1|   ------------AG----M----GRSAYLC-PSSGCLQAAQR-KDRLGRAL-------KA-N-VPVEIYQTL-WQR----L-STAAT---IE--------
ref|WP_151695992.1|   ------------QG----M----GRSAYIC-PQPNCLQLAQQ-KNRLGRAL-------KA-N-IPETLYESL-QER----I-GTAKQ---GGPNQD----
ref|WP_267384513.1|   ------------KG----M----GRSAYIC-PHESCLYNASK-KNRLGKVL-------RT-N-IPDNIYRTL-WNR----W-HRLNNAESISTKPPNS--
gb|NJO80776.1|        ------------QG----M----GRSVYLC-PQESCLQAAQK-KDRLGRSL-------KT-A-VPEEIYQTL-WQR----L-TPSPGAAPKADLDQIYG-
gb|NEO34688.1|        ------------QG----M----GRSAYIC-PSAQCIQAAQK-KNRLGRVL-------KT-S-IPPSVYDEL-ANR----L-V-----------------
ref|WP_190359885.1|   ------------QG----M----GRAAYIC-PQPNCLQLARQ-KNRLGRAL-------KA-N-IPETLYESL-QER----L-GTAKQ---GGPNQD----
ref|WP_308256896.1|   ------------------M----GRSAYLC-PQLECLNLAKK-KKRLSRAL-------KA-E-ITAEIYEQL-TKR----L-L-----------------
gb|MDS3859689.1|      -----------------GQ----GRSAYLC-PQASCLHQAQK-KNHLGRVL-------KA-P-IPPEIYATL-HSR----L-VARPE---HPLTLPPPVS
gb|MBR8831668.1|      ------------QG----M----GRSAYLC-PSPTCLKLAQQ-KNRLGRAL-------KT-H-IPETIYQTL-WKR----L-AEN-------QNN-----
ref|WP_317267730.1|   ------------EG----M----GRSAYVC-QTAECLMVAQK-KNRLGRAL-------RA-Q-VPEKVFQIL-WQR----L-ATDSQ---PDS-------
gb|REJ59451.1|        ------------QG----M----GRAAYIC-PQPNCLQLARQ-KNRLGRAL-------KA-N-IPETLYESL-QER----L-GTAKQ---GRPNQD----
gb|MBP0000506.1|      ------------RG----M----GRSAYLC-PRADCLKVAQK-KNRLGRAL-------KA-T-VPPSVYEIL-WQR----L-STATV---PSPPANCER-
ref|WP_297050161.1|   ------------RG----M----GRSAYLC-PTAECLKVAKQ-KKRLARSL-------RC-P-IPEDVFTIL-AAR----L-DAHRN-------------
tpg|HAZ43175.1|       ------------QG----M----GRSAYLC-RDTGCLLAAQK-KDRLGRAL-------KA-H-VPSEIYQEL-WQR----L-STQPD-------------
gb|MBW4619585.1|      ------------------I----GRSAYLC-PQKSCLDLAKK-KNRLGRSL-------KA-N-VPEEIYQKL-GQR----L-N-----------------
ref|WP_298613741.1|   ------------QG----M----GRSAYLC-RTADCLKVAKQ-KKRLARSL-------RC-P-IPEEIFTTL-AAR----L-NAHPN-------------
gb|MCL1465298.1|      ------------RG----E----GRSAYLC-PTEICLQAAQK-KNRLGHAL-------KT-A-VSPEIYQTL-WKR----L-SSPQV-------------
ref|WP_298975829.1|   ------------QG----M----GRSAYLC-RTADCLKVAKQ-KKRLARSL-------RC-P-IPEEIFTTL-AAR----L-NAHPD-------------
ref|WP_168571252.1|   ------------RG----M----GRSAYIC-LSEQCVRAAQK-KNRLGRSL-------KA-T-VPPQLYETL-RQR----V-VREGS-------------
ref|WP_287729185.1|   ------------QG----M----GRAAYIC-PQPNCLQLARQ-KNRLGRAL-------KA-N-IPETVYESL-QER----L-GPAGQ---LRPNQD----
ref|WP_190764830.1|   ------------EG----M----GRSAYLC-PTAQCLQAAQK-KNRLGRAL-------KA-Q-VDNSLYQAL-SDR----L-SQK---------------
```

```
gb|NJL20540.1|        ------------DG----M----GRSAYLC-PTAQCLQAAQK-KNRLGRAL-------KA-P-VDNRLYQIL-SDR----I-SPK---------------
gb|MDA0866786.1|      ------------AG----M----GRSAYLC-PKRSCLQAAQR-KKQLSRAL-------RT-P-VQAEIYKAL-AQR----L-G--RA-------------
gb|MDX2230996.1|      ------------QG----M----GRSAYLC-PEETCLKAAQK-KNRLGKAL-------KA-T-VPDSVYHEL-TCR-------LR---------------
gb|MBE9043893.1|      ------------RG----M----GRSAYLC-PNEQCFTQARL-KNRLKNAL-------RA-K-IPEHIYQNL-QER----L-S-----------------
tpg|HAO10320.1|       ------------HG----M----GRSAYLC-PNPDCLRVAQK-KNRLGHAL-------RT-S-VSEEVYQSI-RQR----L-D-----------------
ref|WP_317034091.1|   ------------EG----M----GRSAYIC-PQAACLRQAQK-KGRLGRAL-------KA-N-VPEELYQTL-WQR----L-EGADQ-------------
gb|MBW4678534.1|      ------------QG----M----GRSAYLC-SNESCLSAAQK-KNRLGRAL-------KA-S-VPIEIYQIL-WQR----L-SAM-----QADASKQLTA
gb|MDJ0570890.1|      --------R-LDQG----M----GRSAYLC-ANANCLNQANL-KNLLSRAL-------RV-K-VPEYIYQNL-QER----L-ND---------------V
ref|WP_287659929.1|   ------------QG----M----GRAAYIC-PQPNCLQLARQ-KNRLGRAL-------KA-N-IPETVYESL-QER----L-GPAGQ---LRPNQD----
ref|WP_190774570.1|   ------------EG----T----GRSAYLC-PNEACLRVAQK-KNALGRSL-------KV-P-VSPETYQML-WQR----L-SEAE----HDQTKLSQG-
ref|WP_310800080.1|   ------------QG----M----GRSAYLC-PTAECLKVAKQ-KKRLARSL-------RC-P-IPEDIFTTL-AAR----L-DAHRN-------------
ref|WP_002801176.1|   ------------QG----M----GRAAYIC-PQPNCLQLARQ-KNRLGRAL-------KA-N-IPETVYESL-QER----L-GPAGQ---FRPNQD----
ref|WP_190450280.1|   ------------SG----M----GRSAYLC-PQTTCLQLAQK-KNRLGRAL-------KA-A-VADELFQAL-WQR----L---ANS---SEAT-QHEVN
gb|MBR8827281.1|      ------------RG----M----GRSAYIC-PQESCLQRARQ-KNKLGRSL-------KT-I-VPEQIYQRL-WER----L-SDSKT---ETIPRDIK--
gb|MDC0831746.1|      ------------RG----M----GRSAYIC-PRADCLKTAQK-KNRLGRTL-------RA-S-VPPSVYSTL-WQR----L-SEIRE---RRPPNTDES-
ref|WP_007306607.1|   ------------KG----M----GRSAYLC-PRESCLTLASK-KNRLGRRL-------KA-P-IPDSIYQEL-WER----L-SKFVP---EQ----ELS-
gb|AJD57472.1|        ------------QG----E----GRSAYLC-PTVACFNQGRR-KNRLGRAL-------RS-A-IPDSVWVAL-EAR----L-SVTICPSLSEGSDG----
ref|WP_079676917.1|   ------------HG----M----GRSAYLC-PNPDCLKAAQK-KNRLGHAL-------RI-S-VSEEVYQSL-WQR----L-D-----------------
ref|WP_082901633.1|   ------------RG----M----GRSAYIC-PRADCLKTAQK-KNRLGRTL-------RA-S-VPPSVYSTL-WQR----L-SEIRE---RRPPKTDES-
ref|WP_190341424.1|   ------------HG----F----GRSTYLC-PQPTCLQLVQK-KNKLSRAL-------KA-A-VPKQIYEHL-WQQ----L-TPDSL-------------
ref|WP_149974369.1|   ------------QG----M----GRAAYIC-PQPHCLQLARQ-KNRLGRAL-------KA-N-IPETIYESL-QER----L-GTAGQ---FRPNQD----
ref|WP_019500197.1|   ------------------M----GRSAYIC-PTAACLQQAQK-KNRLGRSL-------RA-P-IPAEIWSQL-ESR----L-EAGTK---KYISI-----
gb|MBV5261761.1|      ------------RG----M----GRSAYLC-PCRSCLDIAQR-KDRLGRSL-------KK-N-VPPNIYQAL-QQR----L-DALTL-------------
ref|WP_324706802.1|   ------------VG----M----GRSAYVC-QTDECLKGAQK-KNRLGRAL-------KA-P-VPEELFQIL-WRR----L-AARSQ---QDG-------
ref|WP_293094764.1|   ------------EG----E----GRSAYIC-PQVECLKIADK-KNKLGRSL-------KA-Y-VPTEIYQSL-WQR----L-E--NS---SSKATESHY-
ref|WP_287453144.1|   ------------TG----M----GRSAYLC-PTADCLKAVQK-KDRLSRVL-------KA-P-VPPEIYQTL-QPS----L-QPKTK-------------
ref|WP_083622039.1|   ------------QG----M----GRSAYLC-PNPDCLRVAQK-KNRLGHAL-------RT-S-VPEELYQSL-WQR----L-ANPSE---MTSPCGDVSE
gb|NJP09111.1|        ------------RG----M----GRSAYLC-PQKSCLKLAQK-KNRLGRTL-------KA-F-VPESIYHEL-WRR----L-------------------
gb|MBW4420472.1|      ------------QG----M----GRSAYLC-PQSGCLQIARK-KDRLSRVL-------KA-P-VPESVYQAL-TQQ----L-ESHPL---VKRQQLGNDR
ref|WP_159249505.1|   ------------QG----M----GRAAYIC-PEPHCLQLARQ-KNRLGRAL-------KA-N-IPEKIYESL-QER----L-GPAGQ---FRPNQD----
ref|WP_083622686.1|   ------------HG----M----GRSAYLC-PNPDCLKLAQK-KNRLGHAL-------RT-S-VSEEVYQSL-WER----L-D-----------------
gb|MDX2215915.1|      ------------EG----M----GRSVYLC-PQASCLQTAQK-KDRLGRSL-------KA-P-IPESIYQIL-WQR----L-SGVAT---IEEAGLK---
ref|WP_046662795.1|   ------------QG----M----GRAAYIC-PQPHCLQLARQ-KNRLGRAL-------KA-N-IPEKIYESL-QER----L-GPAGQ---FRPNQD----
ref|WP_316429393.1|   ------------QG----M----GRSVYLC-RQASCLQAAQK-KDRLGRSL-------KA-S-VPEQIYQTL-WQR----L-VVEADRPQLDPDSDEIVR
ref|WP_040054730.1|   ------------------I----GRSAYLC-PNETCLAIASQ-KNRLGRGL-------RT-S-IPQDIYKKL-WKR----L-ESESA-------------
gb|MBF2048868.1|      ------------QG----M----GRSVYLC-RQASCLQAAQK-KDRLGRSL-------KA-S-VPEQIYQTL-WQR----L-VVEADRPQLDPDSDEIVR
ref|WP_190504524.1|   ------------SG----M----GRSAYLC-PQASCLQSAQK-KDRLGRSL-------KA-T-VPESIYQIL-WQR----L-SGVAT---IEKAGLK---
gb|MBW4692942.1|      ------------QG----M----GRSAYLC-PQANCLQLAQK-KDRLGRAL-------KA-P-VADQLYKTL-WQR----L-SDQPT---QLIDTST---
gb|MDR9403844.1|      ------------QG----M----GRSAYVC-PCPSCIKTAQH-KNRLGRAL-------KV-K-IPDDIYQGL-WER----I-KDVSE-------------
ref|WP_008203458.1|   ------------QG----M----GRAAYIC-PQSHCLQLARQ-KNRLGRAL-------KA-N-IPETLYDSL-QER----L-GPGEQ---GRANQD----
ref|WP_190759246.1|   ------------LG----M----GRSAYLC-SNESCLSAAQK-KNRLGRAL-------KA-S-VPIEIYQIL-WQR----L-SAM-----QADASKQLAA
tpg|HAX74477.1|       ------------QG----M----GRSAYLC-RDSRCLLAAQK-KDRLGRAL-------KA-P-VTPQIYQKL-WQR----L-STQPD-------------
gb|MCD8489703.1|      ------------EG----M----GRSAYLC-PTADCLRVAQK-KNRLGRSL-------KA-A-VAPIIYQAL-WQR----L-AEPQA---SPPPASDADL
ref|WP_069966942.1|   ------------EG----M----GRSAYLC-PTADCLRVAQK-KNRLGRSL-------KA-A-VAPIIYQAL-WQR----L-AEPQA---NPPPASDADL
ref|WP_204141672.1|   ------------QG----M----GRSAYLC-PNVTCLSLAQK-KNRLSRSL-------RA-P-VPQAIYQQL-GRR-------LQSA-------------
ref|WP_141724342.1|   ------------EG----M----GRSAYLC-PTADCLRVAQK-KNRLGRSL-------KA-A-VAPIIYQAL-WQR----L-AEPQA---NPPPASDADL
ref|WP_011244384.1|   ------------QG----E----GRSAYLC-PTVACFNQGRR-KNRLGRAL-------RS-A-IPDSVWVAL-EAR----L-SVTICPSLSEGSDG----
gb|NEQ33015.1|        ------------QG----M----GRSAYLC-PQVACLQAAQK-KKRLGRSL-------RT-S-VSPEIYELL-WQR----L-APLKS-------------
gb|MBW4699545.1|      ------------QG----M----GRSAYLC-PREECLRLAQR-KNRLARAL-------KA-R-VGKAVYEQL-RSE----L-AKIVK---PKASATASSE
ref|WP_168535677.1|   ------------QG----M----GRSAYIC-PQASCLQAAQK-KNRLGRCLWPAKRSLHG-T-VPETLYQTL-WQR----L-NHSNPQ----------DR
ref|WP_200669078.1|   ------------RG----M----GRSAYIC-LSEQCVRAAQK-KNRLGRSL-------KA-T-VSPQLYETL-RQR----V-VGETS-------------
ref|WP_250683588.1|   ------------QG----M----GRSAYLC-RTAECLKVAKQ-KKRLARSL-------RC-P-IPKDIFTTL-AAR----L-DAHRN-------------
ref|WP_024544928.1|   ------------RG----M----GRSAYLC-QDHRCLEMAQR-KDRLGRSL-------KK-N-VPPTIYQTL-QER----L-SSLD--------------
```

```
gb|MBE9010997.1|       ------------DG----M----GRSAYLC-PTPNCIQTAQR-KDRLSRAL-------KA-P-VPPIIYQTL-KQK----Q-TDTSS-------------
ref|WP_193855116.1|    ------------------I----GRSAYLC-PNANCLGAAKK-KNRLAKAL-------RT-A-VPSEIYQTL-ENR----L-QSQSS---GKLG------
gb|NJN19948.1|         ------------AG----M----GRSAYLC-PQRTCLQTAQK-KKQLSRAL-------RA-P-VNGGIYMAL-AQR----L-K--DL-------------
gb|MDJ0515540.1|       ------------EG----E----GRSAYIC-PQVECLKIADQ-KNKLGRSL-------KA-Y-VPVGIYQSL-WQR----L-E--NS---SSKATESHY-
gb|MDY7006698.1|       ------------EG----E----GRSAYIC-PQTECFQIARK-KNKLGRSL-------KT-Y-VSPEIYQAL-SER----L-K--NY---SSKKTKDDR-
ref|WP_190649792.1|    ------------TG----M----GRSAYLC-PTADCLKAVQK-KDRLSRVL-------KA-P-VPPEIYQTL-QPS----L-QPKTK-------------
ref|WP_036044569.1|    ------------TG----I----GRSAYLC-PTADCLKAVQK-KDRLSRVL-------KA-P-VPPEIYQTL-QPS----L-QPKTK-------------
ref|WP_058999772.1|    ------------HG----M----GRSAYLC-PNSNCIQAAQK-KDRLSRVL-------KA-P-VPPQIYQTL-KQKQ-AET-SSLT--------------
gb|MDJ0898873.1|       ------------QV----I----GRSAYIC-PNSDCLQKARR-KNLLKRAL-------RV-M-VPAQIYDLL-LKR----L-A--AE---I---------
gb|MBW4665085.1|       ------------RG----M----GRSAYLC-PQVDCLLAAQK-KNRLGRSL-------RT-S-IPPGVYDTL-WQR-------LKNN-------------
gb|MDJ0556426.1|       ------------------E----GRAAYIC-PQPDCLKIAHK-KNKLGRSL-------KA-Y-ISPEIYQSL-WER----F-N-----------------
gb|MBC8121277.1|       ------------RG----M----GRSAYLC-PREDCLRQAQR-KNRLGRAL-------KA-T-VGVHVYAML-HQK----L-TALGQ---LDQPTGDVSE
ref|WP_017287872.1|    ------------TG----I----GRSAYLC-PTADCLKAVQK-KDRLSRVL-------KA-P-VPPEIYQTL-QPS----L-QPKTK-------------
ref|WP_190376153.1|    ------------TG----M----GRSAYLC-PSPSCIQAAQK-KGRLSRAL-------KA-T-VPTELYQHL-QQK----Q-VEVHA-------------
gb|MBC7972109.1|       ------------SG----M----GRSVYLC-PHATCLQSAQK-KNRLGRAL-------KA-A-VADHLYKTL-WQR----L-SDEAM---VQPDSSVQAP
gb|MDY6937449.1|       ------------KG----M----GRSAYIC-VRAECLKIAQK-KNRLGNAL-------KV-T-IPPQLYQTL-WQR----L-ASIEDGESSLGHP-----
ref|WP_106237396.1|    --------Q-LDGG----M----GRSAYLC-PNIHCLTTAKS-KNRLKGAL-------RT-K-VPDHIYQNL-QAR----L-S-----------------
gb|MBD1842922.1|       ------------TG----M----GRSAYLC-PNPNCVQAAQK-KGRLSRAL-------KA-T-VPSEIYQHLLQQK----Q-GEVQA-------------
ref|WP_323217923.1|    ------------QG----M----GRSAYLC-RNAECLKAAQK-KNRLGRSL-------KA-A-IPEEIYQIL-WQR----A-TSE---------------
gb|NEQ45539.1|         ------------QG----M----GRSAYLC-PDMLCLKVAQK-KNRLGKAL-------RA-P-VPDAIYQQL-WQR----L-------------------
gb|MCY6491807.1|       ------------TG----M----GRSAYLC-PTADCLKAVQK-KDRLSRVL-------KA-P-VPPEIYQTL-HPS----L-QPNTK-------------
ref|WP_012954106.1|    ------------------M----GRSVYLC-PKKACLAVASQ-KNRLERGL-------KT-S-IPQNIYKKL-WER----L-ENESI-------------
ref|WP_065713394.1|    ------------RG----M----GRSAYLC-QCHHCLEMAQR-KDRLGRSL-------RK-N-VSPDIYQTL-KER----L-NSLD--------------
dbj|BAC90592.1|        ------------EG----M----GRSAYLC-PREECLRLAQR-KNRLARAL-------KA-R-VDEAVYEQL-RSE----L-AKIVK---PKASATASSE
ref|WP_012306383.1|    ------------RG----M----GRSAYLC-QCHHCLEMAQR-KDRLGRSL-------KK-N-VSPEIYQTL-KER----L-NSLN--------------
gb|MBW4471632.1|       ------------------M----GRSAYLC-PQVRCLQAAQK-KDRLGRVL-------KA-P-IAEQLYKTL-WER----L-TDAQP---IP-----PTR
ref|WP_326498313.1|    ------------TG----M----GRSAYLC-PTADCLKAVQK-KDRLSRVL-------KA-P-VPPEIYQTL-HPS----L-QPNTK-------------
ref|WP_071782506.1|    ------------HG----M----GRSAYLC-PNPDCLKVAQK-KNRLGHAL-------RI-S-VSEEVYQGL-WQR----L-D-----------------
tpg|HIK25275.1|        ------------RG----M----GRSAYLC-RTAECLKVAKQ-KKRLARSL-------RC-P-IPEEIFTTL-AAR----L-DPPPN-------------
gb|NJK36543.1|         ------------IG----M----GRSAYLC-PTTDCLQIAQR-KDRLGRAL-------KA-K-VPESIYGAL-WER----L-AP--D---SGSDLAQKTS
gb|NEP78655.1|         ------------EG----E----GRSAYIC-PQTECFQIARK-KNKLGRSL-------KT-Y-VSPEIYQAL-WER----L-K--NS---SSNTKDNC--
gb|NCO76359.1|         ------------NG----M----GRSAYIC-PQAQCLNLAQK-KKRLSRVL-------KA-Q-IPPEIYQQL-WQR----L-TD----------------
ref|WP_017293473.1|    ------------EG----M----GRSAYLC-PQAQCLNLAQK-KKRLPRAL-------KT-D-IPSEIYEQL-WQK----L-HEKEL---R---IEN---
ref|WP_104387117.1|    ------------QG----M----GRSAYIC-RQASCLQLAQK-KNRLGRCLWPARRSLQG-K-VPDTLYQTL-WQR----L-NHSDS-------------
gb|TAD79778.1|         ------------HG----M----GRSAYLC-PCTDCLRAAQR-KDRIGRSL-------KI-A-VSDSVYAQL-WQRIAPPTDSQGNPNTTGSNRPHRES-
ref|WP_066345584.1|    ------------------M----GRSAYIC-PQAQCLNLAQK-KKRLPRAL-------KT-D-IPLEIYERL-WQK----L-EYQEK---MDK-------
gb|MCL2931732.1|       ------------EG----E----GRSAYIC-GKIECLKVAHK-KNKLGRAL-------KT-Y-VPIEIYQSL-WQR----L-ESL---------------
ref|WP_046277840.1|    ------------QG----M----GRSAYLC-RNAECLKAAQK-KNRLGRSL-------KA-A-IPEEIYQIL-WQR----A-TSE---------------
gb|NJK54848.1|         --------Q-LDRG----M----GRSAYLC-PNIHCLTTAKS-KNRLKGAL-------RT-K-VPDQIYQNL-QAR----L-S-----------------
gb|MDX2244890.1|       ------------EG----M----GRSAYLC-REEACLKAAQK-KNRLGKAL-------KA-T-VPEATYHEL-TYR----L-SKDYF-------------
ref|WP_215607325.1|    ------------------M----GRSAYLC-PQVDCLNSAHK-KNRLGRTL-------KT-S-IPEDIYRQL-HQN----M-K-----------------
ref|WP_315862757.1|    ------------QG----M----GRSAYLC-PTAACLEAAQK-RDRLSRML-------KV-K-VPASIYEAL-WQR----V-QSPPAPVASTPARGDTPP
gb|MCL2923276.1|       ------------KG----E----GRSAYIC-GKIECLRAAHK-KNKLGRSL-------KA-N-VPLEIYRSL-WQK----L-ENL---------------
gb|NJL46250.1|         ------------WG----M----GRSAYLC-PQVGCLRLAQK-KNRLGRAL-------KV-Q-VPDSVWVTL-WQR----L-EASTV---GQTQLEEGQV
gb|MBW4534770.1|       --------Q-LDRG----M----GRSAYLC-PNIHCLTTAKS-KNRLKGAL-------RT-K-VPDDIYQNL-QER----L-S-----------------
ref|WP_015165576.1|    ------------------M----GRSAYIC-RNADCLKQAQK-KNRLARVL-------RA-P-VEPTIYERL-QAR----L-V-----------------
gb|NJN60647.1|         ------------NG----M----GRSAYLC-PSAECLKIAQK-KNRLSRAL-------RT-T-VGDDLYQSL-WRR----L-SLANR---SEATVNHLHG
ref|WP_190497765.1|    ------------KG----M----GRSAYLC-PDAGCLRLAQR-KNRLGRAL-------KV-P-IPDSLWPLL-WQR----L-DASTS---PTPLAVE---
gb|TAF56653.1|         ------------NG----M----GRSAYLC-PSAECLKIAQK-KNRLSRAL-------RT-T-VGEDLYQSL-WRR----L-SLANR---SEATVNHLHG
ref|WP_300873631.1|    ------------RG----M----GRSAYLC-RTAECLKVAKQ-KKRLARSL-------RC-P-IPEEIFTTL-AAR----L-DPPPN-------------
gb|MBW4581861.1|       ------------QG----M----GRSVYLC-PQATCLKLAQK-KDRLGRAL-------KA-P-VADQLYHTL-WQR----L-SSDQ----ARHQPEQTLQ
ref|WP_193935386.1|    ------------QG----M----GRSAYIC-RQASCLQLAQK-KNRLGRCLWPAQRSLQG-K-VPDTVYQTL-WQR----L-NHSDS-------------
ref|WP_124145084.1|    ------------EG----E----GRSAYIC-PQTECFQIARK-KNKLSRSL-------KT-H-VSPEIYQAL-WQR----L-K--NS---SSKK------
```

```
gb|NJK99433.1|         ------------RG----M----GRSAYLC-PQASCLQAAQR-RDRLGRSL-------KT-A-VPPALYQQL-WER----L-ALSPP---EPPVTRSQAA
gb|WRH67481.1|         ------------QG----M----GRSAYLC-PNPDCLKVAQK-KNRLGHAL-------RI-S-VSEEVYQGL-WQR----L-D-----------------
ref|WP_071789028.1|    ------------------S----GRSAYLC-PQLDCLNLAHK-KNRLGRAL-------RI-A-IPEEVYEEL-YKQ----I-QG----------------
ref|WP_072041293.1|    ------------QG----L----GRSAYLC-PTTSCLSMAQK-KNRLSKAL-------RV-S-VSETIYQQL-WQR----L-QSSSS-------------
gb|MBD1821041.1|       ------------TG----M----GRSAYLC-PSPSCIQAAQK-KGRLSRAL-------KA-T-VPTELYQHL-QQK----Q-VEVHA-------------
ref|WP_293079557.1|    ------------QG----E----GRSAYIC-PQTECFQIARK-KNKLGRSL-------KA-Y-VSPEIYQEL-WQR----L-K--NS---SSKNTKDDG-
ref|WP_310422503.1|    NTFV--------------Q----GRSAYLC-PTASCLQIAQK-KNRLGKSL-------KA-Q-VRDDIYQQL-ELL----------K---FTR-------
gb|MDJ0591465.1|       ------------GG----M----GRSAYLC-PNINCLTQAKL-KNRLRSSL-------RT-K-VPEQIYQNL-QAR----L-SG-------------SG-
ref|WP_023172451.1|    ------------RG----M----GRSAYLC-PCEECLRQAQR-KNRLARAL-------KA-R-VDETFYELL-RCE----L-AKIAK---PET-------
ref|WP_036533928.1|    ------------QG----M----GRSAYLC-PQQNCLQTAQK-KKRLERTL-------RV-K-VSQEVYQQL-WHR----L-EGSIQ-------------
ref|WP_162422901.1|    ------------QG----I----GRSAYLC-PQAECLRSVRK-KDRLSRAL-------KA-P-VPESLYQQL-LQQ----L-VVSPS---R---------
dbj|GGA05509.1|        ------------RG----E----GRSAYIC-PEIECLQIARK-KNKLGRSL-------KT-Y-VSPEIYQEL-WQR----L-K--KS---SSKNTKDDD-
ref|WP_066120908.1|    ------------------M----GRSAYLC-PQYQCLNLAQK-KKRLSRVL-------KI-D-VPSEIYQQL-WQK----L-EQTKS---QT--------
gb|UFP97105.1|         ------------EG----M----GRSAYLC-PREECLRLAQR-KNRLARAL-------KA-R-VDEAVYEQL-RSE----L-AKIVK---PQASATVSSE
ref|WP_019504603.1|    --------Q-LDRG----M----GRSAYLC-PNIDCLNQVRF-NNRLSRSL-------KA-R-VPEDIYQNL-QAR----L-KN---------------V
dbj|BAU14687.1|        ------------QG----M----GRSAYLC-PTPNCIQTAQK-KDRLSRAL-------KA-T-VPLSIYQIL-KQK----Q-AEISS-------------
gb|MCY7272735.1|       ------------QG----T----GRSAYLC-PQSECLKAAQK-KDRLSRVL-------KA-S-VSDQIYKAL-WQR----L-SE----TTSLPTTGDLGL
ref|WP_202219931.1|    ------------RG----E----GRSAYIC-PEIECLQIARK-KNKLGRSL-------KT-Y-VSPEIYQEL-WQR----L-K--KS---SSKNTKDDD-
ref|WP_310488256.1|    DT----------WM----Q----GRSAYLC-PTASCLQLAQK-KNRLGRSL-------KA-A-IPDTTYQQL-GTT----A-F--CK---SK--------
gb|MBE9099686.1|       ------------EG----M----GRSAYLC-QTEACLQTAQK-KKAIRRSL-------RV-P-ERPDIYQLL-WER----L-DKGQ--------------
gb|MBU6228775.1|       ------------QG----M----GRSAYIC-RQADCLRLAQR-KNRLGRAL-------KA-I-APAGVFDRL-WQQ----L-ATGGS-----VDL-----
gb|NKB17092.1|         ------------------M----GRSAYIC-QNASCLQQAHK-KNRLGRSL-------RS-Q-VPPEIFTQL-ELR----L-R--TN---PPIS------
gb|NES69150.1|         ------------RG----E----GRSAYIC-PQTECLQIARK-KNKLGRSL-------KT-Y-VSPEIYQAL-WQR----L-K--NS---SSQNPKDDG-
gb|MBW4553259.1|       ------------QG----A----GRSAYLC-PDASCLQSAQK-KNRLARCL-------KA-A-IPESIYTAL-WQQ----I-N--QR---EFAGQEALAT
gb|PSP19040.1|         ------------SG----M----GRSAYLC-PQRPCLQAARR-KKRLARAL-------KA-P-VPERIDRAL-EQH----L-AAR-------------E-
ref|WP_106312640.1|    NA----------VI----Q----GRSAYLC-PTASCLQIAQK-KNRIGRSL-------KA-K-ISEEIYQQL-ESI------------------------
ref|WP_204104579.1|    ------------KG----M----GRSAYLC-PQRDCLEAAQK-KNRLARSL-------KA-N-APQEVYQNI-YQT----L-RSRLA---A---IDKRN-
gb|MCU0552418.1|       ------------TG----M----GRSAYLC-PTPDCIKAVQK-KDRLSRIL-------KA-P-VPAEIYQTL-LAA----S-SSKSK-------------
gb|MBW4442872.1|       ------------LG----M----GRSAYLC-PTPNCIQAAQK-KDRLGRAL-------KV-A-IPAEIYQGL-KQK----Q-AEFHS-------------
ref|WP_106254952.1|    ------------SG----M----GRSAYLC-PQADCLKLAQK-KDRLARAL-------KA-T-VSESVYKTL-RQR----L-SSQQT-------------
emb|CAA9550821.1|      ------------QG----M----GRSAYLC-PQTNCLQAARK-KNRLGRAL-------KT-P-VPEDIYQTL-WQR----L-PGLP--------------
tpg|HCF29536.1|        ------------RG----M----GRSAYIC-PQASCLQLAQK-KNRLGRSL-------RVKA-IPEEIYQTL-WQR----L-PANQ--------------
gb|MCA1903264.1|       ------------QG----M----GRSVYVC-RQGNCLAEALK-KNRLGKAL-------RQ-P-IPEVLLAEL-QRR----V-SSSQ--------------
gb|NEQ23162.1|         ------------VG----M----GRSAYLC-PNAECLQAAQK-KNRLGRAL-------KA-P-VANSVYQAL-SDR----L-SPDQ--------------
ref|WP_293133712.1|    ------------QG----E----GRSAYIC-EKIECLKVAEQ-KNKLGRSL-------KA-Y-VPPEIYRLL-WQR----L-EKLK--------------
ref|WP_015157060.1|    ------------RG----M----GRSAYIC-PQASCLQVAQK-KNRLGRSL-------KV-KTVSEVLYQTL-LQR----L-VTEP--------------
ref|WP_121971475.1|    ------------EG----T----GRSAYLC-PQASCLEAAQK-KNRLSRML-------RT-Q-VPPEIYQAL-SQR----L-MEDLD---RGGPDKNSY-
dbj|BAZ46326.1|        ------------QG----I----GRSAYIC-PNAQCLVKAKY-KNRLNSCL-------KA-K-VPEHIYQTL-QER----L-N-----------------
gb|MBW4527443.1|       ------------LG----M----GRSAYLC-PTPNCIQAAQK-KDRLGRAL-------KV-A-IPAEIYQGL-KQK----Q-AEFHS-------------
gb|NEN88813.1|         ------------QG----E----GRSAYIC-PKTECFQIARK-KNKLGRSL-------KT-Y-VSPEIYQAL-WQR----L-K--DS---SSKNTQNDR-
gb|MDX2273513.1|       ------------LG----M----GRSAYLC-RQSRCLQDAQK-KNRLGRAL-------RA-S-IPPHIYEVL-WQR----L-ASPH--------------
ref|WP_293062802.1|    ------------QG----E----GRSAYIC-PQTECFQIARK-KNKLGRSL-------KT-Y-VSPEIYQAL-WQR----L-K--DS---SSKNTQNDR-
ref|WP_218081389.1|    ------------------M----GRSAYLC-PNASCLKQAQR-KDRLAKVL-------KA-R-VPPEVYAAL-WTR----L-QVKGQ---GTSAQVDRRR
ref|WP_015168983.1|    -----------------GM----GRSAYIC-PNLNCLQLAQK-KNRLGRSL-------RT-Q-INPEIYEIL-KQR----L-SNA---------------
ref|WP_287129737.1|    ------------------M----GRSAYLC-PTLPCLKQAQR-KDRLSKVL-------KA-R-VPTEVYNAL-WMR----L-GVSNT---TPAGG-----
gb|MCL2927522.1|       ------------QG----E----GRSAYICCGQIECLKIAHK-KNKLGRAL-------KA-Y-VSPEIYQSL-WQR----L-ESL---------------
ref|WP_192153424.1|    ------------RG----M----GRSAYIC-PQASCLQVAQK-KNRLGRSL-------KV-KTVSEVLYQTL-LQR----L-ATEP--------------
gb|NJK69321.1|         ------------SG----M----GRSAYLC-PEAECLKAAQK-KIDWGERY-------KA-P--------------------------------------
gb|MBP0016486.1|       ------------KG----M----GRSAYLC-PRRECLEAAQK-KNRLARSL-------KA-N-APPEVYQDL-YQT----L-RSRLA---A---IEERD-
gb|PZO20693.1|         ------------DG----M----GRSAYLC-RNPQCLENAQK-KNRLGKAL-------RS-P-VPPEIYKEL-HNL----L-IQRSS---EQTDLSSISS
gb|MCH9055185.1|       ------------EG----M----GRSAYLC-PTSECLKLAQQ-KKRLGRSL-------RC-A-VPDTVFSTL-KAC----L-KSSQA---DGNAVPPPET
ref|WP_099799890.1|    ------------EG----M----GRSAYLC-PTSECLKLAQQ-KKRLGRSL-------RC-A-VPDTVFSTL-KAC----L-KSSQA---DGNAVPPPET
gb|MDJ0773676.1|       ------------WG----M----GRSAYIC-QNPNCLQAAGK-KNRLGR-----------------------------------------------
```

```
gb|OWY68048.1|        ------------WG----M----GRSAYIC-PQASCLQVAQK-KNRLGRSL-------KV-KTVSEVLYHTL-LQR----L-VTEP--------------
gb|MCX7595673.1|      ------------EG----M----GRSAYIC-PQASCLSVAQK-KNK------------------------------------------------------
gb|NEQ50465.1|        ------------TG----M----GRSAYLC-PHLECLSSAQK-KNRLSRAL-------KA-S-IPEEIYQQL-HQR----I-R-----------------
gb|NJN30919.1|        ------------EG----M----GRSAYLC-PTADCLKLARK-KRRLGRSL-------RC-A-MDDGIYDLL-EAR----L-EPGQS---APD-------
gb|MDX2254087.1|      K---------FTIQLDHGM----GRSAYLC-PNASCLQIAQK-KNRLGRAL-------RM-P-IPPEVLEVL-RSR----L-ANQST---DQ--------
gb|AFY96277.1|        ------------LG----KTVIQGRSAYLC-PTASCLQIAQK-KNRIGKSL-------KA-K-ISEEIYQQL-ESI----A-ID----------------
ref|WP_229640858.1|   ------------RG----M----GRSAYIC-PDINCLTQAKT-KNRLSSSL-------RT-K-VPEHIYQDL-QER----L-RG----------------
ref|WP_068815920.1|   ------------QG----M----GRSAYLC-PKPECLTAAQK-KDRLSRVL-------KA-A-VSDQIYKAL-WHR----L-SE----TTSPPTTGDLGL
gb|MCY7336460.1|      DK----------TL----Q----GRSAYLC-PTAICLQIAQK-KNRLGRSL-------KA-K-VDEGIYQQL-GLL----K-------------------
gb|MBS9769718.1|      ------------QG----E----GRSAYICCGQIECLKIAHK-KNKLGRSL-------KA-Y-VSPEIYQSL-WQR----L-ESL---------------
ref|WP_190352545.1|   -----------------GM----GRSAYVC-QNLDCLQIAQK-KNRLGRSL-------KM-P-IPKEIFEIL-KSR----L-L-----------------
gb|PZV14006.1|        -----FQIQ-LDHG----M----GRSAYLC-QKLECLKIAQK-KNRLGRSL-------RT-P-IPTEIFEIL-KSR----M-------------------
gb|MDE5085023.1|      ------------QG----E----GRSAYIC-RKIECLKIAHK-KNKLGRSL-------KA-Y-VPPEMYTSL-WQK----L-ENL---------------
ref|WP_293122704.1|   ------------QG----E----GRSAYIC-PQTECFQIARK-KNKLGRSL-------KT-Y-VSPEIYQAL-WQR----L-K--DS---SSKNTQNDR-
ref|WP_319421161.1|   --------Q-LDRG----M----GRSAYLC-CNIKCLNQAKI-KNRLSSSL-------RT-K-VPEGIYQKL-QER----L-S-----------------
ref|WP_295619447.1|   NI----------LM----Q----GRSAYLC-PTDSCLQSAKK-KNRLSKSL-------KA-T-VTDEIYQQL-QLL----I-TSRSE-------------
gb|MCL2936457.1|      ------------QG----E----GRSAYIC-RKIECLKVAHK-KNKLGRSL-------KA-Y-IPPEMYTSL-WQK----L-ENL---------------
gb|MDE5075942.1|      ------------QG----E----GRSAYIC-RKIECLKIAHK-KNKLGRSL-------KA-Y-IPPEMYTSL-WQK----L-ENL---------------
ref|WP_293158219.1|   ------------EG----E----GRSAYIC-PQTECFQIARK-KNKLSRSL-------KT-H-VSPEIYQAL-WQR----L-K--NY---SSKK------
gb|OLP17560.1|        ------------QG----M----GRSAYLC-PQESCLQTAQK-KNRLSRSL-------KT-Q-VPQDFYQLL-WHA----L-CVLDS-------------
gb|NJM49111.1|        --------A-LDCG----M----GRSAYLC-PTLQCLQGALK-KDRLSRML-------RT-T-VPTDIYDQL-RSR----L-EG---------------P
ref|WP_244349359.1|   ------------EG----M----GRSAYLC-RQLDCLQAAQR-KQRLNKAL-------KA-S-VPDELFQRL-EAQ----L-LSHQP---SG--------
ref|WP_232432138.1|   ------------LG----KTVIQGRSAYLC-PTASCLQIAQK-KNRIGKSL-------KA-K-ISEEIYQQL-ESI----A-ID----------------
ref|WP_193989930.1|   ------------------M----GRSAYLC-PQVECLSVAHK-KNRLGRAL-------KA-M-VPDAVYQNL-YQV----I-ASAD--------------
ref|WP_235280105.1|   ------------EG----M----GRSAYLC-RQFGCLQAAQR-KQRLNKAL-------KA-P-VPDELFQRL-EAQ----L-VSNQP---SD--------
ref|WP_166274652.1|   ------------------M----GRSAYLC-PTLDCLQVARK-KRRLERSL-------RV-T-VPPQIYEAL-ENQ----L-QNPPI---SST-------
ref|WP_255131202.1|   ------------------M----GRSAYLC-PTQQCLEEARK-RRKLQRAL-------RV-N-VEDAIYSAL-ASR----L-GPEGI---SLKGI-----
ref|WP_264324473.1|   ------------QG----M----GRSVYLC-RNWDCLKLAQK-KDRLSRAL-------KV-K-VSTVLYQSL-EQR---LE-SQQQT-------------
ref|WP_310410468.1|   ------------LGTILMQ----GRSAYLC-PTAECLQLAQK-KNRLSRSL-------KA-A-VAEDIYHQL--------------GL-------------
ref|WP_259727590.1|   ------------------M----GRSAYLC-PASSCLADAKR-HRRLQRAL-------RC-E-VVPSIYAAL-DQR----L-V-----------EA----
ref|WP_115093230.1|   ------------------M----GRSAYLC-PEENCLEEATR-RKRLQKAL-------RC-Q-VPETVLAML-KQR----L-IQTPG---ESAEAD----
gb|MAH58821.1|        ------------------M----GRSAYLC-PKEECFEEARK-RKRLQKAL-------RC-Q-VPDAVLEAL-KQR----L-SSTKA---ESAEAN----
ref|WP_259703109.1|   ------------------M----GRSAYLC-PASSCLADAKR-HRRLQRAL-------RC-E-VVLSIYAAL-DQR----L-V-----------EA----
gb|NDC14930.1|        ------------------M----GRSAYLC-PNQVCLDEAKR-RKRLQRAL-------RC-Q-VAETIVETL-EQR----L-I-----------------
ref|WP_309742497.1|   ------------LGTILMQ----GRSAYLC-PTAQCLQLAQK-KNRLSRSL-------KA-A-VAEDIYHQL--------------GL-------------
ref|WP_036487578.1|   ------------RG----M----GRSAYIC-PQKTCIRQAFS-KQRLARSL-------RT-K-VSQSSSQNL-QER---LL-DRVSS-------------
gb|NJN76171.1|        --------A-LQSG----M----GRSAYIC-RNKNCLEIALT-KKRLNRAL-------KA-P-IPDSLMKQL-W-K----I-IA---------------S
gb|MBJ7364496.1|      ------------------M----GRSAYLC-PTKQCLEEARK-RRKLQRAL-------RV-N-VEDAIYSAL-ASR----L-GPEGI---SLKGI-----
gb|NBV69567.1|        ------------QG----M----GRSAYLC-PTQPCLEDARK-RRKLQRAL-------RV-N-VEDAVYSTL-AAR----L-GQRDS---AFSATLQAI-
gb|NJM97597.1|        ------------KG----M----GRSAYLC-PQRSCLNAAQK-KKRLARVL-------KA-H-VPPDIYQIL-AQR----L-TV-KD---SKNPIHPSP-
gb|MCY7407389.1|      ------------HG----M----GRSAYIC-QTEKCLLAADK-KDRLSRAL-------KT-A-VPKDLYQTL-KKT----L-STRPS-------------
ref|WP_254933944.1|   ------------QG----M----GRSAYLC-PNQECFEEAKR-RKRLQRAL-------RC-Q-VAETIVETL-EQR----L-NTSVD-------------
ref|WP_017327771.1|   ------------RG----M----GRSAYIC-RELDCLRRSQK-KNRLGKAL-------RT-P-VPPEIYNEL-QML----F-VLDSQ-------------
ref|WP_309728133.1|   HT----------WM----Q----GRSAYLC-PTASCLQLAQK-KNRIGRSL-------KA-A-ISPDIYHQL-GTL----T-F--DR---T---------
gb|MCY7367819.1|      DTLH--------------Q----GRSVYLC-PTAECLQLAQK-KNRLSKSL-------KT-Q-VSQDIYQAL-SFR----I-------------------
gb|MEC8605584.1|      ------------EG----M----GRSAYVC-ASHTCLEDARK-RRKLQRAL-------RV-P-VADSVYDAL-AKR----L-E--PE---A---------
gb|MBW4662466.1|      -------------G----M----GRSVYLC-RKPTCLELAQK-KDRLGRSL-------KA-V-VPPEIYHGL-WQH----L-SASDT--IVGST-----A
ref|WP_190401439.1|   -----FQIQ-LDHG----M----GRSAYLC-QNLDCLQIAQK-KNRLGRAL-------RT-Y-IPPEIFDIL-KSR----LETL----------------
ref|WP_161823453.1|   ------------YG----M----GRSAYLC-AQSECLQQAQK-KKRLERAL-------RT-A-IAPEIYQIL-WQR----L-SAPSV-------------
gb|MCS6943628.1|      ------------KG----M----GRSAYIC-PRQDCLEIAQK-KKRLAKAL-------KI-P-IPPEIFQQL-WQR----L-QQDPTKGYN---------
ref|WP_255116400.1|   ------------QG----M----GRSAYLC-PTQACLEEAKR-RKKLQKGL-------RC-Q-VADSIYAAL-EQR----L-NTTP----V---------
gb|AFZ48415.1|        ------------QG----M----GRSAYIC-PNSECIAIAKK-KKRLGRAL-------RT-M-VSAEIYEEL-KTR-------C----------------
gb|NUN65110.1|        AT----------RGFQIQLDCGIGRSAYLC-QSLSCLQIAQK-KNRLGRSL-------RT-P-IPPEIFEIL-KSR----L-QLTENIAVASQVRT----
```

```
ref|WP_115022254.1|  ------------------M----GRSAYLC-PEENCLEEATR-RKRLQKAL-------RC-Q-VPETVLAVL-KQR----L-IQTPG---ESAEAD----
gb|MCX5962897.1|     ------------HG----M----GRSAYIC-QTEKCLLAAEK-KDRLSRAL-------KT-A-VPKDLYQTL-KKM----L-SALPP-------------
gb|MBM5807801.1|     ------------QG----M----GRSAYLC-PTEACLEEAKR-RKKLQKAL-------RC-Q-VADSIYAAL-EQR----L-HATPS---AGSEAR----
gb|MBV2350088.1|     ------------QG----M----GRSAYLC-PTRDCLEEARR-RKKLQKGL-------RC-Q-VADSIYAAL-EQR----L-HATSS---AGSEAR----
gb|MCG9892687.1|     ------------EG----M----GRSAYLC-PQKSCVAIAPK-KKRLERAL-------KA-V-VPPEVYEQL-EQL----I-Q--EG-------------
gb|MBJ7492561.1|     ------------AG----M----GRSAYLC-PRQDCLSEAKR-RRKLQRAL-------RC-Q-VADSIYAAL-EGR----L-------------------
emb|CAI8160962.1|    ------------------M----GRSAYLC-PNEDCLEEARR-RKRLQKAL-------RC-Q-VHDAVLEAL-DQR----L-RHSTD---ESAEAK----
gb|PSI02062.1|       ------------QG----M----GRSAYLC-QTQPCLEDARK-RRKLQRAL-------RV-N-VEEAVYSTL-AAR----L-GQRDS---AFSATLQAI-
gb|OON12454.1|       ------------QG----M----GRSAYLC-QTQPCLEDARK-RRKLQRAL-------RV-N-VEDAVYSTL-AAR----L-GQRDS---AFSATLQAI-
gb|NBO27849.1|       ------------QG----M----GRSAYLC-QTQPCLEDARK-RRKLQRAL-------RV-N-VEDTVYSTL-AAR----L-GQRQS---AFSATLQAI-
ref|WP_255008768.1|  ------------AG----M----GRSAYLC-PNAACLEEARR-RKRLQRAL-------RC-Q-VADSIYASL-GQR----L-DTDPV-------------
ref|WP_255098738.1|  ------------QG----M----GRSAYLC-QTQPCLEDARK-RRKLQRAL-------RV-N-VEEAVYSTL-AAR----L-GQRDS---AFSATLQAI-
ref|WP_255096693.1|  ------------QG----M----GRSAYLC-QTQPCLEDARK-RRKLQRAL-------RV-N-VEDAVYSTL-AAR----L-GQRDS---AFSATLQAI-
ref|WP_296444202.1|  ------------QG----M----GRSAYLC-PSRDCLEEARR-RKKLQKGL-------RC-Q-VADSIYAAL-EQR----L-HATSS---AGSEAR----
ref|WP_211167718.1|  -----FQIQ-LDDG----M----GRSAYLC-KKLECLQIAQK-KNRLGRSL-------RT-Q-IPSEIFDIL-KSR----LSSTNSQKYDNIFVN-----
gb|NMF59328.1|       -----FQIQ-LDDG----M----GRSAYLC-KKLECLQIAQK-KNRLGRSL-------RT-Q-IPSEIFDIL-KSR----LSSTNSQKYDNIFVN-----
ref|WP_174235275.1|  KQ---FQIQ-LDDG----I----GRSAYLC-KNLDCLQIAQK-KNRLGRSL-------RT-H-IPPEIFDIL-KAR----L-------------------
gb|MBM5799360.1|     ------------RG----E----GRSAYLC-PSTTCLDDAKR-RKRLQRAL-------RC-Q-VADSILNCL-ETR----L-HGN------------G-
ref|WP_142655983.1|  ------QIQ-LDHG----M----GRSAYLC-KKLECLQLAQK-KNRLGRSL-------RT-Q-IPPEIFDIL-KSR----L-------------------
gb|MCP4972515.1|     ------------------M----GRSAYLC-PSENCLDEARR-RKRLHKAL-------RC-Q-VPENVLEVL-QER----L-----N---SVAEAR----
ref|WP_043692418.1|  ------------------M----GRSAYLC-PTEDCLEEARR-RKRLQKAL-------RC-Q-VPDAVITVL-QER----F-SPGTG---VSAEAN----
ref|WP_011936384.1|  ------------EG----M----GRSAYVC-ASHACVEDARK-RRKLQRAL-------RV-P-VADSVYDAL-AKR----L-E--PE---A---------
gb|MCA6503255.1|     ------QIQ-LDHG----M----GRSAYLC-KKLECLQLAQK-KNRLGRSL-------RT-Q-IPPEIFDIL-KSR----L-------------------
ref|WP_131456745.1|  ------------------M----GRSAYLC-PQEACLEEARR-RKRLQKAL-------RC-Q-VPDSVVEVL-QER----L-TLQRD---TAAEAR----
gb|MCB4394176.1|     ------------------M----GRSAYLC-PEGNCLEEATR-RKRLQKAL-------RC-Q-VPETVLAVL-KQR----L-IQTPG---ESAEAD----
gb|MCS5706208.1|     ------------QG----M----GRSAYLC-QTHCCLDDARR-KKKLQRAL-------RC-Q-VADSIVEAL-EQR----L-TDGG----A---------
gb|MEB3105445.1|     ------------QG----M----GRSAYLC-QTHCCLDDARR-KKKLQRAL-------RC-Q-VADSIVEAL-EQR----L-TDGG----A---------
gb|MCL1492198.1|     ------QIQ-LDHG----M----GRSAYLC-KKLECLQLAQK-KNRLGRSL-------RT-Q-IPPEIFDIL-KSR----L-------------------
ref|WP_255087149.1|  ------------------M----GRSAYLC-PTRSCLDDAKR-QKRLQRAL-------RC-E-VAASIYLAL-ERR----L-D----------GA-----
ref|WP_269621837.1|  ------------------M----GRSAYLC-PQENCLKEAWH-RKRLQKAL-------RC-Q-VDVSVIEVL-QDR----L-NHCND---SITKAR----
ref|WP_310483930.1|  NT----------LM----H----GRSAYLC-PTDSCLQSAKK-KNRLSKSL-------KA-T-VNDDIYQQL-GLL----L-TSRSE-------------
ref|WP_156796781.1|  ------------RG----M----GRSAYLC-PRRECLDEARR-RKRLQRAL-------GC-P-VADPVLEAL-EQR----L-ACQPGSTGTQPASVATP-
gb|MDA0716923.1|     ------------------M----GRSAYLC-PNQVCLEEAKR-RKRLQRAL-------RC-Q-VADAIVETL-EKR----L-I-----------------
tpg|HJN36909.1|      ------------------M----GRSAYLC-QSEACLEEARR-RKRLQKAL-------RC-K-VPDDVMEAL-QKR----L-LNHHD---ADAEAR----
gb|MCS5691210.1|     ------------DG----M----GRSAYLC-PDPACLEEARR-RKRLQRAL-------RC-Q-VADSIISTL-EQR----L-DGIA----P---------
gb|NBQ18864.1|       ------------QG----M----GRSAYLC-QTQPCLEDARK-RRKLQRAL-------RV-N-VEDTVYITL-AAR----L-GQRQS---AFSATLQAI-
ref|WP_255613922.1|  ------------------M----GRSAYLC-PEENCLEEATR-RKRLQKAL-------RC-Q-VPETVLAVL-KQR----L-IQTPG---ESAEAD----
ref|WP_201324463.1|  ------QIQ-LDHG----M----GRSAYLC-KKLECLQLAQK-KNRLGRSL-------RT-Q-IPPEIFDIL-KSR----L-------------------
ref|WP_011125798.1|  ------------------M----GRSAYLC-PSENCLKEAWQ-RKRLQKAL-------RC-Q-VNVSVLEVL-QNR----L-NHCND---SITKAI----
gb|MBM5803857.1|     ------------QG----Q----GRSAYLC-PTPLCFEEARR-HKRLQRAL-------RV-P-VADSILNRL-ETR----L-SNS-------------A-
ref|WP_094591796.1|  ------------GG----M----GRSAYLC-PSRTCLDEARR-RRKLQKAL-------RC-Q-VADSIYAAL-EQR----L-LSAP----A---------
ref|WP_197212874.1|  ------------RG----M----GRSAYLC-PSRDCLEEARR-RKRLQKGL-------RC-P-VPEAVLDGL-AAR----L-AEAG----G---------
ref|WP_115120514.1|  ------------RG----M----GRSAYLC-PTEACLDEARR-RKRLNKAL-------RC-Q-VPDSVIATL-QER----L-SFQRD---SR--------
gb|MBM5785306.1|     ------------QG----M----GRSAYLC-PRRDCLEEARR-RKKLQKGL-------RC-Q-VADSIYAAL-EQR----L-HATSS---AGSEAR----
ref|WP_038023507.1|  ------------RG----M----GRSAYLC-PTEACLDEARR-RKRLNKAL-------RC-Q-VPDSVITVL-QER----L-SFERN---CR--------
ref|WP_254938812.1|  ------------AG----M----GRSAYLC-PKAACLEEARR-RKRLQRAL-------RC-Q-VADSIYASL-GQR----L-DTNPG-------------
ref|WP_038553465.1|  ------------------M----GRSAYLC-PEDNCLEEATR-RKRLQKAL-------RC-Q-VPETVFAML-KQR----L-NQTTG---ESAEAD----
gb|MDT7945133.1|     ------------EG----M----GRSAYVC-RRLECLQAARR-KQQISRAL-------KA-P-VPDSLFQRL-LDS----L-EQDSQ---GH--------
gb|MEB3326908.1|     ------------------M----GRSAYLC-PKQECLEEAKR-RKRLQRAL-------RC-Q-VADSIVASL-ERR----L-T-----------------
ref|WP_130129912.1|  ------------------M----GRSAYLC-PKETCLEEARR-RKRLQKAL-------RC-Q-VPDSVVEVL-QER----L-SLHQG---TAAEAR----
ref|WP_038001092.1|  ------------------M----GRSAYLC-PKETCLEEARR-RKRLQKAL-------RC-Q-VPDNVVEVL-QER----L-SLHQG---TAAEAR----
ref|WP_258040662.1|  QN----QIQ-LDHG----M----GRSAYLC-QKLECLQLAQK-KNRLGRSL-------RT-H-IPPEIFDIL-KSR----L-------------------
gb|PSP26132.1|       ------------SG----V----GRSAYLC-P-NSSCPNFHR-TTAVRN---------------------------------------------------
```

```
gb|MEB3304480.1|     ------------RG----M----GRSAYLC-PEVDCLEEARR-KRRLQRAL-------RC-A-VDDTILKSL-EQR----L-PEL--PPLRQDDQWPQL-
gb|NDC16240.1|       ------------QG----M----GRSAYLC-PQQSCLEEARK-RRKLQRAL-------RV-P-VADAIYEHL-AER----L-GL----------------
tpg|HBC43141.1|      -----FQIQ-LDDG----M----GRSAYLC-KKLECLQIAQK-KNRLGRSL-------RT-Q-IPSEIFDIL-KSR----L-------------------
ref|WP_206339282.1|  ------------------M----GRSAYLC-PEGNCLEEATR-RKRLQKAL-------RC-Q-VPETVLAVL-KQR----L-IQTPG---ESAEAD----
ref|WP_063406354.1|  ------------------M----GRSAYLC-PNEACLEEALR-RKRLQKAL-------RC-Q-VPNNVVEVL-QKR----L-NHSFD---SAAEAK----
gb|NJM75400.1|       ------------RG----M----GRSAYLC-WSFDCIHQAQK-KKRLERAL-------RS-P-VSPDLYQVL-EKN----V-RERLG-------------
ref|WP_259734545.1|  ------------------M----GRSAYLC-PNQECLEEAKR-RKRLQRAL-------RC-Q-VADSIVASL-ERR----L-T-----------------
gb|MBF2058462.1|     ------------KG----E----GRSAYVC-PNSDCINIARK-KKRLGRCL-------RA-P-VSPAIYDDL-QHI----L-D--K--------------
ref|WP_197151498.1|  ------------------M----GRSAYLC-PNQECLEEAKR-RKRLQRAL-------RC-Q-VADSIVASL-ERR----L-T-----------------
ref|WP_254928579.1|  ------------------M----GRSAYLC-PSQVCLDEAKR-RKRLQRAL-------RC-Q-VADSIVETL-ERR----L-T-----------------
ref|WP_323259706.1|  QH----QIQ-LDHG----M----GRSAYIC-QKLDCIQIAQK-KNRLGRSL-------RT-H-IPPEIFDIL-KSR----L-------------------
gb|MBD2423618.1|     ------------------M----GRSAYLC-PSQVCLDEAKR-RKRLQRAL-------RC-Q-VADSIVEAL-ERR----L-T-----------------
gb|MBM5795377.1|     ------------QG----M----GRSAYLC-PRRDCLEEARR-RKKLQKGL-------RC-Q-VADSIYAAL-EQR----L-HATSS---AGSEAR----
ref|WP_281008927.1|  QH----QIQ-LDHG----M----GRSAYIC-QKLDCIQIAQK-KNRLGRSL-------RT-H-IPPEIFDIL-KSR----L-------------------
gb|MCH2566529.1|     ------------------M----GRSAYLC-PSEACLEEALR-RKRLQKAL-------RC-Q-VPNSVVEVL-QKR----L-NHSFD---SAAEAK----
gb|MEB3307264.1|     ------------------M----GRSAYLC-CSHRCLDEARR-RKRLQRAL-------RC-Q-VADHVLDML-ERR----L-P-----------------
tpg|HBH73361.1|      ------------QG----M----GRSAYLC-PSGGCLDDARR-KKRLQRAL-------RC-Q-VADSIVDAL-ELR----L-RTSA----A---------
ref|WP_255146391.1|  ------------EG----M----GRSAYLC-PTRACFDEAKR-RKRLQRAL-------RC-T-VAESTYAAL-EQR----L-NRAP----A---------
ref|WP_271253996.1|  KQ---FQIQ-LDDG----M----GRSAYLC-KKLDCLQIAQK-KNRLGRSL-------RT-H-IPPEIFDIL-KAR----LS------------------
gb|MEB3234896.1|     ------------QG----Q----GRSAYLC-PTAACLEDARK-HKRLQRGL-------RA-Q-VADSILKRL-ESR----L-AGN-------------A-
gb|MDA0886513.1|     ------------------M----GRSAYLC-PNQECLEEAKR-RKRLQRAL-------RC-Q-VADSIVASL-ERR----L-T-----------------
ref|WP_115019618.1|  ------------RG----M----GRSAYLC-PTEACLDEARR-RKRLNKAL-------RC-Q-VPDSVIATL-QER----L-SFQRD---SR--------
ref|WP_011130895.1|  ------------------M----GRSAYLC-PNEACLEEALR-RKRLQKAL-------RC-Q-VPNSVVEVL-QKR----L-NHSFD---SAAEAK----
gb|NDG24387.1|       ------------QG----M----GRSAYVC-RNHTCLEEARR-RKKLQKAL-------RC-Q-VADSIYAAL-EQR----L-RTSP----S---------
gb|MBM5797070.1|     ------------QG----Q----GRSAYLC-PTAACLDDARR-HKRLQRAL-------RG-Q-VADSILKRL-ESR----L-SGN-------------A-
gb|MCA6523574.1|     -----FQIQ-LDDG----M----GRSAYLC-KKLECLQIAQK-KNRLGRSL-------RT-Q-IPSEIFDIL-KSR----L-------------------
gb|MCX5946264.1|     ------------AG----M----GRSAYLC-PRQDCLSEAKR-RRKLQRAL-------RC-Q-VADSIYAAL-EGR----L-------------------
gb|MAV11321.1|       ------------AG----M----GRSAYLC-KQEECLEEARR-RKRLQKAL-------RC-Q-VPDALLATL-QER----L-SRKT--------------
ref|WP_320673936.1|  ------------------M----GRSAYLC-PCEDCLEEARR-RKRLQKAL-------RC-Q-VPESVFEVL-QKR----L-SQSID---EAAEAI----
gb|MCP9817453.1|     ------------GG----M----GRSAYLC-PSRTCLEEARR-RRKLQKAL-------RC-Q-VADSIYAAL-EQR----L-LRAP----A---------
gb|NJK35889.1|       ------------SG----M----GRSAYVC-RCLECLTLAQK-KQRLGRSL-------RA-Q-ISPDLFAEL-HRR----L-SIAPP---EGLG-----P
gb|NJL99010.1|       QPVGKPLIQ-LDQG----Q----GRSAYLC-PQISCLVDAQK-RKRLERSL-------RT-Q-VPAHIYDTL-ASR----L-LP---------------A
ref|WP_193800382.1|  ------------QG----M----GRSAYVC-PRLECITIAQK-KKRLGRTL-------RT-T-MPPEIYDQL-KSR----C-------------------
ref|WP_063403536.1|  ------------------M----GRSAYIC-PSEACLEEALR-RKRLQKAL-------RC-Q-VPNSVVEVL-QKR----L-NHSFD---SAAEAK----
gb|RMD73144.1|       ------------------M----GRSAYIC-KNVDCVAIAEK-K-RLAKAL-------KT-S-IPSDIYEQL-WQK----L-KETKT-------------
gb|MCY7333721.1|     -----FQIQ-LDHG----M----GRSAYIC-QKLECLQLAQK-KNRLGRSL-------RT-Q-IPLKVFDIL-KSR----L-------------------
gb|MDG2329318.1|     ------------------M----GRSAYLC-PKEECLEEARR-RKRLQKAL-------RC-Q-VPDAVLTTL-NER----L-SASTG---ESAEAN----
gb|MCX5941757.1|     ------------AG----M----GRSAYLC-PRQDCLSEAKR-RRKLQRAL-------RC-Q-VADSIYAAL-EGR----L-------------------
tpg|HJN33659.1|      ------------------M----GRSAYLC-PNEACLEEALR-RKRLQKAL-------RC-Q-VPKSVVEVL-QKR----L-NHSFD---SAAEAK----
tpg|HIK19498.1|      ------------EG----M----GRSAYVC-RRLECLQAARR-KQQINRAL-------KA-A-VPDSLFQRL-LDS----L-EQDPQ---GH--------
tpg|HCX54226.1|      ------------TG----M----GRSAYLC-PTEDCLEEARR-RKRLQKAL-------RC-Q-VP----------------------------------
ref|WP_063399518.1|  ------------------M----GRSAYLC-PSEACLEEALR-RKRLQKAL-------RC-Q-VPISVVEVL-QKR----L-NHSFD---SAAEAK----
ref|WP_011359406.1|  ------------------M----GRSAYLC-PKEECLEEARR-RKRLQKAL-------RC-Q-VPDAVLTTL-NGR----L-SASTG---ESAEAN----
gb|MEB3185113.1|     ------------QG----M----GRSAYLC-RNRSCLDDARR-KKKLQRAL-------RC-Q-VADSIVEAL-EQR----L-ANGG----A---------
ref|WP_186470267.1|  -------------G----V----GRSAYLC-PKESCLEEAQR-RKRLPKAL-------RC-Q-VPDSVLEEL-RER----L-ISDTE---SDAEAR----
ref|WP_009628060.1|  QH---FQIQ-LDQG----M----GRSAYLC-QKLDCLQIAQK-KNRLGRSL-------RT-H-IPPEIFDIL-KSR----LELINKNHQTQ---------
gb|MBE67324.1|       ------------------M----GRSAYLC-PKEECLEEARR-RKRLQKAL-------RC-Q-VPDAVLTTL-NER----L-SASTG---ESAEAN----
ref|WP_272159772.1|  ------------------M----GRSAYLC-PKEECLEEARR-RKRLQKAL-------RC-Q-VPDAVLTTL-NGR----L-SASTG---ESAEAN----
ref|WP_063419582.1|  ------------------M----GRSAYLC-PNEGCLEEALR-RKRLQKAL-------RC-Q-VPNSVVEVL-QKR----L-NHSFD---SAAEAK----
gb|MEB3240836.1|     ------------QG----M----GRSAYLC-PEQKCFDEARK-RRKLQRAL-------RV-P-VADSVYDAL-ATR----L-T--PD---ARKALRQDKD
gb|NBW63894.1|       ------------RG----M----GRSAYVC-PNPVCVEEARR-RRKLQKAL-------RC-Q-VADSIIAAL-EQR----L-QSTP----A---------
gb|MBM5788852.1|     ------------AG----M----GRSAYLC-RDHACLEEARR-RKKLQKAL-------RC-Q-VADSIYAAL-EQR----L-HTSP----S---------
gb|MEB3262778.1|     ------------RG----M----GRSAYLC-ANRTCLDDARR-KKKLQRAL-------RC-Q-VADSIVEAL-ERR----L-ASGG----A---------
```

```
ref|WP_296365315.1|   ------------KG----M----GRSAYLC-PSQSCLEEAKR-RKKLQKAL-------RC-Q-VADSIYAAL-EQR----L-HATSS---AGSEAR----
gb|PZU97808.1|        KGGQSFQIQ-LDHG----M----GRSAYLC-KSLECLQLAQK-KNRLGRSL-------RT-Q-VPVEILEVL-KLR----LQTQNVNPPV----------
gb|MAN18293.1|        -------------G----M----GRSAYLC-PKESCLEEAQR-RKRLPKAL-------RC-Q-VPDSVLEEL-RER----L-ISDTE---SDAEAR----
ref|WP_186583364.1|   ------------------M----GRSAYLC-PQESCLEEAQR-RKRLQKAL-------RC-Q-VPDSVMATL-KQR----L-FPDKE---TVAEAR----
gb|MCS5699361.1|      ------------EG----M----GRSAYLC-REPACLEEARR-RKRLQRAL-------RC-Q-VADSIISTL-EQR----I-EGIA----P---------
ref|WP_186570491.1|   ------------------M----GRSAYLC-PEENCLEEATR-RKRLQKAL-------RC-Q-VPETVLAVL-KQR----L-NQTNG---ESAEAD----
ref|WP_186501173.1|   -------------G----M----GRSAYLC-PKESCLEEAQR-RKRLPKAL-------RC-Q-VPDSVLEEL-RER----L-ISDTE---SDAEAR----
gb|MBL6794407.1|      ------------------M----GRSAYLC-PKEECLEEARR-RKRLQKAL-------RC-Q-VPDAVLTTL-NER----L-SASTG---ESAEAK----
gb|MBL6880295.1|      ------------------M----GRSAYLC-PKEECLEEARR-RKRLQKAL-------RC-Q-VPDAVLTTL-NER----L-SASTG---ESAEAN----
gb|MDB4653764.1|      ------------------M----GRSAYLC-PKEECLEEARR-RKRLQKAL-------RC-Q-VPDAVLTTL-NER----L-SASTG---VSAEAN----
gb|RPF82291.1|        ------------------M----GRSAYLC-PKEACLEDARR-RKRLQKAL-------RC-Q-VPDAVLSTM-QKR----L-SSTTG---ESAEAN----
gb|MCH1457301.1|      ------------------M----GRSAYLC-PKEECLEEARR-RKRLQKAL-------RC-Q-VPDAVLTTL-NKR----L-SASTG---VSAEAN----
gb|MED5164851.1|      ------------------M----GRSAYLC-PNEACLEEALR-RKRLQKAL-------RC-Q-VPNSVVEVL-QKR----L-NHSFD---SAAEAK----
gb|PZO43371.1|        KQ---FQIQ-LDDG----M----GRSAYVC-KKLDCLQLAQK-KNRLGRSL-------RT-S-IPAAIFDIL-KSR----L-------------------
ref|WP_028952666.1|   ------------------M----GRSAYLC-PKEACLEDARR-RKRLQKAL-------RC-Q-VPDAVLATM-QKR----L-SSTTR---ESAEAN----
gb|MAI96565.1|        ------------------M----GRSAYLC-VEQNCLEEATR-RKRLQKAL-------RC-Q-VPESIFVAL-KQR----L-NSAMG---ESAEAD----
ref|WP_037988505.1|   ------------------M----GRSAYLC-PKEECLEEARR-RKRLQKAL-------RC-Q-VPDAVLTTL-NER----L-SASTG---VSAEAN----
ref|WP_029553103.1|   ------------QG----M----GRSAYLC-PKRDCLEEARR-RKKLQKGL-------RC-Q-VADSIYAAL-EQR----L-HATSS---AGSEAR----
ref|WP_106502372.1|   ------------------M----GRSAYLC-PNQACLEEAKR-RKRLQRAL-------RC-Q-VADAIVETL-EKR----L-T-----------------
dbj|GDX71743.1|       ------------QG----M----GRSAYLC-RDHTCLEEARR-RKKLQKAL-------RC-Q-VADSIYAAL-EQR----L-QASP----S---------
ref|WP_036911320.1|   ------------------M----GRSAYLC-PNEACLEEALR-RKRLQKTL-------RC-Q-VPKSVVEVL-QKR----L-NHSFD---SAAEAK----
ref|WP_011825091.1|   ------------------M----GRSAYLC-PNEACLEEALR-RKRLQKAL-------RC-Q-VPNSVVEVL-QKR----L-NHSFD---SAAEAK----
gb|MBL6803472.1|      ------------------M----GRSAYLC-QREECLEEARR-RKRLQKAL-------RC-Q-VPDEVLAAL-EQR----L-RPATE---VTAEAN----
ref|WP_048017238.1|   ------------------M----GRSAYLC-PSPACLDEARR-RKRLQRAL-------RC-P-VPDAILDAL-DRR----L-S-----------------
gb|MAV12819.1|        ------------------M----GRSAYLC-QREECLEEARR-RKRLQKAL-------RC-Q-VPDEVLAAL-EQR----L-RPATE---VSAEAN----
ref|WP_115010044.1|   ------------------M----GRSAYLC-REENCLEEATR-RKRLQKAL-------RC-P-VPETVLAVL-KQR----L-NQSVG---ESAEAD----
ref|WP_006851947.1|   ------------------M----GRSAYLC-REENCLEEATR-RKRLQKAL-------RC-Q-VPETVFAVL-KQR----L-NQSVG---ESAEAD----
gb|NDG75793.1|        ------------QG----M----GRSAYVC-QNHVCLDEARR-RKKLQKAL-------RC-Q-VADSFYAAL-EQR----L-PPAPP---AGSEAR----
gb|MEC8441465.1|      ------------------M----GRSAYLC-QREECLEEARR-RKRLQKAL-------RC-Q-VPDDVLAAL-EQR----L-RPSTE---VSAEAN----
ref|WP_069789292.1|   ------------EG----M----GRSAYVC-PTLECITIAQK-KKRLGRTL-------RT-S-VDPEIYDQL-KSR----C-------------------
gb|WRL41094.1|        ------------------M----GRSAYIC-KNVDCVAIAEK-K-RLAKAL-------KT-S-IPSDIYQQL-WQK----L-KETKN---TVK-------
ref|WP_320002208.1|   ------------------M----GRSAYIC-KNVDCVAIAEK-K-RLAKAL-------KT-S-IPSDIYQQL-WQK----L-KETKN---TVK-------
ref|WP_205909688.1|   ------------------M----GRSAYLC-PNEGCLEEALR-RKRLQKAL-------RC-Q-VPNSVVEVL-QKR----L-NHSFD---SAAEAK----
ref|WP_011432759.1|   ------------GG----M----GRSAYVC-RRLECLQAARR-KQQINRAL-------KA-P-VPDSLFQRL-LDS----L-EQDSQ---GH--------
gb|MDA7432491.1|      ------------------M----GRSAYVC-PKEECLEEARR-RKRLQKAL-------RC-Q-VPDAVLTTL-NER----L-SASTG---ESAEAN----
gb|MEB3202259.1|      ------------------M----GRSAYLC-PNQACLEEAKR-RKRLQRAL-------RC-Q-VADSIVAGL-ERR----L-T-----------------
gb|OIP77665.1|        KPRREKSNQHFQVQLDRGM----GRSAYLC-KSWECLQIAQK-KNRLGRSL-------RI-Q-IPAEIWEIL-KSR----L-PT----------------
gb|MEB3159801.1|      ------------RG----M----GRSAYLC-PTEACLDEALR-RKRLNKAL-------RC-Q-VPDSVLAVL-QER----L-SSETN---GR--------
gb|MED5263527.1|      ------------------M----GRSAYLC-TNEACLEEALR-RKRLQKAL-------RC-Q-VPNSVVEVL-QKR----L-NHSFD---SAAEAK----
ref|WP_036919232.1|   ------------------M----GRSAYLC-PNENCLKEAWQ-RKRLQKAL-------RC-Q-VNVSVIEVL-QNR----L-NHCND---SITKAI----
gb|MBM5790812.1|      ------------QG----M----GRSAYLC-PSQRCLEEAKR-RKRLQKAL-------RC-Q-VADSIYAAL-EQR----L-HPTSS---AGAEAR----
gb|MED5384943.1|      ------------------M----GRSAYLC-PEESCLEDARR-RKRLQKAL-------RC-Q-VPETVLSTM-QER----L-SSTTG---ESAEAN----
ref|WP_199310374.1|   QH----QIQ-LDQG----M----GRSAYIC-QKLDCIQIAQK-KNRLGRSL-------RT-H-IPPEIFDIL-KSR----L-------------------
gb|MBD2316621.1|      QH----QIQ-LDQG----M----GRSAYIC-QKLDCIQIAQK-KNRLGRSL-------RT-H-IPPEIFDIL-KSR----L-------------------
ref|WP_038546212.1|   ------------------M----GRSAYLC-PEESCLEDARR-RKRLQKAL-------RC-Q-VPETVLSTM-QER----L-SSTTG---ESAEAD----
ref|WP_322771268.1|   ------------QG----M----GRSAYLC-PSESCLEEAKR-RKKLQKGL-------RC-H-VADSIYAAL-EQR----L-HATSS---AGSEAR----
gb|WVL01204.1|        ------------------M----GRSAYIC-KNVDCVAIAEK-K-RLAKAL-------KT-P-ISSDIYQQL-WQK----L-KEMKN---TVK-------
ref|WP_015218854.1|   ------------------M----GRSAYIC-KNVDCVAIAEK-K-RLAKAL-------KT-S-IPSDIYQQL-WQK----L-KETKN---TVK-------
ref|WP_055075296.1|   QH---FQIQ-LDQG----M----GRSAYLC-QKIDCLQIAQK-KNRLGRSL-------RT-H-IPPEIFDIL-KSR----LELINKNHQTQ---------
gb|MEB3276547.1|      ------------QG----Q----GRSAYLC-PTPACFDDARR-HKRLQRAL-------RG-Q-VADSILKRL-ESR----L-NGV-------------A-
gb|MCB4428669.1|      ------------------M----GRSAYLC-REENCLEEATR-RKRLHKAL-------RC-Q-VPETVLAVL-KQR----L-NQSVG---ESAEAD----
ref|WP_094510206.1|   ------------QG----M----GRSAYLC-PTRACFDEAKR-RKRVQRAL-------RC-A-VAESTYAAL-EQR----L-NRSP----A---------
gb|MDP6171750.1|      ------------------M----GRSAYLC-QNEACLDEAHR-RKRLQKAL-------RC-Q-VPNSVMEVL-QKR----L-NQSFD---SASEAR----
```

```
ref|WP_223805489.1|  QH---FQIQ-LDQG----M----GRSAYLC-QKLDCLQIAQK-KNRLGRSL-------RT-H-IPPDIFDIL-KSR----LELINKNHQTQ---------
ref|WP_011365029.1|  ------------------M----GRSAYLC-REENCLQEATR-RKRLQKAL-------RC-Q-VPETVLAVL-KQR----L-NQSVG---ESAEAD----
ref|WP_214339661.1|  ------------QG----M----GRSAYLC-PSESCLEEAKR-RKKLQKGL-------RC-H-VADSIYAAL-EQR----L-HATSS---AGSEAR----
ref|WP_255099870.1|  ------------QG----M----GRSAYLC-PTRACFDEAKR-RKRVQRAL-------RC-A-VAESTYAAL-EQR----L-NRSP----A---------
gb|TYQ31904.1|       QH---FQIQ-LDQG----M----GRSAYLC-QKLDCLQIAQK-KNRLGRSL-------RT-H-IPPDIFDIL-KSR----LELINKNHQTQ---------
ref|WP_320666922.1|  ------------------M----GRSAYLC-PTETCFKESYR-RKRLQKAL-------RC-Q-VPASIVEML-QKR----L-NHCID---ADIEAI----
gb|MEB3297319.1|     ------------------M----GRSAYLC-RQRSCLEDARK-KKRLQRAL-------RC-Q-IADCFLERL-EQR----L-Q-----------------
ref|WP_198953901.1|  ------------------M----GRSAYLC-PNPVCLEEARR-RKRLQRAL-------RC-Q-VSDSIVETL-QKR----L-T-----------------
ref|WP_186595138.1|  ------------------M----GRSAYLC-PQESCLEEAQR-RKRLQKAL-------RC-Q-VPDSVMATL-KQR----L-FPDKE---TVAEAR----
ref|WP_186493755.1|  ------------------M----GRSAYLC-REENCLEEATR-RKRLQKAL-------RC-Q-VPETVLAVL-KQR----L-NQSVG---ESAEAD----
gb|MEC7393122.1|     ------------------M----GRSAYLC-CEENCLEEATR-RKRLQKAL-------RC-Q-VPETVLAVL-KQR----L-NQSVG---ESAEAD----
gb|QNJ16179.1|       ------------------M----GRSAYLC-QREACLEEARR-RKRLQKAL-------RC-Q-VPDDVLTAL-EQR----L-RDATD---VTAEAN----
gb|QNI91005.1|       ------------------M----GRSAYLC-QREACLEEARR-RKRLQKAL-------RC-Q-VPDDVLAAL-EQR----L-RDATD---VTAEAN----
gb|NJK59609.1|       ------------QGDRRAM----GRAAYVC-PCLPCVAEAQK-KRRLERAL-------KT-P-VPATLWPDL-WAQ----V-QG--------------QV
ref|WP_259728879.1|  ------------QG----M----GRSAYLC-PTRACFDEAKR-RKRLQRAL-------RC-A-VAESTYAAL-EQR----L-NRSP----A---------
gb|AUC59923.1|       ------------EG----M----GRSAYVC-PTLECITIAQK-KKRLGRTL-------RT-S-VDPEIYDQL-KSR----C-------------------
ref|WP_185186942.1|  ------------EG----M----GRSAYLC-PTESCFEEAKR-RKRLQKAL-------RC-Q-VSDSIYAAL-EQR----L-NATS--------------
ref|WP_255599781.1|  ------------------M----GRSAYLC-REENCLEEATR-RKRLHKAL-------RC-Q-VPETVLAVL-KQR----L-NQSVG---ESAEAD----
ref|WP_012196167.1|  ------------EG----M----GRSAYLC-PNEACLEEALR-RKRLQKAL-------RC-Q-VQMSILEVL-QNR----L-NSCN----D---------
ref|WP_115080914.1|  ------------------M----GRSAYLC-REENCLEEATR-RKRLQKAL-------RC-Q-VPETVLAVL-KQR----L-NQSVG---ESAEAD----
gb|MBR75694.1|       ------------------M----GRSAYLC-QREACLEEARR-RKRLQKAL-------RC-Q-VPDDVLTAL-EQR----L-RDATD---VSAEAN----
gb|RNC90972.1|       ------------------M----GRSAYLC-QREACLEEARR-RKRLQKAL-------RC-Q-VPDDVLAAL-EQR----L-RNATD---VTAEAN----
gb|MCB4389873.1|     ------------------M----GRSAYLC-REENCLEEATR-RKRLQKAL-------RC-Q-VPETVLAVL-KQR----L-NQSIG---ESAEAD----
ref|WP_303534944.1|  ------------------M----GRSAYLC-REENCLEEATR-RKRLQKAL-------RC-Q-VPETVLAVL-KQR----L-NQSVG---ESAEAD----
ref|WP_186495900.1|  ------------------M----GRSAYLC-REENCLEEATR-RKRLQKAL-------RC-Q-VPEKVLAVL-KQR----L-NQSVG---ESAEAD----
ref|WP_320676133.1|  ------------------M----GRSAYLC-PNEDCLNEAHR-RKRLQKAL-------RC-Q-VPENIVKVL-KQR----V-NHCND---LPAEAR----
ref|WP_011933969.1|  ------------------M----GRSAYLC-PQESCLEEAQR-RKRLQKAL-------RC-Q-VPDTVMATL-KQR----L-VPDQE---TVAEAR----
ref|WP_011127466.1|  ------------------M----GRSAYLC-QREACLEEARR-RKRLQKAL-------RC-Q-VPDDVLAAL-EQR----L-RNATD---VTAEAN----
gb|MCX5948231.1|     ------------------M----GRSAYLC-PDRACLEEARR-RKRLQRSL-------RC-Q-IADSIIAAL-DAR----L-GTDDA---ATVKA-----
gb|MCS6960746.1|     ------------QG----M----GRSVYVC-RQGSCLQEALK-KNRLSKAL-------RT-P-LPPIVVAEL-RQR----A-ASSAQ---ANSSLSIDNS
ref|WP_217901569.1|  QH----QIQ-LDQG----M----GRSAYVC-QKLECLQIAQK-KNRLGRSL-------RT-H-IPPEIFDIL-KSR----LQNP----------------
gb|OYQ64342.1|       QH----QIQ-LDQG----M----GRSAYVC-QKLECLQIAQK-KNRLGRSL-------RT-H-IPPEIFDIL-KSR----LQNP----------------
ref|WP_190398881.1|  QQ---FQVS-LDEG----M----GRSAYLC-KKLECLQIAQK-KNRLGRSL-------RT-H-IPNEIFDIL-KSR----L-------------------
ref|WP_114989435.1|  ------------------M----GRSAYLC-REENCLEEATR-RKRLQKAL-------RC-Q-VPETVLAVL-KQR----L-NQSVG---ESAEAD----
gb|MCB4407378.1|     ------------------M----GRSAYLC-REENCLEEATR-RKRLQKAL-------RC-Q-VPEAVVAVL-KQR----L-NQSVG---ESAEAD----
ref|WP_186479837.1|  -------------G----M----GRSAYLC-PQESCLEEAQR-RKRLQKAL-------RC-Q-VPDSVLEVL-RER----L-KPDTG---SDAEAR----
ref|WP_247910210.1|  ------------QG----M----GRSAYLC-PTESCFEEAKR-RKRLQKAL-------RC-Q-VSDSIYSAL-EQR----L-NATS--------------
dbj|GCE64220.1|      ------------DG----M----GRSAYLC-PNGPCLEEARR-RKRLQKAL-------RC-Q-VPEDVVALL-QQR----L-NSSRR-------------
ref|WP_099812864.1|  ------------QG----M----GRSAYLC-RRLECLRAARR-KHQMSRAL-------KA-A-VPDSLWQLL-EQQ----L-LRDSE---AQ--------
gb|MBT65681.1|       ------------------M----GRSAYLC-ANEACLEEARR-RKRLQKSL-------RC-Q-VPDDLMTAL-QER----L-TQHRV---ADAEAK----
gb|MEB3183095.1|     ------------AG----M----GRSAYLC-PTVHCLEEARR-RKRLQRSL-------RT-P-VADSIVAGL-QAR----I-EASG----P---------
ref|WP_254995692.1|  ------------------M----GRSAYLC-PNPVCLEEATR-RKRLQRAL-------RC-Q-VADSIVETL-QKR----L-T-----------------
ref|WP_150884534.1|  -------------G----F----GRSAYLC-RSTECLGEAIR-RKRLQRAL-------RC-P-VPQAVIAAL-EER----L-RADAT---ADAEAR----
ref|WP_006043237.1|  ------------------M----GRSAYLC-PQESCLEEAQR-RKRLQKAL-------RC-Q-VPDSVMATL-KQR----L-FSEKE---TVAEAR----
gb|NDC34872.1|       ------------GG----Q----GRSAYLC-RHATCLEAARR-HKRLQRAL-------RC-Q-VAESILKCL-ESR----L-AGND----P---------
ref|WP_255613750.1|  ------------------M----GRSAYLC-REENCLEEATR-RKRLQKAL-------RC-Q-VPETVLAVL-KQR----L-NQSIG---ESAEAD----
gb|OUT75469.1|       ------------EG----M----GRSAYVC-PQNCCLEDARK-RRRLQRAL-------RV-P-VADSVYDAL-AER----L-K--PE---AEKPLRQDED
gb|NCG15537.1|       ------------------M----GRSAYLC-PTEACFEEARR-RKKLQKSL-------RC-Q-VPDDLMTAL-QGR----L-TESRV---AAAEAR----
gb|TGG79053.1|       ------------RG----F----GRSAYLC-PDAACLEDARK-RRRLQRAL-------RC-N-MDDVVYQAL-ESR----L-QHPR----QSCG------
ref|WP_115131857.1|  ------------------M----GRSAYLC-REENCLEEATR-HKRLQKAL-------RC-Q-VPETVLAVL-KQR----L-NQSVG---ESAEAD----
ref|WP_284500670.1|  ------------------M----GRSAYLC-REENCLEEATR-RKRLQKAL-------RC-Q-VPETVLAVL-KQR----L-NQSVG---ESAEAD----
ref|WP_255616060.1|  ------------------M----GRSAYLC-REENCLEEATR-RKRLQKAL-------RC-Q-VPEAVVAVL-KQR----L-NQSVG---ESAEAD----
gb|RCL52291.1|       ------------QG----M----GRSAYLC-PKRCCLEDARK-RRRLQRAL-------RV-P-VADSVYDAL-AQR----L-E--PE---AEKPLRQDED
```

```
tpg|HAN46371.1|       ------------RQEPRTP----GRAAYLC-PCAACLTEAQK-KRRLERAL-------KT-P-VPPTLWPQW-WEQ----L-AALCP-----------QA
ref|WP_186587596.1|   ------------------M----GRSAYLC-RREACLEEARR-RKRLQKAL-------RC-Q-VPDDILAAL-EQR----L-RDSTD---VTAEAN----
ref|WP_011431137.1|   ------------QG----M----GRSAYLC-RRLECLRAARR-KHQMSRVL-------KA-A-VPDSLWQLL-EQQ----L-LRDSE---AQ--------
ref|WP_186498867.1|   ------------------M----GRSAYLC-QREACLEEARR-RKRLQKAL-------RC-Q-VPDDVLAAL-GQR----L-RDATD---VTAEAN----
ref|WP_259720650.1|   ------------RG----M----GRSAYLC-PSRSCLDEAKR-RKRLQRAL-------RT-Q-VADSIIATL-DQR----L-RDDDA-------------
gb|MDP6195789.1|      ------------------I----GRSAYLC-QNEACLDEAHR-HKRLQKAL-------RC-Q-VPKSVVEVL-RKR----L-NHTSE---SAAEAR----
gb|RZO05221.1|        ------------------M----GRSAYLC-RREACLEEARR-RKRLQKAL-------RC-Q-VPDDVLAAL-EQR----L-RDSTD---VTAEAN----
ref|WP_269603352.1|   ------------------M----GRSAYLC-PAESCFEEALK-RKRLQKAL-------RC-A-IDSSIFNML-QKQ----L-NTCIE---SDTEA-----
ref|WP_036900583.1|   ------------KG----M----GRSAYIC-PKKECFEEALR-RKRLQKAL-------RC-Q-VPLTVFDLL-QNR----L-NENK----H---------
ref|WP_048347195.1|   ------------------M----GRSAYLC-PTEACFEEARR-RKRLQKSL-------RC-Q-VPEDVMTAL-QER----L-TEPRV---AAAEAR----
ref|WP_197156911.1|   ------------RG----M----GRSAYLC-PSRSCLDEAKR-RKRLQRAL-------RT-Q-VADSIIATL-NQR----L-RDDDA-------------
gb|MCX5969416.1|      ------------------M----GRSAYLC-RDRACLEEAKR-RKRLQRSL-------RA-Q-IADSIIATL-DAR----L-EGSDA---ATAKA-----
ref|WP_259735584.1|   AT----------GA----M----GRSAYLC-PSSSCIDDARR-RKRLQRSL-------RC-Q-VSDSIYMAL-EER----L-NQ-------------TRA
emb|CAK6691010.1|     ------------RG----M----GRSAYLC-PSRSCLDEAKR-RKRLQRAL-------RT-Q-VADSIIATL-NQR----L-RDDDA-------------
ref|WP_067095672.1|   ------------QG----M----GRSAYLC-RKESCLEEAQR-RKRLHKAL-------RC-Q-VPDSAIEEL-RTR----L-KPNKE---SAAEAR----
ref|WP_269608866.1|   ------------------M----GRSAYLC-PTESCFEEALK-RKRLQKAL-------RC-D-IHSSVFNML-QKE----L-NTCID---SDTEA-----
gb|MBU6251331.1|      ------------GG----E----GRSAYLC-RHATCLESALR-HKRLQRAL-------RC-Q-VAESILKCL-ETR----L-AGDD----P---------
ref|WP_115125622.1|   ------------QG----M----GRSAYLC-RKESCLEEAQR-RKRLHKAL-------RC-Q-VPDSAIEEL-RTR----L-KPDKE---SAAEAR----
gb|MAB54535.1|        ------------------M----GRSAYLC-PRKSCLEETYR-RKRLQKAL-------RC-Q-VPETVLAAL-EQR----L-SQQTE---ESAEAN----
gb|MBM5813821.1|      ------------------M----GRSAYLC-PNPACLEEATR-RKRLQRAL-------RC-Q-VADSIVETL-QKR----L-T-----------------
ref|WP_114994692.1|   ------------QG----M----GRSAYLC-RKESCLEEAQR-RKRLHKAL-------RC-Q-VPDSALEEL-RKR----L-KPNKE---SAAEAR----
gb|MEB3351632.1|      ------------QG----M----GRSAYLC-RNADCLAEARR-RRRFQRSL-------RC-P-VADQLILTL-ETR----L-AIPG----P---------
ref|WP_186538343.1|   ------------QG----M----GRSAYLC-RKESCLEEAQR-RKRLHKAL-------RC-Q-VPDSALEEL-RQR----L-KPNKE---SAAEAR----
ref|WP_038654169.1|   ------------------M----GRSAYLC-PTESCFEEALK-RKRLQKAL-------RC-D-VHSSVFNML-QKQ----L-NTCIY---SDTEA-----
ref|WP_011620279.1|   ------------------M----GRSAYLC-PTEACFEEARR-RKRLQKSL-------RC-Q-VSEELMTAL-QER----L-TEPRV---AAAEAR----
ref|WP_186589231.1|   ------------------M----GRSAYLC-PTEACFEEARR-RKKLQKSL-------RC-Q-VSDDLMTAL-QGR----L-TESRV---AAAEAR----
ref|WP_074162604.1|   ------------QG----I----GRSAYLC-RKESCLEEAQR-RKRLHKAL-------RC-Q-VPDSALEEL-RQR----L-KPHKE---SAAEAR----
gb|MAD68593.1|        ------------QG----M----GRSAYLC-RKESCFEEAQR-RKRLHKAL-------RC-Q-VPDSALEEL-RKR----L-NPNKE---SAAEAR----
ref|WP_110861757.1|   ------------------M----GRSAYLC-PTESCFEEALK-RKRLQKSL-------RC-D-IHSSVFNML-QKQ----L-NTCID---SDTEA-----
ref|WP_322782688.1|   VT----------GA----M----GRSAYVC-PSSSCIEEARR-RKRLQRSL-------RC-Q-VSDSIYRAL-EER----L-SE-------------SLA
ref|WP_197162564.1|   VT----------GA----M----GRSAYVC-PSSSCIEEARR-RKRLQRSL-------RC-Q-VSDSIYRAL-EER----L-SE-------------SLA
gb|MAF41027.1|        ------------QG----T----GRSAYLC-RRESCLEEAQR-RKRLHKAL-------RC-Q-VPDSAIEEL-RRR----L-KPNKE---SAAEAR----
ref|WP_186523834.1|   ------------------M----GRSAYLC-PTEACFEEARR-RKKLQKSL-------RC-Q-VSDDLMTAL-QER----L-TEPRV---AAAEAR----
gb|PZO47380.1|        ------------EG----M----GRSAYLC-PQASCLTST------------------------------------------------------------
gb|MEB3350462.1|      ------------------M----GRSAYLC-PDADCLAEAKR-RRRLQRSL-------RC-P-VEETLLITL-EKR----L-A-----------------
ref|WP_254968188.1|   ------------QG----M----GRSAYLC-PDPACLEEARR-RRRLQRAL-------RC-P-VDDGLLDQL-EQR----L-------------------
ref|WP_011295369.1|   ------------------M----GRSAYLC-PTESCFEEALK-RKRLQKSL-------RC-D-IHSSVFNML-QKQ----L-NTCID---SDTEA-----
ref|WP_029626136.1|   ------------QG----M----GRSAYLC-PTESCFEEAKR-RKRLQKAL-------RC-Q-VSDSIYAAL-AQR----L-NATSY-------------
gb|MAK15572.1|        ------------------M----GRSAYLC-PTEACFEEARR-RKRLQXSL-------RC-Q-VSDDLMTAL-QER----L-TEXRV---AAAEAR----
ref|WP_186490350.1|   ------------QG----M----GRSAYLC-RKESCFEEAQR-RKRLHKAL-------RC-Q-VPDSALEEL-RKR----L-NPNKE---SAAEAR----
gb|MEB3169508.1|      ------------------M----GRSAYLC-PSEACLTEARR-RRRLQRSL-------RC-P-VDETVYRQL-EQR----L-P-----------------
ref|WP_255141823.1|   ST----------RA----M----GRSAYVC-PSSSCIEDARR-RKRLQRSL-------RC-Q-VSDSIYTAL-EER----L-SE-------------SLA
ref|WP_036905622.1|   ------------------M----GRSAYLC-PTESCFEEALK-RKRLQKSL-------RC-D-IHSSVFNML-QKQ----L-NTCID---SDTEA-----
gb|EAQ76225.1|        ST----------RA----M----GRSAYIC-PSSSCIEDARR-RKRLQRSL-------RC-Q-VSDSIYTAL-EER----L-SE-------------SLA
ref|WP_011824427.1|   ------------------M----GRSAYLC-PTESCFQEALK-RKRLQKSL-------RC-D-IHSSVFNML-QKQ----L-NTCID---SDTEA-----
gb|MDP7995305.1|      ------------------M----GRSAYLC-PTEACFEEARR-RKRLQKSL-------RC-Q-VSDDLMTAL-QER----L-TEPRV---AAAEAR----
gb|MCX5957055.1|      ------------------M----GRSAYLC-PSPTCLDEARR-RKRLQRAL-------RC-Q-VSDSIFATL-EAR----L-P-----------------
ref|WP_037979888.1|   ST----------RA----M----GRSAYIC-PSSSCIEDARR-RKRLQRSL-------RC-Q-VSDSIYTAL-EER----L-SE-------------SLA
ref|WP_106220260.1|   ------------GG----M----GRSAYLC-PQPSCLEEARR-RRRLQRGL-------RC-A-VSDTILASL-EAR----L-ERSPPRPLRQDEPWSPR-
ref|WP_006854377.1|   ------------------M----GRSAYLC-PTEACFEEARR-RKRLQKSL-------RC-Q-VSDDLLTAL-QER----L-TEPRV---AAAEAR----
ref|WP_228007088.1|   ------------QG----M----GRSAYLC-PCSDCLEEARR-RRRLQRSL-------RC-Q-GADALLDVL-RER----L-DATP-----QPAPG----
ref|WP_271488669.1|   ------------------M----GRSAYLC-PTEACFEEARR-RKRLQKSL-------RC-Q-VSEGLMTAL-KER----L-TEPRV---AAAEAR----
ref|WP_186516396.1|   ------------------M----GRSAYLC-PTEACFEEARR-RKRLQKSL-------RC-Q-VSEGLMTAL-KER----L-TEPRV---AAAEAR----
```

```
ref|WP_322775930.1|  ST----------RA----M----GRSAYVC-PSSSCIEDARR-RKRLQRSL-------RC-Q-VSDSIYTAL-EER----L-SE-------------SLA
ref|WP_257473614.1|  ------------------M----GRSAYLC-PTESCFEEALK-RKRLQKSL-------RC-D-IHSSVFNML-QKQ----L-NTCID---SDTEA-----
gb|QNI69741.1|       ------------QG----M----GRSAYLC-PCRDCLEEARR-RRRLQRSL-------RC-Q-GADALLDVL-RER----L-DATP-----QPAPG----
gb|MDA7433454.1|     ------------------M----GRSAYLC-PTEACFEEARR-RKRLQKSL-------RC-Q-VSEAVMTAL-QER----L-TETRV---AAAEAR----
ref|WP_269623649.1|  ------------------M----GRSAYLC-PTESCFEEALK-RKRLQKAL-------RS-D-IHSSVFKML-QKQ----L-NACID---SDTEA-----
gb|MAR07732.1|       ------------------M----GRSAYLC-TKETCMEEALR-RKRLQKAL-------RC-Q-VPDSILAEL-QER----V-CREAD---ESAEAN----
ref|WP_254954616.1|  ------------RG----M----GRSAYLC-PEPSCLEEARR-RRRLQRGL-------RC-A-VSDAIIVSL-EAR----L-ERSAPQPLRQDESWSQR-
ref|WP_286160945.1|  ------------GG----M----GRSAYLC-PQPSCLEEARR-RRRLQRGL-------RC-A-VSDTILASL-EAR----L-ERSPPRPLRQDEPWSPR-
ref|WP_269611454.1|  ------------------M----GRSAYLC-PTESCFEEALK-RKRLQKAL-------RC-D-IHSSVFNML-QKQ----L-NTCID---SDSEA-----
gb|MEB3173427.1|     ------------GG----Q----GRSAYLC-PTQACFDEARR-HKRLQRAL-------RV-P-VADSILNRL-ESR----L-SGS-------------A-
ref|WP_225875770.1|  ------------QG----M----GRSAYLC-PCRDCLEEARR-RRRLQRSL-------RC-Q-GADALLDVL-RER----L-DATP-----QPAPG----
gb|MBD2718795.1|     ------------GG----M----GRSAYLC-PQPSCLEEARR-RRRLQRGL-------RC-A-VSEAILASL-EAR----L-ERSAPRPLRQDEPWSPR-
ref|WP_254992200.1|  ------------QG----M----GRSAYLC-PEPSCLEEARR-RRRLQRGL-------RC-A-VSEAILASL-EAR----L-ERSAPRPLRQDEPWSPR-
gb|TVS05339.1|       ------------RG----M----GRSAYLC-PSPACLQEARR-RRRLQRAL-------RC-P-VEDSLLDQL-EQR----L-L---------AG------
gb|QNI50382.1|       ------------------M----GRSAYLC-QREACLEEARR-RKRLQKAL-------RC-Q-VPDDVLAAL-EQR----L-RNATD---VTAEAD----
gb|QBE69991.1|       ------------------M----GRSAYLC-PKETCLEEARR-RKRLQKAL-------RC-Q-VPDSVVEVL-QER----L-SLHQG---TAAEAR----
gb|MBM5821121.1|     ------------GG----M----GRSAYLC-PQPSCLEEARR-RRRLQRGL-------RC-A-VSEAILASL-EAR----L-ERSAPRPLRQDEPWSPR-
gb|EAQ68383.1|       ------------------M----GRSAYLC-PKETCLEEARR-RKRLQKAL-------RC-Q-VPDNVVEVL-QER----L-SLHQG---TAAEAR----
ref|WP_254963331.1|  ------------RG----M----GRSAYLC-PEPSCVEEARR-RRRLQRGL-------RC-A-VSDAIIASL-EAR----L-ERSAPQPLRQDEPWSPR-
ref|WP_186544284.1|  ------------------M----GRSAYLC-PTEACFEEARR-RKRLQKSL-------RC-Q-VSDDLLTAL-QER----L-TEPRV---AAAEAR----
gb|MEB3171165.1|     ------------QG----M----GRSAYLC-PTLPCLEDTRR-RRRLQRAL-------RC-P-VPDSVLTAL-ADR----L-EPGTP---AGAEAR----
gb|MBW4530560.1|     ------------RG----M----GRSAYLC-PEPSCLEEARR-RRRLQRGL-------RC-A-VSDAIIASL-EAR----L-ERSAPQPLRQDEPWSPR-
ref|WP_254957545.1|  ------------RG----M----GRSAYLC-PEPSCLEEARR-RRRLQRGL-------RC-A-VSDAIIASL-EAR----L-ERSAPRPLRQDEPWSPR-
gb|EAU74800.1|       ------------RG----M----GRSAYLC-PTEACLDEARR-RKRLNKAL-------RC-Q-VPDSVITVL-QER----L-SFERN---CR--------
ref|WP_015110535.1|  ------------GG----M----GRSAYLC-PQPSCLEEARR-RRRLQRGL-------RC-A-VSETILAIL-EAR----L-ERSAPRPLRQDEPWSPR-
gb|QNJ13310.1|       ------------------M----GRSAYLC-QREACLEEARR-RKRLQKAL-------RC-Q-VPDDVLAAL-EQR----L-RNATD---VTAEAN----
ref|WP_254944301.1|  ------------RG----M----GRSAYLC-PDPSCLEEARR-RRRLQRGL-------RC-A-VSDAIIASL-EAR----L-ERSAPQPLRQDESWSPR-
gb|MBD2549645.1|     ------------AG----M----GRSAYLC-PQSSCLEDARR-RRRLQRGL-------RC-A-VSEAILASL-EAR----L-ERSAPRPLRQDEPWSPR-
gb|MCT0206289.1|     ------------GG----M----GRSAYLC-PQPSCLEEARR-RRRLQRGL-------RC-A-VSDAIIASL-EAR----L-ERSAPQPLRQDEPWSPR-
ref|WP_159820065.1|  ------------RG----M----GRSAYLC-PQPSCLEEARR-RRRLQRGL-------RC-A-VSDTILASL-EAR----L-ERSPPRPLRQDEPWSPR-
ref|WP_286194360.1|  ------------RG----M----GRSAYLC-PEPSCLEEARR-RRRLQRGL-------RC-A-VSDAIIASL-EAR----L-ERSAPQPLRQDEPWSPR-
ref|WP_254980322.1|  ------------RG----M----GRSAYLC-PNPACLEEARR-RRRLQRAL-------RC-P-VDDTLLDQL-EQR----L-------------------
ref|WP_158467192.1|  ------------------M----GRSAYLC-PTESCFEVALK-RKRLQKAL-------RC-E-IHSSFFNML-QKQ----L-NTCID---SDTEA-----
gb|QVL53487.1|       ------------QG----M----GRSAYVC-PQLSCLEETRR-RRRLQRAL-------RC-P-VDESLVETL-MQR----L-SNEAL-------------
gb|KAF0652084.1|     ------------RG----M----GRSAYLC-PQPSCLEEARR-RRRLQRGL-------RC-A-VSDTILASL-EAR----L-ERSPPRPLRQDEPWSPR-
ref|WP_255092431.1|  ET----------RA----M----GRSAYVC-PSSSCIEDARR-RKRLQRSL-------RC-Q-VSDSIYTAL-EER----L-SE-------------PLH
gb|MEB3361199.1|     ------------------M----GRSAYLC-PSEACLQDARR-RRRLQRAL-------RC-P-VEEAVYRQL-EER----L-H-----------------
ref|WP_198949670.1|  KT----------RA----M----GRSAYVC-PTTTCIEDARR-RKRLQRSL-------RC-Q-VSDSIYTAL-EER----L-RE-------------PLH
ref|WP_323356953.1|  ------------QG----M----GRSAYLC-PEPSCLEEARR-RRRLQRGL-------RC-A-VSDAIIASL-EAR----L-ERSAPQPLRQDEPWSPH-
gb|MEB3266211.1|     ------------------M----GRSAYLC-PDPACLADAKR-RRRLQRSL-------RC-P-VNDAVLITL-EKR----L-D-----------------
ref|WP_255103146.1|  DT----------RA----M----GRSAYVC-PSSSCIEEAKR-RKRLQRSL-------RC-Q-VSDSIYTAL-EER----L-SE-------------PLH
ref|WP_255110457.1|  TT----------KA----M----GRSAYVC-PSSRCIEDARR-RKRLQRAL-------RC-Q-VSDSIYAAL-VER----L-SE-------------SHA
ref|WP_257473432.1|  ------------------M----GRSAYLC-PTESCFEVALK-RKRLQKAL-------RC-E-IHSSFFNML-QKQ----L-NTCID---SDTEA-----
gb|MBM5793042.1|     ------------QG----M----GRSAYLC-PRRDCLEEARR-RKKLQKGL-------RC-Q--------------------------------------
gb|MCX5932016.1|     TT----------RA----M----GRSAYVC-PSSRCIEDARR-RKRLQRAL-------RC-Q-VSDSIYAAL-VER----L-SE-------------SHA
gb|MEB3351395.1|     ------------RG----M----GRSAYLC-PQPDCLNEARR-RRRLQRAL-------RC-A-VDDTLLNTL-EQR----L-PRQ--HALRQDDQWPQP-
gb|MBE9154035.1|     ------------QG----M----GRSAYLC-PCSDCLEEARR-RRRLQRSL-------RC-Q-GADALLDVL-RER----L-DATP-----QPAPG----
gb|MBM5825515.1|     ------------RG----M----GRSAYLC-PRHSCVEEARR-RRRLQRAL-------RC-P-VTDAILDAL-EQR----L-EAGLT-------------
gb|MCU0530038.1|     ------------QG----M----GRSAYLC-PHPACLDEARR-RRRLQRAL-------RC-A-LDDALLDEL-GQR----L-------------------
gb|MEB3156001.1|     ------------AG----M----GRSAYLC-PDPACLEDARR-RRRLQRSL-------RC-A-VEETIFNTL-EGR----L-PRR--HPLRQDDQWPQP-
gb|MBM5816260.1|     ------------AG----M----GRSAYLC-PDPACLEDARR-RRRLQRSL-------RC-A-VEETIFNTL-EGR----L-PRR--HPLRQDDKWPQP-
gb|MCF8131934.1|     ------------SG----M----GRSAYVC-RRSACLDEARR-RRRLQRAL-------RV-P-VPDAILDLL-VER----L-SLEGD---PPLRQDEYRP
gb|MDM7936623.1|     TT----------RA----M----GRSAYVC-PSSRCIEDARR-RKRLQRAL-------RC-Q-VSDSIYAAL-VER----L-SE-------------SHA
```

```
gb|MEB3260791.1|     ------------AG----M----GRSAYLC-PEPACLEDARR-RRRLQRSL-------RC-A-VEETIFTTL-EGR----L-PPQR--------------
gb|MEB3353798.1|     ------------QG----M----GRSAYLC-PQHSCLEEARR-RRRLQRAL-------RC-P-VDESILERL-AQR----L-GA----------------
gb|MEB3334793.1|     ------------RG----M----GRSAYLC-PHPDCFNEARR-RRRLQRSL-------RC-A-VDDGILNTL-ERR----L-PRL--HPLRQDEQWPQL-
tpg|HYP03616.1|      ------------QG----M----GRSAYIC-PSRECLEEARR-RRRLPRAL-------RC-P-IPENILASL-EAR----L-SG-DA---PASH------
ref|WP_221629786.1|  SP----------RA----M----GRSAYVC-PSPSCIEEARR-RKRLQRSL-------RC-P-VSDSIYTAL-DQR----L-RE-------------SRA
gb|MBC1260749.1|     SP----------RA----M----GRSAYVC-PSPSCIEEARR-RKRLQRSL-------RC-P-VSDSIYTAL-DQR----L-RE-------------SRA
gb|MEB3176206.1|     ------------QG----M----GRSAYLC-PEQNCFDEARK-RRKLQRAL-------RV-P-VADSVYDAL-ATR----L-E--PD---AGKALRQDKD
ref|WP_259738767.1|  SP----------RA----M----GRSAYVC-PSPSCIEEARR-RKRLQRSL-------RC-P-VSDSIYTAL-DQR----L-HE-------------SRA
gb|MEB3195012.1|     ------------RG----M----GRSAYLC-PEPACLDEARR-RRRLQRSL-------RC-A-VDDTILHTL-ETR----L-PAL--HALRQDDQWPQP-
gb|MEB3257996.1|     ------------RG----M----GRSAYLC-PQPDCLDEARR-RRRLQRAL-------RC-A-VDDTILNTL-ELR----L-PRL--HPLRQDDQWPQL-
gb|MBO6974532.1|     -------------G----T----GRSAYIC-KSKKCYADSKI-KKKLQKAL-------KT-F-FEPEFFDIF-EKE----I-TSYND---NPNKGI----
gb|MEB3257676.1|     ------------RG----M----GRSAYLC-PQPACLEEARR-RRRLQRSL-------RC-A-VDDSILHTL-EER----L-PRL--HPLRQDDQWPQL-
gb|MEB3165236.1|     ------------------M----GRSAYLC-PDPACLEEARR-RRRLQRAL-------RC-Q-VADSIISTL-EER----L-LSDPS---AGVEA-----
gb|MEB3349663.1|     ------------AG----M----GRSAYLC-RQQSCLDEARR-RRRLQRAL-------RC-P-VDDGVLVTL-EQR----L-ANDGP-------------
ref|WP_209040626.1|  -------------G----T----GRSAYIC-KSNKCYSDSKI-KKKLQKAL-------KT-S-LEPEFIEIF-EKE----I-TSYNN---NPNKGI----
gb|MCH9714882.1|     ------------QG----M----GRSAYLC-RQASCLEEARR-RRRLQRAL-------RC-P-VSEAVLDAL-EQR----L-TNAAQ-------------
ref|WP_245156268.1|  -------------G----T----GRSAYIC-KSNKCYSDSKI-KKKLQKAL-------KT-S-LEPEFIEIF-EKE----I-TSYNN---NPNKGI----
gb|MDX1977370.1|     DR----------DL----M----GRSAYVC-RHQTCLDLARK-KKAWGRSL-------KT-A-VDQELLQQL-HQL----Q-AQL-------------H-
gb|QNI77496.1|       ------------------M----GRSAYLC-PTEACFEEARR-RKKLQKSL-------RC-Q-VSDDLMTAL-QGR----L-TESRV---AAAEAR----
gb|EAU72346.1|       ------------------M----GRSAYLC-PKEECLEEARR-RKRLQKAL-------RC-Q-VPDAVLTTL-NER----L-SASTG---VSAEAN----
gb|MBL6801298.1|     ------------------M----GRSAYLC-PQESCLEEAQR-RKRLQKAL-------RC-Q-VPDSVMATL-KQR----L-FPDKE---TVAEAR----
ref|WP_082303561.1|  -------------G----M----GRSAYIC-KLKKCYSDSKI-KKKLQKAL-------KT-P-LEPEFIDIF-EKE----M-TSYND---YPN-------
gb|MCX5959459.1|     ------------------M----GRSAYLC-PSPTCLDEARR-RKRLQRAL-------RC-Q-VSDSIFATL-AAR----L-P-----------------
gb|QNI89774.1|       ------------------M----GRSAYLC-PTEACFEEARR-RKRLQKSL-------RC-Q-VSDDLLTAL-QER----L-TEPRV---AAAEAR----
gb|QNJ32650.1|       ------------------M----GRSAYLC-PTEACFEEARR-RKRLQKSL-------RC-Q-VSEGLMTAL-KER----L-TEPRV---AAAEAR----
tpg|HAS93740.1|      ---PNS------TT----F----GRSVYLC-YNKACIETAFK-KNKIGKHL-------KA-T-IPNELKGQL-LDE----L-RNS---------------
gb|MBE7709406.1|     ---GGP------KI----F----GRSAYLC-YNNSCIENALK-KNKLQKAL-------KV-P-VTQELKGKL-LNE----L-------------------
tpg|HBH17627.1|      S-----------KY----F----GRSSYLC-YNKECLKDAIK-KKRFQRTF-------KK-E-ISDSTFEHL-ENI----I-NK----------------
gb|MEB3319688.1|     ------------QG----M----GRSAYLC-PCSACLEDARR-RRRVVRAL-------RC-P-VDDAVFSEL-ERR----LGDDSRRPSPP---------

                             210       220       230
                     ....|....|....|....|....|....|....|.
ref|WP_010871325.1|  ------------------------------------
ref|WP_190597250.1|  ------------------------------------
ref|WP_028947476.1|  ------------------------------------
gb|MEB3228520.1|     ------------------------------------
gb|MEB3311284.1|     ------------------------------------
gb|MEB3122126.1|     ------------------------------------
ref|WP_075904775.1|  QPNLG-N-----------------------------
gb|NEQ20425.1|       ---KST------------------------------
gb|UXE61785.1|       V-----------------------------------
ref|WP_293043112.1|  QPN---------------------------------
gb|NEO94735.1|       KTNLG-N-----------------------------
ref|WP_293076746.1|  QPNLG-N-----------------------------
ref|WP_071107114.1|  KPNLG-N-----------------------------
ref|WP_044492698.1|  KPNLG-N-----------------------------
ref|WP_293017770.1|  QPNLG-N-----------------------------
ref|WP_293114070.1|  QPNLG-D-----------------------------
gb|PZV27550.1|       ------------------------------------
ref|WP_070395889.1|  KPNLG-N-----------------------------
gb|NEQ64070.1|       KAESPDKEHPK-------------------------
gb|MEB3190993.1|     V-----------------------------------
gb|MBW4545396.1|     ---RST------------------------------
```

```
gb|MBE9166779.1|      ANKT--------------------------------
gb|NES99769.1|        ------------------------------------
gb|NER21363.1|        KAESPNKEHPK-------------------------
ref|WP_096621468.1|   ------------------------------------
ref|WP_192221722.1|   ---RST------------------------------
gb|NES19959.1|        KAESPNTEHP--------------------------
ref|WP_193875631.1|   ASPPN-SVANPSGAKQV-------------------
gb|NEP56036.1|        KAESPHTKNP--------------------------
gb|NET56888.1|        KAEIPHKTASSINVKYFQ------------------
ref|WP_272066750.1|   ESKKS-P-----------------------------
ref|WP_008308507.1|   ------------------------------------
gb|PSR15992.1|        ------------------------------------
gb|PSN10559.1|        ------------------------------------
gb|MBD0344432.1|      ------------------------------------
gb|NEP16756.1|        ------------------------------------
ref|WP_080808818.1|   ------KGSE----------------RSSP--D---
ref|WP_015954383.1|   ------------------------------------
ref|WP_290221549.1|   ------------------------------------
gb|MBJ7899262.1|      ------------------------------------
ref|WP_094348539.1|   ------TQNQ-I------------------------
gb|TVQ19707.1|        S-----------------------------------
ref|WP_300417529.1|   ------TQNR-I------------------------
ref|WP_110985927.1|   ILKPL-PVSKN-------------------------
ref|WP_114081474.1|   ------TQNQ-I------------------------
gb|MCA1994153.1|      LENDK-G-----------------------------
ref|WP_190490988.1|   SDATHVKQSLKSK-----------------------
ref|WP_272127684.1|   QSSSTQTPTQDTD-----------------------
ref|WP_015183318.1|   IEP---------------------------------
ref|WP_190712634.1|   ------------------------------------
gb|NJM61055.1|        PSDLQTPELSYRE-----------------------
gb|MDJ0704997.1|      ------------------------------------
ref|WP_292849343.1|   ------TQNQNQI-----------------------
ref|WP_169155198.1|   ------------------------------------
tpg|HSM83622.1|       SPSP--KETIAQIDRPTKNFCK--------------
gb|MEB3274626.1|      ------------------------------------
ref|WP_324983291.1|   ------TQNE--I-----------------------
ref|WP_292866613.1|   ------TQNQ-I------------------------
ref|WP_022606905.1|   AAER--------------------------------
ref|WP_292789808.1|   ------TQNQ-I------------------------
ref|WP_096570286.1|   ------------------------------------
ref|WP_012595979.1|   ------------------------------------
ref|WP_324629747.1|   --PHPQ------------------------------
ref|WP_190731269.1|   ------TQNQ-I------------------------
tpg|HAA30330.1|       SDQK---REGGV------------------------
ref|WP_298906330.1|   ------T-----------------------------
dbj|BAZ68765.1|       ------LQN---------------------------
gb|MBW4433734.1|      ------LQN---------------------------
ref|WP_181927846.1|   ------TQNQNQI-----------------------
ref|WP_012412015.1|   ------TQNQ-I------------------------
ref|WP_322677349.1|   ------TQNQ-I------------------------
ref|WP_015139046.1|   ------PQNQNLG-----------------------
ref|WP_104905836.1|   ------TQNQ-I------------------------
ref|WP_206814010.1|   ------------------------------------
```

```
gb|MDJ0601942.1|      ------------------------------------
ref|WP_292717392.1|   ------TQNQ-I------------------------
ref|WP_179066568.1|   ------TQNQ-I------------------------
ref|WP_322722179.1|   ------TQNQ-I------------------------
ref|WP_185581622.1|   ------TQNQ-I------------------------
ref|WP_292822895.1|   ------TQNK-I------------------------
ref|WP_196512348.1|   ------TQNQ-I------------------------
gb|OKH54711.1|        ------------------------------------
ref|WP_292785937.1|   ------TQNQ-I------------------------
gb|TAE56869.1|        ------------------------------------
gb|PZV08645.1|        Q-----------------------------------
ref|WP_292751094.1|   ------TQNK-I------------------------
ref|WP_073597769.1|   -----------SGREDI-------------------
ref|WP_073595002.1|   SKID--------------------------------
gb|MCG8363753.1|      ------------------------------------
gb|PSO56092.1|        STERQQ------------------------------
gb|MBF2088208.1|      S-----------------------------------
gb|NES80898.1|        ------------------------------------
gb|PSO96923.1|        STER--------------------------------
dbj|BAZ39493.1|       SEEVN-SSNATEIST---------------------
ref|WP_322731565.1|   ------TQNP-I------------------------
ref|WP_193978934.1|   SAIDDGY--------WDKWTRRPGDKETREQQEN--
ref|WP_300507896.1|   ------------------------------------
gb|TVQ45794.1|        ------------------------------------
gb|MBW4655594.1|      -----------NVRSDEESLPLPSQRPQA-------
ref|WP_204152300.1|   S-----------------------------------
ref|WP_292734233.1|   ------TQNQ-I------------------------
gb|PSN19469.1|        GDSPRMARSS--------------------------
ref|WP_094331183.1|   ------TQNQ-I------------------------
gb|MDW8178835.1|      -DAQSGPQCPESGGRA--------------------
ref|WP_223049825.1|   QAFKGTRRQGADLTPAN-------------------
dbj|BDA66563.1|       ------------------------------------
ref|WP_194027457.1|   ---PDN------------------------------
gb|NEO98984.1|        DLIKVKSLQAQPKQVKASNR----------------
gb|MDZ8004527.1|      ------TQNQ-IY-----------------------
gb|KAB8333374.1|      ------------------------------------
gb|MCY7392750.1|      EV----------------------------------
gb|MBE9078818.1|      NGVSAGAAAPQP------------------------
ref|WP_322669753.1|   ------TQNQ-I------------------------
ref|WP_100898624.1|   ------TQNQ-I------------------------
ref|WP_229551764.1|   ------TQNQ-I------------------------
gb|MBD0304031.1|      ------TQNQ-I------------------------
ref|WP_169218276.1|   ------SQKQT-------------------------
gb|NEO32345.1|        EHLRVKSLKAQPKQVNPSNS----------------
ref|WP_190814021.1|   --------SSPQGRSLPSASDLK-QPPE--------
ref|WP_196527153.1|   ------TQNQ-I------------------------
ref|WP_323196799.1|   ------TQNQN-------------------------
gb|PSO55897.1|        STERQQ------------------------------
ref|WP_229482756.1|   ------TQNQ-I------------------------
ref|WP_292709822.1|   ------TQNQ-I------------------------
ref|WP_069073458.1|   ------TQNENQI-----------------------
dbj|BAU66516.1|       -------KCHN-------------------------
ref|WP_229454557.1|   ------TQNQ-I------------------------
```

```
ref|WP_322697626.1|   ------------------------------------
ref|WP_273766710.1|   ------T-----------------------------
gb|MBE7381418.1|      -----GGAD---------------------------
gb|RCJ24275.1|        ------TQNQ-I------------------------
ref|WP_015187990.1|   PN--------------------------KN------
gb|NET06949.1|        EHLRVKSLKAQPKQVNPSNS----------------
ref|WP_224409244.1|   GHSSDRAS----------------------------
ref|WP_155748928.1|   ------------------------------------
ref|WP_229547107.1|   ------TQNK-I------------------------
gb|MDW8401950.1|      -DAQSGPQCPESGGRA--------------------
ref|WP_107668609.1|   ------------------------------------
ref|WP_008232964.1|   ------DDTSS-------------------------
gb|MEC4893587.1|      GHSSDRAS----------------------------
ref|WP_109009916.1|   ------TQNQ-I------------------------
gb|MEC4816215.1|      ------------------------------------
gb|MBW4486969.1|      VNGLK--LNRALE-----------------------
tpg|HZG41009.1|       ------------------------------------
ref|WP_322643731.1|   ------TQNQ-I------------------------
ref|WP_118168122.1|   ------TQNQ-I------------------------
ref|WP_194042652.1|   ------TQNQ-I------------------------
ref|WP_086688430.1|   ------TQNQILV-----------------------
gb|MEB3161329.1|      ------------------------------------
gb|MBF2099090.1|      ------------------------------------
gb|MBE8968595.1|      ------TQNQ-I------------------------
ref|WP_152589415.1|   ------TQNQ-I------------------------
gb|MCT7960255.1|      ------------------------------------
gb|MBW4576792.1|      PP----------------------------------
ref|WP_233748074.1|   ----DSES----------------------------
gb|MDJ0730709.1|      ------------------------------------
gb|NEO25536.1|        ------------------------------------
ref|WP_190559635.1|   ------PQNQI-------------------------
ref|WP_190600959.1|   ------PQT---------------------------
ref|WP_289796853.1|   ------TQNQ-I------------------------
ref|WP_194057957.1|   ------------------------------------
ref|WP_016866716.1|   ------------------------------------
gb|MCS6814375.1|      SS--V--AEQTQS-----------------------
tpg|HBB35302.1|       ------------------------------------
gb|MDJ0715684.1|      ------------------------------------
gb|NEP12133.1|        KAESPCTEHNQSKLNTFGEHK---------------
gb|NJP17789.1|        -----------DNRSDPSLKT---------------
gb|KAB8319469.1|      ------------------------------------
ref|WP_190413342.1|   --------SSPQERCLPSASDLK-QPSE--------
gb|MBR8834127.1|      ------------------------------------
gb|MCC5607216.1|      ------TQNQ-I------------------------
ref|WP_168729992.1|   ------NHKQV-------------------------
ref|WP_280654349.1|   ------PQNQN-------------------------
gb|MBD2016732.1|      I-----------------------------------
ref|WP_099069649.1|   ------SQNQQI------------------------
ref|WP_190533136.1|   --------SSPQERCLPSASDLK-QPSE--------
gb|MCU0541201.1|      PSDLQTPELTYRELPIAPAPKPQSDLPTSLKGFVSD
ref|WP_193872423.1|   ----DSQSAS----------------GSKIPPQK--
ref|WP_061545594.1|   ------NHKQG-------------------------
ref|WP_102179423.1|   ------------------------------------
```

```
tpg|HLP90034.1|         ------------------------------------
ref|WP_190956491.1|     ------TQNQ-I------------------------
ref|WP_102175048.1|     ------------------------------------
gb|NJO42002.1|          ------------------------------------
ref|WP_006275620.1|     ------NHNQG-------------------------
gb|MBU7582629.1|        -------QNQISG-----------------------
gb|MCG6133583.1|        ------TPNQN-------------------------
ref|WP_016872536.1|     ------YQNQIEF-----------------------
ref|WP_015117518.1|     ------------------------------------
tpg|HBE20201.1|         ------------------------------------
gb|MBW4500045.1|        ------------------------------------
ref|WP_103135735.1|     Q-------KQI-------------------------
ref|WP_105221837.1|     PN--------------------------KN------
ref|WP_190434246.1|     VDGLK--PDRALE-----------------------
ref|WP_283760063.1|     ------------------------------------
gb|MDZ8167634.1|        ------TQNQ-I------------------------
ref|WP_193879029.1|     -------QNQISG-----------------------
ref|WP_190591144.1|     -------QNQISG-----------------------
gb|MDY6781740.1|        N-----------------------------------
gb|MDZ8106853.1|        ------TQNQ-I------------------------
ref|WP_013325296.1|     ------------------------------------
ref|WP_171573341.1|     ---K--------------------------------
ref|WP_190938724.1|     ------TQNQI-------------------------
ref|WP_086768880.1|     ------NQNQILV-----------------------
ref|WP_251960204.1|     ------TQNQI-------------------------
ref|WP_009545653.1|     ------------------------------------
ref|WP_127085004.1|     ------------------------------------
tpg|HIK08066.1|         Q-------KQI-------------------------
gb|MDZ7979263.1|        ------NQNQILV-----------------------
gb|MCC5635819.1|        ------------------------------------
ref|WP_239731788.1|     ------PQNQN-------------------------
ref|WP_193897322.1|     -------QNQISG-----------------------
ref|WP_187706878.1|     ------NHNQV-------------------------
ref|WP_214438763.1|     ------PQNQV-------------------------
ref|WP_073632737.1|     ------------------------------------
gb|MDZ8241182.1|        ------TQNQ-T------------------------
ref|WP_073548075.1|     PN--------------------------KK------
gb|MBD2163199.1|        ------TQNQILI-----------------------
gb|MBO1346139.1|        TTHEMQR-ATRTTQETLAAGPGDLSPKQPDCSRSTV
gb|MBW4600386.1|        ------------------------------------
gb|MBC6453437.1|        ---------TRTTQETPAAGPGDLSPKQPDCR----
ref|WP_289789294.1|     ------FQNQIEF-----------------------
ref|WP_167722165.1|     ------TQNQILILRNL-------------------
ref|WP_095719821.1|     NNS-----------------------HTNT------
ref|WP_096644639.1|     SNSS--------------------------------
gb|NJM70351.1|          ------SQKQT-------------------------
ref|WP_082127362.1|     ------------------------------------
ref|WP_073610176.1|     ----DSIAT---------------------------
gb|MDZ8028666.1|        ------TQNQ-T------------------------
gb|MDZ8019070.1|        ------TQNQ-T------------------------
ref|WP_017742689.1|     ------NHSSN-------------------------
ref|WP_045868281.1|     ------TQNQILI-----------------------
ref|WP_208344959.1|     ------------------------------------
```

```
gb|MBW4560223.1|      ------PQNQILI-----------------------
gb|PSB49245.1|        --------GSTVPKSDNSSASTV-------------
ref|WP_038087798.1|   ------------------------------------
ref|WP_190427456.1|   --------SSPQERCLPSASDLK-QPSE--------
ref|WP_261224160.1|   -----------------------------PPPVLSK
gb|ARV61523.1|        ------NQKQT-------------------------
gb|MCU0523589.1|      ---Q-RGKN---------------------------
ref|WP_089091493.1|   ------PPNQN-------------------------
gb|MBF2080108.1|      ------------------------------------
ref|WP_268609846.1|   ------------------------------------
gb|MBC6418410.1|      ARRDPRSGS---------------------------
ref|WP_200324434.1|   -AKNQI------------------------------
ref|WP_015202566.1|   ------------------------------------
ref|WP_194023660.1|   PGQPLTQ-----------------------------
dbj|BBD59214.1|       -------QNQISG-----------------------
ref|WP_096579096.1|   -------QNQISG-----------------------
ref|WP_315785635.1|   ------------------------------------
gb|MDP8965164.1|      ------------------------------------
ref|WP_015194852.1|   -------KCQNK------------------------
ref|WP_006528507.1|   ------------------------------------
gb|MBC6423737.1|      ---------TRT-QETPPAGPGDLSPKQPDCR----
gb|MBW4517440.1|      ------------------------------------
ref|WP_102221303.1|   ------------------------------------
gb|MBE9051640.1|      ------TQNQN-------------------------
gb|MEB3268310.1|      EYANK-P-----------------------------
dbj|BAY27136.1|       SNFSLGNLWR--------------------------
ref|WP_190744460.1|   SNSL--------------------------------
gb|MCS6791070.1|      ------------------------------------
ref|WP_015144681.1|   -----------SGREDI-------------------
tpg|HYW19585.1|       ------PQNQN-------------------------
gb|MCJ8278778.1|      ---------DAQE-----------------------
gb|RMF20653.1|        ------------------------------------
gb|NRB09489.1|        ------EEQSSPLA----------------------
gb|MDJ0736158.1|      ------P-----------------------------
gb|NJL87205.1|        ---A--------------------------------
ref|WP_261207255.1|   ------------------------------------
ref|WP_008274651.1|   ------------------------------------
gb|MDJ0845508.1|      ------------------------------------
ref|WP_062247903.1|   ------------------------------------
gb|MDF5707201.1|      ------TQNQ-I------------------------
ref|WP_190555052.1|   VDGLK--PDRALE-----------------------
gb|TAD81818.1|        ------------------------------------
gb|MBF2035119.1|      PSSQASPPVQGLG-----------------------
gb|MCY7283860.1|      ------------------------------------
ref|WP_106289357.1|   ------------------------------------
ref|WP_200989746.1|   ------SQNQN-------------------------
ref|WP_026079831.1|   VPVVK-S-----------------------------
ref|WP_196521436.1|   ------TQNH-I------------------------
gb|NJK27659.1|        ------------------------------------
ref|WP_297080144.1|   TQKPD-SLGQAR------------------------
gb|NJL10525.1|        ------------------------------------
ref|WP_124976615.1|   ------------------------------------
gb|MBW4673665.1|      ------TQNK-I------------------------
```

```
ref|WP_261198198.1|  ------------------------------------
ref|WP_015128246.1|  ------PPNQI-------------------------
gb|TAF21607.1|       ------------------------------------
ref|WP_009458837.1|  ------------------------------------
ref|WP_190980837.1|  -------QNQISG-----------------------
ref|WP_283761350.1|  ---SHSQPRPKS-EP---------------------
gb|MBU6184880.1|     EATTIQF-----------------------------
ref|WP_193197802.1|  ------------------------------------
ref|WP_015114806.1|  -------QNQIPG-----------------------
ref|WP_011318669.1|  Q-------NQN-------------------------
gb|MBC6477661.1|     ---------KSTTQETPAAGPGDLSPKQPDCR----
gb|NDJ22983.1|       ------------------------------------
gb|RAM50821.1|       ------LQN---------------------------
gb|MCZ0898804.1|     --------GLAGRKTDNSSASTV-------------
gb|MBW4554427.1|     ------TQNQN-------------------------
ref|WP_179047786.1|  ------PQNQI-------------------------
ref|WP_071599862.1|  ------------------------------------
ref|WP_207087020.1|  ------------------------------------
gb|MDZ8042175.1|     ------TQNP-I------------------------
ref|WP_102207255.1|  ------------------------------------
ref|WP_066384474.1|  ------PQNQNLG-----------------------
gb|PSB20742.1|       ------------------------------------
ref|WP_190697022.1|  -------Q----------------------------
gb|MCT7955770.1|     ------------------------------------
ref|WP_254010714.1|  KLSKNTN-----------------------------
gb|NBD15363.1|       ------------------------------------
ref|WP_283755094.1|  KS-LPERES---------------------------
gb|MDJ0534388.1|     ------------------------------------
gb|MBW4482298.1|     --PPPPGPKV--------------------------
gb|MBD3880571.1|     ANLES-LKQAQE------------------------
ref|WP_271731773.1|  ------AQNQN-------------------------
gb|MBW4634383.1|     ------------------------------------
ref|WP_193999244.1|  ------TQNKI-------------------------
gb|NJN72043.1|       ------------------------------------
ref|WP_016950665.1|  ------TQNQI-------------------------
ref|WP_191758834.1|  ------------------------------------
gb|MBD2237840.1|     ------TQNQILI-----------------------
ref|WP_283767879.1|  ---SLTQPRAKS-ES---------------------
ref|WP_172358412.1|  TQELD-SLGQTQ------------------------
gb|MCP6758672.1|     ------LQN---------------------------
ref|WP_015228055.1|  Q-----------------------------------
ref|WP_015210357.1|  ------------------------------------
ref|WP_104545615.1|  SN--------------------------KN------
ref|WP_237991081.1|  Q-------KEI-------------------------
ref|WP_190882003.1|  ------TQNK-I------------------------
gb|MBF2073651.1|     ESRTVEDSSTPAGSTASTVAQQATSESIS-------
ref|WP_323312969.1|  ------------------------------------
gb|MBF2001555.1|     ETRTVEDSSTPAGSTASTVAQQATSESIS-------
ref|WP_193967039.1|  ------------------------------------
ref|WP_190462952.1|  ------PQNQI-------------------------
gb|MBV6625219.1|     ------------------------------------
ref|WP_265262465.1|  VPVVK-S-----------------------------
ref|WP_281485672.1|  ------PQNQV-------------------------
```

```
ref|WP_127053480.1|  ------TQNQI-------------------------
tpg|HAC62310.1|      ------------------------------------
ref|WP_277772165.1|  IHEPD-SLGQTR------------------------
ref|WP_190570712.1|  Q-------NQN-------------------------
tpg|HIK13958.1|      ------------------------------------
ref|WP_169264435.1|  ------------------------------------
ref|WP_106455349.1|  ------------------------------------
ref|WP_044206893.1|  ------------------------------------
ref|WP_071188254.1|  ------TQNQI-------------------------
gb|MEC4803202.1|     ESLKL-------------------------------
ref|WP_015135874.1|  ------------------------------------
gb|NER81563.1|       LADE-AGAD---------------------------
ref|WP_247215999.1|  -GAPPGRQYPESGDHE--------------------
dbj|BAY28476.1|      ------TQNQILI-----------------------
gb|MCL6749841.1|     ------TQNQ-I------------------------
ref|WP_190685510.1|  -------QNQISG-----------------------
gb|MBF2065700.1|     -QPEVSL-----------------------------
ref|WP_010997971.1|  Q-------NQN-------------------------
gb|MBD2409277.1|     ------TQNQ-I------------------------
ref|WP_006196598.1|  ------PQNQN-------------------------
ref|WP_088239620.1|  -KSS--------------------------------
gb|MBP5972342.1|     ------------------------------------
ref|WP_193997393.1|  ------PQNQN-------------------------
tpg|HLO50431.1|      GLLALTPLPGSKELPVAPTPKPRSDPLTSLERFVSD
ref|WP_146296814.1|  --AR--------------------------------
gb|MBW4616862.1|     ------------------------------------
ref|WP_323360326.1|  ------PQNQI-------------------------
ref|WP_254564844.1|  ------------------------------------
gb|MDZ7956785.1|     ------TQNQILI-----------------------
gb|NJK29495.1|       GELLSIADAQIPPA----------------------
ref|WP_190965760.1|  -------QNQISG-----------------------
ref|WP_261235470.1|  ------------------------------------
ref|WP_261891940.1|  ID----------------------------------
ref|WP_190435367.1|  ETSSQEPPLVPGP-----------------------
ref|WP_190465083.1|  SEPDAGQNCTRCHSGFHRGTANPNSAEQQGECQNSK
ref|WP_299492025.1|  TD----------------------------------
gb|MDF2387367.1|     ------PQNQN-------------------------
ref|WP_265235232.1|  --------RSTWRQSDNSSASTV-------------
ref|WP_293148638.1|  --------ESIGHQTDNSSASTV-------------
ref|WP_015151790.1|  ------------------------------------
ref|WP_271764063.1|  ------AQNQN-------------------------
ref|WP_190473724.1|  Q-------NQN-------------------------
gb|MEB3313608.1|     KT----------------------------------
gb|TAG88461.1|       ------------------------------------
gb|MBF2015645.1|     ------------------------------------
gb|KOP28074.1|       ------LQN---------------------------
ref|WP_190516861.1|  ----NSVAGQ----------------QSLPGRQA--
dbj|BAY09723.1|      ------NQNQILV-----------------------
ref|WP_300634909.1|  ------------------------------------
gb|TVQ05455.1|       ------------------------------------
ref|WP_026732821.1|  ------------------------------------
ref|WP_318699068.1|  ---PPSRPKSEP------------------------
ref|WP_015195815.1|  -QPQLLDIT---------------------------
```

```
gb|MEB3178814.1|      AFPNKDH-----------------------------
ref|WP_096832064.1|   ------------------------------------
gb|MBD2774873.1|      ------------------------------------
ref|WP_224341437.1|   ------------------------------------
ref|WP_035987080.1|   ------------------------------------
ref|WP_084783000.1|   TQKPD-SLGQAR------------------------
gb|MBV9387306.1|      ------------------------------------
ref|WP_017652092.1|   ------TQNQI-------------------------
ref|WP_322663056.1|   ------------------------------------
ref|WP_054468962.1|   ------------------------------------
gb|MDJ0660438.1|      ------------------------------------
gb|MDY6802921.1|      KTY---------------------------------
ref|WP_280651647.1|   ------PQNQNLN-----------------------
gb|MBD1868348.1|      ------------------------------------
tpg|HAG85663.1|       K-----------------------------------
ref|WP_071593325.1|   ------------------------------------
gb|TVQ51200.1|        WLG---------------------------------
ref|WP_190487963.1|   ------------------------------------
gb|MEC4984856.1|      ------------------------------------
ref|WP_190426410.1|   ETSSQEPPIVPGP-----------------------
ref|WP_220611023.1|   ------PQNQI-------------------------
ref|WP_190391042.1|   ------------------------------------
ref|WP_212664564.1|   ------------------------------------
gb|NJR53419.1|        GELLSIADAQIPPASRLASSHSLHSSSAPLRNG---
ref|WP_318728151.1|   ---PSSLPKSEP------------------------
gb|TAE00837.1|        ------------------------------------
gb|NJR38792.1|        ------------------------------------
dbj|BAY85099.1|       ------------------------------------
gb|NJK66758.1|        NKSSVSSV----------------------------
ref|WP_169616303.1|   IGSC--ADP---------------------------
dbj|BAZ53121.1|       ------PQNKI-------------------------
ref|WP_144864508.1|   ------------------------------------
gb|MBW4669271.1|      ------PHSSNLTGTT--------------------
ref|WP_193933574.1|   PN--------------------------KN------
gb|MBD2363150.1|      Q-------KQI-------------------------
ref|WP_271941877.1|   ---PPSLPKSEP------------------------
ref|WP_190409163.1|   Q-------NPK-------------------------
ref|WP_198126109.1|   ------PQNKI-------------------------
ref|WP_137906635.1|   I-----------------------------------
ref|WP_275521808.1|   EPFGQTPSRGSEELSAAPAPKPQSDPPTSLKRFVSD
gb|MBW4594787.1|      ------------------------------------
gb|MDJ0678327.1|      IASQQ-RN----------------------------
gb|QSJ17240.1|        ------PQNKI-------------------------
gb|NEO84548.1|        ---YF-------------------------------
gb|OCQ94358.1|        --------YQPNGQEIPNPNRVNSQPPI--------
ref|WP_012168150.1|   ID----------------------------------
gb|PSO47335.1|        --KR--------------------------------
ref|WP_193847003.1|   I-----------------------------------
dbj|BAZ90194.1|       ------NQNQG-------------------------
tpg|HAT12760.1|       ------------------------------------
gb|MCY7384728.1|      --------GSTLPKSDNSSASTV-------------
ref|WP_015226430.1|   S-----------------------------------
ref|WP_236507116.1|   ------------------------------------
```

```
ref|WP_012164444.1|  ------------------------------------
gb|TAF06084.1|       ------PQNQI-------------------------
gb|NJL79274.1|       ------------------------------------
ref|WP_315863139.1|  HD----------------------------------
gb|MBD2100510.1|     VDQGRTGEGLL-------------------------
ref|WP_028082426.1|  I-----------------------------------
gb|MCL6433641.1|     ADHESEQA----------------------------
ref|WP_193925389.1|  ------------------------------------
gb|PSB25882.1|       --------GSTVPKSDNSSASTV-------------
ref|WP_194064403.1|  --------GSTGRKTDKSSVSTV-------------
ref|WP_170189095.1|  ------------------------------------
gb|MBW4686284.1|     ------------------------------------
ref|WP_271796938.1|  L-----------------------------------
ref|WP_071515609.1|  IATGSQPHT---------------------------
gb|TVP65994.1|       IGSC--ADS---------------------------
gb|MBI1240017.1|     ------PQNQNLN-----------------------
ref|WP_252659375.1|  ------------------------------------
gb|EKQ70938.1|       PSKV--------------------------------
gb|NJR17659.1|       NNT-----------------------QTNT------
ref|WP_168492396.1|  IQS---------------------------------
ref|WP_010471496.1|  ------------------------------------
ref|WP_313949248.1|  VDQGRTGEGLL-------------------------
ref|WP_138499475.1|  ------TQNQ-I------------------------
tpg|HLO85610.1|      ------------------------------------
ref|WP_190625876.1|  ----NSTVI---------------------------
ref|WP_250121821.1|  -----TEAVAPHLNSC--------------------
ref|WP_044105921.1|  ------------------------------------
gb|MBW4519521.1|     ----SE------------------------------
dbj|BAZ29108.1|      ------------------------------------
ref|WP_071992767.1|  ------RQNQI-------------------------
ref|WP_013190105.1|  -QENQI------------------------------
ref|WP_015214913.1|  ------------------------------------
gb|MDJ0796006.1|     ------DEQSSSNQPEST------------------
gb|MDY6901569.1|     ------------------------------------
gb|MBW4650502.1|     ------------------------------------
ref|WP_214431970.1|  ------------------------------------
ref|WP_007355211.1|  GPFTQTPSRGSEEFPAEPAPKPRSDPPTSLKRFVSD
ref|WP_163668145.1|  ------------------------------------
gb|MBW4642782.1|     ------PHNQI-------------------------
gb|MCM0590902.1|     ------TQNQI-------------------------
gb|RMF69880.1|       TQRSD-LSG---------------------------
ref|WP_199250663.1|  ------------------------------------
tpg|HIK55694.1|      AVASTSVDS---------------------------
gb|MBD0267699.1|     ------------------------------------
ref|WP_009344612.1|  ------NHNQG-------------------------
ref|WP_190700710.1|  ----NSVAGQ----------------QSLPGRQV--
ref|WP_168646496.1|  ------RQNQI-------------------------
gb|MEB3164071.1|     ------------------------------------
gb|NJL35745.1|       QS----------------------------------
gb|KPQ38479.1|       ------------------------------------
ref|WP_190679736.1|  ------------------------------------
ref|WP_159789286.1|  ------------------------------------
ref|WP_190641945.1|  ------------------------------------
```

```
ref|WP_015083448.1|  ------RQDRI-------------------------
ref|WP_281153046.1|  ------------------------------------
ref|WP_198807873.1|  LPSR--------------------------------
ref|WP_072033681.1|  ------RQNQI-------------------------
tpg|HIK31517.1|      ------------------------------------
ref|WP_293246005.1|  --------ESIGHQTDKSSASTV-------------
gb|MBD1833872.1|     ETSSQEPPLVPGP-----------------------
ref|WP_277865451.1|  ------------------------------------
ref|WP_168505662.1|  I-----------------------------------
gb|NEQ95601.1|       ------------------------------------
gb|MBW4459453.1|     ----SSVVVQ----------------QPLPGRQA--
gb|MDJ0617177.1|     ------DEQSSSNQPEST------------------
gb|MCL1475321.1|     AANTT-PHNQIRSDLESH------------------
ref|WP_272035636.1|  GPFTQTPFRGSEEFPAEPAPKPRSDPPTSLKRFVSD
ref|WP_106367757.1|  ------------------------------------
ref|WP_096670737.1|  ------RQNQI-------------------------
ref|WP_099701299.1|  PN--------------------------KN------
ref|WP_224085747.1|  Q-------NSRYS-----------------------
gb|TAE05943.1|       --------ESIGHQTDNSSASTV-------------
ref|WP_067774615.1|  Q-------NSN-------------------------
ref|WP_015178990.1|  --------GRAGRETDNSSASTV-------------
gb|MCC5897105.1|     Y-----------------------------------
gb|NER36886.1|       ------------------------------------
gb|MEB3252158.1|     ------------------------------------
ref|WP_230402764.1|  -NAQSDPQYRESGGHG--------------------
ref|WP_099532894.1|  -----GNPTPPE------------------------
ref|WP_190534658.1|  -----GNPTPPE------------------------
gb|NJN88730.1|       NIAASNSDLQHCDPSNPL------------------
ref|WP_193975681.1|  --------GSTGRKTDKSSVSTV-------------
ref|WP_053538412.1|  ------RQNRI-------------------------
ref|WP_072620812.1|  PQN---------------------------------
gb|MDD1421071.1|     ------RQDRI-------------------------
ref|WP_271948199.1|  ------------------------------------
gb|NES70347.1|       ------------------------------------
gb|NBO30352.1|       EATTIQL-----------------------------
gb|MBD1828535.1|     --------ASTGRKTDNSSASTV-------------
gb|PZV12745.1|       SKT---------------------------------
gb|PZO41402.1|       SKT---------------------------------
ref|WP_015172566.1|  NEDTTR------------------------------
ref|WP_190754133.1|  ----NSVAAQ----------------QPLPGRQA--
gb|UNU24151.1|       --------GSPGRKTDNGSASTV-------------
gb|MBS3030443.1|     ------HQDRI-------------------------
gb|MBI4782862.1|     ---TDSAITESAKP------------DSSP------
gb|MDJ0724125.1|     ------------------------------------
gb|MBD1815077.1|     --------GRAGRKTDSSSVSTV-------------
gb|EGK88241.1|       --------GRAGRKTDNSSVSTV-------------
ref|WP_072207768.1|  ------------------------------------
gb|MCG9884819.1|     ------------------------------------
ref|WP_190528272.1|  -----GNPTPPE------------------------
ref|WP_011057771.1|  HD----------------------------------
ref|WP_264321713.1|  ------------------------------------
ref|WP_029634746.1|  ------------------------------------
ref|WP_026099823.1|  ------------------------------------
```

```
gb|MBW4568874.1|      ------------------------------------
gb|MDY7022637.1|      ------------------------------------
gb|MBD1213631.1|      I-----------------------------------
ref|WP_071527241.1|   PAP---------------------------------
gb|PZU98241.1|        --SPPPVPKV--------------------------
tpg|HIK45324.1|       DPPPPPTAEEDDR-----------------------
gb|OBQ35981.1|        I-----------------------------------
gb|NCR11955.1|        ------------------------------------
ref|WP_228383259.1|   ------------------------------------
gb|MBW4611665.1|      ------------------------------------
ref|WP_028091624.1|   I-----------------------------------
gb|MEB3290003.1|      PRDHR-VSE---------------------------
gb|MBD0262109.1|      ------------------------------------
ref|WP_035737147.1|   ------------------------------------
ref|WP_024124972.1|   CNHD--------------------------------
gb|OKH17997.1|        ------------------------------------
tpg|HBK98803.1|       ------------------------------------
ref|WP_006512183.1|   ------IAEM--------------------------
ref|WP_072044931.1|   ------------------------------------
ref|WP_023068840.1|   ------------------------------------
gb|MBF2027289.1|      LG----------------------------------
ref|WP_272116663.1|   PQN---------------------------------
ref|WP_193960470.1|   ------------------------------------
gb|RFP56458.1|        -----GNPTPPE------------------------
ref|WP_125732142.1|   ------------------------------------
ref|WP_006621525.1|   ------------------------------------
ref|WP_190386203.1|   ------RQDRI-------------------------
ref|WP_163518983.1|   ------------------------------------
dbj|BAI90404.1|       ------------------------------------
ref|WP_190574353.1|   ------------------------------------
gb|MDJ0508716.1|      ------------------------------------
gb|TVR11480.1|        ------------------------------------
ref|WP_310454240.1|   YD----------------------------------
gb|MBC6434993.1|      ------------------------------------
ref|WP_002798549.1|   ------------------------------------
dbj|BAZ16285.1|       -QKEILV-----------------------------
gb|MBW4493842.1|      ------------------------------------
gb|EKV02414.1|        ------------------------------------
ref|WP_190665968.1|   KLPNQNQIASEPKP------------QTRPPTVL--
ref|WP_089128587.1|   ------------------------------------
ref|WP_293126645.1|   --------GSIGHQTDKSSVSTV-------------
gb|MEB3224903.1|      ------------------------------------
gb|NET30621.1|        ------------------------------------
gb|QSF50481.1|        YD----------------------------------
ref|WP_149818242.1|   YD----------------------------------
ref|WP_263749268.1|   ------------------------------------
ref|WP_073069683.1|   SLSKT-QH----------------------------
ref|WP_271954606.1|   ------------------------------------
tpg|HIK12112.1|       TGGDGKTLSHRP------------------------
gb|MBW4506515.1|      ------PPSFD-------------------------
ref|WP_287684335.1|   ------------------------------------
ref|WP_225875273.1|   ------------------------------------
gb|TRV14446.1|        ------------------------------------
```

```
gb|MBW4477482.1|      ------------------------------------
gb|NJK40366.1|        ------------------------------------
gb|TRU22031.1|        ------------------------------------
ref|WP_002768359.1|   ------------------------------------
gb|MDJ0546172.1|      ------------------------------------
ref|WP_287078697.1|   ------------------------------------
ref|WP_226583963.1|   GANPT-PST---------------------------
gb|MBL1176422.1|      ------VE----------------------------
ref|WP_159293949.1|   ------------------------------------
ref|WP_071776908.1|   -----F------------------------------
ref|WP_190792225.1|   ------------------------------------
ref|WP_271991901.1|   ------------------------------------
ref|WP_110577867.1|   ------------------------------------
ref|WP_286826366.1|   ------------------------------------
gb|NCR41277.1|        ------------------------------------
gb|NCR55568.1|        ------------------------------------
ref|WP_190367604.1|   -------QQLS-------------------------
ref|WP_150976468.1|   ------------------------------------
ref|WP_149986802.1|   ------------------------------------
gb|MDX2096196.1|      ------------------------------------
ref|WP_002738851.1|   ------------------------------------
ref|WP_194012660.1|   --------GSTGRKTDKSSVSTV-------------
ref|WP_072721595.1|   ------------------------------------
ref|WP_108935646.1|   ------------------------------------
ref|WP_272080703.1|   ------------------------------------
gb|OUC16623.1|        ------------------------------------
ref|WP_271926239.1|   ------------------------------------
gb|MBW4467673.1|      HSNLSSSEDWLKDR------------QEQN------
gb|NCQ85689.1|        ------------------------------------
ref|WP_286626612.1|   YD----------------------------------
ref|WP_009782890.1|   ------------------------------------
ref|WP_181496891.1|   HD----------------------------------
gb|AFY59850.1|        PTRQKDGLSQPPDHAGLGHCP---------------
gb|MBC7883588.1|      LSGGPEDDKRQDAKQG-ENL----------------
ref|WP_009631737.1|   ------QVNAQY------------------------
gb|MBW4539957.1|      LAIE--------------------------------
ref|WP_287969625.1|   ------------------------------------
ref|WP_024970683.1|   ------------------------------------
gb|MBX2866220.1|      ------------------------------------
gb|NJK61661.1|        ------------------------------------
ref|WP_310743590.1|   YD----------------------------------
ref|WP_119259560.1|   ------------------------------------
ref|WP_172191971.1|   ----------------------A-------------
ref|WP_322876998.1|   ------------------------------------
ref|WP_310744398.1|   YD----------------------------------
ref|WP_088891762.1|   ------------------------------------
ref|WP_151695992.1|   ------------------------------------
ref|WP_267384513.1|   ------------------------------------
gb|NJO80776.1|        ----AAKERQGMG-----------------------
gb|NEO34688.1|        ------------------------------------
ref|WP_190359885.1|   ------------------------------------
ref|WP_308256896.1|   ------------------------------------
gb|MDS3859689.1|      ------------------------------------
```

```
gb|MBR8831668.1|     ------------------------------------
ref|WP_317267730.1|  ------------------------------------
gb|REJ59451.1|       ------------------------------------
gb|MBP0000506.1|     -HDEPTTRQGHSDDDIRG------------------
ref|WP_297050161.1|  HD----------------------------------
tpg|HAZ43175.1|      KANPT-ASTLSEH-----------------------
gb|MBW4619585.1|     ----DEL-----------------------------
ref|WP_298613741.1|  HD----------------------------------
gb|MCL1465298.1|     AANTT-PQNQITSDLESH------------------
ref|WP_298975829.1|  HD----------------------------------
ref|WP_168571252.1|  AAEES-ISSNDPERESGSLLRSPNP---------RD
ref|WP_287729185.1|  ------------------------------------
ref|WP_190764830.1|  ------------------------------------
gb|NJL20540.1|       ------------------------------------
gb|MDA0866786.1|     C-----------------------------------
gb|MDX2230996.1|     ------------------------------------
gb|MBE9043893.1|     ------------------------------------
tpg|HAO10320.1|      ------------------------------------
ref|WP_317034091.1|  AAIAL-LPQALSPSSPQP------------------
gb|MBW4678534.1|     KLPNQNQIASEPKP------------QTRPPTVL--
gb|MDJ0570890.1|     ------------------------------------
ref|WP_287659929.1|  ------------------------------------
ref|WP_190774570.1|  GAVTLMNDLNR-------------------------
ref|WP_310800080.1|  YD----------------------------------
ref|WP_002801176.1|  ------------------------------------
ref|WP_190450280.1|  QTNL----PD--------------------------
gb|MBR8827281.1|     ------------------------------------
gb|MDC0831746.1|     -QDESTIRQGQANDDIRG------------------
ref|WP_007306607.1|  ------------------------------------
gb|AJD57472.1|       ------------------------------------
ref|WP_079676917.1|  ------------------------------------
ref|WP_082901633.1|  -QDESTIRQGQANDDIRG------------------
ref|WP_190341424.1|  SSNL--------------------------------
ref|WP_149974369.1|  ------------------------------------
ref|WP_019500197.1|  -----T------------------------------
gb|MBV5261761.1|     PP----------------------------------
ref|WP_324706802.1|  ------------------------------------
ref|WP_293094764.1|  ------------------------------------
ref|WP_287453144.1|  I-----------------------------------
ref|WP_083622039.1|  STTCSRSQPL--------------------------
gb|NJP09111.1|       ------VKEV--------------------------
gb|MBW4420472.1|     PTIE--------------------------------
ref|WP_159249505.1|  ------------------------------------
ref|WP_083622686.1|  ------------------------------------
gb|MDX2215915.1|     ------------------------------------
ref|WP_046662795.1|  ------------------------------------
ref|WP_316429393.1|  --------ITAKGSTEKDSAEG--------------
ref|WP_040054730.1|  ------------------------------------
gb|MBF2048868.1|     --------ITAKDSTKRDSAEG--------------
ref|WP_190504524.1|  ------------------------------------
gb|MBW4692942.1|     ------------------------------------
gb|MDR9403844.1|     LGSVG-RSCNRNSRKL--------------------
ref|WP_008203458.1|  ------------------------------------
```

```
ref|WP_190759246.1|  KLPNQNQIASEPKP------------QTRPPTVL--
tpg|HAX74477.1|      GANPT-PENATASTPSD-------------------
gb|MCD8489703.1|     PPRTPPLEI---------------------------
ref|WP_069966942.1|  PPRTPPLEI---------------------------
ref|WP_204141672.1|  SHQGA-TSSSTATC--D-------------------
ref|WP_141724342.1|  PPRTPPLEI---------------------------
ref|WP_011244384.1|  ------------------------------------
gb|NEQ33015.1|       ------------------------------------
gb|MBW4699545.1|     SSGGPVDDQREYARKN--------------------
ref|WP_168535677.1|  IQS---------------------------------
ref|WP_200669078.1|  AAEES-ISSNDSERESGSLLRSPNP---------RD
ref|WP_250683588.1|  HD----------------------------------
ref|WP_024544928.1|  ------------------------------------
gb|MBE9010997.1|     H------PQPELY-----------------------
ref|WP_193855116.1|  ------------------------------------
gb|NJN19948.1|       SNLPD-------------------------------
gb|MDJ0515540.1|     ------------------------------------
gb|MDY7006698.1|     ------------------------------------
ref|WP_190649792.1|  I-----------------------------------
ref|WP_036044569.1|  I-----------------------------------
ref|WP_058999772.1|  ------------------------------------
gb|MDJ0898873.1|     ------------------------------------
gb|MBW4665085.1|     IPH---------------------------------
gb|MDJ0556426.1|     ------------------------------------
gb|MBC8121277.1|     PSGGPVDDKRKSTEQN-QNL----------------
ref|WP_017287872.1|  I-----------------------------------
ref|WP_190376153.1|  -------TQPELY-----------------------
gb|MBC7972109.1|     ------------------------------------
gb|MDY6937449.1|     ------------------------------------
ref|WP_106237396.1|  ------------------------------------
gb|MBD1842922.1|     -------TQPELY-----------------------
ref|WP_323217923.1|  ------------------------------------
gb|NEQ45539.1|       ------------------------------------
gb|MCY6491807.1|     I-----------------------------------
ref|WP_012954106.1|  ------------------------------------
ref|WP_065713394.1|  ------------------------------------
dbj|BAC90592.1|      SSGGPVDDQREHARKN--------------------
ref|WP_012306383.1|  ------------------------------------
gb|MBW4471632.1|     AG----KQVF--------------------------
ref|WP_326498313.1|  I-----------------------------------
ref|WP_071782506.1|  ------------------------------------
tpg|HIK25275.1|      HD----------------------------------
gb|NJK36543.1|       G-----------------------------------
gb|NEP78655.1|       ------------------------------------
gb|NCO76359.1|       ------------------------------------
ref|WP_017293473.1|  ------------------------------------
ref|WP_104387117.1|  ------------------------------------
gb|TAD79778.1|       ------------------------------------
ref|WP_066345584.1|  ------------------------------------
gb|MCL2931732.1|     ------------------------------------
ref|WP_046277840.1|  ------------------------------------
gb|NJK54848.1|       ------------------------------------
gb|MDX2244890.1|     AKNEH-RR----------------------------
```

```
ref|WP_215607325.1|  ------------------------------------
ref|WP_315862757.1|  IASPASLGSQAASSPAAKPGTSPACEQSAPQSRSPK
gb|MCL2923276.1|     ------------------------------------
gb|NJL46250.1|       AQMHGKTTV---------------------------
gb|MBW4534770.1|     ------------------------------------
ref|WP_015165576.1|  ----AEQAK---------------------------
gb|NJN60647.1|       TTNI----MM--------------------------
ref|WP_190497765.1|  PPSHNLPTLG--------------------------
gb|TAF56653.1|       TTNI----MM--------------------------
ref|WP_300873631.1|  HD----------------------------------
gb|MBW4581861.1|     ------------------------------------
ref|WP_193935386.1|  ------------------------------------
ref|WP_124145084.1|  ------------------------------------
gb|NJK99433.1|       DGRS--------------------------------
gb|WRH67481.1|       ------------------------------------
ref|WP_071789028.1|  ------------------------------------
ref|WP_072041293.1|  ----LETGSN----------------RSMT--QS--
gb|MBD1821041.1|     -------TQPELY-----------------------
ref|WP_293079557.1|  ------------------------------------
ref|WP_310422503.1|  ------------------------------------
gb|MDJ0591465.1|     ------------------------------------
ref|WP_023172451.1|  ------------------------------------
ref|WP_036533928.1|  AGKG--------------------------------
ref|WP_162422901.1|  ------------------------------------
dbj|GGA05509.1|      ------------------------------------
ref|WP_066120908.1|  ------------------------------------
gb|UFP97105.1|       SSGGPVDDQREHAREN--------------------
ref|WP_019504603.1|  ------------------------------------
dbj|BAU14687.1|      L------TQPELY-----------------------
gb|MCY7272735.1|     ITSPDLPTA---------------------------
ref|WP_202219931.1|  ------------------------------------
ref|WP_310488256.1|  HISA--------------------------------
gb|MBE9099686.1|     PS----------------------------------
gb|MBU6228775.1|     ------------------------------------
gb|NKB17092.1|       ------------------------------------
gb|NES69150.1|       ------------------------------------
gb|MBW4553259.1|     HPSE-SWQNG--------------------------
gb|PSP19040.1|       ATEQNRRPAPDGED----------------ASLA--
ref|WP_106312640.1|  ------------------------------------
ref|WP_204104579.1|  --SHPP------------------------------
gb|MCU0552418.1|     INPL--------------------------------
gb|MBW4442872.1|     L------TQPELY-----------------------
ref|WP_106254952.1|  LPQPP-TNAQPT------------------------
emb|CAA9550821.1|    ------------------------------------
tpg|HCF29536.1|      -----PRA----------------------------
gb|MCA1903264.1|     ------------------------------------
gb|NEQ23162.1|       ------------------------------------
ref|WP_293133712.1|  ------------------------------------
ref|WP_015157060.1|  ------------------------------------
ref|WP_121971475.1|  ------------------------------------
dbj|BAZ46326.1|      ------------------------------------
gb|MBW4527443.1|     L------TQPELY-----------------------
gb|NEN88813.1|       ------------------------------------
```

```
gb|MDX2273513.1|      ------------------------------------
ref|WP_293062802.1|   ------------------------------------
ref|WP_218081389.1|   TDEISTNL----------------------------
ref|WP_015168983.1|   ------------------------------------
ref|WP_287129737.1|   ------------------------------------
gb|MCL2927522.1|      ------------------------------------
ref|WP_192153424.1|   ------------------------------------
gb|NJK69321.1|        ------------------------------------
gb|MBP0016486.1|      --SHSP------------------------------
gb|PZO20693.1|        K-----------------------------------
gb|MCH9055185.1|      ------------------------------------
ref|WP_099799890.1|   ------------------------------------
gb|MDJ0773676.1|      ------------------------------------
gb|OWY68048.1|        ------------------------------------
gb|MCX7595673.1|      ------------------------------------
gb|NEQ50465.1|        ------------------------------------
gb|NJN30919.1|        ------------------------------------
gb|MDX2254087.1|      ------------------------------------
gb|AFY96277.1|        ------------------------------------
ref|WP_229640858.1|   ------------------------------------
ref|WP_068815920.1|   IASPNLPTN---------------------------
gb|MCY7336460.1|      ------------------------------------
gb|MBS9769718.1|      ------------------------------------
ref|WP_190352545.1|   ------------------------------------
gb|PZV14006.1|        ------------------------------------
gb|MDE5085023.1|      ------------------------------------
ref|WP_293122704.1|   ------------------------------------
ref|WP_319421161.1|   ------------------------------------
ref|WP_295619447.1|   ---L--TD----------------------------
gb|MCL2936457.1|      ------------------------------------
gb|MDE5075942.1|      ------------------------------------
ref|WP_293158219.1|   ------------------------------------
gb|OLP17560.1|        ----TSCS----------------------------
gb|NJM49111.1|        ------------------------------------
ref|WP_244349359.1|   --AP--TEFEISR-----------------------
ref|WP_232432138.1|   ------------------------------------
ref|WP_193989930.1|   ------------------------------------
ref|WP_235280105.1|   --EP--TETVVHIG----------------------
ref|WP_166274652.1|   ------------------------------------
ref|WP_255131202.1|   ---PSGNMNTVP-SL---------------------
ref|WP_264324473.1|   SASAN-LQSGDCGVTAIN------------------
ref|WP_310410468.1|   LKLID-K-----------------------------
ref|WP_259727590.1|   ---PIAASEAR-------------------------
ref|WP_115093230.1|   ------------------------------------
gb|MAH58821.1|        ------------------------------------
ref|WP_259703109.1|   ---PIAASEAR-------------------------
gb|NDC14930.1|        ---TSAGRASEA-R----------------------
ref|WP_309742497.1|   LKLID-N-----------------------------
ref|WP_036487578.1|   QESKN-LKS-DGNLPGI-------------------
gb|NJN76171.1|        YDET--------------------------------
gb|MBJ7364496.1|      ---PSGNMNTVP-SL---------------------
gb|NBV69567.1|        ------------------------------------
gb|NJM97597.1|        ------------------------------------
```

```
gb|MCY7407389.1|      ----EI------------------------------
ref|WP_254933944.1|   RASKA-I-----------------------------
ref|WP_017327771.1|   IGDR--------------------------------
ref|WP_309728133.1|   ------------------------------------
gb|MCY7367819.1|      ------------------------------------
gb|MEC8605584.1|      ------------------------------------
gb|MBW4662466.1|      ---SKGSVVESGK-----------------------
ref|WP_190401439.1|   ------------------------------------
ref|WP_161823453.1|   ----DESGGS----------------RSNS--Q---
gb|MCS6943628.1|      ------------------------------------
ref|WP_255116400.1|   -AGSEAR-----------------------------
gb|AFZ48415.1|        ------------------------------------
gb|NUN65110.1|        ------------------------------------
ref|WP_115022254.1|   ------------------------------------
gb|MCX5962897.1|      ----EI------------------------------
gb|MBM5807801.1|      ------------------------------------
gb|MBV2350088.1|      ------------------------------------
gb|MCG9892687.1|      ------------------------------------
gb|MBJ7492561.1|      ------PSTSPSGAEAR-------------------
emb|CAI8160962.1|     ------------------------------------
gb|PSI02062.1|        ------------------------------------
gb|OON12454.1|        ------------------------------------
gb|NBO27849.1|        ------------------------------------
ref|WP_255008768.1|   AVSEA-R-----------------------------
ref|WP_255098738.1|   ------------------------------------
ref|WP_255096693.1|   ------------------------------------
ref|WP_296444202.1|   ------------------------------------
ref|WP_211167718.1|   ------------------------------------
gb|NMF59328.1|        ------------------------------------
ref|WP_174235275.1|   ------------------------------------
gb|MBM5799360.1|      DACSEAR-----------------------------
ref|WP_142655983.1|   ------------------------------------
gb|MCP4972515.1|      ------------------------------------
ref|WP_043692418.1|   ------------------------------------
ref|WP_011936384.1|   ------------------------------------
gb|MCA6503255.1|      ------------------------------------
ref|WP_131456745.1|   ------------------------------------
gb|MCB4394176.1|      ------------------------------------
gb|MCS5706208.1|      -SGSEAR-----------------------------
gb|MEB3105445.1|      -SGSEAR-----------------------------
gb|MCL1492198.1|      ------------------------------------
ref|WP_255087149.1|   ---SAAASEAR-------------------------
ref|WP_269621837.1|   ------------------------------------
ref|WP_310483930.1|   ---I--IEPI--------------------------
ref|WP_156796781.1|   KASSEAR-----------------------------
gb|MDA0716923.1|      ---PDANLASEA-R----------------------
tpg|HJN36909.1|       ------------------------------------
gb|MCS5691210.1|      -AAAEAR-----------------------------
gb|NBQ18864.1|        ------------------------------------
ref|WP_255613922.1|   ------------------------------------
ref|WP_201324463.1|   ------------------------------------
ref|WP_011125798.1|   ------------------------------------
gb|MBM5803857.1|      EASSEAR-----------------------------
```

```
ref|WP_094591796.1|   -AGAEAR-----------------------------
ref|WP_197212874.1|   -ASVEAR-----------------------------
ref|WP_115120514.1|   ------------------------------------
gb|MBM5785306.1|      ------------------------------------
ref|WP_038023507.1|   ------------------------------------
ref|WP_254938812.1|   AVSEA-R-----------------------------
ref|WP_038553465.1|   ------------------------------------
gb|MDT7945133.1|      --LL--QKPEP-------------------------
gb|MEB3326908.1|      ---PGADLAFEA-R----------------------
ref|WP_130129912.1|   ------------------------------------
ref|WP_038001092.1|   ------------------------------------
ref|WP_258040662.1|   ------------------------------------
gb|PSP26132.1|        ------------------------------------
gb|MEB3304480.1|      GCGSVATLAPPDRR----------------PE----
gb|NDC16240.1|        ----G-G-----------------------------
tpg|HBC43141.1|       ------------------------------------
ref|WP_206339282.1|   ------------------------------------
ref|WP_063406354.1|   ------------------------------------
gb|NJM75400.1|        V------LNPR-------------------------
ref|WP_259734545.1|   ---PGADLAFKA-R----------------------
gb|MBF2058462.1|      ------------------------------------
ref|WP_197151498.1|   ---PGADLAFKA-R----------------------
ref|WP_254928579.1|   ---SGADLASEA-R----------------------
ref|WP_323259706.1|   ------------------------------------
gb|MBD2423618.1|      ---SGADLASEA-R----------------------
gb|MBM5795377.1|      ------------------------------------
ref|WP_281008927.1|   ------------------------------------
gb|MCH2566529.1|      ------------------------------------
gb|MEB3307264.1|      ---IGDAPASEA-R----------------------
tpg|HBH73361.1|       -SGSEAR-----------------------------
ref|WP_255146391.1|   -AASEAR-----------------------------
ref|WP_271253996.1|   ------------------------------------
gb|MEB3234896.1|      DACSEAR-----------------------------
gb|MDA0886513.1|      ---PGADLAFKA-R----------------------
ref|WP_115019618.1|   ------------------------------------
ref|WP_011130895.1|   ------------------------------------
gb|NDG24387.1|        -ADAEAR-----------------------------
gb|MBM5797070.1|      DACTEAR-----------------------------
gb|MCA6523574.1|      ------------------------------------
gb|MCX5946264.1|      ------PTDTPTGSEAR-------------------
gb|MAV11321.1|        ------------------------------------
ref|WP_320673936.1|   ------------------------------------
gb|MCP9817453.1|      -AGAEAR-----------------------------
gb|NJK35889.1|        LQATLEGG-------QELGNSLRLGQDGIG------
gb|NJL99010.1|        A--SPPST----------------------------
ref|WP_193800382.1|   ------------------------------------
ref|WP_063403536.1|   ------------------------------------
gb|RMD73144.1|        ------------------------------------
gb|MCY7333721.1|      ------------------------------------
gb|MDG2329318.1|      ------------------------------------
gb|MCX5941757.1|      ------PKA---------------------------
tpg|HJN33659.1|       ------------------------------------
tpg|HIK19498.1|       --LL--QKPEP-------------------------
```

```
tpg|HCX54226.1|      ------------------------------------
ref|WP_063399518.1|  ------------------------------------
ref|WP_011359406.1|  ------------------------------------
gb|MEB3185113.1|     -SGSEAR-----------------------------
ref|WP_186470267.1|  ------------------------------------
ref|WP_009628060.1|  ------------------------------------
gb|MBE67324.1|       ------------------------------------
ref|WP_272159772.1|  ------------------------------------
ref|WP_063419582.1|  ------------------------------------
gb|MEB3240836.1|     CPSL--------------------------------
gb|NBW63894.1|       -AGAEAR-----------------------------
gb|MBM5788852.1|     -AGAEAR-----------------------------
gb|MEB3262778.1|     -SGSEAR-----------------------------
ref|WP_296365315.1|  ------------------------------------
gb|PZU97808.1|       ------------------------------------
gb|MAN18293.1|       ------------------------------------
ref|WP_186583364.1|  ------------------------------------
gb|MCS5699361.1|     -AAAEAR-----------------------------
ref|WP_186570491.1|  ------------------------------------
ref|WP_186501173.1|  ------------------------------------
gb|MBL6794407.1|     ------------------------------------
gb|MBL6880295.1|     ------------------------------------
gb|MDB4653764.1|     ------------------------------------
gb|RPF82291.1|       ------------------------------------
gb|MCH1457301.1|     ------------------------------------
gb|MED5164851.1|     ------------------------------------
gb|PZO43371.1|       ------------------------------------
ref|WP_028952666.1|  ------------------------------------
gb|MAI96565.1|       ------------------------------------
ref|WP_037988505.1|  ------------------------------------
ref|WP_029553103.1|  ------------------------------------
ref|WP_106502372.1|  ---PGADLASEA-R----------------------
dbj|GDX71743.1|      -AGAEAR-----------------------------
ref|WP_036911320.1|  ------------------------------------
ref|WP_011825091.1|  ------------------------------------
gb|MBL6803472.1|     ------------------------------------
ref|WP_048017238.1|  ---ADQPPASEA-R----------------------
gb|MAV12819.1|       ------------------------------------
ref|WP_115010044.1|  ------------------------------------
ref|WP_006851947.1|  ------------------------------------
gb|NDG75793.1|       ------------------------------------
gb|MEC8441465.1|     ------------------------------------
ref|WP_069789292.1|  ------------------------------------
gb|WRL41094.1|       ------------------------------------
ref|WP_320002208.1|  ------------------------------------
ref|WP_205909688.1|  ------------------------------------
ref|WP_011432759.1|  --LL--QKPEP-------------------------
gb|MDA7432491.1|     ------------------------------------
gb|MEB3202259.1|     ---PGADLAFKA-R----------------------
gb|OIP77665.1|       ------------------------------------
gb|MEB3159801.1|     ------------------------------------
gb|MED5263527.1|     ------------------------------------
ref|WP_036919232.1|  ------------------------------------
```

```
gb|MBM5790812.1|      ------------------------------------
gb|MED5384943.1|      ------------------------------------
ref|WP_199310374.1|   ------------------------------------
gb|MBD2316621.1|      ------------------------------------
ref|WP_038546212.1|   ------------------------------------
ref|WP_322771268.1|   ------------------------------------
gb|WVL01204.1|        ------------------------------------
ref|WP_015218854.1|   ------------------------------------
ref|WP_055075296.1|   ------------------------------------
gb|MEB3276547.1|      DACTEAR-----------------------------
gb|MCB4428669.1|      ------------------------------------
ref|WP_094510206.1|   -AASEAR-----------------------------
gb|MDP6171750.1|      ------------------------------------
ref|WP_223805489.1|   ------------------------------------
ref|WP_011365029.1|   ------------------------------------
ref|WP_214339661.1|   ------------------------------------
ref|WP_255099870.1|   -AASEAR-----------------------------
gb|TYQ31904.1|        ------------------------------------
ref|WP_320666922.1|   ------------------------------------
gb|MEB3297319.1|      ---TGDAPASEA-R----------------------
ref|WP_198953901.1|   ---PGADLASKA-R----------------------
ref|WP_186595138.1|   ------------------------------------
ref|WP_186493755.1|   ------------------------------------
gb|MEC7393122.1|      ------------------------------------
gb|QNJ16179.1|        ------------------------------------
gb|QNI91005.1|        ------------------------------------
gb|NJK59609.1|        GDSPVCNPN---------------------------
ref|WP_259728879.1|   -AASEAR-----------------------------
gb|AUC59923.1|        ------------------------------------
ref|WP_185186942.1|   ------------------------------------
ref|WP_255599781.1|   ------------------------------------
ref|WP_012196167.1|   -ASSEAI-----------------------------
ref|WP_115080914.1|   ------------------------------------
gb|MBR75694.1|        ------------------------------------
gb|RNC90972.1|        ------------------------------------
gb|MCB4389873.1|      ------------------------------------
ref|WP_303534944.1|   ------------------------------------
ref|WP_186495900.1|   ------------------------------------
ref|WP_320676133.1|   ------------------------------------
ref|WP_011933969.1|   ------------------------------------
ref|WP_011127466.1|   ------------------------------------
gb|MCX5948231.1|      -----R------------------------------
gb|MCS6960746.1|      CHQQ-CGHN---------------------------
ref|WP_217901569.1|   ------------------------------------
gb|OYQ64342.1|        ------------------------------------
ref|WP_190398881.1|   ------------------------------------
ref|WP_114989435.1|   ------------------------------------
gb|MCB4407378.1|      ------------------------------------
ref|WP_186479837.1|   ------------------------------------
ref|WP_247910210.1|   ------------------------------------
dbj|GCE64220.1|       YGR---------------------------------
ref|WP_099812864.1|   --SP--PETSAPGASALDKT----------------
gb|MBT65681.1|        ------------------------------------
```

```
gb|MEB3183095.1|      -ATPEAR-----------------------------
ref|WP_254995692.1|   ---PGADLASKA-R----------------------
ref|WP_150884534.1|   ------------------------------------
ref|WP_006043237.1|   ------------------------------------
gb|NDC34872.1|        -ACSEAR-----------------------------
ref|WP_255613750.1|   ------------------------------------
gb|OUT75469.1|        CLSP-VE-----------------------------
gb|NCG15537.1|        ------------------------------------
gb|TGG79053.1|        -RVAEAR-----------------------------
ref|WP_115131857.1|   ------------------------------------
ref|WP_284500670.1|   ------------------------------------
ref|WP_255616060.1|   ------------------------------------
gb|RCL52291.1|        SPSH-AE-----------------------------
tpg|HAN46371.1|       AKEPSAVLSLDLETVAFTPDPSSRTEVES-------
ref|WP_186587596.1|   ------------------------------------
ref|WP_011431137.1|   --SP--PETSAPGASALDKT----------------
ref|WP_186498867.1|   ------------------------------------
ref|WP_259720650.1|   TTSEA-R-----------------------------
gb|MDP6195789.1|      ------------------------------------
gb|RZO05221.1|        ------------------------------------
ref|WP_269603352.1|   ------------------------------------
ref|WP_036900583.1|   -SNSEER-----------------------------
ref|WP_048347195.1|   ------------------------------------
ref|WP_197156911.1|   TTSEA-R-----------------------------
gb|MCX5969416.1|      -----R------------------------------
ref|WP_259735584.1|   AASEAR------------------------------
emb|CAK6691010.1|     TTSEA-R-----------------------------
ref|WP_067095672.1|   ------------------------------------
ref|WP_269608866.1|   ------------------------------------
gb|MBU6251331.1|      -ACSEAR-----------------------------
ref|WP_115125622.1|   ------------------------------------
gb|MAB54535.1|        ------------------------------------
gb|MBM5813821.1|      ---PGADLASKA-R----------------------
ref|WP_114994692.1|   ------------------------------------
gb|MEB3351632.1|      -SASEAD-----------------------------
ref|WP_186538343.1|   ------------------------------------
ref|WP_038654169.1|   ------------------------------------
ref|WP_011620279.1|   ------------------------------------
ref|WP_186589231.1|   ------------------------------------
ref|WP_074162604.1|   ------------------------------------
gb|MAD68593.1|        ------------------------------------
ref|WP_110861757.1|   ------------------------------------
ref|WP_322782688.1|   AASEAR------------------------------
ref|WP_197162564.1|   AASEAR------------------------------
gb|MAF41027.1|        ------------------------------------
ref|WP_186523834.1|   ------------------------------------
gb|PZO47380.1|        ------------------------------------
gb|MEB3350462.1|      ---THPAPVSEA-N----------------------
ref|WP_254968188.1|   ------AVAPRS------------------------
ref|WP_011295369.1|   ------------------------------------
ref|WP_029626136.1|   QALRQ-DK----------------------------
gb|MAK15572.1|        ------------------------------------
ref|WP_186490350.1|   ------------------------------------
```

```
gb|MEB3169508.1|      ---AATSACVEA-R----------------------
ref|WP_255141823.1|   AASEAR------------------------------
ref|WP_036905622.1|   ------------------------------------
gb|EAQ76225.1|        AASEAR------------------------------
ref|WP_011824427.1|   ------------------------------------
gb|MDP7995305.1|      ------------------------------------
gb|MCX5957055.1|      ---TAGIASPEA-R----------------------
ref|WP_037979888.1|   AASEAR------------------------------
ref|WP_106220260.1|   ACGPGALSAPPYGD----------------LSE---
ref|WP_006854377.1|   ------------------------------------
ref|WP_228007088.1|   -ASSEAR-----------------------------
ref|WP_271488669.1|   ------------------------------------
ref|WP_186516396.1|   ------------------------------------
ref|WP_322775930.1|   AASEAR------------------------------
ref|WP_257473614.1|   ------------------------------------
gb|QNI69741.1|        -ASSEAR-----------------------------
gb|MDA7433454.1|      ------------------------------------
ref|WP_269623649.1|   ------------------------------------
gb|MAR07732.1|        ------------------------------------
ref|WP_254954616.1|   VCGPGVLSAPPYGD----------------LSE---
ref|WP_286160945.1|   ACGPGALSAPPYGD----------------LSE---
ref|WP_269611454.1|   ------------------------------------
gb|MEB3173427.1|      DASSEAR-----------------------------
ref|WP_225875770.1|   -ASSEAR-----------------------------
gb|MBD2718795.1|      ACGPGVLSAPPYGD----------------LSE---
ref|WP_254992200.1|   ACGPGALSAPPYGD----------------LSE---
gb|TVS05339.1|        -AAADSPH----------------------------
gb|QNI50382.1|        ------------------------------------
gb|QBE69991.1|        ------------------------------------
gb|MBM5821121.1|      ACGPGVLSAPPYGD----------------LSE---
gb|EAQ68383.1|        ------------------------------------
ref|WP_254963331.1|   VCDPGVLSAPPYGD----------------LS----
ref|WP_186544284.1|   ------------------------------------
gb|MEB3171165.1|      ------------------------------------
gb|MBW4530560.1|      ACGPGALSAPPYGD----------------LSE---
ref|WP_254957545.1|   VCGPGVLSAPPYGD----------------LSE---
gb|EAU74800.1|        ------------------------------------
ref|WP_015110535.1|   ACGPGALSAPPYGD----------------LSE---
gb|QNJ13310.1|        ------------------------------------
ref|WP_254944301.1|   ACGPGVLSAPPYGD----------------LSE---
gb|MBD2549645.1|      ACGPGVLSAPPYGD----------------LSE---
gb|MCT0206289.1|      ACGPGVLSAPPYGD----------------LSE---
ref|WP_159820065.1|   ACGPGALSAPPYGD----------------LSE---
ref|WP_286194360.1|   ACGPGALSAPPYGD----------------LSE---
ref|WP_254980322.1|   ------LAGERR------------------------
ref|WP_158467192.1|   ------------------------------------
gb|QVL53487.1|        R------PTPP-------------------------
gb|KAF0652084.1|      ACGPGALSAPPYGD----------------LSE---
ref|WP_255092431.1|   AASEAR------------------------------
gb|MEB3361199.1|      ---WLSTASADS-KR---------------------
ref|WP_198949670.1|   AASEAR------------------------------
ref|WP_323356953.1|   ACGPGALSAPPYGD----------------LSE---
gb|MEB3266211.1|      ---AHPAPVSEA-D----------------------
```

```
ref|WP_255103146.1|  AASEAR------------------------------
ref|WP_255110457.1|  AASEAR------------------------------
ref|WP_257473432.1|  ------------------------------------
gb|MBM5793042.1|     ------------------------------------
gb|MCX5932016.1|     AASEAR------------------------------
gb|MEB3351395.1|     GCGSEATVAPPDRR----------------PE----
gb|MBE9154035.1|     -ASSEAR-----------------------------
gb|MBM5825515.1|     PVDRR-PPPP--------------------------
gb|MCU0530038.1|     ------AEGPRS------------------------
gb|MEB3156001.1|     GCGPVASVAPARSE----------------T-----
gb|MBM5816260.1|     GCGPEAFVAPARSE----------------T-----
gb|MCF8131934.1|     VQLTGTEA-------PRAPSIGDLHDERRQSPDL--
gb|MDM7936623.1|     AASEAR------------------------------
gb|MEB3260791.1|     -APTEAR-----------------------------
gb|MEB3353798.1|     -VDQR-------------------------------
gb|MEB3334793.1|     GCGPVASVAPPDRR----------------PE----
tpg|HYP03616.1|      ------------------------------------
ref|WP_221629786.1|  AASEAR------------------------------
gb|MBC1260749.1|     AASEAR------------------------------
gb|MEB3176206.1|     CPSL--------------------------------
ref|WP_259738767.1|  AASEAR------------------------------
gb|MEB3195012.1|     GCGSVATVAPPDRR----------------PE----
gb|MEB3257996.1|     GCGSVAIVAPPDRR----------------PE----
gb|MBO6974532.1|     ------------------------------------
gb|MEB3257676.1|     GCGSVATVAPPDRR----------------PE----
gb|MEB3165236.1|     -----R------------------------------
gb|MEB3349663.1|     R------PSPP-------------------------
ref|WP_209040626.1|  ------------------------------------
gb|MCH9714882.1|     Q------TSPP-------------------------
ref|WP_245156268.1|  ------------------------------------
gb|MDX1977370.1|     N-----------------------------------
gb|QNI77496.1|       ------------------------------------
gb|EAU72346.1|       ------------------------------------
gb|MBL6801298.1|     ------------------------------------
ref|WP_082303561.1|  ------------------------------------
gb|MCX5959459.1|     ---TAGIASPEA-R----------------------
gb|QNI89774.1|       ------------------------------------
gb|QNJ32650.1|       ------------------------------------
tpg|HAS93740.1|      ------------------------------------
gb|MBE7709406.1|     ------------------------------------
tpg|HBH17627.1|      ------------------------------------
gb|MEB3319688.1|     ------------------------------------
```
